# The Role of c-Jun Signaling in Cytidine Analog-Induced Cell Death in Melanoma

**DOI:** 10.1101/2025.04.23.650054

**Authors:** Shayne Sensenbach, Han G. Ngo, Sreyashi Ghosh, Prashant Karki, Vahideh Angardi, Mehmet A. Orman

## Abstract

Melanoma stands as an increasingly pressing health concern. Enhanced mitochondrial metabolism has been reported in melanoma cells that survived treatment with traditional therapeutics, including cytidine analogs like gemcitabine (GEM). These findings suggest that chemotherapeutic drugs may play dual roles in promoting both cell survival and cell death, although the underlying mechanisms require further investigation. Herein, we conducted proteomics analysis on GEM-treated melanoma cells and found a drug-induced activation of DNA damage response and apoptosis, along with cell cycle arrest. Additionally, GEM treatment significantly altered protein networks related to mitochondrial ribosomal activity, the electron transport chain, and translation. Furthermore, we reported an upregulation of the JNK/c-Jun network in connection with the apoptotic proteins. Co-treatment with a Jun N-terminal Kinase (JNK) inhibitor, JNK-IN-8 (JNK_i_), significantly increased cell survival, suggesting the involvement of c-Jun signaling in GEM-induced cell death. Additionally, proteomics analysis revealed that JNK_i_ downregulated apoptosis in co-treated cells, highlighting the potential role of the JNK/c-Jun network inhibition in chemotherapeutic tolerance. Collectively, our findings bridge gaps in understanding how melanoma cells respond to cytidine analogs by demonstrating the multifaceted effects of these agents in 1) inducing JNK-mediated apoptotic cell death, and 2) promoting a state of cell cycle inhibition.

## Introduction

Melanoma poses a significant health concern globally, with an estimated 100,640 cases projected for the United States in 2024, resulting in 8,290 deaths [1]. Notably, the incidence of melanoma has been steadily increasing over the past few decades [1]. Despite promising survival rates for early-stage melanoma, advanced metastatic melanoma presents a formidable challenge, with a 5-year survival rate of only 35% [1]. Unfortunately, many metastatic melanoma patients, and cancer patients in general, experience relapse, or recurrence of the disease. Upon studying 451 metastatic melanoma patients between 1994 and 2014, Koolen *et al*. estimated that the recurrence rate of metastatic melanoma is over 40% [2]. Traditional chemotherapeutics, such as dacarbazine, have historically been employed in treating metastatic melanoma [3] but have shown limited efficacy [4], [5], [6], [7]. In recent years, targeted therapeutics for melanoma have emerged and exhibited improved response rates, but these drugs still have their limitations [8], [9]. Enhancing treatment strategies for metastatic melanoma, by improving the efficacy of drugs which are already available and approved, holds significant promise for reducing recurrence rates and improving overall survival.

A critical factor in cancer recurrence and the emergence of drug-resistant mutations is the existence of tolerant cells, such as persisters [10], [11], [12], [13], [14] which transiently adopt a slow-growing and highly drug-tolerant state, contributing to cancer relapse [10], [11], [12], [13], [14] [15], [16]. Cancer cells exhibit significant heterogeneity, with diverse and complex mechanisms enabling their survival under adverse conditions. Drug treatments can further modify these mechanisms and often serve dual roles. While they trigger cell death pathways, they may also induce the formation of drug-tolerant cell subpopulations, possibly through cell-cycle arrest. Unfortunately, understanding how drug treatments alter cancer cell physiology remains challenging due to the intricate genetic diversity of cancer cells.

This study aimed to comprehensively characterize the molecular mechanisms behind melanoma cell death and survival upon chemotherapeutic treatment. Although phenotypic changes, *e.g.*, metabolic alterations, have been identified in drug-tolerant cancer cells, the specific upstream signaling pathways triggering these changes remain unknown. In our previous study, we probed the metabolic effects of traditional chemotherapeutic drugs on A375 melanoma cells, with a focus on gemcitabine (GEM), and observed increased mitochondrial activity in drug-tolerant melanoma cells [15]. GEM is a cytidine analog whose chemical structure closely resembles cytidine, but with two fluorine atoms attached to the 2’ carbon of the sugar group [17]. The primary mechanism of GEM involves its incorporation into DNA during replication, leading to disrupted replication, DNA damage, and eventual cell death [17]. The cytidine analog drugs azacitidine (AZA) and cytarabine (CYT) are similar to GEM in structure and function [18], [19], and these three drugs are used to treat various cancers [17], [18], [19]. In this report, we further examined the effects of GEM, as well as AZA and CYT, on A375 melanoma cells, and expanded our study to include four metastatic melanoma cell lines: RPMI-7951, SH-4, SK-Mel-3, and SK-Mel-24. Throughout our experiments, notable increases in c-Jun expression and phosphorylation were detected in response to GEM treatment. Further experiments involving JNK-IN-8 (JNK_i_), which prevents the activation of c-Jun by Jun N-terminal Kinase (JNK), demonstrated the pivotal role of c-Jun in melanoma cell death following chemotherapeutic treatment. The c-Jun-mediated cell death induced by cytidine analogs appears to be a conserved phenomenon, as evidenced by its observation in both a lung cancer cell line, H1975, and a non-cancerous human cell line, HEK-293. Additionally, our proteomics analyses provided a deeper understanding of how the inhibition of JNK signaling influences cellular responses to chemotherapy, while also shedding light on the complex molecular pathways involved in GEM-induced cell cycle arrest and survival mechanisms. Our comprehensive analysis of melanoma cell responses to cytidine analogs and our discovery of the critical role of c-Jun signaling in this process represent novel contributions to the field.

## Results

### Gemcitabine Activates Protein Networks Associated with the DNA Damage Response, Cell Cycle Arrest, and Apoptosis in Melanoma Cells

To address the limited efficacy of chemotherapeutic drugs in melanoma treatment [4], [5], [6], [7], and to better understand their impact on melanoma cell physiology, we first employed an untargeted proteomics approach. This analysis was conducted on two groups: solvent (DMSO)-treated A375 cells and A375 cells treated with 20 nM GEM – consistent with our previous study [15] – for 24 hours (see **Supplemental Tables 1 and 2** for cell lines and drug concentrations used). We followed established proteomics data processing steps, including data transformation, normalization, imputation, and statistical analysis, as previously described [20]. Subsequently, we performed pathway enrichment analysis on upregulated and downregulated proteins using STRING (see **Supplemental Tables 3 and 4**) [21]. With a standard threshold P value of 0.05, the 24-hour GEM treatment caused significant changes in the expression of over 2000 proteins. To refine and focus our analysis, we chose a P value of 0.01, which brought the number of proteins of interest under 1000 (see **Supplemental Tables 5 and 6**). STRING enabled us to group these proteins into clusters, which provide insights into changes in cellular behavior and signaling pathways by highlighting interactions among upregulated and downregulated proteins (see **Supplemental Tables 3 and 4**).

GEM treatment affected numerous DNA damage response (DDR) and cell cycle regulation-related proteins, resulting in both upregulation and downregulation (**Figure 1A, B**). Many of the downregulated proteins in these groups are positive regulators of cell cycle progression, DNA replication, and mitosis (**Figure 1B**; **Supplemental Tables 4 and 6**), indicating cell cycle arrest due to DNA damage caused by GEM treatment. This may also explain the formation of previously reported dormant cell subpopulations in melanoma [15]. Additionally, GEM treatment led to the downregulation of proteins associated with RNA processing and metabolism, tRNA aminoacylation, ribosome biogenesis and translation initiation, and Golgi body vesicular activity (**Figure 1B**; **Supplemental Tables 4 and 6**). Enrichment analysis confirmed the statistical significance of these downregulations [selected clusters from the Biological Processes enrichment analysis (Gene Ontology) were plotted using STRING; **Figure 1C, D**]. These observed downregulations logically stem from GEM’s role as a DNA-damaging agent, activating the DDR. GEM-induced upregulations of proteins involved in DDR, cell cycle regulation, and apoptosis were observed, as expected (**Figure 1A**; **Supplemental Tables 3 and 5**). These pathways are interconnected and are well-known to be activated by traditional chemotherapeutic drugs, as they elicit cytotoxicity by causing DNA damage, which halts the cell cycle and induces apoptosis [22], [23], [24]. Cytidine analog drugs are known to cause apoptotic cell death [25], [26], [27], and we have already shown this with apoptosis assays in GEM-treated melanoma cells in our previous study [15]. This alignment with established knowledge strengthens the validity of our experimental methods and data analysis strategy. Furthermore, we observed upregulations in proteins associated with mitosis, electron transport chain (ETC), mitochondrial ribosomal activity, protein folding and processing, mRNA splicing, and multivesicular body (MVB) transport (**Figure 1A**; **Supplemental Tables 3 and 5**). These effects were confirmed with statistically significant upregulations in our enrichment analysis for Biological Processes (Gene Ontology) (**Figure 1D**). Cumulatively, these changes in protein expression indicate GEM-induced cell cycle arrest and dormancy, with the activation of DDR and cell cycle regulatory proteins being especially strong indicators [22], [23], [24]. These findings are aligned with our previous study in which we showed increased mitochondrial activity in GEM-tolerant melanoma persisters [15], and with the upregulated and downregulated protein networks, we offer additional insight into the mechanisms behind GEM-induced melanoma cell dormancy.

**Figure 1.**
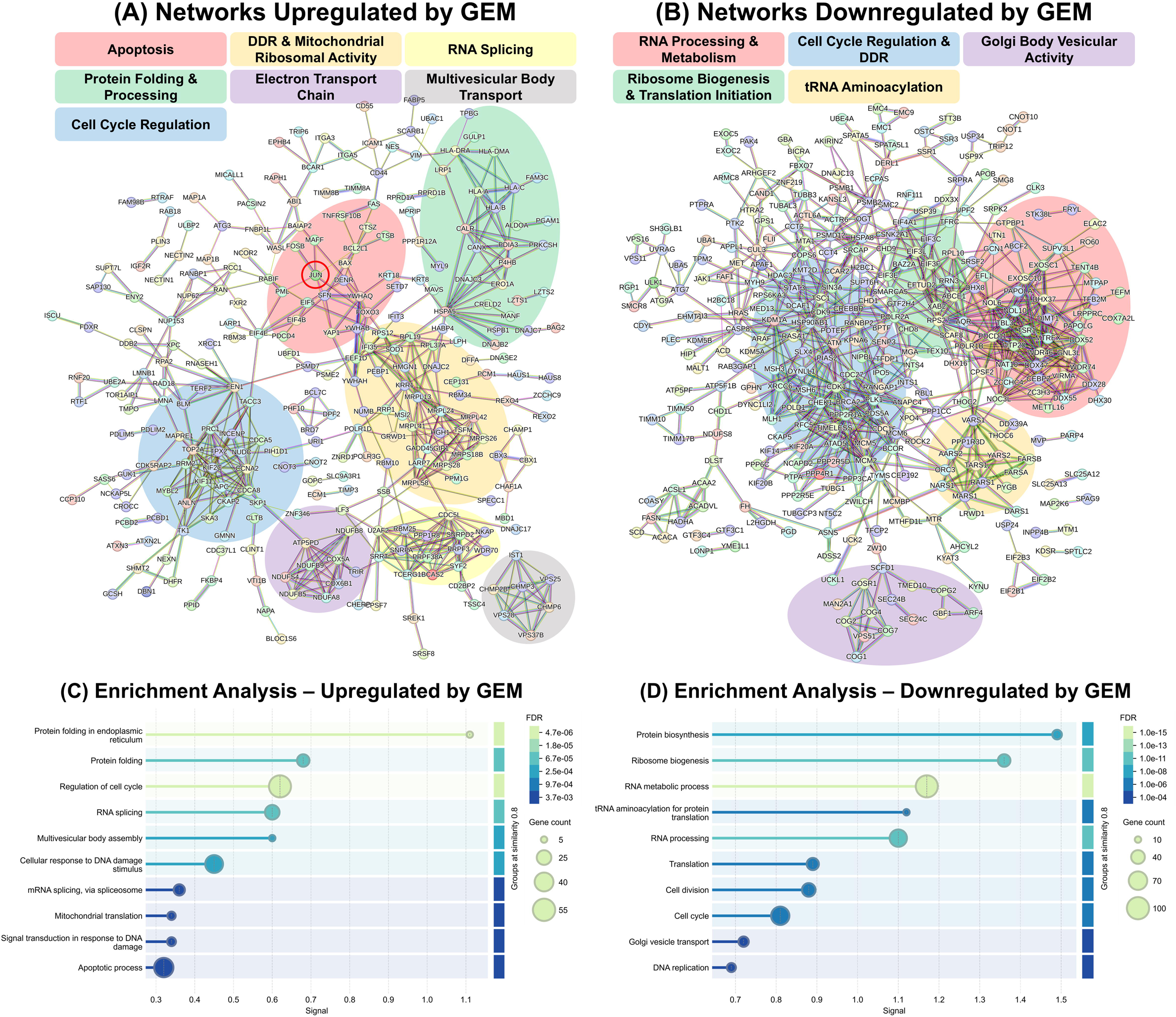
Untargeted proteomics analysis of A375 cells. Changes in protein expression were assessed upon 24 hours of treatment with 20 nM gemcitabine (GEM). (A-B) Protein networks which were significantly (P<0.01) (A) upregulated and (B) downregulated upon GEM treatment. Untreated and treated cells were collected after 24 hours, and LC-MS was used to quantify differences in protein expression between the two groups. STRING was used to depict networks of interacting proteins, indicating altered cellular activities. Significantly changed protein networks are highlighted and described using a color code. The JUN protein (i.e., c-Jun) is highlighted with a red circle in the apoptotic subnetwork. (C-D) Visualization of functional enrichment in the proteins which were (C) upregulated and (D) downregulated by GEM treatment. The protein subnetworks or clusters identified in (A) and (B) were further analyzed through Gene Ontology Biological Process enrichment analysis and visualized in (C) and (D). **Gene count:** the number of proteins related to each term which were identified as upregulated or downregulated. **FDR (False Discovery Rate):** A measure of the significance of enrichment. P values are shown, corrected for multiple testing for each category with the Benjamini-Hochberg procedure. **Signal:** a weighted average between the ratio of observed vs expected proteins and −log(FDR), balancing both these values and allowing intuitive sorting of enrichment values. N = 3.

Of particular interest were the pathways related to cell death and survival. Notably, several proteins involved in apoptosis, such as BAX, BCL2L1, FAS, FOXO3, PDCD4, TNFRSF10B, and YAP1 were upregulated (**Figure 1A**; **Supplemental Tables 3 and 5**). Interestingly, JUN (encoding the c-Jun protein; hereafter referred to as c-Jun) was also upregulated, and interactions between c-Jun and these apoptosis-related proteins were highlighted (**Figure 1A**; **Supplemental Tables 3 and 5**). c-Jun is a transcription factor known for its roles in many complex cellular pathways including cell death, survival, and proliferation [28], [29], [30], [31], [32]. Given its dual role in cancer biology—promoting apoptosis or facilitating cell survival depending on the context—its role in melanoma treated with GEM remains unclear. However, the observed upregulation of apoptotic proteins alongside c-Jun and their interactions (**Figure 1A**) suggest that c-Jun plays a significant role in mediating apoptotic signaling in response to GEM treatment.

### Gemcitabine Activates the JNK Signaling Pathway

To further investigate the signaling pathways affected by GEM, we employed antibody-sandwich-based screening experiments that detect phosphorylated proteins in specific signaling pathways, including MAPK, NF-κB, TGF-β, AKT, and JAK/STAT, to compare GEM-treated cells with control cells (**Figure S1**). As these are high-throughput screening experiments, we focused on observing clear changes, even if they lacked statistical significance (**Figure 2A-E**). We observed significant downregulation of heat shock protein 27 (HSP27, P<0.001) and tumor protein p53 (P<0.001) in the MAPK array (**Figure 2A**), and of the tyrosine kinase Src (P<0.05) in the JAK/STAT array (**Figure 2E**). From the NF-κB array, we observed an upregulation in histone deacetylase 2 (HDAC2, P<0.01) (**Figure 2B**). Most interestingly, the TGF-β array revealed a statistically significant upregulation of c-Jun (P<0.001) and a clear upregulation of activating transcription factor 2 (ATF-2, insignificant) (**Figure 2C**), consistent with our proteomics data. While the upregulation in ATF-2 was statistically insignificant (P=0.0568), it is easily identified in the images used for analysis, similar to c-Jun (**Figure 2F-H; Figure S1**). Both c-Jun and ATF-2 are members of the AP-1 family of transcription factors, and these proteins are known to interact with each other and influence the expression of numerous genes [28]. We corroborated the upregulation of c-Jun through Western blot experiments, showing a significantly stronger signal for c-Jun in GEM-treated samples compared to the DMSO control (**Figure 2I**). Moreover, we observed increased levels of N-terminally-phosphorylated c-Jun (Serine 73). N-terminal phosphorylation activates c-Jun, and dimers of active c-Jun and ATF-2 stimulate the expression of c-Jun itself [33]. Therefore, it is logical to observe concurrent increases in both the expression and phosphorylation of c-Jun. The increased phosphorylation of ATF-2 observed in the immunodetection array experiments (**Figure 2C**) also aligns with these findings.

**Figure 2.**
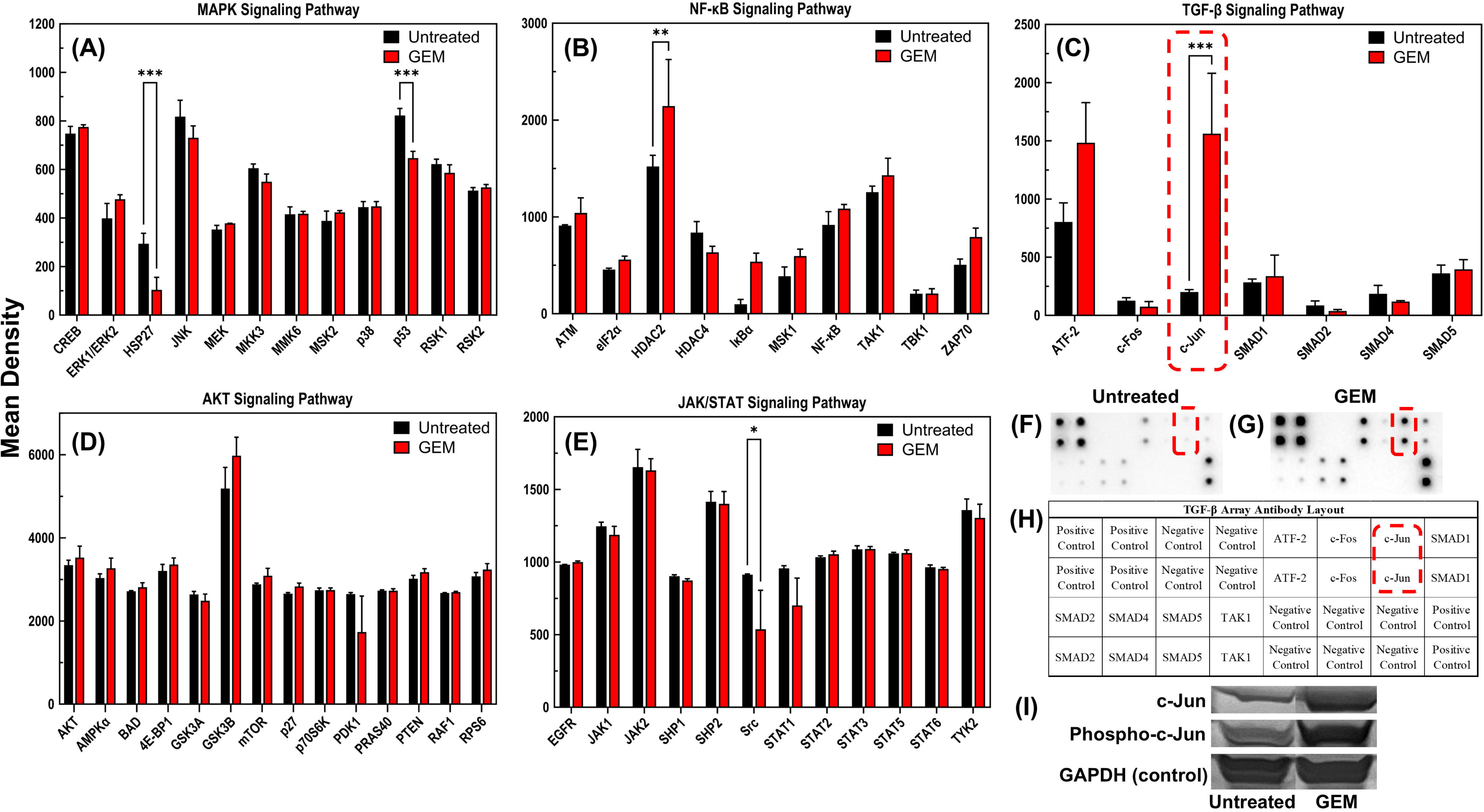
Targeted analysis of protein expression and phosphorylation in A375 cells, comparing untreated cells to those treated with 20 nM gemcitabine (GEM) for 24 hours. (A-E) Quantified levels of A375 protein phosphorylation in the MAPK, NF-κB, TGF-β, AKT, and JAK/STAT signaling pathways. Cell lysates were analyzed using immunodetection arrays which were printed with antibodies of interest. Increased phosphorylation of c-Jun is highlighted in red in (C). For (A-E), statistical significance was assessed using two-way ANOVA with Sidak’s multiple comparisons post hoc testing. (*P<0.05, **P<0.01, ***P<0.001). N = 3. (F-G) Representative images of TGF-β immunodetection arrays which were incubated with lysates from untreated and GEM-treated cells. The detection areas corresponding to c-Jun are highlighted in red. (H) The layout of printed antibodies on the TGF-β immunodetection arrays, corresponding to the images in F and G. (I) Representative images of western blot results, comparing c-Jun protein expression and phosphorylation (at Serine 73) in untreated and GEM-treated A375 cells. The western blot loading control (GAPDH) is shown on the bottom row. N = 4.

Western blot experiments for c-Jun were also conducted on A375 cells treated with other cytidine analogs, AZA and CYT, to determine whether c-Jun activation is specific to GEM. Treatment concentrations were determined using concentration-dependent kill curves (**Figure S2**) (for further explanation, see Materials and Methods). Similarly, we observed upregulations in c-Jun expression and phosphorylation in samples treated with 100 μM AZA and 500 nM CYT compared to the DMSO-treated control (**Figure 3A**). These results further highlight the crucial role of c-Jun in cytidine analog mediated-cell death signaling in melanoma.

**Figure 3.**
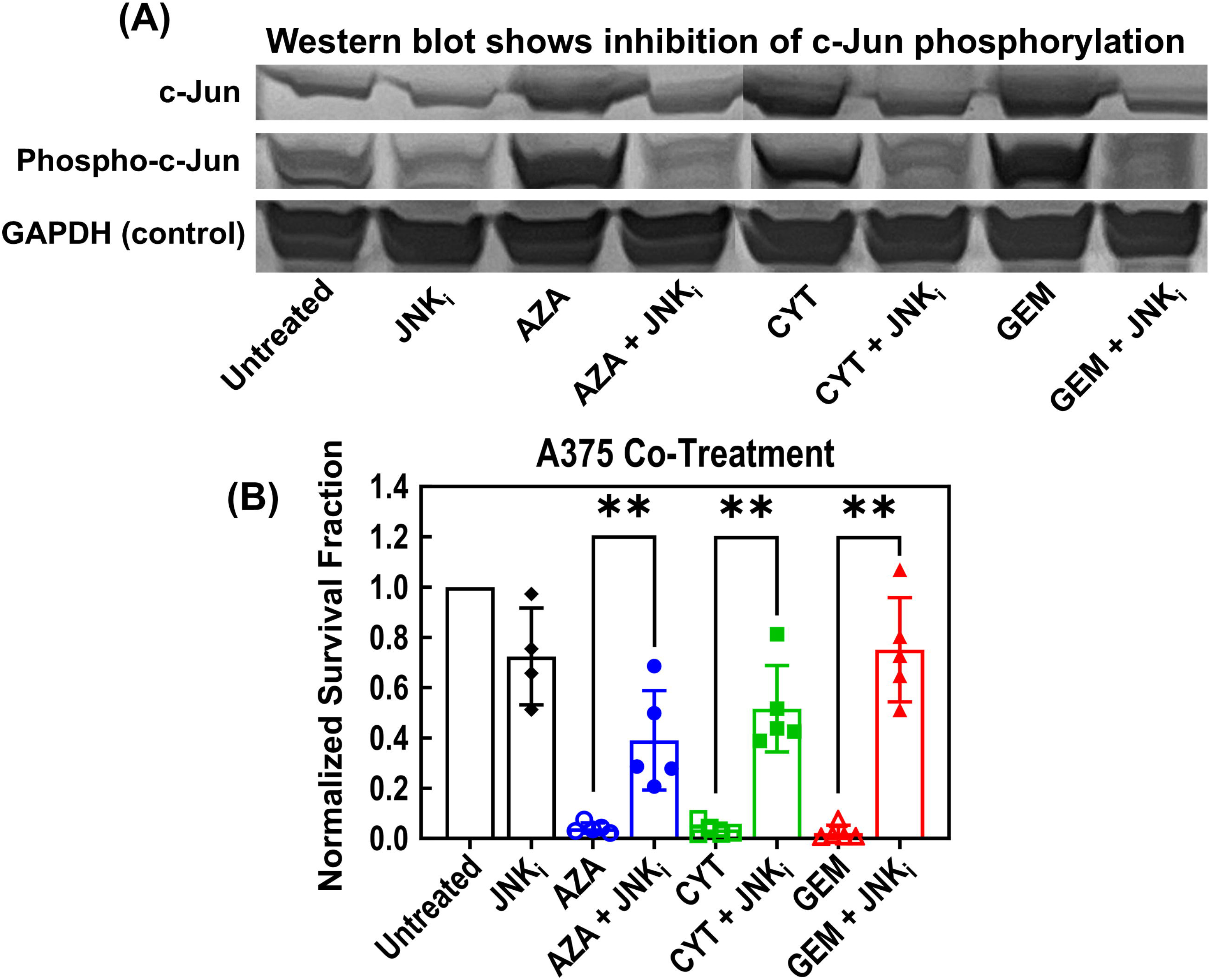
Effects on A375 cells of co-treatment with cytidine analogs, AZA, CYT, and GEM, plus JNK_i_. See Supplemental Table 2 for treatment concentrations. (A) Representative western blot image comparing A375 cell lysates from cells treated under the indicated conditions for 24 hours. The top row shows c-Jun protein levels, the middle row shows phosphorylated (Serine 73) c-Jun levels, and the bottom row shows GAPDH (loading control) protein levels. N = 4. (B) A375 survival fractions were measured with flow cytometry after 72 hours of treatment under the indicated conditions and normalized to the untreated cell number. Pair-wise t-tests were used to determine statistical significance. (**P<0.01). N = 4 (or greater).

### JNK-inhibition Dampens the Toxicity of Cytidine Analogs in Multiple Melanoma Cell Lines

To further explore the role of c-Jun-mediated cell death, we investigated the effects of inhibiting c-Jun. Initially, we utilized siRNA to downregulate c-Jun levels, achieving a 30-50% reduction, but this was insufficient to impact GEM-mediated cell-killing (**Figure S3**). The JNK/c-Jun pathway is a complex signaling cascade within the mitogen-activated protein kinase (MAPK) family, where the phosphorylation of c-Jun by JNK enzymes (JNK-1, JNK-2, and JNK-3) is necessary for its activation [34]. Thus, inhibiting this phosphorylation process collectively may be necessary to observe an impact. To achieve this, we conducted experiments using JNK-IN-8 (JNK_i_), an irreversible inhibitor of JNK-1, JNK-2, and JNK-3 [35]. JNK_i_ is commonly used in cancer research, and it has been shown to synergize with some anticancer drugs [36] while preventing toxicity of others [37] by suppressing JNK signaling. Therefore, JNK_i_ is suitable for these experiments, as we aim to directly target JNK. Concentration-dependent kill curve experiments were used to determine the JNK_i_ treatment concentration; we chose the maximum concentration that did not result in a drastic decrease in cell survival fraction (**Figure S2**).

Next, we analyzed A375 cells treated with a cytidine analog (100 μM AZA, 500 nM CYT, or 20 nM GEM), either alone or in combination with 5 μM JNK_i_, using Western blot. The results were compared to control groups, including a DMSO-treated group and a group treated with JNK_i_ alone (**Figure 3A; Figure S4; Figure S5**). The results showed a strong inhibition of both c-Jun expression and phosphorylation in the co-treated samples compared to the corresponding single treatment samples (**Figure 3A; Figure S4; Figure S5**). While each cytidine analog upregulated c-Jun, co-treatment with JNK_i_ reduced c-Jun expression and phosphorylation, effectively restoring them to untreated baseline levels. Also, single treatment with JNK_i_ led to a decrease in both expression and phosphorylation of c-Jun (**Figure 3A)**. The decreased total protein level is likely due to the decreased activity of c-Jun, which controls its own expression [33].

Co-treatment viability assays were performed on A375 samples using the same treatment conditions. Interestingly, the samples co-treated with each cytidine analog plus JNK_i_ demonstrated significantly higher survival fractions than their corresponding single-treatment samples (**Figure 3B**). These data once again highlight the importance of c-Jun in cell death signaling during cytidine analog treatment. Inhibiting c-Jun phosphorylation, and therefore curtailing its signaling activity, prevents cytidine-analog-induced cell death in A375 melanoma cells.

To gain a broader perspective, we extended these co-treatment experiments to include six additional cell lines. In addition to A375, we tested four metastatic melanoma cell lines: RPMI-7951, SH-4, SK-Mel-3, and SK-Mel-24. *BRAF* V600E mutations are present in each of these melanoma cell lines, along with various mutations in the *CDKN2A*, *PTEN*, and *TP53* genes (see **Supplemental Table 1**). Each melanoma cell line was treated with the same drug combinations as A375. We also tested HEK-293 (human kidney cells, non-cancerous) and H1975 (a lung cancer cell line) under treatment with GEM, as well as GEM plus JNK_i_. Concentration-dependent kill curves were used to determine the treatment concentrations for each cell line (**Figure S2; Figure S6; Supplemental Table 2**). We found that while the melanoma cell lines displayed varying tolerance levels to the cytidine analog drugs (**Figure S2**), co-treatment with JNK_i_ consistently improved the viability of each melanoma cell line when treated with each individual cytidine analog (**Figure 4A-D**). This trend held true across the set of melanoma cell lines, despite varying treatment concentrations due to differences in their tolerance levels. Co-treatment with JNK_i_ also improved viability in HEK-293 and H1975 cells during GEM treatment (**Figure 4E-F; Figure S6**). These findings suggest a potentially conserved mechanism: JNK-mediated apoptosis may be the primary cell death pathway induced by cytidine analogs in many cancer cell types, as well as in non-cancerous (but proliferative) cells

**Figure 4.**
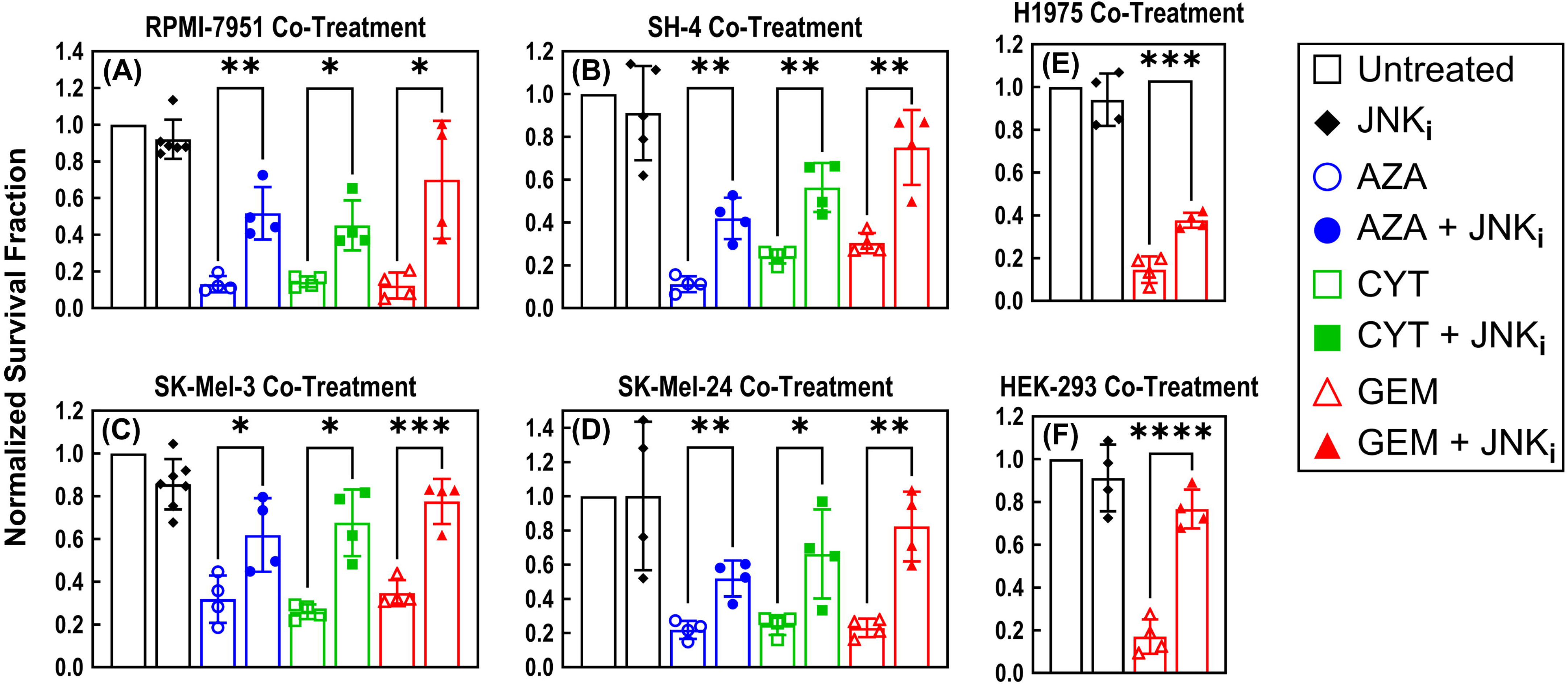
Co-treatment of multiple cell lines with cytidine analogs, AZA, CYT, and GEM, plus JNK_i_. See Supplemental Table 2 for treatment concentrations. (A-F) correspond to co-treatment of RPMI-7951, SH-4, SK-Mel-3, SK-Mel-24, H1975, and HEK-293, respectively. Survival fractions were measured by flow cytometry after 72 hours of treatment under the indicated conditions and normalized to the untreated cell number. Pair-wise t-tests were used to determine statistical significance. (*P<0.05, **P<0.01, ***P<0.001, ****P<0.0001). N = 4 (or greater).

### JNK-inhibition Downregulates Networks Associated with Cell Cycle Arrest and Apoptosis

After consistently observing the rescuing effect of JNK_i_ during cytidine analog treatment across multiple cell lines, we sought to investigate this phenomenon further. Using untargeted proteomics, we analyzed A375 cells treated with GEM, JNK_i_, or their combination, under the previously described concentrations and conditions. Data processing followed the methods outlined earlier, with further details provided in the Materials and Methods section. Analysis of the data revealed that JNK_i_ alone had a less pronounced effect on the cells compared to GEM treatment, as expected. Specifically, utilizing our established threshold of P values less than 0.01, we observed that GEM treatment impacted the expression of 920 proteins (see **Supplemental Tables 5 and 6**), whereas JNK_i_ treatment affected the expression of only 111 proteins (see **Supplemental Tables 9 and 10**). Additionally, the STRING analyses illustrated that the cells exhibited lower reactivity to JNK_i_ (**Figure S7; Supplemental Tables 7 and 8**) compared to GEM (**Figure 1A and B; Supplemental Tables 3 and 4**). These findings align with previous experiments, as JNK_i_ did not drastically alter the viability or growth rates of the melanoma cell lines, unlike the pronounced effects observed with cytidine analog drugs. Furthermore, JNK_i_ treatment resulted in a decrease in the expression of c-Jun, albeit with a P value of 0.0254, which fell just outside the defined threshold for inclusion in the STRING analysis. This observation is logical, considering that JNK_i_ inhibits c-Jun signaling, but this effect is not as severe if c-Jun is not already hyperactive. Consequently, the addition of JNK_i_ was anticipated to exert a more substantial impact on c-Jun expression in GEM-treated cells compared to untreated cells, and this expected effect held true (P value of 0.0003 vs. 0.0254, respectively; **Supplemental Table 14**). Beyond the confirmation of c-Jun downregulation, no noteworthy effects of JNK_i_ treatment emerged from the proteomics dataset. The STRING analysis did not reveal the same complex and interconnected protein networks as observed in the GEM treatment dataset (**Figure S7**, in comparison to **Figure 1A and B**). These results are encouraging, affirming that the treatment with JNK_i_ alone serves as a control experiment, verifying that JNK_i_ does not markedly affect untreated cells while exerting a significant impact on cells treated with cytidine analogs.

The analysis of proteomics data from A375 cells co-treated with JNK_i_ and GEM revealed significant shifts in protein expression (see **Supplemental Tables 11-14**). To uncover these changes, we identified proteins that exhibited substantial differences in expression levels (P values less than 0.01) in the co-treated cells compared to those treated with GEM alone. This approach enabled us to focus on the proteins influenced by the addition of JNK_i_ during GEM treatment. Employing the same data analysis protocol as previously outlined, we detected 298 proteins with significant alterations in expression, with a predominant majority (193 out of 298) exhibiting downregulation (see **Supplemental Tables 13 and 14**). Consistent with this numerical distribution, the analysis of upregulated proteins (due to JNK_i_ co-treatment) in STRING did not show extensive clusters of interacting proteins (**Figure 5A; Supplemental Tables 11 and 12**).

**Figure 5.**
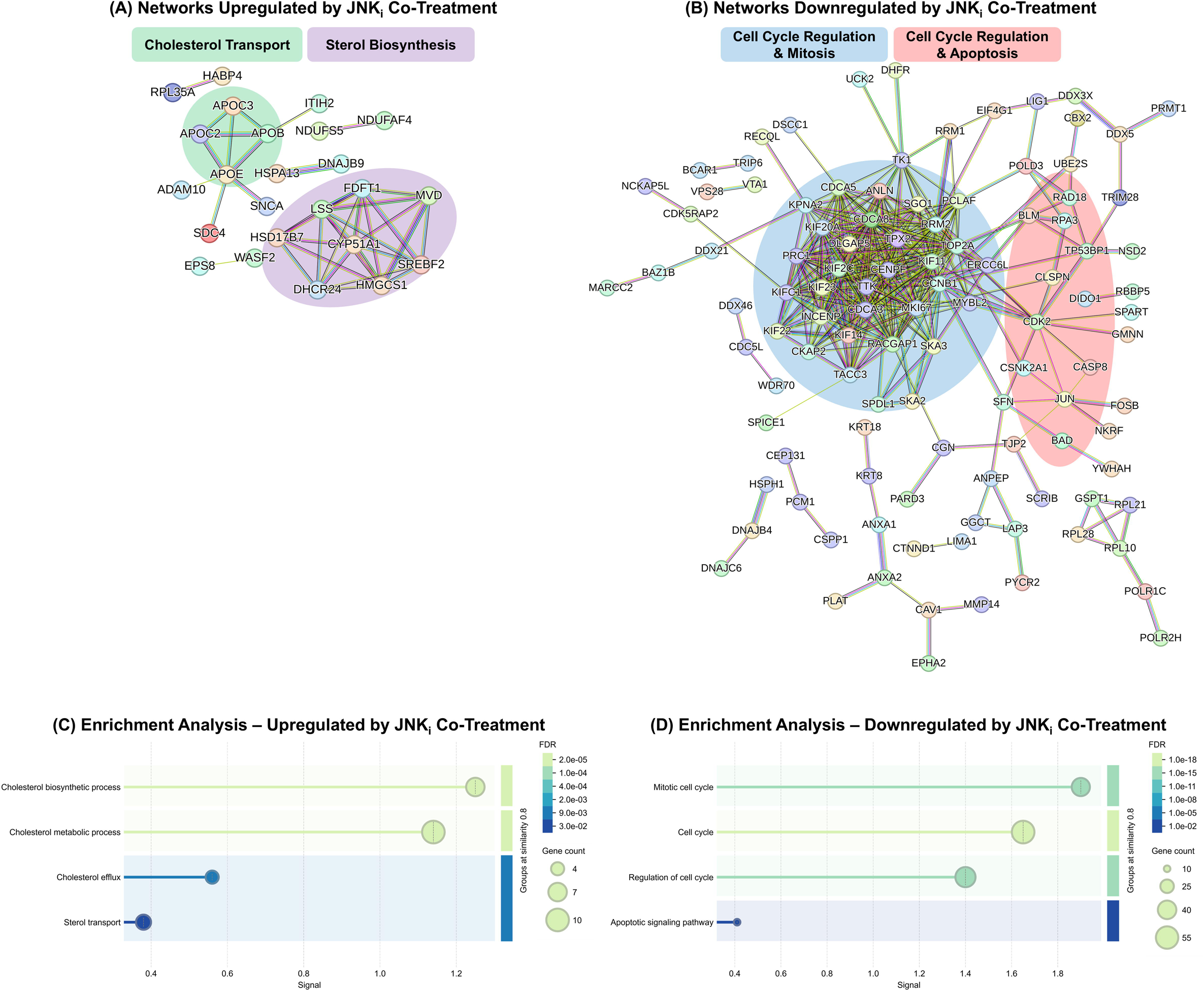
Untargeted proteomics analysis of A375 cells. Changes in protein expression were assessed upon 24 hours of treatment, comparing co-treatment with GEM plus JNK_i_ to single GEM treatment. Treatment concentrations for GEM and JNK_i_ were 20 nM and 5 μM, respectively. (A-B) Significantly (P<0.01) (A) upregulated and (B) downregulated protein networks are presented similar to Figure 1. In this case, upregulated and downregulated protein networks are due to the addition of JNK_i_. (C-D) Visualization of functional enrichment in the proteins which were (C) upregulated and (D) downregulated by JNK_i_ co-treatment. The protein subnetworks or clusters identified in (A) and (B) were further analyzed through Gene Ontology Biological Process enrichment analysis and visualized in (C) and (D). See Figure 1 for descriptions of the terms in these figures. N = 3.

However, both the upregulated and downregulated datasets provided interesting insights into anticipated effects on cell death and proliferation pathways. The upregulated dataset highlighted key changes in sterol biosynthesis and cholesterol transport (**Figure 5A; Supplemental Table 11**). In the downregulated dataset, we identified decreased expression of proteins related to apoptosis, cell cycle regulation, and mitosis (**Figure 5B; Supplemental Table 12**). Statistically significant alterations in these protein networks were observed in our enrichment analysis of the Biological Processes (Gene Ontology) group (**Figure 5C and D**). In the STRING protein map, c-Jun is positioned on the right side of the downregulated protein map, forming connections with multiple apoptosis-related proteins, including BAD, CASP8, and DIDO1 (**Figure 5B**). The large cluster on the middle-left side of the STRING figure contains proteins related to cell cycle regulation and mitosis. Based on these changes, we conclude that c-Jun mediates growth arrest and cell death signals during GEM treatment. Inhibiting c-Jun in co-treatment preserves sterol biosynthesis and transport pathways, promoting survival and proliferation. Altogether, these proteomics findings clearly demonstrate that inhibiting the JNK/c-Jun pathway reduces the apoptotic impact of GEM treatment in co-treated cultures, thereby supporting our previous results.

## Discussion

This study provides insights into how melanoma cells respond to cytidine analog drugs. GEM treatment induced upregulation of apoptosis, inhibition of cell growth, and alterations in mitochondrial metabolism. Although JNK inhibition alone does not significantly affect the global proteomic landscape of melanoma cells, its inhibition in GEM-treated cells markedly attenuated the induction of apoptosis and other proteomic changes, which contributes to enhanced cell survival. The untargeted proteomics experiments generated extensive data, providing a comprehensive view of altered protein levels and pathway enrichments. Due to the volume of these results and analyses (**Supplemental Tables 3-14**), much of the proteomics discussion is included in the supplemental file. Below, we focus on the most significant results.

### c-Jun is a Crucial Apoptotic Signaling Molecule

The mechanisms involved in cell death are critical in the context of cancer therapy. c-Jun has been extensively studied as a critical signaling molecule; it has influence on cell fate in terms of death or survival, as well as proliferation [28], [29], [30], [31], [32]. Both the overexpression of c-Jun and its activation by JNK have been linked to the prevention of apoptosis during chemotherapeutic treatment, suggesting c-Jun as a target for cancer treatment [31]. Also, activation of the JNK pathway has been shown to enhance DNA repair and improve survival of cancer cells during chemotherapeutic treatment [38]. However, studies have also shown the opposite effect, with either the expression or phosphorylation of c-Jun promoting apoptosis, while the inhibition of c-Jun prevents it [37], [39], [40], [41]. The effects of stimulating or suppressing the c-Jun/JNK pathway vary depending on the state of the cells and the environment in which they exist [32]. For melanoma specifically, c-Jun primarily exerts oncogenic effects [42], for example, by enhancing the MAPK and PI3K signaling pathways, promoting proliferation and survival [43]. *PTEN* mutations may also be important in determining the role of c-Jun activation in melanoma cells, as c-Jun can reverse the tumor suppressive effects of *PTEN*. However, in this work, we consistently observed c-Jun activation and expression being strongly associated with apoptosis, while inhibiting c-Jun restored viability and prevented apoptosis. The observed apoptosis-inducing effect of c-Jun activation was consistent across five melanoma cell lines with various mutations, including cells with or without mutations in the *PTEN*, *TP53*, and *CDKN2A* genes (see **Supplemental Table 1**). Therefore, in the context of melanoma cells being treated with cytidine analogs, the primary effect of c-Jun activation is promotion of apoptotic death.

### Cytidine Analog Mediated Apoptosis is Prevented by JNK-inhibition

We demonstrated that the apoptotic death induced by three cytidine analog drugs – AZA, CYT, and GEM – is inhibited by co-treatment with JNK_i_. This phenomenon was observed across five melanoma cell lines, one lung cancer cell line, and a non-cancerous HEK-293 cell line. Therefore, these findings are not specific to melanoma, nor are they specific to cancer in general. However, previous studies have demonstrated that c-Jun can have contradictory effects in different contexts [31], [37], [38], [39], [40], [41]. Interestingly, the importance of JNK activity during GEM treatment has been demonstrated previously using a lung cancer cell line [41]. There have also been previous reports highlighting the importance of the JNK pathway in melanoma cells treated with paclitaxel [44], docetaxel [45], and doxycycline [46], but not those treated with agents which directly target DNA. Interestingly, another study reported that JNK-inhibition led to apoptosis in melanoma cells [47]. The role of c-Jun and the JNK pathway clearly depends on many factors, including the specific drugs used for treatment. In the case of DNA-damaging cytidine analogs, our findings consistently demonstrate that JNK activity plays a pivotal role in apoptotic signaling. This is particularly evident in melanoma cells, as we have observed this phenomenon across five different melanoma cell lines.

These findings confirm the importance of the JNK signaling pathway in cancer treatment, which has been reported for multiple cancer types and multiple types of drug treatments, albeit with various implications in the clinical context. However, we studied combinations of cell lines and drugs which are distinct to the existing literature, meaningfully expanding on current knowledge in the field. Additionally, our extensive proteomics analyses provide connections between the JNK pathway and several other pathways which are important to melanoma chemotherapeutic tolerance. We also offer insight into the mechanisms by which JNK-inhibition prevents apoptosis, which is clinically relevant in the context of chemotherapeutic resistance and tolerance in melanoma.

### Potential Therapeutic Applications

Our extensive proteomics analyses identified pathways activated or inactivated by GEM treatment, as well as those further affected by JNK_i_ co-treatment. While these pathways are discussed in detail in the supplemental file, some may represent potential therapeutic targets. From single GEM treatment, we highlighted upregulations in ETC activity (mitochondrial metabolism), mitochondrial ribosomes, and HSPs. Mitochondrial metabolism and mitochondrial ribosomal activity are directly linked, and these findings are further supported by our previous study [15]. Also, mitochondrial ribosomes have been studied as a target for cancer treatment (see references [48], [49], [50] and supplemental file), so these effects are intriguing in a therapeutic context. Mitochondrial inhibition, with antibiotics targeting bacterial ribosomes, for example, holds promise for eradicating melanoma persisters, thereby enhancing melanoma treatment with traditional chemotherapeutics. Moreover, specific HSPs are known to be important in cancer development and chemotherapeutic response; therefore, the specific HSPs and molecular chaperones which were upregulated serve as potential targets to sensitize melanomas to chemotherapeutics; the proteins we identified include HSPA5 and HSPB1, among others.

Golgi body vesicular transport was downregulated by GEM treatment. Due to its connections with cell death pathways, the Golgi body is also an interesting target. JNK_i_ co-treatment downregulated several apoptotic proteins, including BAD, CASP8, and DIDO1. The inactivation of these apoptotic proteins resulted in a drug-resistant phenotype. Therefore, targeting these proteins may lead to the antagonization of drug resistance mechanisms, thereby increasing the efficacy of chemotherapeutic treatment. However, direct activation of apoptosis may be overly toxic to melanoma patients, so targeting these proteins may lead to complications.

Apolipoprotein expression was upregulated by JNK_i_ co-treatment, and this shift also partially explains the drug-resistant phenotype we observed. Apolipoproteins, as well as the upregulated sterol biosynthesis proteins, are important in cancer cell growth. Further, as the inhibition of apolipoproteins has been used for the treatment of hyperlipidemia [51], these molecules serve as realistic targets to improve the response of melanoma to chemotherapeutic treatments, without exerting additional toxicity on patients.

## Conclusion

Our study addresses gaps in understanding the response of metastatic melanoma to chemotherapy. Using a panel of five melanoma cell lines, we probed the mechanisms behind the death and survival of melanoma cells being treated with three cytidine analog chemotherapeutic drugs. Several screening methods revealed c-Jun as a key molecule in the signaling response to these treatments, and using a JNK-inhibitor, we demonstrated a consistent prevention of apoptotic death by preventing the activation of c-Jun. Proteomics analysis provided a comprehensive description of the effects of single GEM treatment, as well as JNK-inhibition during GEM treatment, revealing multiple potential therapeutic targets for metastatic melanoma. The targets we find particularly intriguing include mitochondrial metabolism (ETC activity), mitochondrial ribosomes, and apolipoprotein expression. These targets should be further investigated in future research focusing on melanoma treatment. Uncovering additional information about the behaviors and effects of the JNK/c-Jun pathway in various cellular contexts, as well as various treatment methods, remains an important point of emphasis for future studies. Due to our findings, we believe studying the effects of JNK inhibitors and similar chemicals will continue to make meaningful contributions to melanoma research. Analyzing the proteomes of melanoma cells, and how they respond to JNK inhibition during various additional chemical treatments, will lead to novel discoveries in the mechanisms behind drug-induced cell death, drug tolerance, and drug resistance.

## Materials and Methods

### Cell Lines and Materials

The following human melanoma cell lines were used:

- A375
- RPMI-7951
- SH-4
- SK-Mel-3
- SK-Mel-24

Note: these melanoma cell lines all have *BRAF* V600E mutations. Also, A375 is a primary tumor cell line, whereas the others were isolated from metastatic tumors. Additional information on each of the cell lines we used can be found in the Supplemental File (see Supplemental Table 1). The non-melanoma cell lines used are below:

- H1975 – a human lung cancer cell line
- HEK-293 – a human embryonic kidney cell line

Cell lines were purchased from ATCC (Manassas, VA). Culture media, materials, antibodies, and chemicals were obtained from Fisher Scientific (Atlanta, GA) and Thermo Fisher Scientific (Waltham, MA). Cell culture flasks and plates were purchased from VWR International (Pittsburgh, PA). Drugs were stored as stock solutions in dimethyl sulfoxide (DMSO) and diluted accordingly for treatment. The volume percentage of DMSO was consistent between treated and untreated control groups.

### Cell Culture Conditions and Techniques

All the above mammalian cell lines except SK-Mel-3 were cultured in Dulbecco’s Modification of Eagle’s Medium (DMEM), supplemented with 10% fetal bovine serum (FBS) and 100 units of penicillin and 100 μg streptomycin per milliliter. The SK-Mel-3 cell line was similarly cultured in McCoy’s 5A Medium with identical FBS and antibiotic supplements. Cells were grown in T-75 flasks with 15 milliliters of complete culture medium, starting with a seeding density of roughly 200,000 cells per milliliter. Actively growing cell cultures were kept in a humidified incubator operating at 37 °C and 5% carbon dioxide. The culture medium was discarded and replaced for each flask every 3-4 days.

Adherent cells were used for this study, so 0.25% trypsin-EDTA (Ethylenediaminetetraacetic acid) solution was used to detach the cells when they reached 80-90% confluence. To detach cells from a T-75 flask, the culture medium was discarded, 5 milliliters of Dulbecco’s Phosphate Buffered Saline (DPBS) solution (without calcium chloride or magnesium chloride) was used to wash the surface of the flask; then, the DPBS was discarded and 2 milliliters of 0.25% trypsin-EDTA solution was added. The cells were then incubated at 37 °C for 2 minutes. Once the cells had visibly detached from the surface, 8 milliliters of DMEM were used to wash the surface and remove the cells. The trypsin-EDTA solution was removed by centrifugation (unless otherwise stated, all centrifugations took place at 800 RPM for 5 minutes), and then the cells were resuspended in DMEM and seeded appropriately. To assess the cell density, 0.4% trypan blue solution was used to stain a cell culture sample (one-to-one volume ratio of cell culture to trypan blue solution), and the cell density and viability were calculated using an Invitrogen Countess II cell counter (catalog # AMQAX1000, Thermo Fisher Scientific, Waltham, MA).

### Cell Collection and Analysis for Proteomics Experiments

A375 melanoma cells were seeded in T-75 flasks, with 2.5 million cells in each flask, and incubated for 24 hours to allow the cells to attach and begin growing. Then, the medium was removed, and replaced with fresh medium, plus either 20 nM GEM, 5 μM JNK_i_, both (co-treatment), or DMSO (control). After 24 hours of treatment, the cells were collected and resuspended in DMEM, and counted with trypan blue staining. Aliquots containing 15 million cells were taken from each sample; they were centrifuged, resuspended in 1 milliliter of DMEM, and chilled on ice for 5 minutes. The chilled suspensions were centrifuged at 4 °C, and the pellets were resuspended in ice-cold DMEM, **without** FBS or antibiotics. This washing step was repeated three times, and finally, the suspensions were centrifuged at 4 °C, the supernatants were removed, and the pellets were transferred to −80 °C storage.

These samples were delivered to the University of Houston Mass Spectrometry Laboratory (UH-MSL), where the proteins were isolated from the cells, processed into peptides, and analyzed using liquid chromatography-mass spectrometry (LC-MS). Proteins were extracted and digested with the Sample Preparation by Easy Extraction and Digestion (SPEED) method, as previously described [52]. Briefly, the cells were centrifuged, and the supernatant was removed. Then, 10 μL of trifluoroacetic acid was added, and the samples were vortexed thoroughly. 100 μL of tris base (2 M concentration) was added, and then a mixture of CAA (40 mM) and TCEP (10 mM) was added. Samples were heated for 5 minutes at 95 °C, and then 500 μL of HPLC grade water was added. Trypsin was added (w/w 1:50) and samples were incubated overnight at 37 °C. Next, the peptides were desalted with C18 ziptips, and a CentriVap (Labconco Corporation, Kansas City, MO) was used to vacuum dry them.

LC-MS was carried out as previously established [53], with modifications. In this case, a NanoElute LC system was used, and it was coupled to a timsTOF Pro (Bruker Daltonics, Germany) with a CaptiveSpray source. 100 ng samples were loaded onto a packed column (75 mm x 15 cm, 1.9 mm ReproSil-Pur C18 particle, from Dr. Maisch GmbH, Germany) at 40 °C. The mobile phases included buffer A and buffer B (0.1% FA in either water or ACN, respectively). The short gradient, from 2% B to 30% B, was 17.8 minutes, with 18.3 minutes to 95% B, and maintained for 2.4 additional minutes. For data acquisition, a data-independent acquisition parallel accumulation-serial fragmentation (diaPASEF) scheme with 16 m/z and ion mobility windows was used. 1.6 kV was used for the electrospray voltage, and 180 °C was used for the ion transfer tube temperature. Full MS scans, over the m/z range of 100-1700, were acquired. The collision energy was linearly ramped as a function of the mobility from 27 eV at 1/K0 = 0.85 Vscm-2 to 45 eV at 1/K0 = 1.3 Vscm-2.

These data were analyzed using the software DIANN version 1.8 [54]. The software’s default settings were used for peptide and protein identification from diaPASEF data. The UniProt-SwissProt Homo Sapiens database (Taxon ID 9606, downloaded on 1/20/2023, 20,404 entries) was used to generate an in-house spectral library. Cysteine carbamidomethylation was included as a fixed modification, with methionine oxidation and acetylation as variable modifications. The false discovery rate (FDR) was controlled at <1% at both peptide and protein levels.

### Protein Isolation for Western Blot and Immunodetection Array Experiments

A375 melanoma cells were cultured and treated similarly to the proteomics cell collection methods. However, in this case, cells were either singularly treated with 100 μM AZA, 500 nM CYT, or 20 nM GEM, or co-treated with each cytidine analog plus 5 μM JNK_i_. DMSO-treated and JNK_i_-treated control samples were also collected. After 24 hours of treatment, the cells were trypsinized, collected, centrifuged, resuspended in DPBS, and counted with trypan blue staining. Aliquots containing 10 million cells were taken from each sample and centrifuged. These pellets were resuspended in 500 microliters of ice-cold M-PER lysis buffer (Fisher Scientific catalog # PI78503), with the addition of 1x Halt Protease and Phosphatase Inhibitor Cocktail (Fisher Scientific catalog # PI78441), and agitated for 30 minutes at 4 °C. To ensure adequate lysis, these samples were subjected to three 30-second cycles of sonication at 20 kHz and 10% amplitude, with incubation in ice for 30 seconds between each sonication step. A Fisher Scientific Model 50 Sonic Dismembrator (Fisher Scientific catalog # FB50110) was used for sonication. After sonication, the cell lysates were centrifuged at 4 °C and 13,000 RPM for 10 minutes, and the protein extracts (in the supernatants) were collected, aliquoted, and stored at −80 °C.

### Western Blot

Lysates from A375 cells which were treated with DMSO, JNK_i_, AZA, CYT, or co-treated with each cytidine analog plus JNK_i_ were used to compare levels of the c-Jun protein, as well as phospho-c-Jun (Serine 73). Protein concentrations in each sample were measured using a BCA assay (Fisher Scientific catalog # FERA65453), following the manufacturer’s instructions. Samples were prepared by mixing protein lysates with Invitrogen Bolt LDS Sample Buffer (4X) (Fisher Scientific catalog # B0007), Bolt Sample Reducing Agent (10X) (Fisher Scientific catalog # B0009), and UltraPure distilled water (used to normalize protein concentration in each sample) (Fisher Scientific catalog # 10977-015). Samples were denatured at 95 °C for 5 minutes, and then loaded in Bolt 1.0mm x 15-well 12% Bis-Tris gels (Fisher Scientific catalog # NW00125BOX). 50-100 μg of protein was loaded into each well. The PageRuler Plus Prestained Protein Ladder was also loaded to guide membrane cutting (Fisher Scientific catalog # 26619). The SuperSignal Molecular Weight Protein Ladder (Fisher Scientific catalog # 84785) was also loaded and visualized in our images after the membranes were stained.

Gels were run at 120 V for 120 minutes in Bolt MES SDS Running Buffer (Fisher Scientific catalog # B0002) supplemented with 1% Bolt Antioxidant (Fisher Scientific catalog # BT0005). After separation, proteins were blotted onto Novex PVDF membranes (0.2 μm pore size) (Fisher Scientific catalog # LC2002). Membranes were first activated with methanol (Fisher Scientific catalog # A412-500) for 30 seconds and rinsed with agitation in DI water for one minute. After the membranes were activated and oriented with the gels, blotting was performed at 90 V for 60 minutes in Bolt Transfer Buffer (Fisher Scientific catalog # BT00061), and the blotting tank was cooled with ice packs. After transferring the proteins to the membranes, the membranes were trimmed according to the appropriate molecular weight range. Membranes were washed with DI water, and then treated with SuperSignal Western Blot Enhancer Antigen Pretreatment Solution (Fisher Scientific catalog # 1862324) for 10 minutes with agitation. The membranes were washed with DI water and then blocked in a 5% milk solution, prepared using instant non-fat dry milk powder (Fisher Scientific catalog # M17200-500.0) in TBS Tween 20 Wash Buffer (Fisher Scientific catalog # 28360). Membranes were blocked for 120 minutes with agitation.

After blocking, the milk solution was removed, and the TBS Tween 20 Wash Buffer was used to wash the membranes 3 times for 10 minutes each with agitation. The membranes were then incubated with primary antibodies diluted 1000-fold in SuperSignal Western Blot Enhancer Primary Antibody Diluent (Fisher Scientific catalog # 1862324). The following primary antibodies were used: Invitrogen c-Jun Recombinant Polyclonal Antibody (Fisher Scientific catalog # PI711202) and Invitrogen Phospho-c-Jun (Ser73) Polyclonal Antibody (Fisher Scientific catalog # 44-292-G). In general, N-terminal phosphorylation activates c-Jun (Ser63 and Ser73), whereas C-terminal phosphorylation inhibits c-Jun (Thr231, Thr239, Ser243, Ser249) [55]. Therefore, detection of c-Jun which has been phosphorylated at Ser73 indicates the activation of c-Jun. Primary antibody incubation took place overnight at 4 °C with agitation and protection from light. Membranes were then washed in wash buffer 4 times for 5 minutes each with agitation. Then, membranes were incubated with a rabbit IgG Horseradish Peroxidase-conjugated Antibody (Fisher Scientific catalog # HAF008) diluted 2000-fold in Blocker FL Fluorescent Blocking Buffer (Fisher Scientific catalog # NC2869193). Secondary antibody incubation took place for 60 minutes with agitation. Then, the secondary antibodies were removed, and the membranes were washed in wash buffer 3 times for 10 minutes each with agitation. The wash buffer was removed, and Pierce 1-step Ultra TMB Blotting Solution (Fisher Scientific catalog # 37574) was added to cover the membranes. Membranes were protected from light for roughly 60 seconds, and then washed twice with DI water and allowed to air dry. Images were taken by cellular camera.

### Immunodetection Array Experiments

The levels of phosphorylation of numerous proteins were compared between A375 cell lysates which were either DMSO-treated or treated with 20 nM GEM for 24 hours. Three biological replicates were performed for each condition. Human phosphorylation arrays, which are printed with antibodies for specific phosphorylated proteins, were purchased from Raybiotech Inc (Peachtree Corners, GA). Arrays were purchased for the following signaling pathways: AKT, JAK/STAT, MAPK, NF-κB, and TGF-β (RayBiotech catalog #’s AAH-AKT-1, AAH-JAKSTAT-1, AAH-MAPK-1, AAH-NFKB-1, and AAH-TGFB-1, respectively). The instructions from the manufacturer were followed for these experiments. Briefly, the arrays were incubated with a blocking buffer at room temperature for 30 minutes, then incubated with the protein extract samples overnight at 4 °C, washed with wash buffers, and incubated with a detection antibody cocktail. After incubation with detection antibodies, the arrays were washed, incubated with the secondary, HRP-conjugated antibodies, washed again, and exposed to the detection buffer. Then, chemiluminescent images of the arrays were captured using a UVP ChemStudio (catalog # 849-97-0928-02, Analytik Jena, Jena, Germany). The signals from these chemiluminescent images were quantified and normalized according to manufacturer instructions, and signals corresponding to each protein were compared between untreated and treated samples.

### Viability Assays Using Flow Cytometry

Cells were harvested from T-75 flasks and 300,000 cells were seeded per well in 6-well plates. After 24 hours of incubation to allow the cells to attach, the medium was removed and replaced with various concentrations of drugs, in either single or co-treatment. After 72 hours of treatment, samples were trypsinized, collected, centrifuged, and resuspended in DPBS. Cell suspensions were stained for 15 minutes at 37 °C with 250 nM of both SYTO60 red (Fisher Scientific catalog # S11342) and SYTOX green (Fisher Scientific catalog # S7020) fluorescent nucleic acid stains. These samples were then analyzed with flow cytometry. Live and dead cell populations are distinct under these conditions, as SYTO60 Red penetrates all cells (live and dead), whereas SYTOX Green can only penetrate cells with compromised membranes (dead cells only). Survival fractions were calculated based on the total number of live cells in each sample, compared to the number of live cells in the DMSO-treated control. Kill curves for each drug were plotted as normalized survival fraction vs. drug concentration. The co-treatment data were analyzed by comparing survival fractions under each treatment condition.

Co-treatment concentrations for AZA, CYT, and GEM were determined from the concentration-dependent kill curve results. Concentrations close to 10-fold above our calculated IC_50_ values were chosen, in the plateau region where the survival fraction was relatively independent of increasing concentration (Figures S2, S6). In cases where the plateau region resulted in almost complete killing of the cells, we chose a slightly lower concentration to increase the number of cells available for analysis. Similarly, JNK_i_ co-treatment concentrations were determined with concentration-dependent kill curves (Figures S2, S6). For JNK_i_, we chose the maximum concentration that did not result in a substantial decrease in cell survival fraction.

### Transfection with c-Jun siRNA

A375 melanoma cells were initially seeded in T-75 flasks at a density of 2.5 million cells per flask and allowed to attach and grow for 24 hours. Subsequently, cells were harvested, and 150,000 cells were seeded per well into 6-well plates using DMEM medium supplemented with 10% FBS, but **without** antibiotics. Transfection of c-Jun siRNA into A375 cells was performed using Lipofectamine RNAiMAX Transfection Reagent, with negative control siRNA used for comparison. RNA-Lipofectamine complexes were prepared in Opti-MEM Reduced Serum Medium by diluting 5 μL or 7.5 μL of Lipofectamine Reagent in 250 μL of Opti-MEM Medium, respectively. Stock solutions of c-Jun siRNA and negative control siRNA (50 μM) were diluted to a working concentration of 10 μM. Solutions of varying siRNA concentrations (10 nM or 20 nM) were prepared by diluting 3 μL or 6 μL of siRNA in 250 μL of Opti-MEM Medium. After mixing with diluted Lipofectamine Reagent in a 1:1 ratio and incubating for 20 minutes at room temperature, the siRNA-lipid complexes were added to cells and incubated at 37 °C. Following 24 hours of incubation, the medium was replaced with fresh medium plus 20 nM GEM for an additional 24-hour treatment. Post-treatment, cells were collected for western blot analysis according to our standard protocols. For viability assays, siRNA-transfected cells were seeded at 300,000 cells per well in new 6-well plates after 24 hours of incubation and treated with 20 nM GEM for 72 hours. Untreated cells were used as controls across all conditions. After the treatment period, samples were collected, and cell viability was assessed to determine survival rates. Results from western blot analysis and cell survival assays are reported specifically for the condition using 10 nM c-Jun siRNA and 7.5 μL Lipofectamine Reagent, as all tested conditions yielded comparable outcomes.

### Proteomics Data Analysis and Statistics

After our proteomics samples were processed, we received the datasets from the University of Houston Mass Spectrometry Laboratory. These data were processed in Microsoft Excel, as described previously [20]. Briefly, Excel was used for several data processing steps, including data transformation and normalization, as well as calculations for fold change and statistical significance (P values). Proteins which were detected and quantified in less than 11 of the 12 samples were excluded from analysis. The dataset was transformed to a normal distribution by logarithmic transformation. Next, the data were normalized with both the average and slope methods to minimize the effects of variation between technical replicates. The Probabilistic Minimum Imputation method was used to replace missing values. Then, the relative ratio of each protein in each treatment group (*e.g.*, untreated, GEM-treated) was determined, and parametric t-tests were used to calculate P values. F-tests were used to determine whether the replicated measurements for each protein exhibited equal or unequal variances, and the results of the F-tests were used to determine the t-test type. Significantly changed protein networks were identified using STRING Version 12.0 [21].

Data from all other experiments were analyzed in Microsoft Excel and GraphPad Prism 8.3.0. All presented figures were produced using GraphPad Prism 8.3.0. Unless otherwise stated, at least four independent biological replicates were performed for each condition of each experiment. All data points represent mean values, and error bars represent standard error values. Asterisks signify statistical significance using pair-wise t-tests or two-way ANOVA with Sidak’s multiple comparisons post hoc testing: *P<0.05, **P<0.01, ***P<0.001, ****P<0.0001.

## Supporting information

Supplemental Proteomics Dataset

## Acknowledgments

The authors thank the members of the Orman Lab for their assistance. Support from University of Houston Startup funds, NSF CAREER Award 2044375, and NIH/NIAID R01-AI143643 is acknowledged. The DoD SMART Program is acknowledged for supporting Shayne Sensenbach, the first author.

## Contributions

S.S., P.K., V.A., and M.A.O. conceived and designed the study. S.S., H.N., S.G., P.K., and V.A. performed the experiments. Proteomics experiments were conducted at the Mass Spectrometry Laboratory of Dr. Chengzhi Cai at the University of Houston, with associated service fees. S.S. analyzed the data and wrote the manuscript. M.A.O. revised the manuscript. All authors have read and approved the final manuscript.

## Data availability

All data in this manuscript can be found in either the Main text or Supplementary file. The mass spectrometry proteomics data have been deposited to the ProteomeXchange Consortium via the PRIDE partner repository with the dataset identifier PXD060382.

Reviewer access details – Log in to the PRIDE website using the following details:

Project accession: PXD060382

Token: e93KHf7dSRad

