## Supporting Information for "The Role of c-Jun Signaling in Cytidine Analog-Induced Cell Death in Melanoma"

**Supplemental Proteomics Dataset**

The proteomics data are provided in an Excel spreadsheet. This spreadsheet was used for data processing as outlined in the Materials and Methods section. (XLSX)

**Supplemental Discussion of Proteomics Data**

***Overview of GEM-induced Protein Upregulations***

Our initial proteomics analysis revealed GEM-induced upregulations in the DDR, cell cycle regulation, mitosis, apoptosis, the ETC, mitochondrial ribosomal activity, protein folding and processing, mRNA splicing, and multivesicular body (MVB) transport.

The upregulated proteins related to the DDR include BLM, CLSPN, DDB2, PML, RAD18, RBM38, RPA2, SOD1, and XPC. Numerous proteins integral to other discussed pathways—such as cell cycle regulation, apoptosis, and protein folding—are interconnected with DDR, due to the intricacies of DDR pathways. Treatment with GEM, and chemotherapeutics in general, is anticipated to trigger extensive upregulation of DDR proteins [22], [23]. Indeed, prior research has demonstrated GEM's notable activation of ATM (ataxia telangiectasia mutated) and ATR (ataxia telangiectasia and Rad3 related), well-established initiators of DDR [56]. Notably, cells with impaired ATM and ATR function display increased sensitivity to GEM treatment [56]. Additionally, an ATR inhibitor was shown to enhance sensitivity to GEM treatment in pancreatic cancer cells, demonstrating the therapeutic promise of ATR inhibition [57]. Interestingly, we observed reduced levels of ATM and ATR proteins following GEM treatment. However, this downregulation does not necessarily denote inactivation (see GEM-induced Downregulations section, below, for further explanation). In fact, the concurrent upregulation of multiple DDR-related proteins suggests hyperactivity of ATM and ATR, given their pivotal roles in initiating

DDR signaling. Considering GEM's inhibitory effects on DNA replication and induction of DNA damage, these findings align with rational expectations.

The cell cycle regulating proteins which were upregulated by GEM treatment include CCNA2, CDK5RAP2, CKAP2, GMNN, MYBL2, and SKP1. Upon DNA damage and activation of the DDR, the upregulation of proteins regulating the cell cycle is expected. Progression of the cell cycle is controlled at specific checkpoints which are influenced by the DDR [58]. For example, the DNA damage induced activation of p53 is known to arrest the cell cycle at multiple points including the G2/M transition, preventing the cell from entering mitosis [59]. Specifically, the p53-induced activation of p21 inhibits CDKs, including CDK1, causing G2/M arrest [60]. The group of upregulated cell-cycle regulating proteins encompasses those involved in G1/S, S, and G2/M checkpoints. As all these transition states are associated with the DDR [22], these results align with current literature.

Moreover, we identified upregulated mitosis-related proteins following GEM treatment, including ANLN, CDCA5, CDCA8, INCENP, KIF11, KIF2C, NUDC, PRC1, TOP2A, and TPX2. These proteins participate in both the regulation and execution of mitosis. The upregulation of these proteins indicates cell cycle arrest at the mitotic spindle checkpoint, and potentially mitotic cell death. Prolonged mitotic arrest can trigger apoptosis, akin to cell cycle arrest at other checkpoints [61]. However, mitotic arrest has been identified as a survival mechanism in cancer cells with damaged DNA and may allow cancers to acquire chemotherapeutic resistance genotypes [62]. Cumulatively, the upregulations of cell cycle-regulating proteins and mitotic proteins indicate arrest throughout several cell cycle checkpoints. Therefore, sub-populations of the GEM-treated cells arrest at various points, all of which can culminate in cell death. The sub-populations which halt at the G1/S and S checkpoints (early in the cycle) may exhibit faster rates of cell death, whereas those proceeding to the G2/M and spindle checkpoints may display slower rates. Given that our analysis focused solely on A375 cells surviving 24 hours of GEM treatment, these differential killing rates may account for the significant upregulation of proteins associated with the G2/M transition and mitosis.

The upregulated proteins related to apoptosis include BAX, BCL2L1, FAS, FOXO3, JUN, PDCD4, TNFRSF10B, and YAP1. Given that we demonstrated the high sensitivity of A375 cells to GEM treatment (Figure 3B), and that GEM elicits toxicity by causing DNA damage, we expected to observe upregulations in apoptotic proteins. These observations also align with the observed upregulations in the DDR and in cell cycle regulation, as these pathways are well-known to be interconnected [22], [23], [24]. In the context of cancer therapy, these pathways are of utmost importance, so we were particularly interested in further describing these mechanisms. Due to the upregulation of c-Jun which we consistently observed across multiple experiments, and due to the crucial role of c-Jun in various cellular signaling activities [28], we focused on investigating the role of c-Jun in this context.

The proteins related to the ETC which were upregulated by GEM treatment include ATP5PD, COX5A, COX6B1, NDUFA8, NDUFB5, NDUFB8, NDUFB9, and NDUFS4. These upregulations show a strong induction of oxidative phosphorylation. This metabolic change may be induced by the treatment itself, or this observation may be due to higher survival rates in cells with highly active oxidative phosphorylation, or both. Metabolic shifts have been previously reported in various cancer cell types in response to treatment with various chemotherapeutic and targeted cancer therapeutic drugs [13], [14]. In our previous in-depth study of A375 persister cell metabolism, we also highlighted consistent upregulations in Krebs cycle activity in response to chemotherapeutic treatment [15]. Therefore, these results are consistent with current literature, and they provide further verification for our previous work.

Several proteins related to mitochondrial ribosomal activity were also upregulated, including MRPL13, MRPL24, MRPL41, MRPL42, MRPL58, MRPS18B, MRPS26, MRPS28, and TSFM. Changes in the expression of mitochondrial ribosomal genes are associated with apoptosis [63], [64], [65], which partially explains these observations. However, using antibiotics to target mitochondrial ribosomes has been shown to 1) reduce the clonogenicity and proliferation of cancer cells [48], and 2) increase the efficacy of chemotherapeutic treatment [49], [50]. The anticancer efficacy of these antibiotics may be related to the increased mitochondrial activity we previously reported in A375 GEM persisters [15]. As A375 GEM persisters have relatively high usage of the Krebs cycle and oxidative phosphorylation, an increase in mitochondrial ribosomal activity could be instrumental in attaining the metabolic shifts observed in cancer persisters.

The proteins related to protein folding and processing which were upregulated by GEM treatment include CALR, CANX, DNAJB2, DNAJC2, DNAJC3, DNAJC7, ERO1A, HLA-A, HLA-B, HLA-C, HLA-DMA, HLA-DRA, HSPA5, HSPB1, and P4HB. This group of proteins encompasses molecular chaperones and HLA proteins, both of which are known to be overexpressed in response to DNA damage [66], [67], [68]. Also, the expression of chaperones, commonly referred to as heat shock proteins (HSPs), is crucial in cancer development and the sensitivity of cancers to treatment [66]. In fact, one such protein, HSP90, has been extensively studied as a target for cancer therapy, as it influences the folding and expression of several oncogenic proteins [69], [70]. Although the upregulation of this group of proteins may be an expected outcome of GEM treatment, the specific HSPs which were upregulated may provide novel targets to improve the chemotherapeutic treatment of melanoma.

The proteins related to mRNA splicing which were upregulated by GEM treatment include BCAS2, CDC5L, PRPF3, PRPF38A, RBM25, SNRPA, SNRPD2, SYF2, and U2AF2. Changes in the expression patterns of proteins related to mRNA splicing are known to be induced by DNA damage, and some of these proteins are associated with chemoresistance in cancer [71], [72], [73]. We also identified proteins related to MVB transport processes which were upregulated by GEM treatment, including CHMP2B, CHMP3, CHMP6, IST1, VPS25, VPS28, and VPS37B. MVB transport processes sort and degrade proteins within endosomes [74]. Interestingly, one MVB related protein, CHMP1A, has been shown to be important in the induction of cell death upon chemotherapeutic treatment [75].

***Overview of GEM-induced Protein Downregulations***

Our proteomics analysis revealed significant downregulations in proteins related to the DDR, cell cycle regulation, RNA processing and metabolism, ribosome biogenesis and translation initiation, tRNA aminoacylation, and Golgi body vesicular transport. Cumulatively, these observations are logical outcomes of the extensive DNA damage caused by treatment with GEM.

The DDR related proteins which were downregulated by GEM treatment include ATAD5, ATM, BRCA2, MLH1, MSH3, MSH6, and XRCC6. Clearly, the DDR is not suppressed, but activated by DNA damage, and the ATM and ATR proteins are known as key DDR activators [22], [23], [56]. Here, we report a decrease in the expression of ATR upon GEM treatment, although this shift was statistically insignificant ($P=0.2795$). We also observed a significant reduction of ATM expression ($P<0.01$) upon GEM treatment. However, while DNA damage activates both ATM and ATR, the expression of these proteins is not necessarily induced by DNA damage [76], [77]. Complex mechanisms which are not fully understood influence the expression of these proteins, including autoregulatory mechanisms [77], [78]. Further, mRNA transcripts of both ATR and ATM have been shown to be significantly lower in blood cancer patients treated with chemotherapy, compared to untreated patients [79].

GEM-induced downregulations were also observed in proteins related to cell cycle regulation, including ANAPC4, CDC16, CDC27, CDK1, CEP192, CHEK1, HSP90AB1, KIF20A, MCM2, MCM5, MCM6, NCAPD2, PDS5A, PLK1, PPP6C, RBL1, RPS6KA3, and ZWILCH. As discussed above, severe DNA damage is expected to halt the cell cycle at specific checkpoints [58]. Many proteins are involved in each checkpoint of the cell cycle; some of these proteins promote, while others inhibit, the progression of the cell cycle. Therefore, severe DNA damage is expected to induce both the activation and inhibition of various cell-cycle-related proteins. For example, p53 activates many proteins [80], but inhibits others by activating p21, which inhibits CDKs [60], [81], [82]. Therefore, it is reasonable to observe downregulated cell-cycle-related proteins due to DNA damage. Interestingly, we observed a downregulation of CDK1 (also known as Cdc2). As CDK1 promotes cell cycle progression through the G2/M transition [83], downregulated CDK1 inhibits cell cycle progression. CDK1 phosphorylates many proteins and plays crucial roles in cell cycle regulation [83], [84], as well as apoptosis [85], [86]. Similar to CDK1, many more of these downregulated proteins have positive regulatory effects on the cell cycle, indicating that they are expected to be downregulated or inhibited while the cell cycle is arrested. Cumulatively, the upregulated and downregulated proteomics data imply drastic arrest and dysregulation of the cell cycle in response to severe DNA damage.

Several proteins involved in RNA processing and metabolism were also downregulated by GEM, including CPSF2, ELAC2, EXOSC1, EXOSC10, DIMT1, GTPBP1, LRPPRC, METTL16, MTPAP, MTREX, NAT10, PAPOLA, PAPOLG, PDCD11, RO60, SUPV3L1, TENT4B, TFB2M, THOC2, THOC6, TSR1, UTP20, VIRMA, WDR74, ZC3H3, and ZCCHC4. Proteins which influence RNA metabolism, such as METTL3, have been shown to be important to cancer [87], and specifically to cancer's response to chemotherapy [88]. Also, downregulations in RNA processing genes have been previously reported in cancer persister cells upon chemotherapeutic treatment [89]. Interestingly, many of the downregulated proteins in this group are related to ribosomal RNA (rRNA) processing and maturation. Decreased production of rRNA is directly related to decreased ribosome biogenesis, translation initiation, and transfer RNA (tRNA) aminoacylation, all of which are discussed below. Therefore, this group of downregulated proteins aligns with current literature as well as the rest of this dataset.

The proteins related to ribosome biogenesis and translation initiation which were downregulated by GEM treatment include EFL1, EIF2B1, EIF2B2, EIF2B3, EIF3C, EIF3F, EIF3L, EIF4A1, PDCD11, RPL10, RPS27, TEX10, TSR1. Ribosome biogenesis is known to be inhibited by treatment with chemotherapeutic drugs [90], [91]. When ribosome biogenesis is inhibited, a decrease in translation initiation is also expected. We also observed downregulations in several proteins related to tRNA aminoacylation, including AARS2, DARS1, FARSA, FARSB, MARS1, NARS1, RARS1, TARS1, VARS1, and YARS2. These cellular activities are directly linked to each other, so each of these sets of downregulated proteins corroborate each other. Taken together, these findings demonstrate significantly decreased levels of translation.

We also identified a group of downregulated proteins related to Golgi body vesicular transport, including ARF4, COG1, COG2, COG4, COG7, COPG2, GBF1, GOSR1, SCFD1, SEC24B, and SEC24C. The Golgi body is known to play a role in apoptotic signaling [92], [93]. In fact, during apoptosis, caspases target multiple proteins in the Golgi body, disrupting its function [92], [93]. Given its involvement in apoptosis, and its connection to the endoplasmic reticulum (ER) and the unfolded protein response (UPR), the Golgi body has become a target for novel cancer therapeutics [94]. As we clearly observed upregulated apoptotic signaling upon GEM treatment, it is rational to observe a concurrent downregulation of Golgi body vesicular transport. However, as the Golgi body is involved in apoptosis, targeting the Golgi body could lead to enhanced apoptotic death during chemotherapeutic treatment.

Considering our analysis of the GEM-induced changes in the proteome of A375 melanoma cells, we offer verification of our previous study [15], and elaborate further on our previous characterization of the GEM persistence state in A375 melanoma cells. Specifically, our findings related to upregulated cell cycle regulation and upregulated ETC activity mirror our previous findings using distinct experimental methods. We also identified several additional phenomena related to A375 GEM persistence, including upregulated mitochondrial ribosomal activity and HSP expression, as well as downregulated RNA processing and Golgi body vesicular transport. These findings provide additional targets for melanoma persisters, which could lead to enhanced methods to treat melanomas with chemotherapeutics. Most interestingly, upregulated mitochondrial ribosomal activity is directly related to ETC activity, which we highlighted in our previous study [15]. As mitochondrial ribosomes have been studied as a target in cancers in previous reports [48], [50], [65] they are therefore a compelling target for eradicating melanoma chemotherapeutic persisters.

***Overview of $JNK_i$ Co-treatment-induced Protein Upregulations and Downregulations***

Our second proteomics dataset compared A375 cells treated with GEM only to those treated with GEM plus $JNK_i$. These data revealed that in comparison to the GEM-treated cells, the co-treated cells exhibited significant upregulations in proteins related to sterol biosynthesis and cholesterol transport. Similarly, significant downregulations were observed in proteins related to cell cycle regulation, mitosis, and apoptosis.

The upregulated proteins related to sterol biosynthesis include CYP51A1, DHCR24, HSD17B7, LSS, MVD, and SREBF2. The cholesterol biosynthesis pathway is well-known to be involved in cancer cell proliferation [95]. In fact, lipid concentrations in blood are used as biomarkers for cancer risk [96]. Directly related to these findings are our observed upregulations in proteins related to cholesterol transport, including APOB, APOC2, APOC3, and APOE. Each of these are apolipoproteins, which are known to promote cancer cell survival and proliferation [97], [98], as well as disease progression [99], [100]. Intriguingly, connections have been previously drawn between JNK activity and the expression of APOC1 [99] and APOE2 [97]. However, while APOC1 was previously shown to inhibit the JNK pathway [99], we report that JNK-inhibition increases apolipoprotein expression during chemotherapeutic treatment. These findings suggest that JNK-inhibition allows A375 melanoma cells to evade apoptosis and maintain proliferative activity partly through the sustained expression of apolipoproteins.

Proteins related to cell cycle regulation which were shown to be downregulated include CCNB1, CDCA3, CDC5L, CDK2, CDK5RAP2, CLSPN, GMNN, MYBL2, RPA3, and TP53BP1. Similarly, mitosis related proteins were downregulated, including ANLN, CDCA5, CDCA8, CENPF, ERCC6L, INCENP, KIF11, KIF14, KIF22, KIF2C, KIF20A, KIF23, KIFC1, PRC1, RACGAP1, SGO1, SKA2, SKA3, SPDL1, SPICE1, TOP2A, and TPX2. Both these groups show striking similarities to the proteins in the same pathways which were upregulated by GEM treatment. We previously concluded that GEM treatment led to cell cycle arrest at multiple checkpoints, including the mitotic spindle checkpoint. Therefore, these downregulated proteins indicate that co-treatment with $JNK_i$ prevents the arrest of the cell cycle caused by GEM. While these findings are logical and aligned with many previously discussed points, they do not necessarily offer insight into the mechanism(s) by which $JNK_i$ attenuates apoptosis.

The proteins related to apoptosis which were downregulated include BAD, CASP8, DIDO1, and JUN. Downregulation of the c-Jun protein due to JNK-inhibition was clearly expected, and this observation serves as a confirmation that the treatment had an on-target effect. As detailed in Results section, the statistical significance of c-Jun suppression was greater in cells treated with GEM plus $JNK_i$ compared to those treated with only $JNK_i$. This quantitative comparison provides further proof of effective JNK-inhibition. However, the rest of these downregulations provide insight into how JNK-inhibition prevents apoptosis under these conditions. CSP8 (caspase-8) is an established apoptosis-inducer, and its activation has been associated with c-Jun activity [101], and similar findings have been reported regarding other caspase proteins [102], [103]. The BAD (BCL2 associated agonist of cell death) protein has been shown to be inhibited by active JNK, thereby preventing apoptosis [104]. However, active c-Jun stimulates BAD protein expression [39]. Under our experimental conditions, BAD is suppressed during JNK-inhibition, contributing to the prevention of apoptosis. Interestingly, the knockdown of the DIDO1 (death inducer obliterator 1) protein has been shown to decrease JNK activity and induce apoptosis [105]. However, we observed JNK-inhibition leading to decreased expression of DIDO1 protein, while preventing apoptosis. Our observations once again emphasize that the effects of JNK/c-Jun activity or inhibition are dependent on many factors. In the case of melanoma cells treated with cytidine analogs, the anti-apoptotic effect of JNK-inhibition is at least partly due to the suppression of signals from BAD, CASP8, and DIDO1. These molecules are potential targets for the enhancement of chemotherapeutic treatment of metastatic melanoma.

In summary, the JNK/c-Jun pathway has been connected to multiple apoptosis-related proteins in multiple cancer and tissue types [39], [101], [102], [103], [104], [105]. However, in some cases, JNK activity prevents apoptosis [104], [106]. Also, while the JNK/c-Jun pathway is well-known to promote apoptosis under certain conditions, mechanistic understanding of how this pathway drives apoptosis is not well-known [106]. Under our experimental conditions, the BAD, CASP8, and DIDO1 proteins appear to be crucial to the apoptotic signals induced by the JNK pathway. Another mechanism by which JNK-inhibition prevents apoptosis and sustains proliferation levels is through enhanced apolipoprotein expression. Our findings related to JNK-inhibition clarify the function of this pathway in the context of melanoma cells treated with cytidine analogs, and provide targets for the enhancement of chemotherapeutic treatment of metastatic melanoma.

### Supplemental Figures

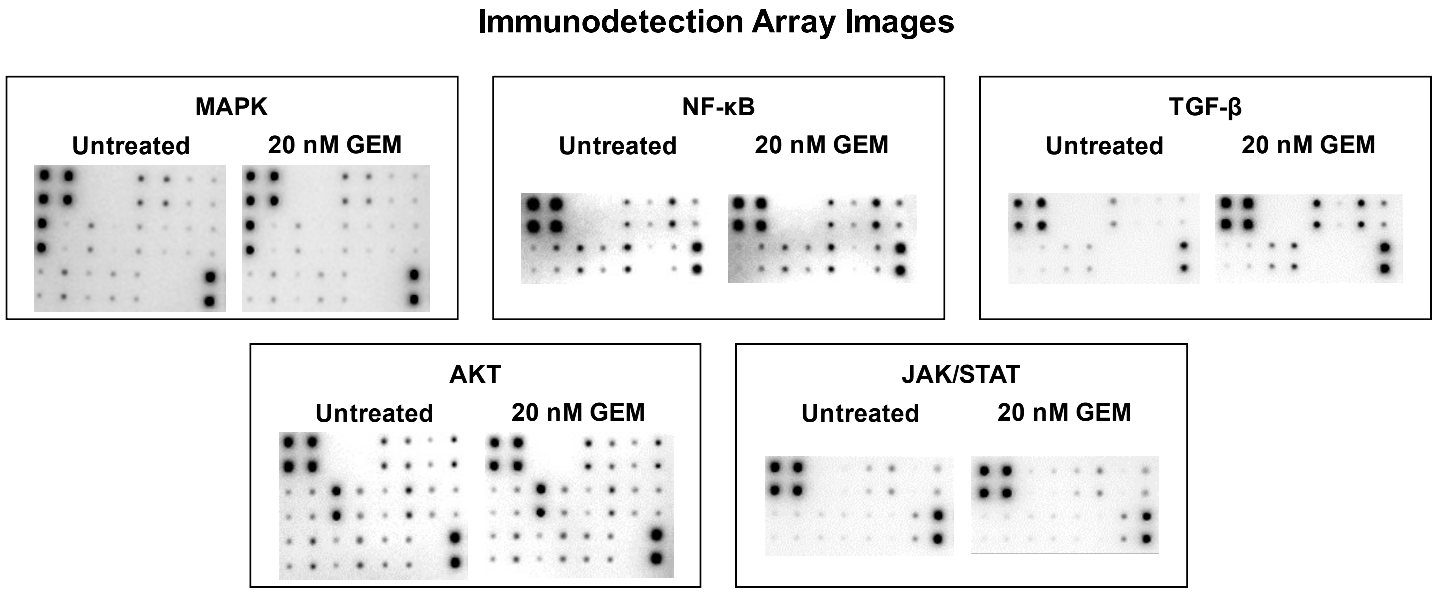


**Figure S1.** Images collected during the immunodetection array experiments. These images were collected and processed with a UVP ChemStudio (catalog # 849-97-0928-02, Analytik Jena, Jena, Germany), resulting in the plotted data shown in Figure 2. Each set of membranes were used to compare A375 cell lysates which were untreated or treated with gemcitabine (GEM) for 24 hours. N = 3.

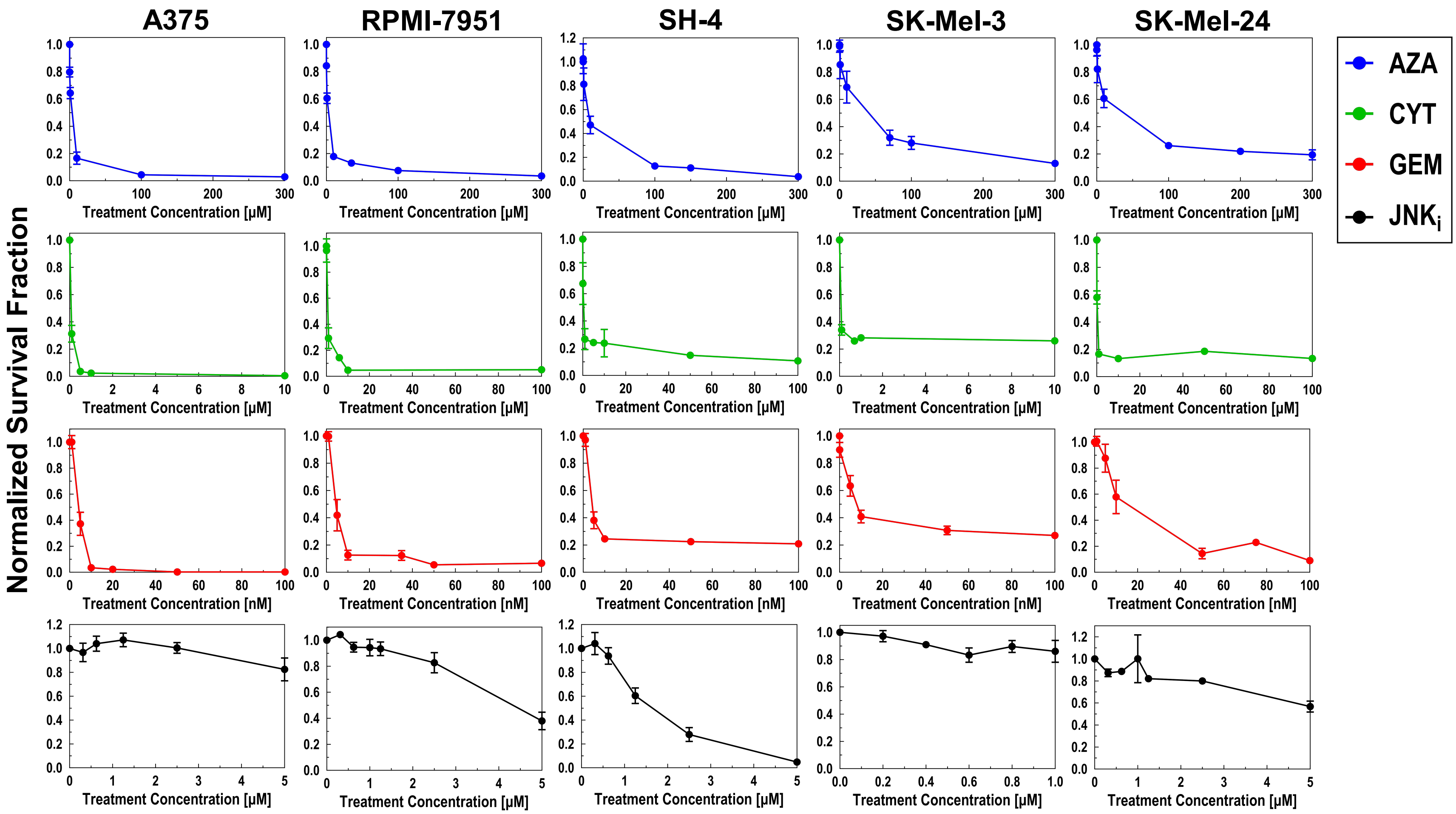


**Figure S2.** Concentration-dependent kill curves for melanoma cell lines. The listed cell lines were treated with varying concentrations of each of the listed drugs: azacitidine (AZA), cytarabine (CYT), gemcitabine (GEM), and JNK-IN-8 (JNK$_i$). Survival fractions were measured by flow cytometry after 72 hours of treatment under the indicated conditions and normalized to the untreated cell number. Treatment with each drug is distinguished using a color code. These concentration-dependent kill curves were used to determine treatment concentrations in other experiments, as detailed in the methods section (see "Viability Assays Using Flow Cytometry"). N = 4.

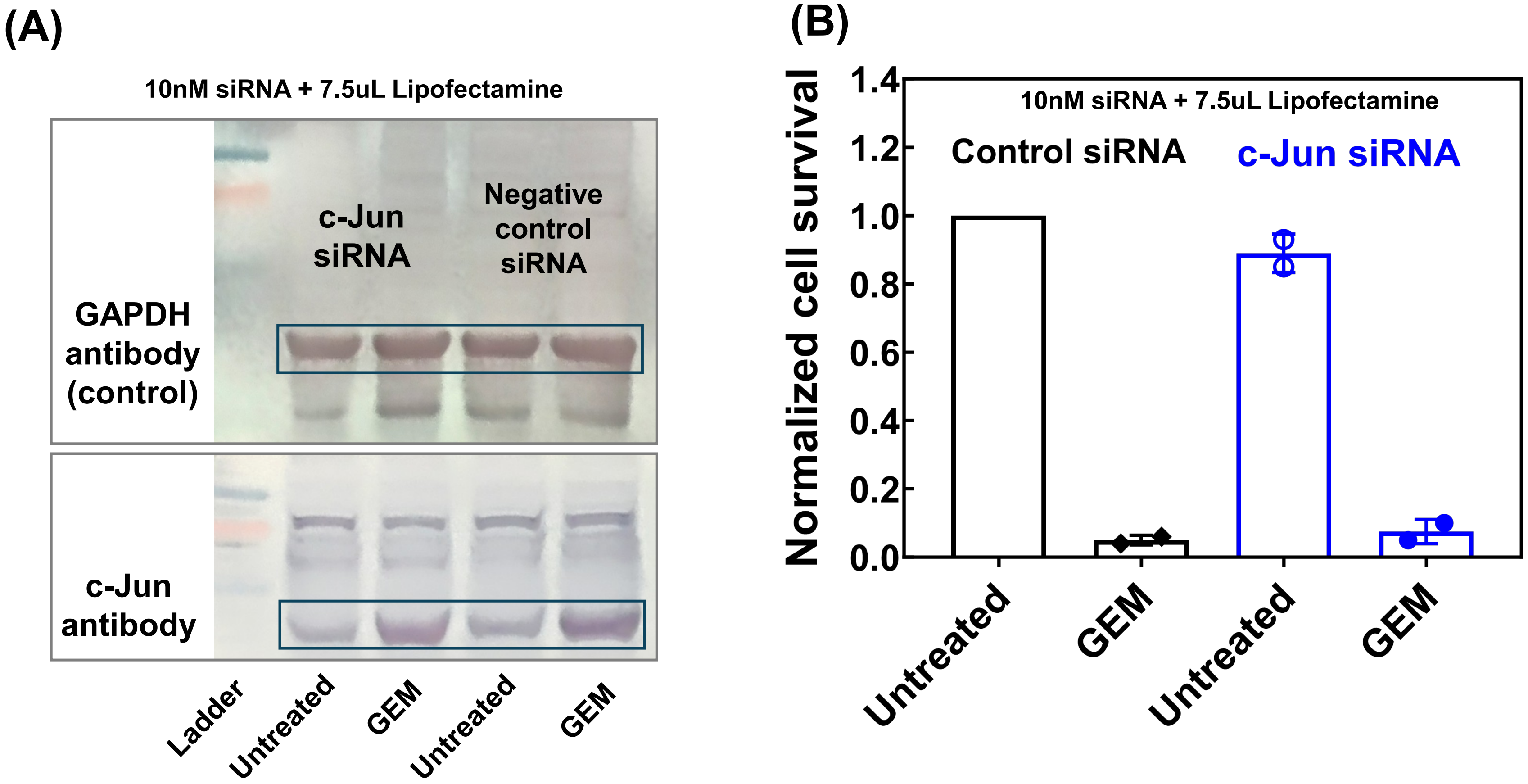


**Figure S3.** Effects of c-Jun targeted knockdown on A375 cells. (A) Representative western blot image comparing c-Jun siRNA-transfected A375 cells with negative control siRNA-transfected cells after 24-hour treatment with 20 nM gemcitabine (GEM). The siRNA-lipid complex was prepared using 7.5 μL Lipofectamine RNAiMAX Transfection Reagent in Opti-MEM medium with a 10 nM siRNA solution. After incubation at 37°C for 24 hours, cells were exposed to 20 nM GEM for an additional 24 hours prior to western blot analysis. GAPDH (loading control) protein levels are shown in the top row, and c-Jun protein levels in the bottom row. (B) A375 survival fractions were assessed by flow cytometry following 72-hour treatment with 20 nM GEM under the described conditions, normalized to untreated negative control siRNA-transfected cells. N = 2.

### Original Western Blot Images – Primary Antibody: c-Jun

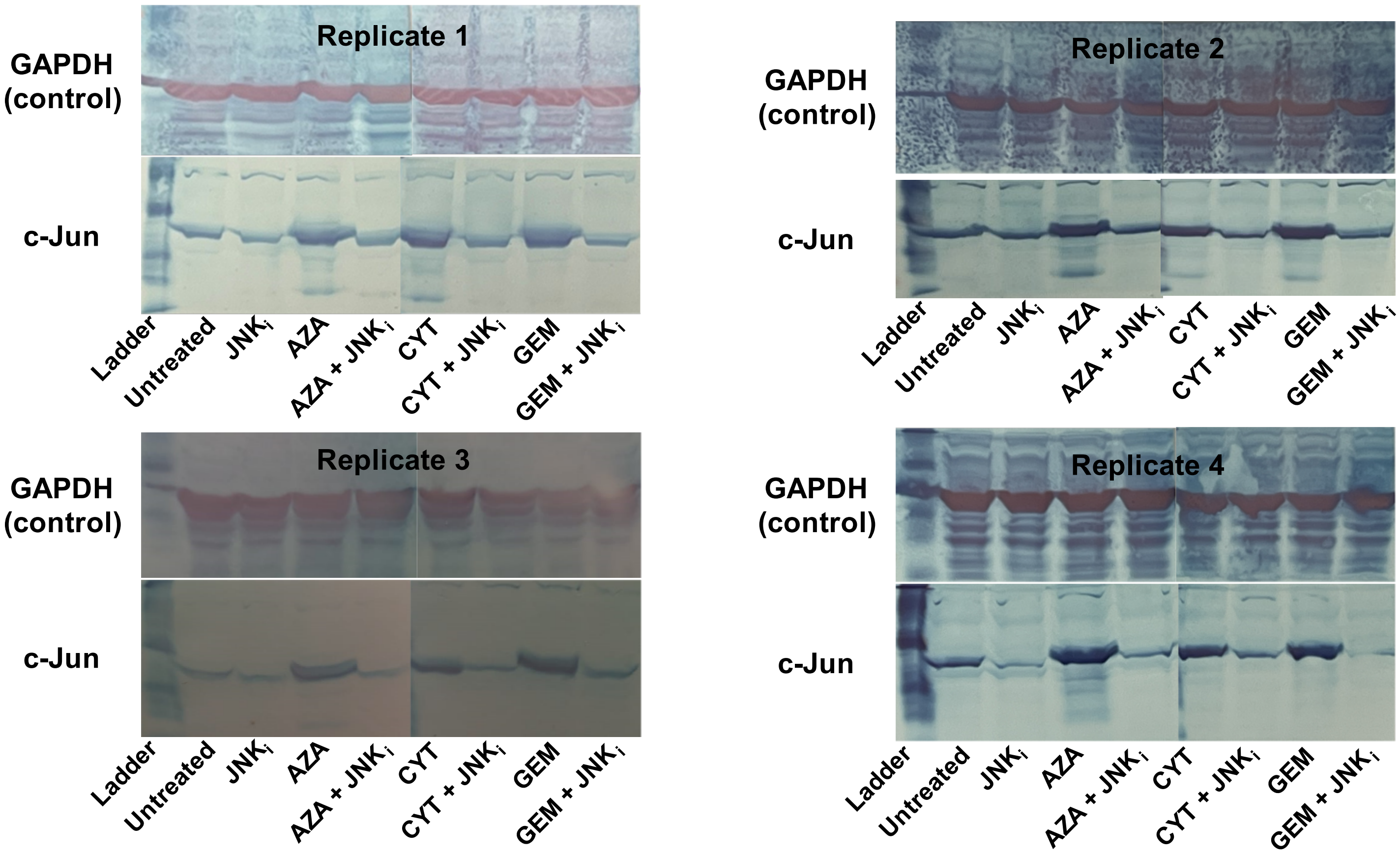


**Figure S4.** Original images collected during western blot experiments. These experiments compare total c-Jun protein expression (primary antibody was specific to human c-Jun) in A375 cells under the indicated treatment conditions. The following drugs were used: azacitidine (AZA), cytarabine (CYT), gemcitabine (GEM), and JNK-IN-8 ($JNK_i$). Treatment concentrations can be found in Supplemental Table 2. Cell lysates were collected after 24 hours of treatment. GAPDH is shown as the loading control for each independent biological replicate. In each set of images, the membranes treated with GAPDH antibodies are shown in the top row, while those treated with c-Jun antibodies are shown in the bottom row. N = 4.

### Original Western Blot Images – Primary Antibody: Phospho -c-Jun

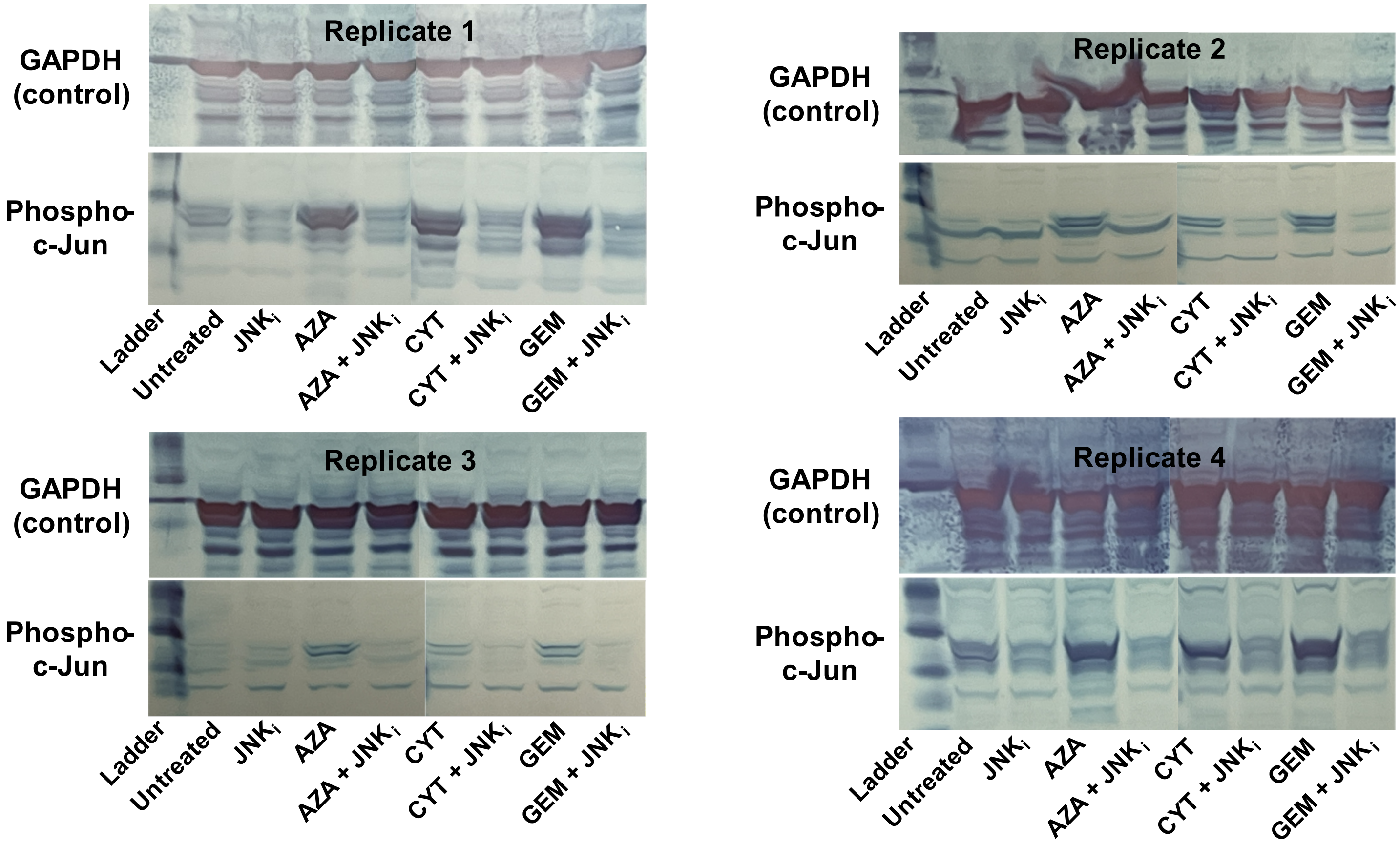


**Figure S5.** Original images collected during western blot experiments. In these experiments, the same treatment conditions (for A375 cells) from Figure S5 were used to compare phosphorylated (Serine 73) c-Jun levels. Treatment concentrations can be found in Supplemental Table 2. Cell lysates were collected after 24 hours of treatment. GAPDH is again shown as the loading control in the top row for each set of images, whereas the bottom row corresponds to membranes treated with phospho-c-Jun antibodies. N = 4.

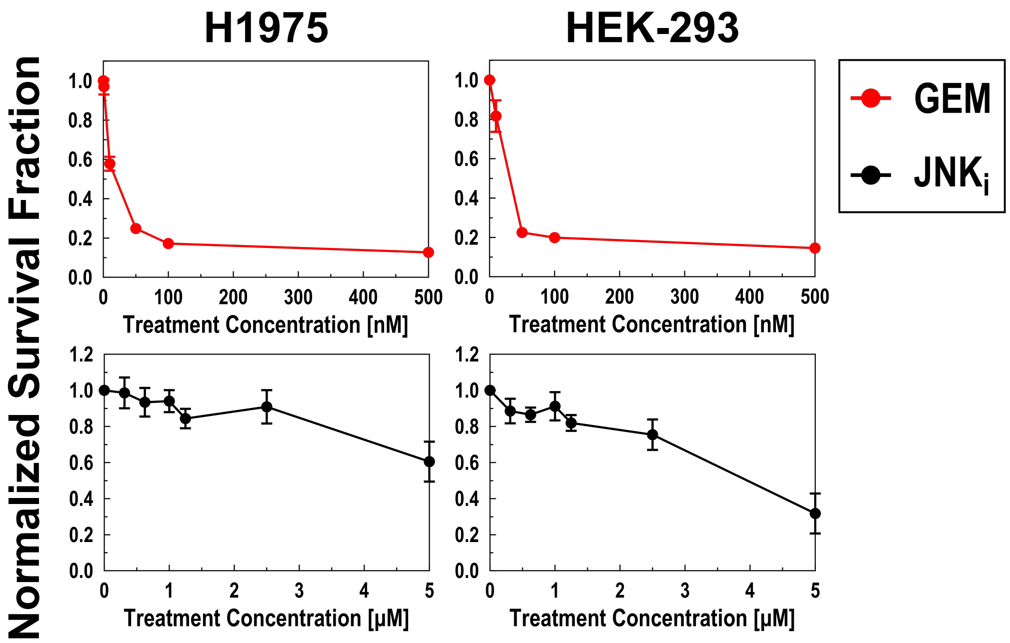


**Figure S6.** Concentration-dependent kill curves for additional (non-melanoma) cell lines. The listed cell lines were treated with varying concentrations of gemcitabine (GEM) and JNK-IN-8 ($JNK_i$). Survival fractions were measured by flow cytometry after 72 hours of treatment under the indicated conditions and normalized to the untreated cell number. A color code is used to distinguish each treatment. Treatment concentrations in other experiments were determined using these plots. N = 4.

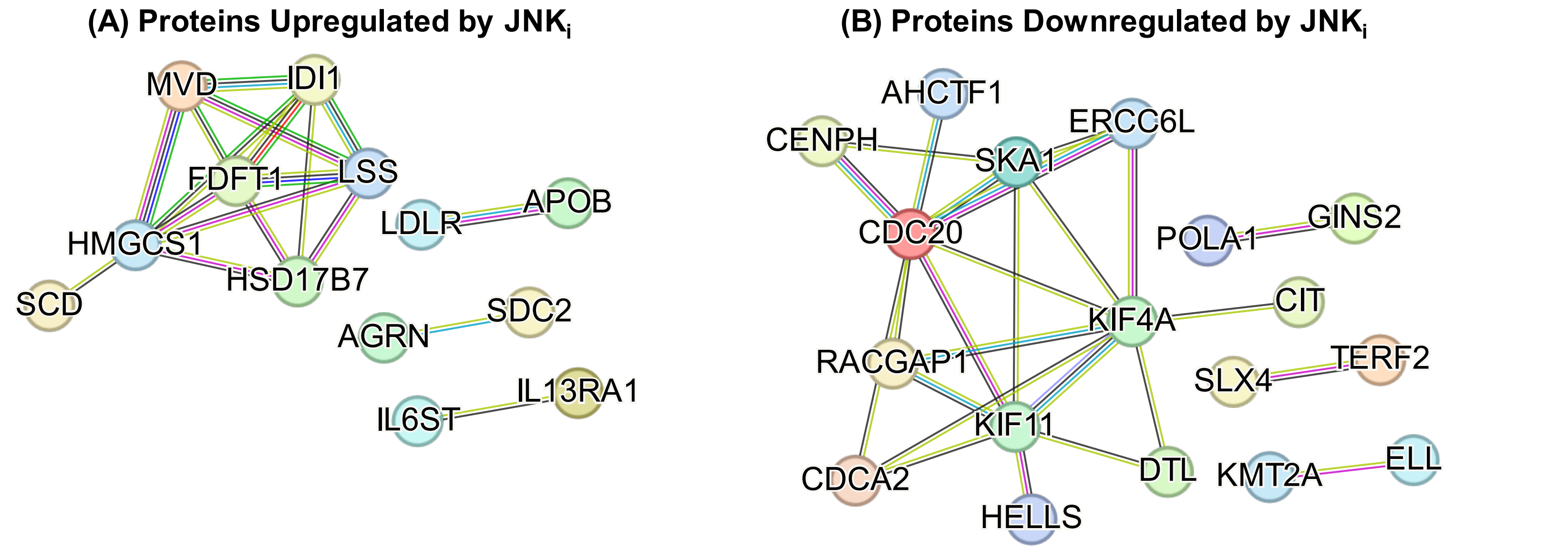


**Figure S7.** Untargeted proteomics analysis of A375 cells. Changes in protein expression were assessed upon 24 hours of treatment with 5 μM $JNK_i$. Significantly ($P<0.01$) (A) upregulated and (B) downregulated protein networks are presented. N = 3.

### Supplemental Tables

**Supplemental Table 1. Information on the cell lines used for this study.**

| Cell Line | Cell Type | Tumor Type/Location | Cell Morphology | Notable Oncogenic Mutations |
|---|---|---|---|---|
| **A375** | Human melanoma | In situ/skin | Epithelial | BRAF (V600E), CDKN2A |
| **RPMI-7951** | Human melanoma | Metastatic/lymph node | Epithelial | BRAF (V600E), PTEN, TP53 |
| **SH-4** | Human melanoma | Metastatic/pleural effusion | Spindle shaped, epithelial | BRAF (V600E) |
| **SK-Mel-3** | Human melanoma | Metastatic/lymph node | Fibroblast | BRAF (V600E), TP53 |
| **SK-Mel-24** | Human melanoma | Metastatic/lymph node | Stellate | BRAF (V600E), TP53 |
| **H1975** | Human non-small cell lung cancer | In situ/lung | Epithelial | EGFR, PIK3CA, TP53 |
| **HEK-293** | Human fetal kidney cell | N/A: non-cancerous | Epithelial | N/A: non-cancerous |

**Supplemental Table 2. Treatment concentrations used for each cell line.** These concentrations were used in the proteomics, immunodetection array, western blot, and co-treatment viability assays.

| Cell Line | Treatment Concentration | | | |
|---|---|---|---|---|
| | AZA | CYT | GEM | $JNK_i$ |
| **A375** | 100 µM | 500 nM | 20 nM | 5 µM |
| **RPMI-7951** | 35 µM | 6 µM | 35 nM | 1 µM |
| **SH-4** | 150 µM | 5 µM | 50 nM | 625 nM |
| **SK-Mel-3** | 70 µM | 700 nM | 50 nM | 1 µM |
| **SK-Mel-24** | 200 µM | 10 µM | 75 nM | 1 µM |
| **H1975** | | | 100 nM | 1 µM |
| **HEK-293** | | | 50 nM | 1 µM |

**Supplemental Table 3. Pathway Analysis for proteins upregulated by GEM treatment in A375 melanoma cells.** Statistical analysis was performed for the entire human genome, and the following pathways were identified. Protein networks and pathways were classified under several frameworks, including gene ontology annotations, KEGG pathways, reactome pathways, subcellular localization, and UniProt annotated keywords. **Count in Network:** the number of proteins identified as upregulated or downregulated in our samples, compared to the total number of proteins in the network. **Strength:** Log10(observed/expected). The strength quantifies the enrichment effect. It is the ratio between the number of proteins identified in a given network and the expected number in a random network with the same number of total proteins. **False Discovery Rate:** A measure of the significance of enrichment. P values are shown, corrected for multiple testing for each category with the Benjamini-Hochberg procedure.

**Biological Process (Gene Ontology)**

| GO-term | description | count in network | strength | false discovery rate |
|---|---|---|---|---|
| GO:0002486 | Antigen processing and presentation of endogenous peptid... | 3 of 3 | 1.65 | 0.0239 |
| GO:0034975 | Protein folding in endoplasmic reticulum | 7 of 11 | 1.46 | 4.78e-05 |
| GO:0090435 | Protein localization to nuclear envelope | 4 of 13 | 1.14 | 0.0417 |
| GO:0051231 | Spindle elongation | 4 of 13 | 1.14 | 0.0417 |
| GO:0044406 | Adhesion of symbiont to host | 4 of 13 | 1.14 | 0.0417 |
| GO:0051255 | Spindle midzone assembly | 4 of 14 | 1.11 | 0.0481 |
| GO:0036258 | Multivesicular body assembly | 7 of 32 | 0.99 | 0.0036 |
| GO:0039702 | Viral budding via host ESCRT complex | 5 of 23 | 0.99 | 0.0335 |
| GO:0007097 | Nuclear migration | 5 of 23 | 0.99 | 0.0335 |
| GO:0007080 | Mitotic metaphase plate congression | 11 of 53 | 0.97 | 7.23e-05 |
| GO:0019076 | Viral release from host cell | 6 of 31 | 0.94 | 0.0176 |
| GO:0051310 | Metaphase plate congression | 12 of 65 | 0.92 | 7.04e-05 |
| GO:0043162 | Ubiquitin-dependent protein catabolic process via the multiv... | 7 of 38 | 0.92 | 0.0079 |
| GO:0045324 | Late endosome to vacuole transport | 6 of 37 | 0.86 | 0.0335 |
| GO:0050000 | Chromosome localization | 13 of 84 | 0.84 | 9.90e-05 |
| GO:0060236 | Regulation of mitotic spindle organization | 7 of 45 | 0.84 | 0.0168 |
| GO:0032509 | Endosome transport via multivesicular body sorting pathway | 6 of 41 | 0.82 | 0.0480 |
| GO:0090307 | Mitotic spindle assembly | 7 of 51 | 0.79 | 0.0286 |
| GO:0051225 | Spindle assembly | 12 of 96 | 0.75 | 0.0012 |
| GO:0007052 | Mitotic spindle organization | 12 of 97 | 0.75 | 0.0012 |
| GO:0031110 | Regulation of microtubule polymerization or depolymerizati... | 11 of 88 | 0.75 | 0.0022 |
| GO:0000070 | Mitotic sister chromatid segregation | 15 of 122 | 0.74 | 0.00015 |
| GO:0000281 | Mitotic cytokinesis | 10 of 84 | 0.73 | 0.0065 |
| GO:0070507 | Regulation of microtubule cytoskeleton organization | 18 of 157 | 0.71 | 4.78e-05 |
| GO:0007098 | Centrosome cycle | 10 of 88 | 0.71 | 0.0086 |
| GO:0000819 | Sister chromatid segregation | 16 of 144 | 0.7 | 0.00019 |
| GO:0061077 | Chaperone-mediated protein folding | 8 of 71 | 0.7 | 0.0335 |
| GO:1902850 | Microtubule cytoskeleton organization involved in mitosis | 14 of 129 | 0.69 | 0.00094 |
| GO:0007051 | Spindle organization | 17 of 160 | 0.68 | 0.00016 |
| GO:0140014 | Mitotic nuclear division | 18 of 175 | 0.67 | 0.00013 |
| GO:0044000 | Movement in host | 14 of 139 | 0.66 | 0.0015 |
| GO:0061640 | Cytoskeleton-dependent cytokinesis | 11 of 112 | 0.65 | 0.0115 |
| GO:0051701 | Biological process involved in interaction with host | 15 of 157 | 0.63 | 0.0014 |
| GO:1901879 | Regulation of protein depolymerization | 9 of 94 | 0.63 | 0.0417 |
| GO:0010965 | Regulation of mitotic sister chromatid separation | 9 of 96 | 0.62 | 0.0458 |
| GO:0006457 | Protein folding | 19 of 215 | 0.6 | 0.00038 |
| GO:0032543 | Mitochondrial translation | 10 of 112 | 0.6 | 0.0350 |
| GO:0097191 | Extrinsic apoptotic signaling pathway | 10 of 113 | 0.6 | 0.0370 |
| GO:0016032 | Viral process | 21 of 242 | 0.59 | 0.00016 |

| | | | | |
|---|---|---|---|---|
| GO:0051983 | Regulation of chromosome segregation | 11 of 128 | 0.59 | 0.0270 |
| GO:0019058 | Viral life cycle | 15 of 178 | 0.58 | 0.0042 |
| GO:0006997 | Nucleus organization | 12 of 147 | 0.56 | 0.0238 |
| GO:0042770 | Signal transduction in response to DNA damage | 11 of 135 | 0.56 | 0.0370 |
| GO:0032984 | Protein-containing complex disassembly | 11 of 137 | 0.56 | 0.0408 |
| GO:0050684 | Regulation of mRNA processing | 11 of 140 | 0.55 | 0.0458 |
| GO:0032886 | Regulation of microtubule-based process | 19 of 261 | 0.52 | 0.0026 |
| GO:2000045 | Regulation of G1/S transition of mitotic cell cycle | 12 of 164 | 0.52 | 0.0460 |
| GO:1903047 | Mitotic cell cycle process | 39 of 537 | 0.51 | 7.73e-07 |
| GO:0033044 | Regulation of chromosome organization | 18 of 253 | 0.51 | 0.0053 |
| GO:0034504 | Protein localization to nucleus | 14 of 195 | 0.51 | 0.0245 |
| GO:0140694 | Non-membrane-bounded organelle assembly | 22 of 314 | 0.5 | 0.0013 |
| GO:0044403 | Biological process involved in symbiotic interaction | 18 of 257 | 0.5 | 0.0063 |
| GO:0051656 | Establishment of organelle localization | 26 of 380 | 0.49 | 0.00040 |
| GO:0008380 | RNA splicing | 25 of 370 | 0.48 | 0.00069 |
| GO:0007059 | Chromosome segregation | 19 of 286 | 0.48 | 0.0071 |
| GO:0000278 | Mitotic cell cycle | 41 of 631 | 0.47 | 4.27e-06 |
| GO:1901990 | Regulation of mitotic cell cycle phase transition | 22 of 332 | 0.47 | 0.0023 |
| GO:0010948 | Negative regulation of cell cycle process | 18 of 272 | 0.47 | 0.0107 |
| GO:0000398 | mRNA splicing, via spliceosome | 16 of 245 | 0.47 | 0.0245 |
| GO:0022402 | Cell cycle process | 54 of 835 | 0.46 | 2.76e-08 |
| GO:0010564 | Regulation of cell cycle process | 45 of 716 | 0.45 | 2.25e-06 |
| GO:0000226 | Microtubule cytoskeleton organization | 34 of 542 | 0.45 | 9.04e-05 |
| GO:0051301 | Cell division | 33 of 527 | 0.45 | 0.00013 |
| GO:1901987 | Regulation of cell cycle phase transition | 27 of 431 | 0.45 | 0.00096 |
| GO:0000280 | Nuclear division | 20 of 323 | 0.44 | 0.0104 |
| GO:0006397 | mRNA processing | 27 of 455 | 0.43 | 0.0018 |
| GO:0051493 | Regulation of cytoskeleton organization | 31 of 541 | 0.41 | 0.00098 |
| GO:0030036 | Actin cytoskeleton organization | 31 of 547 | 0.41 | 0.0012 |
| GO:0007346 | Regulation of mitotic cell cycle | 28 of 493 | 0.41 | 0.0024 |
| GO:0006417 | Regulation of translation | 26 of 456 | 0.41 | 0.0042 |
| GO:2001233 | Regulation of apoptotic signaling pathway | 21 of 365 | 0.41 | 0.0163 |
| GO:0007010 | Cytoskeleton organization | 69 of 1229 | 0.4 | 1.65e-08 |
| GO:0045786 | Negative regulation of cell cycle | 20 of 359 | 0.4 | 0.0293 |
| GO:0016071 | mRNA metabolic process | 33 of 611 | 0.39 | 0.0013 |
| GO:0010608 | Post-transcriptional regulation of gene expression | 30 of 546 | 0.39 | 0.0022 |
| GO:0010639 | Negative regulation of organelle organization | 19 of 351 | 0.39 | 0.0480 |
| GO:0007049 | Cell cycle | 66 of 1246 | 0.38 | 3.70e-07 |
| GO:0033365 | Protein localization to organelle | 37 of 713 | 0.37 | 0.00098 |
| GO:0030029 | Actin filament-based process | 32 of 609 | 0.37 | 0.0025 |
| GO:0051640 | Organelle localization | 27 of 514 | 0.37 | 0.0094 |
| GO:0034248 | Regulation of cellular amide metabolic process | 27 of 516 | 0.37 | 0.0098 |
| GO:1903829 | Positive regulation of protein localization | 24 of 461 | 0.37 | 0.0218 |
| GO:0033043 | Regulation of organelle organization | 61 of 1190 | 0.36 | 4.05e-06 |
| GO:0007017 | Microtubule-based process | 41 of 803 | 0.36 | 0.00051 |
| GO:0032990 | Cell part morphogenesis | 25 of 506 | 0.35 | 0.0304 |
| GO:0051726 | Regulation of cell cycle | 54 of 1108 | 0.34 | 7.20e-05 |
| GO:0006974 | Cellular response to DNA damage stimulus | 36 of 744 | 0.34 | 0.0038 |
| GO:0032989 | Cellular component morphogenesis | 29 of 603 | 0.34 | 0.0180 |
| GO:0120035 | Regulation of plasma membrane bounded cell projection or... | 30 of 635 | 0.33 | 0.0185 |
| GO:0010256 | Endomembrane system organization | 26 of 552 | 0.33 | 0.0417 |
| GO:0070925 | Organelle assembly | 37 of 799 | 0.32 | 0.0063 |
| GO:0080135 | Regulation of cellular response to stress | 33 of 712 | 0.32 | 0.0133 |
| GO:0051276 | Chromosome organization | 44 of 968 | 0.31 | 0.0022 |
| GO:0006396 | RNA processing | 39 of 868 | 0.31 | 0.0071 |
| GO:0061024 | Membrane organization | 35 of 773 | 0.31 | 0.0129 |
| GO:0060341 | Regulation of cellular localization | 43 of 963 | 0.3 | 0.0038 |
| GO:0031175 | Neuron projection development | 30 of 674 | 0.3 | 0.0389 |
| GO:0006886 | Intracellular protein transport | 30 of 683 | 0.3 | 0.0458 |
| GO:0015031 | Protein transport | 51 of 1180 | 0.29 | 0.0017 |
| GO:0051248 | Negative regulation of protein metabolic process | 45 of 1038 | 0.29 | 0.0045 |
| GO:0032880 | Regulation of protein localization | 37 of 852 | 0.29 | 0.0171 |
| GO:0045184 | Establishment of protein localization | 54 of 1272 | 0.28 | 0.0015 |

| | | | | |
|---|---|---|---|---|
| GO:0044087 | Regulation of cellular component biogenesis | 41 of 971 | 0.28 | 0.0137 |
| GO:0051128 | Regulation of cellular component organization | 97 of 2365 | 0.27 | 4.27e-06 |
| GO:0008104 | Protein localization | 80 of 1943 | 0.27 | 6.67e-05 |
| GO:0046907 | Intracellular transport | 57 of 1365 | 0.27 | 0.0014 |
| GO:0006996 | Organelle organization | 140 of 3470 | 0.26 | 2.66e-09 |
| GO:0051649 | Establishment of localization in cell | 71 of 1756 | 0.26 | 0.00040 |
| GO:0006915 | Apoptotic process | 42 of 1041 | 0.26 | 0.0249 |
| GO:0051641 | Cellular localization | 103 of 2677 | 0.24 | 2.23e-05 |
| GO:0090304 | Nucleic acid metabolic process | 86 of 2203 | 0.24 | 0.00013 |
| GO:0043933 | Protein-containing complex organization | 57 of 1465 | 0.24 | 0.0067 |
| GO:0030030 | Cell projection organization | 45 of 1158 | 0.24 | 0.0310 |
| GO:0008219 | Cell death | 43 of 1118 | 0.24 | 0.0458 |
| GO:0044085 | Cellular component biogenesis | 102 of 2702 | 0.23 | 5.44e-05 |
| GO:0022607 | Cellular component assembly | 94 of 2467 | 0.23 | 9.90e-05 |
| GO:0033036 | Macromolecule localization | 88 of 2356 | 0.23 | 0.00040 |
| GO:0016070 | RNA metabolic process | 59 of 1550 | 0.23 | 0.0084 |
| GO:0031399 | Regulation of protein modification process | 59 of 1560 | 0.23 | 0.0097 |
| GO:0033554 | Cellular response to stress | 59 of 1572 | 0.23 | 0.0113 |
| GO:0065003 | Protein-containing complex assembly | 49 of 1303 | 0.23 | 0.0335 |
| GO:0010467 | Gene expression | 78 of 2101 | 0.22 | 0.0015 |
| GO:0010941 | Regulation of cell death | 61 of 1651 | 0.22 | 0.0125 |
| GO:0071840 | Cellular component organization or biogenesis | 203 of 5639 | 0.21 | 1.03e-10 |
| GO:0016043 | Cellular component organization | 196 of 5436 | 0.21 | 2.12e-10 |
| GO:0034641 | Cellular nitrogen compound metabolic process | 125 of 3463 | 0.21 | 1.84e-05 |
| GO:0006139 | Nucleobase-containing compound metabolic process | 99 of 2722 | 0.21 | 0.00026 |
| GO:0051246 | Regulation of protein metabolic process | 94 of 2622 | 0.21 | 0.00078 |
| GO:0019220 | Regulation of phosphate metabolic process | 51 of 1405 | 0.21 | 0.0480 |
| GO:0046483 | Heterocycle metabolic process | 103 of 2891 | 0.2 | 0.00036 |
| GO:0006725 | Cellular aromatic compound metabolic process | 103 of 2936 | 0.2 | 0.00061 |
| GO:0051172 | Negative regulation of nitrogen compound metabolic process | 84 of 2403 | 0.2 | 0.0044 |
| GO:0031324 | Negative regulation of cellular metabolic process | 79 of 2265 | 0.2 | 0.0078 |
| GO:0071705 | Nitrogen compound transport | 56 of 1580 | 0.2 | 0.0458 |
| GO:0048522 | Positive regulation of cellular process | 186 of 5584 | 0.18 | 6.90e-07 |
| GO:0048523 | Negative regulation of cellular process | 158 of 4736 | 0.18 | 1.84e-05 |
| GO:1901360 | Organic cyclic compound metabolic process | 108 of 3181 | 0.18 | 0.0012 |
| GO:0044260 | Cellular macromolecule metabolic process | 85 of 2512 | 0.18 | 0.0100 |
| GO:0009892 | Negative regulation of metabolic process | 97 of 2982 | 0.17 | 0.0114 |
| GO:0010605 | Negative regulation of macromolecule metabolic process | 91 of 2760 | 0.17 | 0.0124 |
| GO:0048519 | Negative regulation of biological process | 171 of 5313 | 0.16 | 3.63e-05 |
| GO:0048518 | Positive regulation of biological process | 197 of 6207 | 0.15 | 4.61e-06 |
| GO:0051171 | Regulation of nitrogen compound metabolic process | 174 of 5734 | 0.14 | 0.00058 |
| GO:0010604 | Positive regulation of macromolecule metabolic process | 109 of 3533 | 0.14 | 0.0244 |
| GO:0031323 | Regulation of cellular metabolic process | 172 of 5681 | 0.13 | 0.00076 |
| GO:0080090 | Regulation of primary metabolic process | 176 of 5899 | 0.13 | 0.0012 |
| GO:0043170 | Macromolecule metabolic process | 173 of 5781 | 0.13 | 0.0012 |
| GO:0060255 | Regulation of macromolecule metabolic process | 185 of 6249 | 0.12 | 0.00096 |
| GO:0051234 | Establishment of localization | 117 of 3946 | 0.12 | 0.0480 |
| GO:0019222 | Regulation of metabolic process | 195 of 6784 | 0.11 | 0.0022 |
| GO:0006807 | Nitrogen compound metabolic process | 189 of 6643 | 0.11 | 0.0055 |
| GO:0044237 | Cellular metabolic process | 182 of 6568 | 0.1 | 0.0270 |
| GO:0044238 | Primary metabolic process | 195 of 7156 | 0.09 | 0.0325 |
| GO:0050789 | Regulation of biological process | 305 of 11655 | 0.07 | 0.0012 |
| GO:0050794 | Regulation of cellular process | 291 of 11025 | 0.07 | 0.0015 |
| GO:0009987 | Cellular process | 375 of 14826 | 0.06 | 7.04e-05 |
| GO:0065007 | Biological regulation | 313 of 12385 | 0.06 | 0.0148 |

### Molecular Function (Gene Ontology)

| GO-term | description | count in network | strength | false discovery rate |
|---|---|---|---|---|
| GO:0045296 | Cadherin binding | 35 of 334 | 0.67 | 3.53e-10 |
| GO:0031072 | Heat shock protein binding | 12 of 126 | 0.63 | 0.0204 |
| GO:0032182 | Ubiquitin-like protein binding | 11 of 117 | 0.63 | 0.0429 |
| GO:0050839 | Cell adhesion molecule binding | 46 of 560 | 0.57 | 2.82e-10 |
| GO:0003779 | Actin binding | 27 of 448 | 0.43 | 0.0027 |
| GO:0003723 | RNA binding | 96 of 1672 | 0.41 | 5.91e-14 |
| GO:0008092 | Cytoskeletal protein binding | 48 of 1002 | 0.33 | 0.00054 |
| GO:0044877 | Protein-containing complex binding | 55 of 1261 | 0.29 | 0.0012 |
| GO:0042802 | Identical protein binding | 90 of 2144 | 0.28 | 3.74e-06 |
| GO:0019899 | Enzyme binding | 87 of 2084 | 0.27 | 7.98e-06 |
| GO:0005515 | Protein binding | 250 of 7242 | 0.19 | 5.91e-14 |
| GO:0003676 | Nucleic acid binding | 139 of 4003 | 0.19 | 1.19e-05 |
| GO:0005488 | Binding | 355 of 12838 | 0.09 | 3.53e-10 |

### Cellular Component (Gene Ontology)

| GO-term | description | count in network | strength | false discovery rate |
|---|---|---|---|---|
| GO:0005880 | Nuclear microtubule | 2 of 2 | 1.65 | 0.0469 |
| GO:0042612 | MHC class I protein complex | 3 of 8 | 1.23 | 0.0299 |
| GO:0042824 | MHC class I peptide loading complex | 3 of 9 | 1.18 | 0.0375 |
| GO:0042406 | Extrinsic component of endoplasmic reticulum membrane | 3 of 9 | 1.18 | 0.0375 |
| GO:0000815 | ESCRT III complex | 3 of 10 | 1.13 | 0.0438 |
| GO:0031089 | Platelet dense granule lumen | 4 of 14 | 1.11 | 0.0125 |
| GO:0071556 | Integral component of lumenal side of endoplasmic reticulu... | 6 of 26 | 1.02 | 0.0021 |
| GO:0036452 | ESCRT complex | 6 of 27 | 1.0 | 0.0024 |
| GO:0035371 | Microtubule plus-end | 4 of 18 | 1.0 | 0.0260 |
| GO:0031941 | Filamentous actin | 6 of 29 | 0.97 | 0.0032 |
| GO:0042611 | MHC protein complex | 5 of 24 | 0.97 | 0.0094 |
| GO:0042641 | Actomyosin | 10 of 72 | 0.8 | 0.00050 |
| GO:0001725 | Stress fiber | 9 of 65 | 0.79 | 0.0013 |
| GO:0030018 | Z disc | 14 of 131 | 0.68 | 0.00017 |
| GO:0005876 | Spindle microtubule | 8 of 76 | 0.68 | 0.0116 |
| GO:0005762 | Mitochondrial large ribosomal subunit | 6 of 56 | 0.68 | 0.0411 |
| GO:0031674 | I band | 15 of 146 | 0.66 | 0.00012 |
| GO:0005761 | Mitochondrial ribosome | 9 of 90 | 0.65 | 0.0082 |
| GO:0030496 | Midbody | 19 of 201 | 0.63 | 2.20e-05 |
| GO:0055038 | Recycling endosome membrane | 9 of 94 | 0.63 | 0.0104 |
| GO:0030864 | Cortical actin cytoskeleton | 7 of 73 | 0.63 | 0.0347 |
| GO:0016363 | Nuclear matrix | 12 of 128 | 0.62 | 0.0021 |
| GO:0031970 | Organelle envelope lumen | 9 of 96 | 0.62 | 0.0116 |
| GO:0005758 | Mitochondrial intermembrane space | 8 of 86 | 0.62 | 0.0233 |
| GO:0005925 | Focal adhesion | 38 of 416 | 0.61 | 1.41e-10 |
| GO:0120111 | Neuron projection cytoplasm | 8 of 89 | 0.61 | 0.0267 |
| GO:0016528 | Sarcoplasm | 8 of 89 | 0.61 | 0.0267 |
| GO:0034399 | Nuclear periphery | 13 of 150 | 0.59 | 0.0021 |
| GO:0005884 | Actin filament | 8 of 99 | 0.56 | 0.0422 |
| GO:0015934 | Large ribosomal subunit | 9 of 113 | 0.55 | 0.0290 |
| GO:0070469 | Respirasome | 8 of 101 | 0.55 | 0.0457 |
| GO:0090734 | Site of DNA damage | 8 of 102 | 0.55 | 0.0472 |
| GO:0030863 | Cortical cytoskeleton | 8 of 102 | 0.55 | 0.0472 |
| GO:0005643 | Nuclear pore | 8 of 102 | 0.55 | 0.0472 |
| GO:0030017 | Sarcomere | 16 of 217 | 0.52 | 0.0021 |
| GO:0000779 | Condensed chromosome, centromeric region | 13 of 175 | 0.52 | 0.0070 |
| GO:0000775 | Chromosome, centromeric region | 18 of 252 | 0.51 | 0.0012 |
| GO:0072686 | Mitotic spindle | 13 of 180 | 0.51 | 0.0084 |
| GO:0000776 | Kinetochore | 12 of 165 | 0.51 | 0.0123 |
| GO:0044391 | Ribosomal subunit | 13 of 187 | 0.5 | 0.0113 |
| GO:0031902 | Late endosome membrane | 11 of 157 | 0.5 | 0.0260 |

| | | | | |
|---|---|---|---|---|
| GO:0000781 | Chromosome, telomeric region | 10 of 143 | 0.5 | 0.0375 |
| GO:0098687 | Chromosomal region | 25 of 365 | 0.49 | 9.12e-05 |
| GO:0015629 | Actin cytoskeleton | 32 of 482 | 0.48 | 7.54e-06 |
| GO:0098798 | Mitochondrial protein-containing complex | 20 of 296 | 0.48 | 0.00091 |
| GO:0005635 | Nuclear envelope | 32 of 487 | 0.47 | 9.05e-06 |
| GO:0000793 | Condensed chromosome | 18 of 275 | 0.47 | 0.0027 |
| GO:0005840 | Ribosome | 15 of 228 | 0.47 | 0.0080 |
| GO:0030027 | Lamellipodium | 13 of 203 | 0.46 | 0.0216 |
| GO:0031227 | Intrinsic component of endoplasmic reticulum membrane | 11 of 175 | 0.45 | 0.0454 |
| GO:0005912 | Adherens junction | 11 of 177 | 0.45 | 0.0472 |
| GO:0016607 | Nuclear speck | 26 of 420 | 0.44 | 0.00027 |
| GO:0005681 | Spliceosomal complex | 12 of 197 | 0.44 | 0.0399 |
| GO:0034774 | Secretory granule lumen | 19 of 321 | 0.43 | 0.0053 |
| GO:0005819 | Spindle | 25 of 425 | 0.42 | 0.00084 |
| GO:0031965 | Nuclear membrane | 18 of 305 | 0.42 | 0.0075 |
| GO:1990904 | Ribonucleoprotein complex | 39 of 687 | 0.41 | 1.35e-05 |
| GO:0005874 | Microtubule | 25 of 453 | 0.39 | 0.0021 |
| GO:0016604 | Nuclear body | 43 of 833 | 0.37 | 3.38e-05 |
| GO:0005813 | Centrosome | 32 of 609 | 0.37 | 0.00054 |
| GO:0099512 | Supramolecular fiber | 51 of 1000 | 0.36 | 5.13e-06 |
| GO:0099513 | Polymeric cytoskeletal fiber | 39 of 757 | 0.36 | 0.00011 |
| GO:0005788 | Endoplasmic reticulum lumen | 16 of 312 | 0.36 | 0.0422 |
| GO:0019866 | Organelle inner membrane | 28 of 566 | 0.35 | 0.0037 |
| GO:0010008 | Endosome membrane | 27 of 540 | 0.35 | 0.0041 |
| GO:0070161 | Anchoring junction | 64 of 1325 | 0.34 | 7.47e-07 |
| GO:0031967 | Organelle envelope | 62 of 1262 | 0.34 | 7.47e-07 |
| GO:0005730 | Nucleolus | 49 of 996 | 0.34 | 2.17e-05 |
| GO:0005815 | Microtubule organizing center | 40 of 825 | 0.34 | 0.00029 |
| GO:0070062 | Extracellular exosome | 99 of 2096 | 0.33 | 1.05e-10 |
| GO:0005856 | Cytoskeleton | 110 of 2369 | 0.32 | 1.38e-11 |
| GO:0015630 | Microtubule cytoskeleton | 63 of 1355 | 0.32 | 3.53e-06 |
| GO:0005911 | Cell-cell junction | 23 of 499 | 0.32 | 0.0260 |
| GO:0005769 | Early endosome | 19 of 411 | 0.32 | 0.0481 |
| GO:0099080 | Supramolecular complex | 62 of 1366 | 0.31 | 9.05e-06 |
| GO:0005743 | Mitochondrial inner membrane | 23 of 502 | 0.31 | 0.0267 |
| GO:0030141 | Secretory granule | 36 of 873 | 0.27 | 0.0104 |
| GO:0043232 | Intracellular non-membrane-bounded organelle | 206 of 5191 | 0.25 | 1.30e-17 |
| GO:0005654 | Nucleoplasm | 163 of 4169 | 0.25 | 4.06e-12 |
| GO:0005740 | Mitochondrial envelope | 32 of 802 | 0.25 | 0.0293 |
| GO:0070013 | Intracellular organelle lumen | 220 of 5660 | 0.24 | 4.00e-18 |
| GO:0031981 | Nuclear lumen | 173 of 4526 | 0.24 | 3.07e-12 |
| GO:0140513 | Nuclear protein-containing complex | 50 of 1290 | 0.24 | 0.0042 |
| GO:0030054 | Cell junction | 80 of 2115 | 0.23 | 0.00012 |
| GO:0005829 | Cytosol | 198 of 5438 | 0.21 | 1.28e-12 |
| GO:0030659 | Cytoplasmic vesicle membrane | 43 of 1190 | 0.21 | 0.0308 |
| GO:0099503 | Secretory vesicle | 38 of 1047 | 0.21 | 0.0462 |
| GO:0032991 | Protein-containing complex | 193 of 5506 | 0.2 | 1.01e-10 |
| GO:0005615 | Extracellular space | 111 of 3247 | 0.19 | 0.00011 |
| GO:0005694 | Chromosome | 63 of 1850 | 0.19 | 0.0133 |
| GO:0031982 | Vesicle | 129 of 3957 | 0.17 | 0.00013 |
| GO:0005634 | Nucleus | 247 of 7672 | 0.16 | 2.88e-11 |
| GO:0031410 | Cytoplasmic vesicle | 78 of 2482 | 0.15 | 0.0264 |
| GO:0005576 | Extracellular region | 127 of 4175 | 0.14 | 0.0029 |
| GO:0005737 | Cytoplasm | 355 of 12056 | 0.12 | 1.29e-16 |
| GO:0031090 | Organelle membrane | 107 of 3673 | 0.12 | 0.0347 |
| GO:0043226 | Organelle | 399 of 14017 | 0.11 | 2.87e-21 |
| GO:0043229 | Intracellular organelle | 381 of 13231 | 0.11 | 1.73e-18 |
| GO:0005622 | Intracellular anatomical structure | 413 of 14891 | 0.1 | 6.90e-22 |
| GO:0043227 | Membrane-bounded organelle | 365 of 13188 | 0.1 | 2.86e-12 |
| GO:0043231 | Intracellular membrane-bounded organelle | 343 of 12149 | 0.1 | 1.62e-11 |
| GO:0012505 | Endomembrane system | 133 of 4721 | 0.1 | 0.0299 |
| GO:0110165 | Cellular anatomical entity | 425 of 18293 | 0.02 | 0.0044 |

### Local network cluster (STRING)

| cluster | description | count in network | strength | false discovery rate |
|---|---|---|---|---|
| CL:14202 | ESCRT III complex, and ESCRT II complex | 4 of 8 | 1.35 | 0.0171 |
| CL:22848 | Filamentous actin, and EF-hand, Ca insensitive | 7 of 16 | 1.29 | 0.0018 |
| CL:14186 | ESCRT complex | 6 of 16 | 1.23 | 0.0024 |
| CL:2264 | Mixed, incl. SWAP/Surp, and TARP syndrome | 4 of 11 | 1.21 | 0.0361 |
| CL:6620 | Mixed, incl. Spindle elongation, and Axon hillock | 4 of 12 | 1.18 | 0.0439 |
| CL:6619 | Mixed, incl. Spindle elongation, and Outer kinetochore | 7 of 24 | 1.12 | 0.0022 |
| CL:22846 | Mixed, incl. Filamentous actin, and PDZ domain-containing ... | 8 of 28 | 1.11 | 0.0018 |
| CL:6617 | Mixed, incl. Spindle elongation, and Polo-like kinase mediate... | 8 of 30 | 1.08 | 0.0018 |
| CL:22845 | Mixed, incl. Filamentous actin, and Glomerulosclerosis | 9 of 44 | 0.96 | 0.0018 |
| CL:14181 | Multivesicular body assembly | 7 of 35 | 0.95 | 0.0088 |
| CL:14176 | Multivesicular body assembly, and BRO1 domain | 9 of 49 | 0.92 | 0.0022 |
| CL:1541 | Mixed, incl. Photodynamic therapy-induced unfolded protein... | 7 of 42 | 0.87 | 0.0171 |
| CL:6606 | Mitotic sister chromatid segregation, and Mitotic spindle ch... | 9 of 61 | 0.82 | 0.0066 |
| CL:25993 | Anchoring of the basal body to the plasma membrane, and ... | 9 of 61 | 0.82 | 0.0066 |
| CL:25994 | Mixed, incl. Centriole replication, and Centriolar subdistal ap... | 7 of 49 | 0.81 | 0.0320 |
| CL:2032 | U2-type spliceosomal complex, and mRNA Splicing - Major ... | 15 of 127 | 0.73 | 0.0018 |
| CL:2035 | U2-type spliceosomal complex, and mRNA Splicing - Major ... | 11 of 106 | 0.67 | 0.0120 |
| CL:560 | Mitochondrial translation initiation | 9 of 87 | 0.67 | 0.0361 |
| CL:6599 | Amplification of signal from the kinetochores, and Mitotic si... | 11 of 113 | 0.64 | 0.0171 |
| CL:6601 | Mixed, incl. Centromere, and Condensin complex | 10 of 104 | 0.64 | 0.0317 |
| CL:16224 | Mixed, incl. MHC classes I/II-like antigen recognition protein... | 10 of 109 | 0.62 | 0.0382 |

### KEGG Pathways

| pathway | description | count in network | strength | false discovery rate |
|---|---|---|---|---|
| hsa05330 | Allograft rejection | 6 of 34 | 0.9 | 0.0141 |
| hsa04612 | Antigen processing and presentation | 11 of 64 | 0.89 | 0.00021 |
| hsa05332 | Graft-versus-host disease | 6 of 36 | 0.87 | 0.0155 |
| hsa04940 | Type I diabetes mellitus | 6 of 38 | 0.85 | 0.0172 |
| hsa05416 | Viral myocarditis | 7 of 55 | 0.76 | 0.0172 |
| hsa05320 | Autoimmune thyroid disease | 6 of 48 | 0.75 | 0.0321 |
| hsa04115 | p53 signaling pathway | 7 of 72 | 0.64 | 0.0397 |
| hsa05169 | Epstein-Barr virus infection | 18 of 192 | 0.62 | 0.00021 |
| hsa04141 | Protein processing in endoplasmic reticulum | 15 of 163 | 0.62 | 0.00096 |
| hsa03040 | Spliceosome | 12 of 132 | 0.61 | 0.0063 |
| hsa04210 | Apoptosis | 10 of 131 | 0.54 | 0.0321 |
| hsa05166 | Human T-cell leukemia virus 1 infection | 14 of 210 | 0.48 | 0.0172 |
| hsa05131 | Shigellosis | 14 of 218 | 0.46 | 0.0198 |
| hsa04144 | Endocytosis | 14 of 241 | 0.42 | 0.0379 |

### Reactome Pathways

| pathway | description | count in network | strength | false discovery rate |
|---|---|---|---|---|
| HSA-9755779 | SARS-CoV-2 targets host intracellular signalling and regulat... | 4 of 12 | 1.18 | 0.0169 |
| HSA-9614399 | Regulation of localization of FOXO transcription factors | 4 of 12 | 1.18 | 0.0169 |
| HSA-75035 | Chk1/Chk2(Cds1) mediated inactivation of Cyclin B:Cdk1 co... | 4 of 13 | 1.14 | 0.0204 |
| HSA-9694676 | Translation of Replicase and Assembly of the Replication Tr... | 4 of 14 | 1.11 | 0.0233 |
| HSA-9735871 | SARS-CoV-1 targets host intracellular signalling and regulat... | 4 of 15 | 1.08 | 0.0284 |
| HSA-111447 | Activation of BAD and translocation to mitochondria | 4 of 15 | 1.08 | 0.0284 |
| HSA-983170 | Antigen Presentation: Folding, assembly and peptide loadin... | 7 of 29 | 1.04 | 0.0019 |
| HSA-174414 | Processive synthesis on the C-strand of the telomere | 4 of 19 | 0.98 | 0.0495 |
| HSA-9615710 | Late endosomal microautophagy | 7 of 34 | 0.97 | 0.0035 |
| HSA-917729 | Endosomal Sorting Complex Required For Transport (ESCRT) | 6 of 31 | 0.94 | 0.0099 |
| HSA-162588 | Budding and maturation of HIV virion | 5 of 28 | 0.9 | 0.0284 |
| HSA-199992 | trans-Golgi Network Vesicle Budding | 9 of 72 | 0.75 | 0.0062 |
| HSA-2995410 | Nuclear Envelope (NE) Reassembly | 9 of 74 | 0.74 | 0.0070 |
| HSA-3371453 | Regulation of HSF1-mediated heat shock response | 8 of 69 | 0.72 | 0.0159 |
| HSA-5389840 | Mitochondrial translation elongation | 10 of 88 | 0.71 | 0.0056 |

| | | | | |
|---|---|---|---|---|
| HSA-3371556 | Cellular response to heat stress | 10 of 89 | 0.7 | 0.0059 |
| HSA-8854518 | AURKA Activation by TPX2 | 8 of 72 | 0.7 | 0.0172 |
| HSA-5625740 | RHO GTPases activate PKNs | 7 of 63 | 0.7 | 0.0308 |
| HSA-2565942 | Regulation of PLK1 Activity at G2/M Transition | 9 of 87 | 0.67 | 0.0155 |
| HSA-5419276 | Mitochondrial translation termination | 9 of 88 | 0.66 | 0.0159 |
| HSA-5368286 | Mitochondrial translation initiation | 9 of 88 | 0.66 | 0.0159 |
| HSA-380259 | Loss of Nlp from mitotic centrosomes | 7 of 69 | 0.66 | 0.0486 |
| HSA-9610379 | HCMV Late Events | 8 of 81 | 0.65 | 0.0291 |
| HSA-114608 | Platelet degranulation | 12 of 126 | 0.63 | 0.0054 |
| HSA-5357801 | Programmed Cell Death | 19 of 206 | 0.62 | 0.00040 |
| HSA-109581 | Apoptosis | 16 of 175 | 0.61 | 0.0015 |
| HSA-9678108 | SARS-CoV-1 Infection | 12 of 140 | 0.59 | 0.0096 |
| HSA-9609646 | HCMV Infection | 11 of 126 | 0.59 | 0.0146 |
| HSA-72163 | mRNA Splicing - Major Pathway | 17 of 203 | 0.58 | 0.0015 |
| HSA-72203 | Processing of Capped Intron-Containing Pre-mRNA | 21 of 281 | 0.53 | 0.0015 |
| HSA-68882 | Mitotic Anaphase | 17 of 232 | 0.52 | 0.0044 |
| HSA-453279 | Mitotic G1 phase and G1/S transition | 11 of 148 | 0.52 | 0.0295 |
| HSA-162587 | HIV Life Cycle | 11 of 149 | 0.52 | 0.0300 |
| HSA-162599 | Late Phase of HIV Life Cycle | 10 of 136 | 0.52 | 0.0488 |
| HSA-69275 | G2/M Transition | 14 of 195 | 0.51 | 0.0146 |
| HSA-69278 | Cell Cycle, Mitotic | 37 of 526 | 0.5 | 2.92e-06 |
| HSA-9705683 | SARS-CoV-2-host interactions | 14 of 199 | 0.5 | 0.0155 |
| HSA-9694516 | SARS-CoV-2 Infection | 20 of 294 | 0.49 | 0.0030 |
| HSA-1640170 | Cell Cycle | 44 of 658 | 0.48 | 7.47e-07 |
| HSA-68886 | M Phase | 25 of 382 | 0.47 | 0.0015 |
| HSA-162906 | HIV Infection | 15 of 229 | 0.47 | 0.0166 |
| HSA-68877 | Mitotic Prometaphase | 13 of 201 | 0.46 | 0.0322 |
| HSA-3700989 | Transcriptional Regulation by TP53 | 21 of 361 | 0.42 | 0.0086 |
| HSA-72766 | Translation | 17 of 290 | 0.42 | 0.0205 |
| HSA-195258 | RHO GTPase Effectors | 17 of 292 | 0.42 | 0.0216 |
| HSA-6798695 | Neutrophil degranulation | 26 of 476 | 0.39 | 0.0051 |
| HSA-8953854 | Metabolism of RNA | 38 of 705 | 0.38 | 0.00060 |
| HSA-9012999 | RHO GTPase cycle | 24 of 449 | 0.38 | 0.0094 |
| HSA-9679506 | SARS-CoV Infections | 22 of 411 | 0.38 | 0.0150 |
| HSA-9716542 | Signaling by Rho GTPases, Miro GTPases and RHOBTB3 | 36 of 688 | 0.37 | 0.0015 |
| HSA-5653656 | Vesicle-mediated transport | 35 of 666 | 0.37 | 0.0015 |
| HSA-194315 | Signaling by Rho GTPases | 35 of 672 | 0.37 | 0.0015 |
| HSA-199991 | Membrane Trafficking | 32 of 626 | 0.36 | 0.0030 |
| HSA-5663205 | Infectious disease | 43 of 917 | 0.32 | 0.0015 |
| HSA-8953897 | Cellular responses to stimuli | 33 of 760 | 0.29 | 0.0164 |
| HSA-2262752 | Cellular responses to stress | 32 of 747 | 0.28 | 0.0216 |
| HSA-168249 | Innate Immune System | 40 of 1041 | 0.24 | 0.0298 |
| HSA-168256 | Immune System | 68 of 1979 | 0.19 | 0.0159 |
| HSA-392499 | Metabolism of proteins | 65 of 1917 | 0.18 | 0.0232 |

**Subcellular localization (COMPARTMENTS)**

| *compartment* | *description* | *count in network* | *strength* | *false discovery rate* |
|---|---|---|---|---|
| GOCC:0000814 | ESCRT II complex | 4 of 8 | 1.35 | 0.0037 |
| GOCC:0042824 | MHC class I peptide loading complex | 4 of 9 | 1.3 | 0.0052 |
| GOCC:0043626 | PCNA complex | 3 of 8 | 1.23 | 0.0379 |
| GOCC:0042612 | MHC class I protein complex | 3 of 8 | 1.23 | 0.0379 |
| GOCC:0031089 | Platelet dense granule lumen | 4 of 14 | 1.11 | 0.0155 |
| GOCC:0071556 | Integral component of lumenal side of endoplasmic reticulu... | 6 of 26 | 1.02 | 0.0024 |
| GOCC:0035371 | Microtubule plus-end | 4 of 17 | 1.02 | 0.0267 |
| GOCC:0042611 | MHC protein complex | 5 of 22 | 1.01 | 0.0089 |
| GOCC:0036452 | ESCRT complex | 6 of 28 | 0.98 | 0.0033 |
| GOCC:0005721 | Pericentric heterochromatin | 4 of 20 | 0.95 | 0.0407 |
| GOCC:0042641 | Actomyosin | 6 of 45 | 0.78 | 0.0221 |
| GOCC:0005876 | Spindle microtubule | 8 of 67 | 0.73 | 0.0078 |
| GOCC:0005925 | Focal adhesion | 31 of 269 | 0.71 | 1.10e-10 |
| GOCC:0055038 | Recycling endosome membrane | 7 of 61 | 0.71 | 0.0193 |

| | | | | |
|---|---|---|---|---|
| GOCC:0030496 | Midbody | 16 of 142 | 0.7 | 2.68e-05 |
| GOCC:0005758 | Mitochondrial intermembrane space | 6 of 55 | 0.69 | 0.0481 |
| GOCC:0072686 | Mitotic spindle | 14 of 133 | 0.68 | 0.00023 |
| GOCC:0005643 | Nuclear pore | 8 of 77 | 0.67 | 0.0155 |
| GOCC:0031902 | Late endosome membrane | 9 of 90 | 0.65 | 0.0101 |
| GOCC:0000775 | Chromosome, centromeric region | 15 of 160 | 0.62 | 0.00037 |
| GOCC:0031227 | Intrinsic component of endoplasmic reticulum membrane | 11 of 120 | 0.62 | 0.0052 |
| GOCC:0000779 | Condensed chromosome, centromeric region | 10 of 112 | 0.6 | 0.0106 |
| GOCC:0030176 | Integral component of endoplasmic reticulum membrane | 10 of 114 | 0.6 | 0.0118 |
| GOCC:0000776 | Kinetochore | 10 of 115 | 0.59 | 0.0124 |
| GOCC:0016607 | Nuclear speck | 16 of 193 | 0.57 | 0.00071 |
| GOCC:0005635 | Nuclear envelope | 25 of 313 | 0.56 | 1.02e-05 |
| GOCC:0031965 | Nuclear membrane | 15 of 184 | 0.56 | 0.0014 |
| GOCC:0005788 | Endoplasmic reticulum lumen | 14 of 172 | 0.56 | 0.0024 |
| GOCC:0070062 | Extracellular exosome | 33 of 428 | 0.54 | 2.59e-07 |
| GOCC:0005819 | Spindle | 22 of 289 | 0.53 | 8.51e-05 |
| GOCC:0098687 | Chromosomal region | 17 of 227 | 0.53 | 0.0012 |
| GOCC:0070161 | Anchoring junction | 40 of 553 | 0.51 | 2.98e-08 |
| GOCC:0055037 | Recycling endosome | 10 of 138 | 0.51 | 0.0379 |
| GOCC:0099513 | Polymeric cytoskeletal fiber | 26 of 382 | 0.49 | 7.08e-05 |
| GOCC:0015629 | Actin cytoskeleton | 24 of 363 | 0.47 | 0.00026 |
| GOCC:0098798 | Mitochondrial protein complex | 20 of 308 | 0.47 | 0.0016 |
| GOCC:0065010 | Extracellular membrane-bounded organelle | 30 of 473 | 0.46 | 4.79e-05 |
| GOCC:0010008 | Endosome membrane | 21 of 330 | 0.46 | 0.0014 |
| GOCC:0005874 | Microtubule | 15 of 233 | 0.46 | 0.0115 |
| GOCC:0000793 | Condensed chromosome | 13 of 205 | 0.46 | 0.0273 |
| GOCC:0005730 | Nucleolus | 38 of 606 | 0.45 | 2.97e-06 |
| GOCC:0016604 | Nuclear body | 24 of 381 | 0.45 | 0.00052 |
| GOCC:0005840 | Ribosome | 12 of 192 | 0.45 | 0.0423 |
| GOCC:0099512 | Supramolecular fiber | 33 of 555 | 0.43 | 5.02e-05 |
| GOCC:1990904 | Ribonucleoprotein complex | 33 of 564 | 0.42 | 6.82e-05 |
| GOCC:0034774 | Secretory granule lumen | 14 of 241 | 0.42 | 0.0379 |
| GOCC:0005813 | Centrosome | 27 of 470 | 0.41 | 0.00069 |
| GOCC:0031967 | Organelle envelope | 46 of 832 | 0.4 | 3.51e-06 |
| GOCC:0009986 | Cell surface | 23 of 438 | 0.37 | 0.0076 |
| GOCC:0005856 | Cytoskeleton | 80 of 1575 | 0.36 | 1.09e-09 |
| GOCC:0015630 | Microtubule cytoskeleton | 49 of 973 | 0.36 | 1.59e-05 |
| GOCC:0099080 | Supramolecular complex | 42 of 844 | 0.35 | 0.00012 |
| GOCC:0043232 | Intracellular non-membrane-bounded organelle | 160 of 3309 | 0.34 | 6.66e-20 |
| GOCC:0030054 | Cell junction | 51 of 1053 | 0.34 | 2.45e-05 |
| GOCC:0070013 | Intracellular organelle lumen | 139 of 2902 | 0.33 | 1.89e-16 |
| GOCC:0031981 | Nuclear lumen | 88 of 1850 | 0.33 | 2.14e-09 |
| GOCC:0005654 | Nucleoplasm | 54 of 1146 | 0.33 | 2.45e-05 |
| GOCC:0005694 | Chromosome | 45 of 951 | 0.33 | 0.00017 |
| GOCC:0005815 | Microtubule organizing center | 29 of 607 | 0.33 | 0.0057 |
| GOCC:0000323 | Lytic vacuole | 27 of 570 | 0.33 | 0.0099 |
| GOCC:0005764 | Lysosome | 26 of 566 | 0.32 | 0.0171 |
| GOCC:0000785 | Chromatin | 22 of 476 | 0.32 | 0.0374 |
| GOCC:0005829 | Cytosol | 134 of 3054 | 0.3 | 6.98e-13 |
| GOCC:0030141 | Secretory granule | 32 of 719 | 0.3 | 0.0086 |
| GOCC:0030659 | Cytoplasmic vesicle membrane | 32 of 723 | 0.3 | 0.0091 |
| GOCC:0005768 | Endosome | 29 of 671 | 0.29 | 0.0209 |
| GOCC:0005773 | Vacuole | 28 of 640 | 0.29 | 0.0212 |
| GOCC:0031982 | Vesicle | 84 of 2125 | 0.25 | 1.78e-05 |
| GOCC:0005615 | Extracellular space | 41 of 1027 | 0.25 | 0.0101 |
| GOCC:0005634 | Nucleus | 182 of 4787 | 0.23 | 6.10e-13 |
| GOCC:0031410 | Cytoplasmic vesicle | 66 of 1738 | 0.23 | 0.00091 |
| GOCC:0005739 | Mitochondrion | 45 of 1182 | 0.23 | 0.0128 |

| | | | | |
|---|---|---|---|---|
| GOCC:0043229 | Intracellular organelle | 335 of 9609 | 0.2 | 7.23e-29 |
| GOCC:0031090 | Organelle membrane | 80 of 2290 | 0.2 | 0.0017 |
| GOCC:0043226 | Organelle | 345 of 10113 | 0.19 | 7.23e-29 |
| GOCC:0043231 | Intracellular membrane-bounded organelle | 280 of 8162 | 0.19 | 1.95e-18 |
| GOCC:0043227 | Membrane-bounded organelle | 304 of 9083 | 0.18 | 7.68e-20 |
| GOCC:0032991 | Protein-containing complex | 181 of 5325 | 0.18 | 1.50e-08 |
| GOCC:0012505 | Endomembrane system | 106 of 3156 | 0.18 | 0.00044 |
| GOCC:0005737 | Cytoplasm | 268 of 8195 | 0.17 | 5.27e-14 |
| GOCC:0005622 | Intracellular | 374 of 11512 | 0.16 | 1.97e-30 |
| GOCC:0110165 | Cellular anatomical entity | 390 of 14060 | 0.1 | 1.89e-16 |

**Annotated Keywords (UniProt)**

| keyword | description | count in network | strength | false discovery rate |
|---|---|---|---|---|
| KW-0490 | MHC I | 3 of 8 | 1.23 | 0.0369 |
| KW-0440 | LIM domain | 14 of 70 | 0.95 | 2.30e-07 |
| KW-0143 | Chaperone | 20 of 197 | 0.66 | 3.89e-06 |
| KW-0747 | Spliceosome | 11 of 138 | 0.55 | 0.0125 |
| KW-0540 | Nuclease | 10 of 127 | 0.55 | 0.0225 |
| KW-0508 | mRNA splicing | 20 of 274 | 0.52 | 0.00034 |
| KW-0493 | Microtubule | 20 of 280 | 0.51 | 0.00043 |
| KW-0498 | Mitosis | 19 of 275 | 0.49 | 0.00092 |
| KW-0689 | Ribosomal protein | 12 of 175 | 0.49 | 0.0225 |
| KW-0132 | Cell division | 26 of 384 | 0.48 | 6.45e-05 |
| KW-0131 | Cell cycle | 42 of 651 | 0.46 | 1.94e-07 |
| KW-0007 | Acetylation | 201 of 3362 | 0.43 | 1.13e-40 |
| KW-1017 | Isopeptide bond | 102 of 1717 | 0.43 | 2.77e-17 |
| KW-0507 | mRNA processing | 21 of 349 | 0.43 | 0.0023 |
| KW-0206 | Cytoskeleton | 72 of 1235 | 0.42 | 2.63e-11 |
| KW-0945 | Host-virus interaction | 31 of 540 | 0.41 | 0.00015 |
| KW-0687 | Ribonucleoprotein | 16 of 278 | 0.41 | 0.0213 |
| KW-0832 | Ubl conjugation | 133 of 2399 | 0.4 | 5.49e-21 |
| KW-0053 | Apoptosis | 29 of 534 | 0.39 | 0.00071 |
| KW-0488 | Methylation | 50 of 973 | 0.36 | 4.28e-06 |
| KW-0694 | RNA-binding | 34 of 686 | 0.35 | 0.00079 |
| KW-0653 | Protein transport | 30 of 617 | 0.34 | 0.0029 |
| KW-0175 | Coiled coil | 102 of 2166 | 0.33 | 4.25e-11 |
| KW-0597 | Phosphoprotein | 329 of 8122 | 0.26 | 3.49e-43 |
| KW-0965 | Cell junction | 32 of 809 | 0.25 | 0.0392 |
| KW-0963 | Cytoplasm | 196 of 5095 | 0.24 | 3.79e-15 |
| KW-0539 | Nucleus | 175 of 5278 | 0.17 | 1.94e-07 |
| KW-0025 | Alternative splicing | 267 of 10313 | 0.07 | 0.0062 |

**Supplemental Table 4. Pathway Analysis for proteins downregulated by GEM treatment in A375 melanoma cells.** See Table 3 for details.

**Biological Process (Gene Ontology)**

| GO-term | description | count in network | strength | false discovery rate |
|---|---|---|---|---|
| GO:0000301 | Retrograde transport, vesicle recycling within Golgi | 4 of 9 | 1.26 | 0.0177 |
| GO:0071044 | Histone mRNA catabolic process | 4 of 11 | 1.17 | 0.0278 |
| GO:0007000 | Nucleolus organization | 5 of 15 | 1.14 | 0.0093 |
| GO:1905214 | Regulation of RNA binding | 4 of 13 | 1.1 | 0.0417 |
| GO:0006418 | tRNA aminoacylation for protein translation | 11 of 43 | 1.02 | 1.17e-05 |
| GO:0008334 | Histone mRNA metabolic process | 5 of 20 | 1.01 | 0.0224 |
| GO:0032211 | Negative regulation of telomere maintenance via telomerase | 5 of 22 | 0.97 | 0.0301 |
| GO:0006270 | DNA replication initiation | 6 of 29 | 0.93 | 0.0153 |
| GO:1904357 | Negative regulation of telomere maintenance via telomere l... | 6 of 30 | 0.91 | 0.0174 |
| GO:0032205 | Negative regulation of telomere maintenance | 7 of 38 | 0.88 | 0.0093 |
| GO:0000959 | Mitochondrial RNA metabolic process | 8 of 44 | 0.87 | 0.0037 |
| GO:0006890 | Retrograde vesicle-mediated transport, Golgi to endoplasmi... | 9 of 51 | 0.86 | 0.0016 |
| GO:0002183 | Cytoplasmic translational initiation | 6 of 34 | 0.86 | 0.0271 |
| GO:2000279 | Negative regulation of DNA biosynthetic process | 7 of 41 | 0.85 | 0.0129 |
| GO:0006378 | mRNA polyadenylation | 6 of 35 | 0.85 | 0.0301 |
| GO:0006891 | intra-Golgi vesicle-mediated transport | 6 of 38 | 0.81 | 0.0410 |
| GO:0000462 | Maturation of SSU-rRNA from tricistronic rRNA transcript (S... | 6 of 39 | 0.8 | 0.0454 |
| GO:0032210 | Regulation of telomere maintenance via telomerase | 8 of 55 | 0.78 | 0.0120 |
| GO:0030490 | Maturation of SSU-rRNA | 8 of 55 | 0.78 | 0.0120 |
| GO:2001251 | Negative regulation of chromosome organization | 12 of 84 | 0.77 | 0.00048 |
| GO:1904356 | Regulation of telomere maintenance via telomere lengthening | 9 of 64 | 0.76 | 0.0067 |
| GO:0006413 | Translational initiation | 10 of 75 | 0.74 | 0.0041 |
| GO:0031124 | mRNA 3-end processing | 7 of 53 | 0.73 | 0.0395 |
| GO:0031057 | Negative regulation of histone modification | 7 of 53 | 0.73 | 0.0395 |
| GO:0051053 | Negative regulation of DNA metabolic process | 19 of 149 | 0.72 | 5.48e-06 |
| GO:0032508 | DNA duplex unwinding | 11 of 89 | 0.71 | 0.0034 |
| GO:0006275 | Regulation of DNA replication | 16 of 136 | 0.68 | 0.00014 |
| GO:0042274 | Ribosomal small subunit biogenesis | 9 of 78 | 0.68 | 0.0201 |
| GO:0042254 | Ribosome biogenesis | 34 of 299 | 0.67 | 6.94e-10 |
| GO:2000278 | Regulation of DNA biosynthetic process | 14 of 124 | 0.67 | 0.00085 |
| GO:0006399 | tRNA metabolic process | 22 of 199 | 0.66 | 4.72e-06 |
| GO:0006401 | RNA catabolic process | 18 of 164 | 0.65 | 7.85e-05 |
| GO:0016239 | Positive regulation of macroautophagy | 8 of 74 | 0.65 | 0.0497 |
| GO:0000956 | Nuclear-transcribed mRNA catabolic process | 11 of 103 | 0.64 | 0.0094 |
| GO:0016072 | rRNA metabolic process | 26 of 250 | 0.63 | 1.02e-06 |
| GO:1903008 | Organelle disassembly | 11 of 105 | 0.63 | 0.0106 |
| GO:0031123 | RNA 3-end processing | 11 of 107 | 0.63 | 0.0120 |
| GO:0006261 | DNA-templated DNA replication | 14 of 137 | 0.62 | 0.0022 |
| GO:0006402 | mRNA catabolic process | 13 of 127 | 0.62 | 0.0038 |
| GO:0006364 | rRNA processing | 22 of 220 | 0.61 | 2.05e-05 |
| GO:0030071 | Regulation of mitotic metaphase/anaphase transition | 9 of 91 | 0.61 | 0.0444 |
| GO:0022613 | Ribonucleoprotein complex biogenesis | 44 of 449 | 0.6 | 4.05e-11 |
| GO:0001824 | Blastocyst development | 11 of 114 | 0.6 | 0.0183 |
| GO:0000910 | Cytokinesis | 11 of 114 | 0.6 | 0.0183 |
| GO:0032204 | Regulation of telomere maintenance | 10 of 103 | 0.6 | 0.0289 |
| GO:0006403 | RNA localization | 17 of 180 | 0.59 | 0.00081 |
| GO:0006260 | DNA replication | 19 of 203 | 0.58 | 0.00029 |
| GO:1905818 | Regulation of chromosome separation | 10 of 109 | 0.58 | 0.0403 |
| GO:0051168 | Nuclear export | 12 of 132 | 0.57 | 0.0166 |
| GO:0000018 | Regulation of DNA recombination | 12 of 132 | 0.57 | 0.0166 |

| | | | | |
|---|---|---|---|---|
| GO:0045931 | Positive regulation of mitotic cell cycle | 11 of 121 | 0.57 | 0.0259 |
| GO:0034660 | ncRNA metabolic process | 47 of 535 | 0.56 | 1.94e-10 |
| GO:0061640 | Cytoskeleton-dependent cytokinesis | 10 of 112 | 0.56 | 0.0468 |
| GO:0051236 | Establishment of RNA localization | 14 of 161 | 0.55 | 0.0088 |
| GO:0006888 | Endoplasmic reticulum to Golgi vesicle-mediated transport | 11 of 126 | 0.55 | 0.0336 |
| GO:0097193 | Intrinsic apoptotic signaling pathway | 14 of 166 | 0.54 | 0.0111 |
| GO:0006412 | Translation | 32 of 389 | 0.53 | 2.60e-06 |
| GO:0048193 | Golgi vesicle transport | 24 of 292 | 0.53 | 0.00012 |
| GO:0033044 | Regulation of chromosome organization | 21 of 253 | 0.53 | 0.00047 |
| GO:0050658 | RNA transport | 13 of 159 | 0.53 | 0.0217 |
| GO:0006997 | Nucleus organization | 12 of 147 | 0.53 | 0.0332 |
| GO:0051052 | Regulation of DNA metabolic process | 44 of 541 | 0.52 | 9.84e-09 |
| GO:0034470 | ncRNA processing | 33 of 409 | 0.52 | 2.55e-06 |
| GO:0031056 | Regulation of histone modification | 15 of 188 | 0.52 | 0.0112 |
| GO:0007051 | Spindle organization | 13 of 160 | 0.52 | 0.0224 |
| GO:0006913 | Nucleocytoplasmic transport | 19 of 248 | 0.5 | 0.0031 |
| GO:0016071 | mRNA metabolic process | 45 of 611 | 0.48 | 9.19e-08 |
| GO:0051301 | Cell division | 39 of 527 | 0.48 | 1.13e-06 |
| GO:0006396 | RNA processing | 62 of 868 | 0.47 | 1.54e-10 |
| GO:0006417 | Regulation of translation | 33 of 456 | 0.47 | 2.05e-05 |
| GO:0006520 | Cellular amino acid metabolic process | 21 of 290 | 0.47 | 0.0027 |
| GO:0071826 | Ribonucleoprotein complex subunit organization | 15 of 211 | 0.47 | 0.0279 |
| GO:0043604 | Amide biosynthetic process | 38 of 536 | 0.46 | 4.27e-06 |
| GO:2000112 | Regulation of cellular macromolecule biosynthetic process | 38 of 539 | 0.46 | 4.72e-06 |
| GO:0006397 | mRNA processing | 32 of 455 | 0.46 | 5.24e-05 |
| GO:0022618 | Ribonucleoprotein complex assembly | 14 of 203 | 0.45 | 0.0497 |
| GO:0034248 | Regulation of cellular amide metabolic process | 34 of 516 | 0.43 | 8.87e-05 |
| GO:0010639 | Negative regulation of organelle organization | 23 of 351 | 0.43 | 0.0043 |
| GO:0022411 | Cellular component disassembly | 21 of 321 | 0.43 | 0.0086 |
| GO:0006338 | Chromatin remodeling | 20 of 303 | 0.43 | 0.0106 |
| GO:0034655 | Nucleobase-containing compound catabolic process | 19 of 287 | 0.43 | 0.0141 |
| GO:0016070 | RNA metabolic process | 99 of 1550 | 0.42 | 4.29e-15 |
| GO:0044265 | Cellular macromolecule catabolic process | 50 of 786 | 0.42 | 8.31e-07 |
| GO:0006281 | DNA repair | 32 of 497 | 0.42 | 0.00027 |
| GO:0043161 | Proteasome-mediated ubiquitin-dependent protein catabolic... | 22 of 347 | 0.42 | 0.0089 |
| GO:0090304 | Nucleic acid metabolic process | 137 of 2203 | 0.41 | 3.32e-21 |
| GO:0051276 | Chromosome organization | 61 of 968 | 0.41 | 2.23e-08 |
| GO:0006259 | DNA metabolic process | 49 of 785 | 0.41 | 1.98e-06 |
| GO:0010608 | Post-transcriptional regulation of gene expression | 34 of 546 | 0.41 | 0.00026 |
| GO:0006518 | Peptide metabolic process | 34 of 547 | 0.41 | 0.00026 |
| GO:0008380 | RNA splicing | 23 of 370 | 0.41 | 0.0084 |
| GO:1901990 | Regulation of mitotic cell cycle phase transition | 21 of 332 | 0.41 | 0.0120 |
| GO:0072594 | Establishment of protein localization to organelle | 21 of 335 | 0.41 | 0.0134 |
| GO:1903311 | Regulation of mRNA metabolic process | 19 of 302 | 0.41 | 0.0224 |
| GO:0051054 | Positive regulation of DNA metabolic process | 19 of 304 | 0.41 | 0.0239 |
| GO:0048511 | Rhythmic process | 17 of 270 | 0.41 | 0.0403 |
| GO:0009057 | Macromolecule catabolic process | 61 of 989 | 0.4 | 4.74e-08 |
| GO:1903047 | Mitotic cell cycle process | 33 of 537 | 0.4 | 0.00045 |
| GO:0001701 | In utero embryonic development | 24 of 393 | 0.4 | 0.0077 |
| GO:0046700 | Heterocycle catabolic process | 20 of 327 | 0.4 | 0.0224 |
| GO:0031647 | Regulation of protein stability | 20 of 328 | 0.4 | 0.0227 |
| GO:0006325 | Chromatin organization | 37 of 612 | 0.39 | 0.00018 |
| GO:0019439 | Aromatic compound catabolic process | 21 of 349 | 0.39 | 0.0206 |
| GO:0044270 | Cellular nitrogen compound catabolic process | 20 of 333 | 0.39 | 0.0266 |
| GO:0006914 | Autophagy | 18 of 303 | 0.39 | 0.0497 |
| GO:0010467 | Gene expression | 124 of 2101 | 0.38 | 3.80e-17 |
| GO:0045184 | Establishment of protein localization | 74 of 1272 | 0.38 | 6.39e-09 |
| GO:0016570 | Histone modification | 23 of 396 | 0.38 | 0.0178 |
| GO:0006139 | Nucleobase-containing compound metabolic process | 155 of 2722 | 0.37 | 4.84e-21 |
| GO:0000278 | Mitotic cell cycle | 36 of 631 | 0.37 | 0.00073 |

| | | | | |
|---|---|---|---|---|
| GO:0043632 | Modification-dependent macromolecule catabolic process | 34 of 601 | 0.37 | 0.0015 |
| GO:0044260 | Cellular macromolecule metabolic process | 141 of 2512 | 0.36 | 3.24e-18 |
| GO:0007049 | Cell cycle | 69 of 1246 | 0.36 | 1.94e-07 |
| GO:0006974 | Cellular response to DNA damage stimulus | 42 of 744 | 0.36 | 0.00018 |
| GO:0033365 | Protein localization to organelle | 40 of 713 | 0.36 | 0.00034 |
| GO:0051603 | Proteolysis involved in protein catabolic process | 36 of 649 | 0.36 | 0.0012 |
| GO:1901987 | Regulation of cell cycle phase transition | 24 of 431 | 0.36 | 0.0218 |
| GO:1901361 | Organic cyclic compound catabolic process | 21 of 377 | 0.36 | 0.0428 |
| GO:0046483 | Heterocycle metabolic process | 159 of 2891 | 0.35 | 2.48e-20 |
| GO:0006725 | Cellular aromatic compound metabolic process | 159 of 2936 | 0.35 | 1.06e-19 |
| GO:0015031 | Protein transport | 65 of 1180 | 0.35 | 7.51e-07 |
| GO:0022402 | Cell cycle process | 46 of 835 | 0.35 | 0.00011 |
| GO:0043603 | Cellular amide metabolic process | 44 of 810 | 0.35 | 0.00025 |
| GO:0030163 | Protein catabolic process | 41 of 755 | 0.35 | 0.00052 |
| GO:0006886 | Intracellular protein transport | 37 of 683 | 0.35 | 0.0015 |
| GO:0019941 | Modification-dependent protein catabolic process | 32 of 589 | 0.35 | 0.0047 |
| GO:0010638 | Positive regulation of organelle organization | 28 of 508 | 0.35 | 0.0100 |
| GO:0007346 | Regulation of mitotic cell cycle | 27 of 493 | 0.35 | 0.0134 |
| GO:0034645 | Cellular macromolecule biosynthetic process | 41 of 778 | 0.34 | 0.00096 |
| GO:0006511 | Ubiquitin-dependent protein catabolic process | 31 of 578 | 0.34 | 0.0074 |
| GO:1901360 | Organic cyclic compound metabolic process | 164 of 3181 | 0.33 | 2.09e-18 |
| GO:0044271 | Cellular nitrogen compound biosynthetic process | 77 of 1494 | 0.33 | 3.59e-07 |
| GO:0031329 | Regulation of cellular catabolic process | 41 of 789 | 0.33 | 0.0013 |
| GO:0051129 | Negative regulation of cellular component organization | 36 of 691 | 0.33 | 0.0038 |
| GO:0034641 | Cellular nitrogen compound metabolic process | 178 of 3463 | 0.32 | 2.39e-20 |
| GO:0010256 | Endomembrane system organization | 28 of 552 | 0.32 | 0.0273 |
| GO:0009896 | Positive regulation of catabolic process | 26 of 513 | 0.32 | 0.0403 |
| GO:0070727 | Cellular macromolecule localization | 97 of 1948 | 0.31 | 1.18e-08 |
| GO:0071705 | Nitrogen compound transport | 79 of 1580 | 0.31 | 7.51e-07 |
| GO:0010564 | Regulation of cell cycle process | 36 of 716 | 0.31 | 0.0069 |
| GO:0008104 | Protein localization | 95 of 1943 | 0.3 | 4.56e-08 |
| GO:0044248 | Cellular catabolic process | 79 of 1620 | 0.3 | 1.98e-06 |
| GO:0046907 | Intracellular transport | 67 of 1365 | 0.3 | 1.85e-05 |
| GO:0009059 | Macromolecule biosynthetic process | 68 of 1394 | 0.3 | 1.85e-05 |
| GO:1901566 | Organonitrogen compound biosynthetic process | 65 of 1338 | 0.3 | 3.81e-05 |
| GO:0009792 | Embryo development ending in birth or egg hatching | 33 of 676 | 0.3 | 0.0184 |
| GO:0043009 | Chordate embryonic development | 32 of 654 | 0.3 | 0.0215 |
| GO:0033036 | Macromolecule localization | 113 of 2356 | 0.29 | 1.84e-09 |
| GO:0033554 | Cellular response to stress | 75 of 1572 | 0.29 | 8.91e-06 |
| GO:0043933 | Protein-containing complex organization | 70 of 1465 | 0.29 | 2.30e-05 |
| GO:0033043 | Regulation of organelle organization | 56 of 1190 | 0.29 | 0.00056 |
| GO:0009894 | Regulation of catabolic process | 47 of 988 | 0.29 | 0.0024 |
| GO:0043069 | Negative regulation of programmed cell death | 42 of 911 | 0.28 | 0.0100 |
| GO:0043170 | Macromolecule metabolic process | 263 of 5781 | 0.27 | 1.52e-26 |
| GO:0006996 | Organelle organization | 158 of 3470 | 0.27 | 1.02e-12 |
| GO:0051641 | Cellular localization | 122 of 2677 | 0.27 | 4.96e-09 |
| GO:0009056 | Catabolic process | 89 of 1976 | 0.27 | 5.26e-06 |
| GO:0043066 | Negative regulation of apoptotic process | 40 of 891 | 0.27 | 0.0210 |
| GO:1901575 | Organic substance catabolic process | 75 of 1686 | 0.26 | 0.00010 |
| GO:0051649 | Establishment of localization in cell | 77 of 1756 | 0.26 | 0.00012 |
| GO:0065003 | Protein-containing complex assembly | 58 of 1303 | 0.26 | 0.0016 |
| GO:0051726 | Regulation of cell cycle | 49 of 1108 | 0.26 | 0.0076 |
| GO:0044237 | Cellular metabolic process | 287 of 6568 | 0.25 | 1.99e-27 |
| GO:0006915 | Apoptotic process | 45 of 1041 | 0.25 | 0.0196 |
| GO:0060548 | Negative regulation of cell death | 44 of 1016 | 0.25 | 0.0216 |
| GO:0018130 | Heterocycle biosynthetic process | 43 of 985 | 0.25 | 0.0216 |
| GO:0034654 | Nucleobase-containing compound biosynthetic process | 40 of 913 | 0.25 | 0.0295 |
| GO:0070647 | Protein modification by small protein conjugation or removal | 39 of 902 | 0.25 | 0.0409 |
| GO:0006807 | Nitrogen compound metabolic process | 281 of 6643 | 0.24 | 1.26e-24 |
| GO:0051246 | Regulation of protein metabolic process | 111 of 2622 | 0.24 | 2.04e-06 |

| | | | | |
|---|---|---|---|---|
| GO:1901576 | Organic substance biosynthetic process | 104 of 2438 | 0.24 | 4.27e-06 |
| GO:0009058 | Biosynthetic process | 106 of 2506 | 0.24 | 4.41e-06 |
| GO:0044249 | Cellular biosynthetic process | 101 of 2362 | 0.24 | 5.78e-06 |
| GO:0071702 | Organic substance transport | 82 of 1957 | 0.24 | 0.00025 |
| GO:0019438 | Aromatic compound biosynthetic process | 42 of 992 | 0.24 | 0.0392 |
| GO:0010243 | Response to organonitrogen compound | 41 of 963 | 0.24 | 0.0403 |
| GO:0044238 | Primary metabolic process | 298 of 7156 | 0.23 | 4.81e-26 |
| GO:0019538 | Protein metabolic process | 162 of 3910 | 0.23 | 9.30e-10 |
| GO:0043412 | Macromolecule modification | 119 of 2898 | 0.23 | 2.55e-06 |
| GO:0044085 | Cellular component biogenesis | 113 of 2702 | 0.23 | 2.59e-06 |
| GO:0031324 | Negative regulation of cellular metabolic process | 93 of 2265 | 0.23 | 0.00011 |
| GO:0045934 | Negative regulation of nucleobase-containing compound m... | 64 of 1562 | 0.23 | 0.0054 |
| GO:1901565 | Organonitrogen compound catabolic process | 46 of 1113 | 0.23 | 0.0361 |
| GO:0008219 | Cell death | 46 of 1118 | 0.23 | 0.0388 |
| GO:0008152 | Metabolic process | 320 of 7988 | 0.22 | 2.00e-26 |
| GO:0071704 | Organic substance metabolic process | 305 of 7522 | 0.22 | 4.11e-25 |
| GO:0036211 | Protein modification process | 107 of 2674 | 0.22 | 4.43e-05 |
| GO:0006508 | Proteolysis | 51 of 1247 | 0.22 | 0.0241 |
| GO:0071840 | Cellular component organization or biogenesis | 223 of 5639 | 0.21 | 2.36e-13 |
| GO:0016192 | Vesicle-mediated transport | 51 of 1298 | 0.21 | 0.0489 |
| GO:0051128 | Regulation of cellular component organization | 91 of 2365 | 0.2 | 0.0015 |
| GO:1901564 | Organonitrogen compound metabolic process | 188 of 4981 | 0.19 | 1.65e-08 |
| GO:0051172 | Negative regulation of nitrogen compound metabolic process | 90 of 2403 | 0.19 | 0.0038 |
| GO:0045935 | Positive regulation of nucleobase-containing compound me... | 78 of 2056 | 0.19 | 0.0086 |
| GO:0006796 | Phosphate-containing compound metabolic process | 71 of 1877 | 0.19 | 0.0173 |
| GO:0044093 | Positive regulation of molecular function | 60 of 1587 | 0.19 | 0.0432 |
| GO:0031327 | Negative regulation of cellular biosynthetic process | 60 of 1592 | 0.19 | 0.0460 |
| GO:0031399 | Regulation of protein modification process | 59 of 1560 | 0.19 | 0.0468 |
| GO:0016043 | Cellular component organization | 200 of 5436 | 0.18 | 2.23e-08 |
| GO:0009892 | Negative regulation of metabolic process | 110 of 2982 | 0.18 | 0.00092 |
| GO:0010605 | Negative regulation of macromolecule metabolic process | 101 of 2760 | 0.18 | 0.0030 |
| GO:0031323 | Regulation of cellular metabolic process | 205 of 5681 | 0.17 | 5.42e-08 |
| GO:0031325 | Positive regulation of cellular metabolic process | 111 of 3114 | 0.17 | 0.0031 |
| GO:0080090 | Regulation of primary metabolic process | 208 of 5899 | 0.16 | 2.69e-07 |
| GO:0019219 | Regulation of nucleobase-containing compound metabolic ... | 143 of 4074 | 0.16 | 0.00036 |
| GO:0051173 | Positive regulation of nitrogen compound metabolic process | 112 of 3166 | 0.16 | 0.0038 |
| GO:0022607 | Cellular component assembly | 86 of 2467 | 0.16 | 0.0376 |
| GO:0051171 | Regulation of nitrogen compound metabolic process | 199 of 5734 | 0.15 | 2.60e-06 |
| GO:0009889 | Regulation of biosynthetic process | 145 of 4205 | 0.15 | 0.00073 |
| GO:0010556 | Regulation of macromolecule biosynthetic process | 136 of 3980 | 0.15 | 0.0023 |
| GO:0019222 | Regulation of metabolic process | 226 of 6784 | 0.14 | 3.47e-06 |
| GO:0060255 | Regulation of macromolecule metabolic process | 208 of 6249 | 0.14 | 2.12e-05 |
| GO:0051179 | Localization | 153 of 4512 | 0.14 | 0.00085 |
| GO:0031326 | Regulation of cellular biosynthetic process | 140 of 4143 | 0.14 | 0.0028 |
| GO:0009893 | Positive regulation of metabolic process | 130 of 3847 | 0.14 | 0.0056 |
| GO:0065009 | Regulation of molecular function | 104 of 3085 | 0.14 | 0.0295 |
| GO:0048523 | Negative regulation of cellular process | 154 of 4736 | 0.13 | 0.0058 |
| GO:0051234 | Establishment of localization | 131 of 3946 | 0.13 | 0.0101 |
| GO:0010604 | Positive regulation of macromolecule metabolic process | 117 of 3533 | 0.13 | 0.0239 |
| GO:0048519 | Negative regulation of biological process | 166 of 5313 | 0.11 | 0.0167 |
| GO:0009987 | Cellular process | 435 of 14826 | 0.08 | 1.37e-14 |

### Molecular Function (Gene Ontology)

| GO-term | description | count in network | strength | false discovery rate |
|---|---|---|---|---|
| GO:0016433 | rRNA (adenine) methyltransferase activity | 4 of 8 | 1.31 | 0.0125 |
| GO:0004652 | Polynucleotide adenylyltransferase activity | 4 of 13 | 1.1 | 0.0468 |
| GO:0004812 | aminoacyl-tRNA ligase activity | 11 of 43 | 1.02 | 6.79e-06 |
| GO:0140658 | ATP-dependent chromatin remodeler activity | 7 of 34 | 0.93 | 0.0046 |
| GO:0003724 | RNA helicase activity | 15 of 76 | 0.91 | 6.11e-07 |
| GO:0070566 | Adenylyltransferase activity | 6 of 31 | 0.9 | 0.0192 |
| GO:0004386 | Helicase activity | 26 of 149 | 0.85 | 1.70e-11 |
| GO:0003678 | DNA helicase activity | 10 of 63 | 0.81 | 0.00098 |
| GO:0008094 | ATP-dependent activity, acting on DNA | 17 of 110 | 0.8 | 1.35e-06 |
| GO:0003743 | Translation initiation factor activity | 8 of 57 | 0.76 | 0.0134 |
| GO:0016887 | ATP hydrolysis activity | 46 of 333 | 0.75 | 2.19e-17 |
| GO:0140101 | Catalytic activity, acting on a tRNA | 17 of 131 | 0.73 | 1.21e-05 |
| GO:0016874 | Ligase activity | 21 of 168 | 0.71 | 8.07e-07 |
| GO:0008135 | Translation factor activity, RNA binding | 12 of 98 | 0.7 | 0.0013 |
| GO:0090079 | Translation regulator activity, nucleic acid binding | 13 of 121 | 0.64 | 0.0020 |
| GO:0140657 | ATP-dependent activity | 55 of 543 | 0.62 | 1.61e-15 |
| GO:0140098 | Catalytic activity, acting on RNA | 39 of 380 | 0.62 | 7.02e-11 |
| GO:0140640 | Catalytic activity, acting on a nucleic acid | 57 of 575 | 0.61 | 9.98e-16 |
| GO:0045182 | Translation regulator activity | 15 of 152 | 0.61 | 0.0013 |
| GO:0017111 | Nucleoside-triphosphatase activity | 61 of 650 | 0.59 | 7.26e-16 |
| GO:0140097 | Catalytic activity, acting on DNA | 20 of 221 | 0.57 | 0.00017 |
| GO:0016462 | Pyrophosphatase activity | 62 of 705 | 0.56 | 5.90e-15 |
| GO:0042393 | Histone binding | 20 of 252 | 0.51 | 0.0010 |
| GO:0043021 | Ribonucleoprotein complex binding | 12 of 152 | 0.51 | 0.0463 |
| GO:0005524 | ATP binding | 116 of 1491 | 0.5 | 1.54e-25 |
| GO:0035639 | Purine ribonucleoside triphosphate binding | 137 of 1834 | 0.49 | 2.29e-28 |
| GO:0032559 | Adenyl ribonucleotide binding | 118 of 1554 | 0.49 | 3.24e-25 |
| GO:0032555 | Purine ribonucleotide binding | 139 of 1903 | 0.48 | 2.29e-28 |
| GO:0000166 | Nucleotide binding | 150 of 2168 | 0.45 | 2.29e-28 |
| GO:0003723 | RNA binding | 115 of 1672 | 0.45 | 3.67e-21 |
| GO:0031267 | Small GTPase binding | 18 of 273 | 0.43 | 0.0197 |
| GO:0043168 | Anion binding | 153 of 2404 | 0.42 | 3.67e-26 |
| GO:0106310 | Protein serine kinase activity | 23 of 361 | 0.42 | 0.0052 |
| GO:0036094 | Small molecule binding | 158 of 2507 | 0.41 | 9.49e-27 |
| GO:0097367 | Carbohydrate derivative binding | 142 of 2278 | 0.41 | 4.14e-23 |
| GO:0005525 | GTP binding | 24 of 381 | 0.41 | 0.0044 |
| GO:0051020 | GTPase binding | 19 of 304 | 0.41 | 0.0239 |
| GO:0003682 | Chromatin binding | 34 of 584 | 0.38 | 0.00067 |
| GO:0004674 | Protein serine/threonine kinase activity | 24 of 434 | 0.36 | 0.0237 |
| GO:0016787 | Hydrolase activity | 114 of 2347 | 0.3 | 1.73e-10 |
| GO:0003676 | Nucleic acid binding | 189 of 4003 | 0.29 | 1.13e-18 |
| GO:0016773 | Phosphotransferase activity, alcohol group as acceptor | 32 of 692 | 0.28 | 0.0483 |
| GO:1901363 | Heterocyclic compound binding | 270 of 5977 | 0.27 | 2.62e-28 |
| GO:0016772 | Transferase activity, transferring phosphorus-containing gro... | 42 of 930 | 0.27 | 0.0134 |
| GO:0097159 | Organic cyclic compound binding | 270 of 6050 | 0.26 | 1.88e-27 |
| GO:0003824 | Catalytic activity | 241 of 5522 | 0.25 | 1.35e-21 |
| GO:0140096 | Catalytic activity, acting on a protein | 94 of 2279 | 0.23 | 5.00e-05 |
| GO:0016740 | Transferase activity | 93 of 2268 | 0.23 | 7.33e-05 |
| GO:0019899 | Enzyme binding | 87 of 2084 | 0.23 | 8.82e-05 |
| GO:0043167 | Ion binding | 222 of 6033 | 0.18 | 1.73e-10 |
| GO:0042802 | Identical protein binding | 78 of 2144 | 0.17 | 0.0237 |
| GO:0005515 | Protein binding | 235 of 7242 | 0.12 | 6.79e-06 |
| GO:0005488 | Binding | 386 of 12838 | 0.09 | 5.29e-11 |

| Cellular Component (Gene Ontology) | | | | |
|---|---|---|---|---|
| GO-term | description | count in network | strength | false discovery rate |
| GO:0005827 | Polar microtubule | 3 of 5 | 1.39 | 0.0260 |
| GO:0005851 | Eukaryotic translation initiation factor 2B complex | 3 of 6 | 1.31 | 0.0323 |
| GO:0031261 | DNA replication preinitiation complex | 4 of 12 | 1.14 | 0.0200 |
| GO:0017119 | Golgi transport complex | 4 of 13 | 1.1 | 0.0245 |
| GO:0042645 | Mitochondrial nucleoid | 13 of 46 | 1.06 | 1.02e-07 |
| GO:0033290 | Eukaryotic 48S preinitiation complex | 4 of 15 | 1.04 | 0.0305 |
| GO:0005852 | Eukaryotic translation initiation factor 3 complex | 4 of 15 | 1.04 | 0.0305 |
| GO:0000177 | Cytoplasmic exosome (RNase complex) | 4 of 16 | 1.01 | 0.0353 |
| GO:0016282 | Eukaryotic 43S preinitiation complex | 4 of 17 | 0.98 | 0.0408 |
| GO:0000159 | Protein phosphatase type 2A complex | 4 of 17 | 0.98 | 0.0408 |
| GO:0005963 | Magnesium-dependent protein serine/threonine phosphatas... | 6 of 29 | 0.93 | 0.0076 |
| GO:1990391 | DNA repair complex | 8 of 43 | 0.88 | 0.0016 |
| GO:0099023 | Vesicle tethering complex | 10 of 65 | 0.8 | 0.00080 |
| GO:0032040 | Small-subunit processome | 6 of 39 | 0.8 | 0.0256 |
| GO:0008287 | Protein serine/threonine phosphatase complex | 8 of 56 | 0.77 | 0.0065 |
| GO:0042470 | Melanosome | 12 of 109 | 0.65 | 0.0018 |
| GO:0030684 | Preribosome | 8 of 76 | 0.64 | 0.0298 |
| GO:0000118 | Histone deacetylase complex | 8 of 80 | 0.61 | 0.0360 |
| GO:0000781 | Chromosome, telomeric region | 14 of 143 | 0.6 | 0.0016 |
| GO:1904813 | ficolin-1-rich granule lumen | 12 of 124 | 0.6 | 0.0050 |
| GO:0000228 | Nuclear chromosome | 23 of 244 | 0.59 | 1.05e-05 |
| GO:1904949 | ATPase complex | 9 of 96 | 0.59 | 0.0303 |
| GO:0101002 | ficolin-1-rich granule | 16 of 185 | 0.55 | 0.0016 |
| GO:0030496 | Midbody | 17 of 201 | 0.54 | 0.0013 |
| GO:0140534 | Endoplasmic reticulum protein-containing complex | 11 of 129 | 0.54 | 0.0226 |
| GO:0098687 | Chromosomal region | 30 of 365 | 0.53 | 2.36e-06 |
| GO:0000779 | Condensed chromosome, centromeric region | 13 of 175 | 0.48 | 0.0245 |
| GO:0005730 | Nucleolus | 71 of 996 | 0.47 | 3.42e-13 |
| GO:0000793 | Condensed chromosome | 20 of 275 | 0.47 | 0.0017 |
| GO:0000776 | Kinetochore | 12 of 165 | 0.47 | 0.0353 |
| GO:0000775 | Chromosome, centromeric region | 17 of 252 | 0.44 | 0.0119 |
| GO:0140513 | Nuclear protein-containing complex | 85 of 1290 | 0.43 | 3.39e-14 |
| GO:0005819 | Spindle | 27 of 425 | 0.42 | 0.00088 |
| GO:0031983 | Vesicle lumen | 19 of 326 | 0.38 | 0.0256 |
| GO:1902494 | Catalytic complex | 88 of 1539 | 0.37 | 2.11e-11 |
| GO:0140535 | Intracellular protein-containing complex | 45 of 784 | 0.37 | 1.99e-05 |
| GO:0034774 | Secretory granule lumen | 18 of 321 | 0.36 | 0.0408 |
| GO:1990904 | Ribonucleoprotein complex | 36 of 687 | 0.33 | 0.0016 |
| GO:0031981 | Nuclear lumen | 233 of 4526 | 0.32 | 3.42e-30 |
| GO:0005654 | Nucleoplasm | 213 of 4169 | 0.32 | 1.06e-26 |
| GO:0005759 | Mitochondrial matrix | 25 of 494 | 0.32 | 0.0270 |
| GO:0005813 | Centrosome | 30 of 609 | 0.31 | 0.0152 |
| GO:1990234 | Transferase complex | 41 of 847 | 0.3 | 0.0022 |
| GO:0016604 | Nuclear body | 40 of 833 | 0.29 | 0.0031 |
| GO:0070013 | Intracellular organelle lumen | 263 of 5660 | 0.28 | 3.65e-29 |
| GO:0032991 | Protein-containing complex | 248 of 5506 | 0.27 | 9.39e-25 |
| GO:0005739 | Mitochondrion | 76 of 1681 | 0.27 | 1.92e-05 |
| GO:0005829 | Cytosol | 238 of 5438 | 0.25 | 1.68e-21 |
| GO:0015630 | Microtubule cytoskeleton | 58 of 1355 | 0.24 | 0.0018 |
| GO:0005815 | Microtubule organizing center | 35 of 825 | 0.24 | 0.0426 |
| GO:0043232 | Intracellular non-membrane-bounded organelle | 214 of 5191 | 0.23 | 2.50e-15 |
| GO:0005694 | Chromosome | 77 of 1850 | 0.23 | 0.00030 |
| GO:0005634 | Nucleus | 293 of 7672 | 0.2 | 6.48e-20 |
| GO:0070062 | Extracellular exosome | 75 of 2096 | 0.17 | 0.0245 |
| GO:0043231 | Intracellular membrane-bounded organelle | 403 of 12149 | 0.13 | 8.38e-24 |
| GO:0031090 | Organelle membrane | 118 of 3673 | 0.12 | 0.0298 |
| GO:0043229 | Intracellular organelle | 418 of 13231 | 0.11 | 4.24e-21 |
| GO:0043227 | Membrane-bounded organelle | 417 of 13188 | 0.11 | 4.89e-21 |
| GO:0005737 | Cytoplasm | 381 of 12056 | 0.11 | 3.01e-15 |
| GO:0005622 | Intracellular anatomical structure | 459 of 14891 | 0.1 | 2.43e-29 |
| GO:0043226 | Organelle | 431 of 14017 | 0.1 | 1.37e-20 |
| GO:0110165 | Cellular anatomical entity | 469 of 18293 | 0.02 | 0.00019 |

### Local network cluster (STRING)

| cluster | description | count in network | strength | false discovery rate |
|---|---|---|---|---|
| CL:13256 | Retrograde transport, vesicle recycling within Golgi | 4 of 8 | 1.31 | 0.0362 |
| CL:1384 | Mitochondrial RNA metabolic process, and Regulation of mi... | 8 of 27 | 1.08 | 0.00076 |
| CL:1385 | Mixed, incl. Mitochondrial transcription, and RAP domain | 6 of 21 | 1.07 | 0.0111 |
| CL:454 | Initiation factor | 8 of 36 | 0.96 | 0.0034 |
| CL:1035 | rRNA modification in the nucleus and cytosol | 10 of 49 | 0.92 | 0.00076 |
| CL:795 | tRNA Aminoacylation, and glutamyl-tRNA(Gln) amidotransfe... | 9 of 48 | 0.89 | 0.0034 |
| CL:797 | tRNA Aminoacylation | 8 of 42 | 0.89 | 0.0082 |
| CL:1037 | Small-subunit processome, and Positive regulation of rRNA ... | 7 of 44 | 0.81 | 0.0491 |
| CL:6455 | DNA replication, and Regulation of DNA-directed DNA polym... | 9 of 60 | 0.79 | 0.0111 |
| CL:895 | rRNA processing, and RNA helicase activity | 26 of 185 | 0.76 | 3.28e-08 |
| CL:897 | rRNA processing, and Ribosomal large subunit biogenesis | 24 of 179 | 0.74 | 2.56e-07 |
| CL:902 | Preribosome, and Ribosome biogenesis | 19 of 144 | 0.73 | 1.91e-05 |
| CL:907 | Preribosome, and Ribosome biogenesis | 16 of 121 | 0.73 | 0.00013 |
| CL:905 | Preribosome, and Ribosome biogenesis | 17 of 133 | 0.72 | 0.00010 |
| CL:13130 | Golgi vesicle transport, and SNARE complex | 14 of 154 | 0.57 | 0.0146 |

### KEGG Pathways

| pathway | description | count in network | strength | false discovery rate |
|---|---|---|---|---|
| hsa00970 | Aminoacyl-tRNA biosynthesis | 10 of 44 | 0.97 | 0.00017 |
| hsa03430 | Mismatch repair | 5 of 23 | 0.95 | 0.0261 |
| hsa01212 | Fatty acid metabolism | 7 of 54 | 0.73 | 0.0261 |
| hsa04110 | Cell cycle | 14 of 120 | 0.68 | 0.00065 |
| hsa03008 | Ribosome biogenesis in eukaryotes | 8 of 77 | 0.63 | 0.0315 |
| hsa03013 | RNA transport | 16 of 161 | 0.61 | 0.00065 |
| hsa04914 | Progesterone-mediated oocyte maturation | 9 of 95 | 0.59 | 0.0315 |
| hsa04114 | Oocyte meiosis | 11 of 121 | 0.57 | 0.0216 |
| hsa05203 | Viral carcinogenesis | 15 of 183 | 0.53 | 0.0070 |
| hsa05132 | Salmonella infection | 14 of 209 | 0.44 | 0.0315 |
| hsa05016 | Huntington disease | 18 of 295 | 0.4 | 0.0261 |

### Reactome Pathways

| pathway | description | count in network | strength | false discovery rate |
|---|---|---|---|---|
| HSA-5423599 | Diseases of Mismatch Repair (MMR) | 3 of 5 | 1.39 | 0.0179 |
| HSA-72731 | Recycling of eIF2:GDP | 3 of 8 | 1.19 | 0.0367 |
| HSA-379716 | Cytosolic tRNA aminoacylation | 8 of 24 | 1.14 | 6.11e-05 |
| HSA-68884 | Mitotic Telophase/Cytokinesis | 4 of 13 | 1.1 | 0.0160 |
| HSA-5358508 | Mismatch Repair | 4 of 15 | 1.04 | 0.0224 |
| HSA-176412 | Phosphorylation of the APC/C | 5 of 20 | 1.01 | 0.0085 |
| HSA-379724 | tRNA Aminoacylation | 10 of 42 | 0.99 | 3.90e-05 |
| HSA-432142 | Platelet sensitization by LDL | 4 of 17 | 0.98 | 0.0297 |
| HSA-6804114 | TP53 Regulates Transcription of Genes Involved in G2 Cell C... | 4 of 18 | 0.96 | 0.0335 |
| HSA-9013422 | RHOBTB1 GTPase cycle | 5 of 23 | 0.95 | 0.0136 |
| HSA-9013418 | RHOBTB2 GTPase cycle | 5 of 23 | 0.95 | 0.0136 |
| HSA-5673000 | RAF activation | 7 of 34 | 0.93 | 0.0022 |
| HSA-9706574 | RHOBTB GTPase Cycle | 7 of 35 | 0.91 | 0.0025 |
| HSA-5620916 | VxPx cargo-targeting to cilium | 4 of 21 | 0.89 | 0.0484 |
| HSA-6790901 | rRNA modification in the nucleus and cytosol | 10 of 60 | 0.84 | 0.00037 |
| HSA-6811440 | Retrograde transport at the Trans-Golgi-Network | 8 of 49 | 0.83 | 0.0026 |
| HSA-6811438 | Intra-Golgi traffic | 7 of 43 | 0.82 | 0.0064 |
| HSA-176187 | Activation of ATR in response to replication stress | 6 of 37 | 0.82 | 0.0141 |
| HSA-69190 | DNA strand elongation | 5 of 32 | 0.81 | 0.0350 |
| HSA-8953750 | Transcriptional Regulation by E2F6 | 5 of 34 | 0.78 | 0.0410 |
| HSA-69618 | Mitotic Spindle Checkpoint | 16 of 111 | 0.77 | 8.46e-06 |
| HSA-5620920 | Cargo trafficking to the periciliary membrane | 7 of 50 | 0.76 | 0.0116 |
| HSA-2500257 | Resolution of Sister Chromatid Cohesion | 17 of 125 | 0.75 | 7.50e-06 |

| | | | | |
|---|---|---|---|---|
| HSA-141444 | Amplification of signal from unattached kinetochores via a ... | 13 of 94 | 0.75 | 0.00012 |
| HSA-9648025 | EML4 and NUDC in mitotic spindle formation | 15 of 116 | 0.72 | 5.90e-05 |
| HSA-6807878 | COPI-mediated anterograde transport | 13 of 101 | 0.72 | 0.00022 |
| HSA-1655829 | Regulation of cholesterol biosynthesis by SREBP (SREBF) | 7 of 55 | 0.72 | 0.0174 |
| HSA-2467813 | Separation of Sister Chromatids | 23 of 189 | 0.7 | 6.92e-07 |
| HSA-69052 | Switching of origins to a post-replicative state | 11 of 91 | 0.7 | 0.0016 |
| HSA-6791312 | TP53 Regulates Transcription of Cell Cycle Genes | 6 of 49 | 0.7 | 0.0367 |
| HSA-176814 | Activation of APC/C and APC/C:Cdc20 mediated degradatio... | 9 of 76 | 0.69 | 0.0066 |
| HSA-9675135 | Diseases of DNA repair | 6 of 50 | 0.69 | 0.0393 |
| HSA-68867 | Assembly of the pre-replicative complex | 13 of 111 | 0.68 | 0.00048 |
| HSA-8852276 | The role of GTSE1 in G2/M progression after G2 checkpoint | 9 of 77 | 0.68 | 0.0069 |
| HSA-5663220 | RHO GTPases Activate Formins | 16 of 139 | 0.67 | 8.25e-05 |
| HSA-68949 | Orc1 removal from chromatin | 8 of 70 | 0.67 | 0.0145 |
| HSA-6781823 | Formation of TC-NER Pre-Incision Complex | 6 of 53 | 0.67 | 0.0486 |
| HSA-69620 | Cell Cycle Checkpoints | 30 of 272 | 0.66 | 4.97e-08 |
| HSA-199977 | ER to Golgi Anterograde Transport | 17 of 154 | 0.66 | 6.71e-05 |
| HSA-176408 | Regulation of APC/C activators between G1/S and early ana... | 9 of 80 | 0.66 | 0.0085 |
| HSA-174184 | Cdc20:Phospho-APC/C mediated degradation of Cyclin A | 8 of 72 | 0.66 | 0.0168 |
| HSA-9707564 | Cytoprotection by HMOX1 | 7 of 63 | 0.66 | 0.0297 |
| HSA-174084 | Autodegradation of Cdh1 by Cdh1:APC/C | 7 of 63 | 0.66 | 0.0297 |
| HSA-68877 | Mitotic Prometaphase | 22 of 201 | 0.65 | 3.81e-06 |
| HSA-6811442 | Intra-Golgi and retrograde Golgi-to-ER traffic | 22 of 201 | 0.65 | 3.81e-06 |
| HSA-69239 | Synthesis of DNA | 13 of 120 | 0.65 | 0.00099 |
| HSA-174178 | APC/C:Cdh1 mediated degradation of Cdc20 and other APC... | 8 of 73 | 0.65 | 0.0179 |
| HSA-380320 | Recruitment of NuMA to mitotic centrosomes | 10 of 93 | 0.64 | 0.0064 |
| HSA-69615 | G1/S DNA Damage Checkpoints | 7 of 66 | 0.64 | 0.0350 |
| HSA-68882 | Mitotic Anaphase | 24 of 232 | 0.63 | 2.82e-06 |
| HSA-5658442 | Regulation of RAS by GAPs | 7 of 67 | 0.63 | 0.0367 |
| HSA-174154 | APC/C:Cdc20 mediated degradation of Securin | 7 of 67 | 0.63 | 0.0367 |
| HSA-5696398 | Nucleotide Excision Repair | 11 of 109 | 0.62 | 0.0055 |
| HSA-6781827 | Transcription-Coupled Nucleotide Excision Repair (TC-NER) | 8 of 78 | 0.62 | 0.0238 |
| HSA-380259 | Loss of Nlp from mitotic centrosomes | 7 of 69 | 0.62 | 0.0410 |
| HSA-69206 | G1/S Transition | 13 of 130 | 0.61 | 0.0019 |
| HSA-9663891 | Selective autophagy | 8 of 80 | 0.61 | 0.0269 |
| HSA-380270 | Recruitment of mitotic centrosome proteins and complexes | 8 of 80 | 0.61 | 0.0269 |
| HSA-948021 | Transport to the Golgi and subsequent modification | 18 of 185 | 0.6 | 0.00013 |
| HSA-69306 | DNA Replication | 15 of 156 | 0.6 | 0.00087 |
| HSA-195253 | Degradation of beta-catenin by the destruction complex | 8 of 82 | 0.6 | 0.0297 |
| HSA-5696399 | Global Genome Nucleotide Excision Repair (GG-NER) | 8 of 83 | 0.6 | 0.0304 |
| HSA-8854518 | AURKA Activation by TPX2 | 7 of 72 | 0.6 | 0.0486 |
| HSA-69017 | CDK-mediated phosphorylation and removal of Cdc6 | 7 of 72 | 0.6 | 0.0486 |
| HSA-72312 | rRNA processing | 19 of 201 | 0.59 | 0.00012 |
| HSA-8868773 | rRNA processing in the nucleus and cytosol | 18 of 191 | 0.59 | 0.00019 |
| HSA-69242 | S Phase | 15 of 162 | 0.58 | 0.0012 |
| HSA-8856688 | Golgi-to-ER retrograde transport | 12 of 133 | 0.57 | 0.0066 |
| HSA-1632852 | Macroautophagy | 12 of 133 | 0.57 | 0.0066 |
| HSA-6811434 | COPI-dependent Golgi-to-ER retrograde traffic | 9 of 99 | 0.57 | 0.0255 |
| HSA-6791226 | Major pathway of rRNA processing in the nucleolus and cyt... | 16 of 181 | 0.56 | 0.0011 |
| HSA-9612973 | Autophagy | 13 of 148 | 0.56 | 0.0054 |
| HSA-390466 | Chaperonin-mediated protein folding | 8 of 91 | 0.56 | 0.0453 |
| HSA-69481 | G2/M Checkpoints | 13 of 149 | 0.55 | 0.0055 |
| HSA-9755511 | KEAP1-NFE2L2 pathway | 9 of 105 | 0.55 | 0.0332 |
| HSA-68886 | M Phase | 32 of 382 | 0.54 | 1.61e-06 |
| HSA-72737 | Cap-dependent Translation Initiation | 10 of 118 | 0.54 | 0.0236 |
| HSA-72766 | Translation | 24 of 290 | 0.53 | 5.95e-05 |
| HSA-195258 | RHO GTPase Effectors | 24 of 292 | 0.53 | 6.12e-05 |
| HSA-69278 | Cell Cycle, Mitotic | 42 of 526 | 0.52 | 5.01e-08 |
| HSA-5617833 | Cilium Assembly | 16 of 200 | 0.52 | 0.0028 |
| HSA-1640170 | Cell Cycle | 51 of 658 | 0.5 | 4.56e-09 |
| HSA-73894 | DNA Repair | 24 of 310 | 0.5 | 0.00013 |

| | | | | |
|---|---|---|---|---|
| HSA-9711123 | Cellular response to chemical stress | 15 of 195 | 0.5 | 0.0062 |
| HSA-69275 | G2/M Transition | 15 of 195 | 0.5 | 0.0062 |
| HSA-1852241 | Organelle biogenesis and maintenance | 22 of 293 | 0.49 | 0.00044 |
| HSA-201681 | TCF dependent signaling in response to WNT | 15 of 201 | 0.49 | 0.0072 |
| HSA-5357801 | Programmed Cell Death | 15 of 206 | 0.48 | 0.0086 |
| HSA-2559583 | Cellular Senescence | 12 of 163 | 0.48 | 0.0238 |
| HSA-8953854 | Metabolism of RNA | 50 of 705 | 0.46 | 5.01e-08 |
| HSA-199991 | Membrane Trafficking | 43 of 626 | 0.45 | 1.29e-06 |
| HSA-983168 | Antigen processing: Ubiquitination & Proteasome degradati... | 21 of 303 | 0.45 | 0.0018 |
| HSA-109581 | Apoptosis | 12 of 175 | 0.45 | 0.0352 |
| HSA-194315 | Signaling by Rho GTPases | 45 of 672 | 0.44 | 1.29e-06 |
| HSA-3700989 | Transcriptional Regulation by TP53 | 23 of 361 | 0.42 | 0.0025 |
| HSA-5684996 | MAPK1/MAPK3 signaling | 18 of 283 | 0.42 | 0.0103 |
| HSA-446203 | Asparagine N-linked glycosylation | 19 of 304 | 0.41 | 0.0088 |
| HSA-983169 | Class I MHC mediated antigen processing & presentation | 23 of 376 | 0.4 | 0.0040 |
| HSA-5673001 | RAF/MAP kinase cascade | 17 of 277 | 0.4 | 0.0181 |
| HSA-72203 | Processing of Capped Intron-Containing Pre-mRNA | 17 of 281 | 0.39 | 0.0206 |
| HSA-2262752 | Cellular responses to stress | 44 of 747 | 0.38 | 2.15e-05 |
| HSA-9012999 | RHO GTPase cycle | 26 of 449 | 0.38 | 0.0034 |
| HSA-6798695 | Neutrophil degranulation | 26 of 476 | 0.35 | 0.0066 |
| HSA-5663205 | Infectious disease | 46 of 917 | 0.31 | 0.00037 |
| HSA-5663202 | Diseases of signal transduction by growth factor receptors ... | 21 of 430 | 0.3 | 0.0491 |
| HSA-1280218 | Adaptive Immune System | 34 of 758 | 0.27 | 0.0179 |
| HSA-392499 | Metabolism of proteins | 85 of 1917 | 0.26 | 1.14e-05 |
| HSA-74160 | Gene expression (Transcription) | 65 of 1476 | 0.26 | 0.00029 |
| HSA-1643685 | Disease | 74 of 1702 | 0.25 | 0.00012 |
| HSA-73857 | RNA Polymerase II Transcription | 54 of 1337 | 0.22 | 0.0084 |
| HSA-168249 | Innate Immune System | 41 of 1041 | 0.21 | 0.0416 |
| HSA-597592 | Post-translational protein modification | 53 of 1405 | 0.19 | 0.0297 |
| HSA-1430728 | Metabolism | 77 of 2092 | 0.18 | 0.0072 |
| HSA-168256 | Immune System | 70 of 1979 | 0.16 | 0.0273 |
| HSA-162582 | Signal Transduction | 85 of 2540 | 0.14 | 0.0352 |

**Subcellular localization (COMPARTMENTS)**

| *compartment* | *description* | *count in network* | *strength* | *false discovery rate* |
|---|---|---|---|---|
| GOCC:0005827 | Polar microtubule | 3 of 5 | 1.39 | 0.0277 |
| GOCC:0031499 | TRAMP complex | 3 of 6 | 1.31 | 0.0394 |
| GOCC:0009317 | acetyl-CoA carboxylase complex | 3 of 6 | 1.31 | 0.0394 |
| GOCC:0005835 | Fatty acid synthase complex | 3 of 6 | 1.31 | 0.0394 |
| GOCC:1990710 | MutS complex | 3 of 7 | 1.25 | 0.0493 |
| GOCC:0032389 | MutLalpha complex | 3 of 7 | 1.25 | 0.0493 |
| GOCC:0032302 | MutSbeta complex | 3 of 7 | 1.25 | 0.0493 |
| GOCC:0032301 | MutSalpha complex | 3 of 7 | 1.25 | 0.0493 |
| GOCC:0017119 | Golgi transport complex | 4 of 11 | 1.17 | 0.0155 |
| GOCC:0000177 | Cytoplasmic exosome (RNase complex) | 4 of 13 | 1.1 | 0.0251 |
| GOCC:0005852 | Eukaryotic translation initiation factor 3 complex | 5 of 19 | 1.03 | 0.0102 |
| GOCC:0031261 | DNA replication preinitiation complex | 4 of 17 | 0.98 | 0.0493 |
| GOCC:0031010 | ISWI-type complex | 4 of 17 | 0.98 | 0.0493 |
| GOCC:0000159 | Protein phosphatase type 2A complex | 4 of 17 | 0.98 | 0.0493 |
| GOCC:0005963 | Magnesium-dependent protein serine/threonine phosphatas... | 6 of 27 | 0.96 | 0.0058 |
| GOCC:0042645 | Mitochondrial nucleoid | 12 of 55 | 0.95 | 5.48e-06 |
| GOCC:0099023 | Vesicle tethering complex | 10 of 60 | 0.84 | 0.00045 |
| GOCC:1990391 | DNA repair complex | 9 of 53 | 0.84 | 0.00099 |
| GOCC:0032040 | Small-subunit processome | 5 of 31 | 0.82 | 0.0496 |
| GOCC:1904949 | ATPase complex | 16 of 120 | 0.74 | 1.50e-05 |
| GOCC:0008287 | Protein serine/threonine phosphatase complex | 7 of 54 | 0.73 | 0.0251 |
| GOCC:0000152 | Nuclear ubiquitin ligase complex | 7 of 60 | 0.68 | 0.0394 |
| GOCC:0030684 | Preribosome | 8 of 72 | 0.66 | 0.0257 |
| GOCC:0070603 | SWI/SNF superfamily-type complex | 10 of 95 | 0.64 | 0.0097 |

| | | | | |
|---|---|---|---|---|
| GOCC:1904813 | ficolin-1-rich granule lumen | 12 of 124 | 0.6 | 0.0054 |
| GOCC:0030496 | Midbody | 13 of 142 | 0.57 | 0.0050 |
| GOCC:0000228 | Nuclear chromosome | 28 of 320 | 0.56 | 2.40e-06 |
| GOCC:0005730 | Nucleolus | 50 of 606 | 0.53 | 4.95e-11 |
| GOCC:0000793 | Condensed chromosome | 16 of 205 | 0.51 | 0.0048 |
| GOCC:0005819 | Spindle | 22 of 289 | 0.49 | 0.00045 |
| GOCC:0098687 | Chromosomal region | 16 of 227 | 0.46 | 0.0115 |
| GOCC:0005874 | Microtubule | 16 of 233 | 0.45 | 0.0148 |
| GOCC:0005802 | trans-Golgi network | 13 of 192 | 0.44 | 0.0477 |
| GOCC:0031981 | Nuclear lumen | 120 of 1850 | 0.43 | 1.66e-20 |
| GOCC:0098791 | Golgi apparatus subcompartment | 16 of 244 | 0.43 | 0.0229 |
| GOCC:0005813 | Centrosome | 30 of 470 | 0.42 | 0.00030 |
| GOCC:1902494 | Catalytic complex | 107 of 1710 | 0.41 | 7.05e-17 |
| GOCC:0005694 | Chromosome | 59 of 951 | 0.41 | 1.94e-08 |
| GOCC:0000785 | Chromatin | 29 of 476 | 0.4 | 0.00086 |
| GOCC:0005759 | Mitochondrial matrix | 23 of 379 | 0.4 | 0.0055 |
| GOCC:0005654 | Nucleoplasm | 69 of 1146 | 0.39 | 1.68e-09 |
| GOCC:0031984 | Organelle subcompartment | 42 of 739 | 0.37 | 6.55e-05 |
| GOCC:0042175 | Nuclear outer membrane-endoplasmic reticulum membrane... | 32 of 560 | 0.37 | 0.00099 |
| GOCC:0005789 | Endoplasmic reticulum membrane | 30 of 538 | 0.36 | 0.0026 |
| GOCC:0070013 | Intracellular organelle lumen | 159 of 2902 | 0.35 | 5.33e-21 |
| GOCC:1990234 | Transferase complex | 49 of 892 | 0.35 | 2.04e-05 |
| GOCC:0005829 | Cytosol | 163 of 3054 | 0.34 | 1.24e-20 |
| GOCC:0015630 | Microtubule cytoskeleton | 52 of 973 | 0.34 | 1.99e-05 |
| GOCC:0005815 | Microtubule organizing center | 32 of 607 | 0.34 | 0.0039 |
| GOCC:1990904 | Ribonucleoprotein complex | 30 of 564 | 0.34 | 0.0053 |
| GOCC:0098827 | Endoplasmic reticulum subcompartment | 27 of 502 | 0.34 | 0.0089 |
| GOCC:0043232 | Intracellular non-membrane-bounded organelle | 171 of 3309 | 0.33 | 1.66e-20 |
| GOCC:0005634 | Nucleus | 234 of 4787 | 0.3 | 9.68e-28 |
| GOCC:0032991 | Protein-containing complex | 250 of 5325 | 0.28 | 8.33e-28 |
| GOCC:0005739 | Mitochondrion | 54 of 1182 | 0.27 | 0.00082 |
| GOCC:0043231 | Intracellular membrane-bounded organelle | 343 of 8162 | 0.24 | 1.67e-36 |
| GOCC:0031090 | Organelle membrane | 96 of 2290 | 0.24 | 1.35e-05 |
| GOCC:0005856 | Cytoskeleton | 65 of 1575 | 0.23 | 0.0020 |
| GOCC:0043229 | Intracellular organelle | 376 of 9609 | 0.21 | 1.67e-36 |
| GOCC:0043227 | Membrane-bounded organelle | 360 of 9083 | 0.21 | 1.73e-34 |
| GOCC:0005622 | Intracellular | 431 of 11512 | 0.19 | 5.09e-48 |
| GOCC:0043226 | Organelle | 386 of 10113 | 0.19 | 2.07e-36 |
| GOCC:0005737 | Cytoplasm | 310 of 8195 | 0.19 | 2.84e-21 |
| GOCC:0098588 | Bounding membrane of organelle | 55 of 1451 | 0.19 | 0.0392 |
| GOCC:0012505 | Endomembrane system | 114 of 3156 | 0.17 | 0.00072 |
| GOCC:0110165 | Cellular anatomical entity | 436 of 14060 | 0.1 | 9.03e-23 |

### Annotated Keywords (UniProt)

| keyword | description | count in network | strength | false discovery rate |
|---|---|---|---|---|
| KW-1135 | Mitochondrion nucleoid | 6 of 17 | 1.16 | 0.00034 |
| KW-0030 | Aminoacyl-tRNA synthetase | 10 of 38 | 1.03 | 6.45e-06 |
| KW-0347 | Helicase | 26 of 141 | 0.88 | 2.65e-12 |
| KW-0396 | Initiation factor | 8 of 47 | 0.84 | 0.0010 |
| KW-0648 | Protein biosynthesis | 21 of 128 | 0.83 | 4.05e-09 |
| KW-1026 | Leukodystrophy | 5 of 35 | 0.77 | 0.0395 |
| KW-0436 | Ligase | 17 of 131 | 0.73 | 4.08e-06 |
| KW-0698 | rRNA processing | 11 of 95 | 0.68 | 0.0010 |
| KW-0647 | Proteasome | 6 of 52 | 0.68 | 0.0395 |
| KW-0090 | Biological rhythms | 15 of 145 | 0.63 | 0.00021 |
| KW-0931 | ER-Golgi transport | 9 of 91 | 0.61 | 0.0115 |
| KW-0511 | Multifunctional enzyme | 7 of 74 | 0.59 | 0.0471 |
| KW-0235 | DNA replication | 9 of 98 | 0.58 | 0.0169 |
| KW-0067 | ATP-binding | 110 of 1379 | 0.52 | 2.89e-25 |
| KW-0507 | mRNA processing | 27 of 349 | 0.5 | 1.46e-05 |
| KW-0547 | Nucleotide-binding | 133 of 1776 | 0.49 | 8.88e-29 |
| KW-0156 | Chromatin regulator | 22 of 292 | 0.49 | 0.00021 |
| KW-0508 | mRNA splicing | 20 of 274 | 0.48 | 0.00069 |
| KW-0137 | Centromere | 10 of 138 | 0.47 | 0.0443 |
| KW-0653 | Protein transport | 43 of 617 | 0.46 | 1.42e-07 |
| KW-0132 | Cell division | 27 of 384 | 0.46 | 7.34e-05 |
| KW-0498 | Mitosis | 19 of 275 | 0.45 | 0.0018 |
| KW-0131 | Cell cycle | 44 of 651 | 0.44 | 2.03e-07 |
| KW-0227 | DNA damage | 26 of 386 | 0.44 | 0.00021 |
| KW-0007 | Acetylation | 221 of 3362 | 0.43 | 5.91e-45 |
| KW-0694 | RNA-binding | 45 of 686 | 0.43 | 2.94e-07 |
| KW-0523 | Neurodegeneration | 23 of 349 | 0.43 | 0.00077 |
| KW-0945 | Host-virus interaction | 35 of 540 | 0.42 | 1.47e-05 |
| KW-0234 | DNA repair | 21 of 325 | 0.42 | 0.0018 |
| KW-0158 | Chromosome | 26 of 433 | 0.39 | 0.0010 |
| KW-0723 | Serine/threonine-protein kinase | 23 of 390 | 0.38 | 0.0030 |
| KW-0342 | GTP-binding | 20 of 342 | 0.38 | 0.0085 |
| KW-0809 | Transit peptide | 30 of 545 | 0.35 | 0.0011 |
| KW-1017 | Isopeptide bond | 89 of 1717 | 0.33 | 1.97e-09 |
| KW-0418 | Kinase | 30 of 634 | 0.29 | 0.0115 |
| KW-0832 | Ubl conjugation | 111 of 2399 | 0.28 | 2.99e-09 |
| KW-0833 | Ubl conjugation pathway | 32 of 695 | 0.28 | 0.0115 |
| KW-0597 | Phosphoprotein | 369 of 8122 | 0.27 | 9.83e-53 |
| KW-0496 | Mitochondrion | 53 of 1181 | 0.27 | 0.00059 |
| KW-0808 | Transferase | 80 of 1820 | 0.26 | 1.22e-05 |
| KW-0378 | Hydrolase | 70 of 1612 | 0.25 | 9.27e-05 |
| KW-0539 | Nucleus | 218 of 5278 | 0.23 | 6.65e-16 |
| KW-0963 | Cytoplasm | 212 of 5095 | 0.23 | 1.08e-15 |
| KW-0206 | Cytoskeleton | 49 of 1235 | 0.21 | 0.0125 |
| KW-0025 | Alternative splicing | 318 of 10313 | 0.1 | 4.62e-08 |

**Supplemental Table 5. Proteins which were upregulated by GEM treatment in A375 melanoma cells.**

| Upregulated proteins after 24 hours of treatment with 20 nM GEM | | | |
|---|---|---|---|
| **Gene name for protein** | **Description** | **Fold change (log2 form)** | **t-test P value** |
| **PCYT1A** | Choline-phosphate cytidylyltransferase A | 1.061138 | 3.74E-05 |
| **TK1** | Thymidine kinase, cytosolic | 1.574312 | 4.18E-05 |
| **CDC37L1** | Hsp90 co-chaperone Cdc37-like 1 | 0.545999 | 7.18E-05 |
| **SHMT2** | Serine hydroxymethyltransferase, mitochondrial | 0.276772 | 7.65E-05 |
| **SFN** | 14-3-3 protein sigma | 1.828153 | 7.95E-05 |
| **FKBP9** | Peptidyl-prolyl cis-trans isomerase FKBP9 | 0.733202 | 7.97E-05 |
| **KRT8** | Keratin, type II cytoskeletal 8 | 0.918275 | 9.45E-05 |
| **SRRT** | Serrate RNA effector molecule homolog | 0.532921 | 0.000101 |
| **DNPH1** | 2'-deoxynucleoside 5'-phosphate N-hydrolase 1 | 0.975685 | 0.000109 |
| **RBM10** | RNA-binding protein 10 | 0.486721 | 0.000122 |
| **IKBIP** | Inhibitor of nuclear factor kappa-B kinase-interacting protein | 0.503438 | 0.000125 |
| **GLG1** | Golgi apparatus protein 1 | 0.48834 | 0.000132 |
| **VTI1B** | Vesicle transport through interaction with t-SNAREs homolog 1B | 0.412797 | 0.000138 |
| **SMTN** | Smoothelin | 1.699071 | 0.000143 |
| **YTHDC1** | YTH domain-containing protein 1 | 0.469841 | 0.000144 |
| **TERF2** | Telomeric repeat-binding factor 2 | 0.429996 | 0.000156 |
| **CTHRC1** | Collagen triple helix repeat-containing protein 1 | 1.23766 | 0.000158 |
| **ATXN3** | Ataxin-3 | 0.39188 | 0.000166 |
| **SYF2** | Pre-mRNA-splicing factor SYF2 | 0.643827 | 0.000176 |
| **PCM1** | Pericentriolar material 1 protein | 0.446632 | 0.000179 |
| **ECM1** | Extracellular matrix protein 1 | 1.203733 | 0.000207 |
| **MICALL1** | MICAL-like protein 1 | 0.757376 | 0.000218 |
| **PNMA8A** | Paraneoplastic antigen-like protein 8A | 0.448057 | 0.000223 |
| **NOL3** | Nucleolar protein 3 | 1.605692 | 0.000238 |

| | | | |
|---|---|---|---|
| **PML** | Protein PML | 0.52529 | 0.000262 |
| **CHMP3** | Charged multivesicular body protein 3 | 1.047321 | 0.000271 |
| **HLA-DMA** | HLA class II histocompatibility antigen, DM alpha chain | 1.441759 | 0.000291 |
| **CDCA5** | Sororin | 1.516139 | 0.000294 |
| **PHLDB1** | Pleckstrin homology-like domain family B member 1 | 0.441239 | 0.000315 |
| **YWHAQ** | 14-3-3 protein theta | 0.331771 | 0.000334 |
| **REPS1** | RalBP1-associated Eps domain-containing protein 1 | 0.354177 | 0.000351 |
| **RAI14** | Ankycorbin | 0.270497 | 0.000358 |
| **UBE2A** | Ubiquitin-conjugating enzyme E2 A | 1.375949 | 0.000364 |
| **RAD18** | E3 ubiquitin-protein ligase RAD18 | 0.352582 | 0.000376 |
| **HLA-B** | HLA class I histocompatibility antigen, B alpha chain | 0.699311 | 0.00038 |
| **REXO2** | Oligoribonuclease, mitochondrial | 0.304103 | 0.000387 |
| **ACP1** | Low molecular weight phosphotyrosine protein phosphatase | 0.396862 | 0.000392 |
| **LARP7** | La-related protein 7 | 0.386477 | 0.000393 |
| **VPS28** | Vacuolar protein sorting-associated protein 28 homolog | 0.928887 | 0.000404 |
| **GRWD1** | Glutamate-rich WD repeat-containing protein 1 | 0.708947 | 0.000414 |
| **CTSB** | Cathepsin B | 1.045409 | 0.000422 |
| **NDUFA8** | NADH dehydrogenase [ubiquinone] 1 alpha subcomplex subunit 8 | 0.894316 | 0.000438 |
| **MRPL42** | 39S ribosomal protein L42, mitochondrial | 1.008335 | 0.000443 |
| **SFSWAP** | Splicing factor, suppressor of white-apricot homolog | 0.296876 | 0.000475 |
| **NOL8** | Nucleolar protein 8 | 0.513434 | 0.00048 |
| **CKAP4** | Cytoskeleton-associated protein 4 | 0.529373 | 0.0005 |
| **VSNL1** | Visinin-like protein 1 | 0.878997 | 0.000503 |
| **TNRC6B** | Trinucleotide repeat-containing gene 6B protein | 0.681316 | 0.000517 |

| | | | |
|---|---|---|---|
| **GOLGA4** | Golgin subfamily A member 4 | 0.28952 | 0.000525 |
| **FOXO3** | Forkhead box protein O3 | 0.554012 | 0.000527 |
| **ILF3** | Interleukin enhancer-binding factor 3 | 0.408037 | 0.000529 |
| **CTSZ** | Cathepsin Z | 1.154656 | 0.000534 |
| **UBE2H** | Ubiquitin-conjugating enzyme E2 H | 0.592209 | 0.000568 |
| **SASS6** | Spindle assembly abnormal protein 6 homolog | 0.942137 | 0.000571 |
| **XRCC1** | DNA repair protein XRCC1 | 0.523168 | 0.000575 |
| **MAP1B** | Microtubule-associated protein 1B | 0.361875 | 0.00058 |
| **LZTS1** | Leucine zipper putative tumor suppressor 1 | 0.462023 | 0.000581 |
| **DNASE2** | Deoxyribonuclease-2-alpha | 0.850023 | 0.0006 |
| **CKAP2** | Cytoskeleton-associated protein 2 | 1.346669 | 0.000608 |
| **PIBF1** | Progesterone-induced-blocking factor 1 | 0.199159 | 0.000675 |
| **TAGLN3** | Transgelin-3 | 0.917039 | 0.00069 |
| **RNF20** | E3 ubiquitin-protein ligase BRE1A | 0.414906 | 0.000701 |
| **CHAF1A** | Chromatin assembly factor 1 subunit A | 0.649914 | 0.000718 |
| **CRELD2** | Protein disulfide isomerase CRELD2 | 1.112964 | 0.000766 |
| **NKAP** | NF-kappa-B-activating protein | 0.533423 | 0.000801 |
| **NUP153** | Nuclear pore complex protein Nup153 | 0.456075 | 0.000825 |
| **ATG3** | Ubiquitin-like-conjugating enzyme ATG3 | 0.408324 | 0.000827 |
| **TACC3** | Transforming acidic coiled-coil-containing protein 3 | 1.634785 | 0.000836 |
| **TMPO** | Lamina-associated polypeptide 2, isoform alpha | 0.276678 | 0.000845 |
| **FOXJ2** | Forkhead box protein J2 | 0.43742 | 0.000848 |
| **CROCC** | Rootletin | 0.211957 | 0.000863 |
| **PLD3** | 5'-3' exonuclease PLD3 | 0.723275 | 0.000866 |
| **NUP62** | Nuclear pore glycoprotein p62 | 0.850618 | 0.000867 |
| **RCN1** | Reticulocalbin-1 | 0.809067 | 0.000887 |

| | | | |
|---|---|---|---|
| **SRSF8** | Serine/arginine-rich splicing factor 8 | 0.717749 | 0.000889 |
| **FAIM** | Fas apoptotic inhibitory molecule 1 | 0.707832 | 0.000902 |
| **MBD1** | Methyl-CpG-binding domain protein 1 | 0.472827 | 0.000905 |
| **CCP110** | Centriolar coiled-coil protein of 110 kDa | 0.887105 | 0.000905 |
| **LIMCH1** | LIM and calponin homology domains-containing protein 1 | 0.758782 | 0.000905 |
| **HSPB1** | Heat shock protein beta-1 | 0.948509 | 0.000949 |
| **TIMM8A** | Mitochondrial import inner membrane translocase subunit Tim8 A | 1.736463 | 0.000959 |
| **LRP1** | Prolow-density lipoprotein receptor-related protein 1 | 0.435394 | 0.000971 |
| **INCENP** | Inner centromere protein | 0.986911 | 0.000977 |
| **NEXN** | Nexilin | 0.78732 | 0.000987 |
| **MPRIP** | Myosin phosphatase Rho-interacting protein | 0.309214 | 0.000998 |
| **ACBD5** | Acyl-CoA-binding domain-containing protein 5 | 0.580858 | 0.001 |
| **NUMB** | Protein numb homolog | 0.320578 | 0.001019 |
| **DNAJB2** | DnaJ homolog subfamily B member 2 | 0.601318 | 0.001026 |
| **APC** | Adenomatous polyposis coli protein | 0.268629 | 0.00104 |
| **JUN** | Transcription factor Jun | 1.67651 | 0.001047 |
| **COX6B1** | Cytochrome c oxidase subunit 6B1 | 1.556949 | 0.001066 |
| **SLC9A3R1** | Na(+)/H(+) exchange regulatory cofactor NHE-RF1 | 0.762379 | 0.001082 |
| **BLM** | RecQ-like DNA helicase BLM | 0.740791 | 0.001108 |
| **CGN** | Cingulin | 0.394074 | 0.001115 |
| **PDCD4** | Programmed cell death protein 4 | 0.74574 | 0.001116 |
| **HLA-C** | HLA class I histocompatibility antigen, C alpha chain | 0.671223 | 0.001129 |
| **TACC1** | Transforming acidic coiled-coil-containing protein 1 | 0.628188 | 0.001153 |
| **BCAR1** | Breast cancer anti-estrogen resistance protein 1 | 0.382141 | 0.001156 |

| | | | |
|---|---|---|---|
| **MPHOSPH6** | M-phase phosphoprotein 6 | 0.928814 | 0.001171 |
| **MYBL2** | Myb-related protein B | 0.81933 | 0.001183 |
| **ITGA3** | Integrin alpha-3 | 0.489185 | 0.001186 |
| **SORBS2** | Sorbin and SH3 domain-containing protein 2 | 0.636972 | 0.001199 |
| **RAN** | GTP-binding nuclear protein Ran | 0.731141 | 0.00124 |
| **RPRD1B** | Regulation of nuclear pre-mRNA domain-containing protein 1B | 0.511633 | 0.001276 |
| **URM1** | Ubiquitin-related modifier 1 | 0.864234 | 0.001341 |
| **SSB** | Lupus La protein | 0.568653 | 0.001352 |
| **ERO1A** | ERO1-like protein alpha | 0.462858 | 0.001372 |
| **MAGED2** | Melanoma-associated antigen D2 | 0.63476 | 0.001396 |
| **TCERG1** | Transcription elongation regulator 1 | 0.429614 | 0.001408 |
| **OGFR** | Opioid growth factor receptor | 0.363672 | 0.001417 |
| **CHAMP1** | Chromosome alignment-maintaining phosphoprotein 1 | 0.506935 | 0.001435 |
| **PRC1** | Protein regulator of cytokinesis 1 | 0.902278 | 0.001469 |
| **PCBD1** | Pterin-4-alpha-carbinolamine dehydratase | 1.107696 | 0.001475 |
| **COTL1** | Coactosin-like protein | 1.277928 | 0.001475 |
| **ATP5PD** | ATP synthase subunit d, mitochondrial | 1.001327 | 0.001519 |
| **SRI** | Sorcin | 0.545876 | 0.001531 |
| **MRPS18B** | 28S ribosomal protein S18b, mitochondrial | 0.366602 | 0.001566 |
| **AKAP2** | A-kinase anchor protein 2 | 0.921541 | 0.001628 |
| **HMGN1** | Non-histone chromosomal protein HMG-14 | 0.316364 | 0.00163 |
| **TOR1AIP2** | Torsin-1A-interacting protein 2 | 0.649261 | 0.001631 |
| **REXO4** | RNA exonuclease 4 | 0.711946 | 0.001631 |
| **GRIPAP1** | GRIP1-associated protein 1 | 0.283344 | 0.001632 |
| **RBM25** | RNA-binding protein 25 | 0.432052 | 0.00166 |
| **TNFRSF10B** | Tumor necrosis factor receptor superfamily member 10B | 0.789069 | 0.001691 |
| **TOP2A** | DNA topoisomerase 2-alpha | 0.3289 | 0.0017 |

| | | | |
|---|---|---|---|
| **WASL** | Actin nucleation-promoting factor WASL | 0.480725 | 0.001716 |
| **MOCS2** | Molybdopterin synthase catalytic subunit | 1.021089 | 0.001743 |
| **GOPC** | Golgi-associated PDZ and coiled-coil motif-containing protein | 0.562253 | 0.001749 |
| **SKP1** | S-phase kinase-associated protein 1 | 1.229828 | 0.001765 |
| **RCC1** | Regulator of chromosome condensation | 0.438823 | 0.001771 |
| **LMNB1** | Lamin-B1 | 0.303848 | 0.001815 |
| **ICAM1** | Intercellular adhesion molecule 1 | 1.009765 | 0.001827 |
| **NCOR2** | Nuclear receptor corepressor 2 | 0.308793 | 0.001846 |
| **BRD8** | Bromodomain-containing protein 8 | 0.570148 | 0.001877 |
| **HLA-DRA** | HLA class II histocompatibility antigen, DR alpha chain | 0.58783 | 0.001932 |
| **CHMP6** | Charged multivesicular body protein 6 | 0.813034 | 0.001941 |
| **PRKCSH** | Glucosidase 2 subunit beta | 0.850675 | 0.00195 |
| **LIMA1** | LIM domain and actin-binding protein 1 | 0.812512 | 0.001973 |
| **NCKAP5L** | Nck-associated protein 5-like | 0.722249 | 0.001989 |
| **LARP1** | La-related protein 1 | 0.332439 | 0.002013 |
| **MAP1A** | Microtubule-associated protein 1A | 0.546537 | 0.002016 |
| **CSRP2** | Cysteine and glycine-rich protein 2 | 0.806611 | 0.002039 |
| **HABP4** | Intracellular hyaluronan-binding protein 4 | 0.379679 | 0.002052 |
| **ECE1** | Endothelin-converting enzyme 1 | 0.996566 | 0.002064 |
| **VPS25** | Vacuolar protein-sorting-associated protein 25 | 0.610776 | 0.002066 |
| **BAX** | Apoptosis regulator BAX | 0.488836 | 0.002076 |
| **MRPS26** | 28S ribosomal protein S26, mitochondrial | 0.438625 | 0.002096 |
| **SAP130** | Histone deacetylase complex subunit SAP130 | 0.341112 | 0.002115 |
| **PEA15** | Astrocytic phosphoprotein PEA-15 | 0.960507 | 0.002205 |
| **VPS37B** | Vacuolar protein sorting-associated protein 37B | 0.373375 | 0.002234 |

| | | | |
|---|---|---|---|
| **FOSB** | Protein FosB | 1.181642 | 0.002234 |
| **TMED8** | Protein TMED8 | 0.250119 | 0.002239 |
| **FTL** | Ferritin light chain | 0.517937 | 0.002267 |
| **RGS3** | Regulator of G-protein signaling 3 | 0.373349 | 0.002287 |
| **BCL7C** | B-cell CLL/lymphoma 7 protein family member C | 0.538613 | 0.002316 |
| **PSME2** | Proteasome activator complex subunit 2 | 1.012727 | 0.002333 |
| **DDB2** | DNA damage-binding protein 2 | 0.680314 | 0.002343 |
| **C1orf109** | Ribosome biogenesis protein C1orf109 | 0.46547 | 0.00235 |
| **NDUFB9** | NADH dehydrogenase [ubiquinone] 1 beta subcomplex subunit 9 | 1.214057 | 0.002353 |
| **CD70** | CD70 antigen | 0.98763 | 0.00238 |
| **FEN1** | Flap endonuclease 1 | 0.711855 | 0.002392 |
| **BCL9** | B-cell CLL/lymphoma 9 protein | 0.32439 | 0.002438 |
| **PRPF3** | U4/U6 small nuclear ribonucleoprotein Prp3 | 0.406686 | 0.002448 |
| **CAVIN3** | Caveolae-associated protein 3 | 1.002002 | 0.00248 |
| **RTF1** | RNA polymerase-associated protein RTF1 homolog | 0.36412 | 0.002512 |
| **EIF4E** | Eukaryotic translation initiation factor 4E | 0.292865 | 0.002513 |
| **URI1** | Unconventional prefoldin RPB5 interactor 1 | 0.349074 | 0.002551 |
| **MRPS28** | 28S ribosomal protein S28, mitochondrial | 0.677315 | 0.002554 |
| **PACSIN2** | Protein kinase C and casein kinase substrate in neurons protein 2 | 0.608188 | 0.002575 |
| **DNAJC2** | DnaJ homolog subfamily C member 2 | 0.374855 | 0.002598 |
| **KIF11** | Kinesin-like protein KIF11 | 0.383667 | 0.002627 |
| **DFFA** | DNA fragmentation factor subunit alpha | 0.865652 | 0.002645 |
| **IFIT3** | Interferon-induced protein with tetratricopeptide repeats 3 | 0.449921 | 0.002653 |
| **ANLN** | Anillin | 1.050883 | 0.002656 |
| **FAM169A** | Soluble lamin-associated protein of 75 kDa | 0.366088 | 0.002669 |

| | | | |
|---|---|---|---|
| **DBN1** | Drebrin | 0.632383 | 0.002676 |
| **HDDC2** | 5'-deoxynucleotidase HDDC2 | 0.446612 | 0.002738 |
| **FAS** | Tumor necrosis factor receptor superfamily member 6 | 1.215144 | 0.00275 |
| **FNBP1L** | Formin-binding protein 1-like | 0.622638 | 0.002824 |
| **CLSPN** | Claspin | 1.134882 | 0.00284 |
| **MEA1** | Male-enhanced antigen 1 | 0.759647 | 0.002894 |
| **PDLIM5** | PDZ and LIM domain protein 5 | 0.563056 | 0.002911 |
| **KANK2** | KN motif and ankyrin repeat domain-containing protein 2 | 0.307249 | 0.002948 |
| **CEMIP2** | Cell surface hyaluronidase | 0.436458 | 0.00295 |
| **GMNN** | Geminin | 1.294287 | 0.002959 |
| **MAFF** | Transcription factor MafF | 1.070628 | 0.002968 |
| **SCARB1** | Scavenger receptor class B member 1 | 0.491139 | 0.003013 |
| **GADD45GIP1** | Growth arrest and DNA damage-inducible proteins-interacting protein 1 | 0.811822 | 0.003026 |
| **SPECC1L** | Cytospin-A | 0.390787 | 0.003029 |
| **NAPA** | Alpha-soluble NSF attachment protein | 0.466118 | 0.003037 |
| **RNASEH1** | Ribonuclease H1 | 0.568631 | 0.003052 |
| **C1orf50** | Uncharacterized protein C1orf50 | 0.364485 | 0.003062 |
| **CLTB** | Clathrin light chain B | 0.741407 | 0.003083 |
| **CDC5L** | Cell division cycle 5-like protein | 0.399504 | 0.003097 |
| **PPP1R8** | Nuclear inhibitor of protein phosphatase 1 | 0.489789 | 0.003097 |
| **SLAIN1** | SLAIN motif-containing protein 1 | 1.022019 | 0.003162 |
| **TIE1** | Tyrosine-protein kinase receptor Tie-1 | 0.738805 | 0.003252 |
| **PCBD2** | Pterin-4-alpha-carbinolamine dehydratase 2 | 0.743147 | 0.003277 |
| **FHL3** | Four and a half LIM domains protein 3 | 1.354208 | 0.00329 |
| **TRIOBP** | TRIO and F-actin-binding protein | 0.584813 | 0.003322 |

| | | | |
|---|---|---|---|
| **CNOT2** | CCR4-NOT transcription complex subunit 2 | 0.205439 | 0.003374 |
| **PEBP1** | Phosphatidylethanolamine-binding protein 1 | 1.173596 | 0.003395 |
| **ZFYVE19** | Abscission/NoCut checkpoint regulator | 0.849389 | 0.00342 |
| **KAZN** | Kazrin | 0.867422 | 0.003454 |
| **TUFT1** | Tuftelin | 0.589861 | 0.003476 |
| **FAM98B** | Protein FAM98B | 0.238381 | 0.003487 |
| **MACROH2A1** | Core histone macro-H2A.1 | 0.433728 | 0.003493 |
| **TOM1** | Target of Myb1 membrane trafficking protein | 0.380499 | 0.003505 |
| **RPA2** | Replication protein A 32 kDa subunit | 0.942145 | 0.003579 |
| **AKAP12** | A-kinase anchor protein 12 | 0.514061 | 0.003582 |
| **PALM2** | Paralemmin-2 | 1.404905 | 0.003598 |
| **RRM2** | Ribonucleoside-diphosphate reductase subunit M2 | 0.86322 | 0.003625 |
| **MSI2** | RNA-binding protein Musashi homolog 2 | 0.433319 | 0.003633 |
| **CCDC115** | Coiled-coil domain-containing protein 115 | 0.420282 | 0.00364 |
| **BLOC1S6** | Biogenesis of lysosome-related organelles complex 1 subunit 6 | 0.730881 | 0.003641 |
| **FDXR** | NADPH:adrenodoxin oxidoreductase, mitochondrial | 0.351431 | 0.003668 |
| **NUDT5** | ADP-sugar pyrophosphatase | 0.400822 | 0.00376 |
| **TMEM30A** | Cell cycle control protein 50A | 0.189425 | 0.003761 |
| **IFI35** | Interferon-induced 35 kDa protein | 0.358646 | 0.003782 |
| **LMO7** | LIM domain only protein 7 | 0.41965 | 0.003784 |
| **FAM117B** | Protein FAM117B | 0.174117 | 0.003798 |
| **PGAM1** | Phosphoglycerate mutase 1 | 0.56646 | 0.003821 |
| **LMCD1** | LIM and cysteine-rich domains protein 1 | 0.660339 | 0.003833 |
| **VIM** | Vimentin | 0.53226 | 0.003873 |
| **RPE** | Ribulose-phosphate 3-epimerase | 0.641112 | 0.003954 |
| **ZNF346** | Zinc finger protein 346 | 0.698994 | 0.003955 |
| **MRPL41** | 39S ribosomal protein L41, mitochondrial | 0.846852 | 0.003976 |

| | | | |
|---|---|---|---|
| **WDR1** | WD repeat-containing protein 1 | 0.303145 | 0.003991 |
| **LRRC59** | Leucine-rich repeat-containing protein 59 | 0.635774 | 0.004044 |
| **MRPL13** | 39S ribosomal protein L13, mitochondrial | 0.367811 | 0.004076 |
| **FAM162A** | Protein FAM162A | 0.574087 | 0.004112 |
| **ULBP2** | UL16-binding protein 2 | 0.45517 | 0.004115 |
| **TIMM8B** | Mitochondrial import inner membrane translocase subunit Tim8 B | 0.691126 | 0.00414 |
| **FXR2** | RNA-binding protein FXR2 | 0.254763 | 0.004154 |
| **PTMA** | Prothymosin alpha | 0.444214 | 0.004161 |
| **CBX1** | Chromobox protein homolog 1 | 0.782293 | 0.004178 |
| **HAUS1** | HAUS augmin-like complex subunit 1 | 0.601134 | 0.004214 |
| **LMNA** | Prelamin-A/C | 0.594805 | 0.004276 |
| **BZW2** | eIF5-mimic protein 1 | 0.20439 | 0.004318 |
| **STXBP4** | Syntaxin-binding protein 4 | 0.281959 | 0.004333 |
| **MAP7D1** | MAP7 domain-containing protein 1 | 0.447239 | 0.004367 |
| **PDIA3** | Protein disulfide-isomerase A3 | 0.397522 | 0.004375 |
| **NDUFB5** | NADH dehydrogenase [ubiquinone] 1 beta subcomplex subunit 5, mitochondrial | 0.52962 | 0.004387 |
| **TTC19** | Tetratricopeptide repeat protein 19, mitochondrial | 0.426162 | 0.004409 |
| **IRF2BP2** | Interferon regulatory factor 2-binding protein 2 | 0.629402 | 0.004534 |
| **RCN3** | Reticulocalbin-3 | 1.233187 | 0.004538 |
| **TPX2** | Targeting protein for Xklp2 | 1.092522 | 0.004558 |
| **GULP1** | PTB domain-containing engulfment adapter protein 1 | 0.23188 | 0.004581 |
| **KRR1** | KRR1 small subunit processome component homolog | 0.425184 | 0.004612 |
| **BSDC1** | BSD domain-containing protein 1 | 0.678958 | 0.004618 |
| **GCSH** | Glycine cleavage system H protein, mitochondrial | 1.0951 | 0.004627 |
| **CD55** | Complement decay-accelerating factor | 0.62929 | 0.004638 |

| | | | |
|---|---|---|---|
| **TRIP6** | Thyroid receptor-interacting protein 6 | 1.05806 | 0.004643 |
| **RAPH1** | Ras-associated and pleckstrin homology domains-containing protein 1 | 0.453109 | 0.004698 |
| **NECTIN2** | Nectin-2 | 0.499258 | 0.004716 |
| **PRPF38A** | Pre-mRNA-splicing factor 38A | 0.591781 | 0.004726 |
| **DNAJC17** | DnaJ homolog subfamily C member 17 | 0.738027 | 0.004739 |
| **COX5A** | Cytochrome c oxidase subunit 5A, mitochondrial | 0.305183 | 0.004772 |
| **RPRD1A** | Regulation of nuclear pre-mRNA domain-containing protein 1A | 0.353741 | 0.004837 |
| **SNRPA** | U1 small nuclear ribonucleoprotein A | 0.540325 | 0.004846 |
| **NES** | Nestin | 0.48867 | 0.00488 |
| **RPL37A** | 60S ribosomal protein L37a | 1.194444 | 0.004936 |
| **PHF10** | PHD finger protein 10 | 0.635085 | 0.004965 |
| **CBX3** | Chromobox protein homolog 3 | 0.733225 | 0.004985 |
| **ALDOA** | Fructose-bisphosphate aldolase A | 0.301877 | 0.004988 |
| **SOD1** | Superoxide dismutase [Cu-Zn] | 0.982132 | 0.004992 |
| **SYNPO** | Synaptopodin | 0.707104 | 0.005006 |
| **YWHAH** | 14-3-3 protein eta | 0.50916 | 0.005022 |
| **GOLIM4** | Golgi integral membrane protein 4 | 0.376983 | 0.005024 |
| **RPL19** | 60S ribosomal protein L19 | 0.476174 | 0.00503 |
| **TIMP3** | Metalloproteinase inhibitor 3 | 0.142731 | 0.005065 |
| **TSFM** | Elongation factor Ts, mitochondrial | 0.256807 | 0.005097 |
| **KIAA1191** | Putative monooxygenase p33MONOX | 0.400414 | 0.005134 |
| **ZNF446** | Zinc finger protein 446 | 0.387877 | 0.005204 |
| **EPHB4** | Ephrin type-B receptor 4 | 0.409151 | 0.005229 |
| **CHERP** | Calcium homeostasis endoplasmic reticulum protein | 0.307021 | 0.005235 |
| **NSMCE4A** | Non-structural maintenance of chromosomes element 4 homolog A | 0.216941 | 0.005241 |

| | | | |
|---|---|---|---|
| **SUPT7L** | STAGA complex 65 subunit gamma | 0.797958 | 0.005242 |
| **BCL2L1** | Bcl-2-like protein 1 | 0.455292 | 0.005279 |
| **RABIF** | Guanine nucleotide exchange factor MSS4 | 0.984147 | 0.005289 |
| **NDUFS4** | NADH dehydrogenase [ubiquinone] iron-sulfur protein 4, mitochondrial | 0.766194 | 0.005303 |
| **SEZ6L2** | Seizure 6-like protein 2 | 0.468942 | 0.005318 |
| **POLR1H** | DNA-directed RNA polymerase I subunit RPA12 | 2.490707 | 0.005322 |
| **EMC8** | ER membrane protein complex subunit 8 | 0.301267 | 0.005353 |
| **CXXC1** | CXXC-type zinc finger protein 1 | 0.356655 | 0.005355 |
| **KIF2C** | Kinesin-like protein KIF2C | 0.360843 | 0.005386 |
| **CALR** | Calreticulin | 0.854097 | 0.0054 |
| **CPSF7** | Cleavage and polyadenylation specificity factor subunit 7 | 0.594996 | 0.005419 |
| **TPBG** | Trophoblast glycoprotein | 0.688034 | 0.005462 |
| **MIA3** | Transport and Golgi organization protein 1 homolog | 0.205787 | 0.005496 |
| **NPTN** | Neuroplastin | 1.244945 | 0.005503 |
| **CANX** | Calnexin | 0.432641 | 0.005505 |
| **TAB3** | TGF-beta-activated kinase 1 and MAP3K7-binding protein 3 | 0.270335 | 0.00553 |
| **SKA3** | Spindle and kinetochore-associated protein 3 | 0.704893 | 0.005571 |
| **CAPS** | Calcyphosin | 0.850416 | 0.005609 |
| **CSRP1** | Cysteine and glycine-rich protein 1 | 0.741394 | 0.005663 |
| **ITFG1** | T-cell immunomodulatory protein | 0.592676 | 0.005681 |
| **MRPL24** | 39S ribosomal protein L24, mitochondrial | 0.215388 | 0.00569 |
| **SETD7** | Histone-lysine N-methyltransferase SETD7 | 0.562426 | 0.005694 |
| **YWHAB** | 14-3-3 protein beta/alpha | 0.460827 | 0.005695 |
| **COA7** | Cytochrome c oxidase assembly factor 7 | 0.873577 | 0.005732 |
| **PITPNB** | Phosphatidylinositol transfer protein beta isoform | 0.210608 | 0.005741 |

| | | | |
|---|---|---|---|
| **HMCES** | Abasic site processing protein HMCES | 0.378889 | 0.005749 |
| **WDR70** | WD repeat-containing protein 70 | 0.31049 | 0.005761 |
| **NUDC** | Nuclear migration protein nudC | 0.356204 | 0.005775 |
| **DPF2** | Zinc finger protein ubi-d4 | 0.536883 | 0.005782 |
| **NDUFB8** | NADH dehydrogenase [ubiquinone] 1 beta subcomplex subunit 8, mitochondrial | 0.648895 | 0.005787 |
| **SIX1** | Homeobox protein SIX1 | 0.736468 | 0.005816 |
| **MAVS** | Mitochondrial antiviral-signaling protein | 0.40986 | 0.005823 |
| **PDCD10** | Programmed cell death protein 10 | 0.590601 | 0.005883 |
| **DNAJC3** | DnaJ homolog subfamily C member 3 | 0.438638 | 0.005901 |
| **IGF2R** | Cation-independent mannose-6-phosphate receptor | 0.20107 | 0.005946 |
| **DNAJC7** | DnaJ homolog subfamily C member 7 | 0.151595 | 0.005983 |
| **UBFD1** | Ubiquitin domain-containing protein UBFD1 | 0.702052 | 0.005983 |
| **NCOA5** | Nuclear receptor coactivator 5 | 0.259839 | 0.006001 |
| **PLIN3** | Perilipin-3 | 0.562882 | 0.006112 |
| **ATXN2L** | Ataxin-2-like protein | 0.396041 | 0.006116 |
| **SLC31A1** | High affinity copper uptake protein 1 | 1.05406 | 0.006133 |
| **HSBP1** | Heat shock factor-binding protein 1 | 0.273827 | 0.006171 |
| **BAG2** | BAG family molecular chaperone regulator 2 | 0.731529 | 0.006197 |
| **TESC** | Calcineurin B homologous protein 3 | 0.526476 | 0.006274 |
| **PIH1D1** | PIH1 domain-containing protein 1 | 0.339163 | 0.006332 |
| **PPID** | Peptidyl-prolyl cis-trans isomerase D | 0.310116 | 0.006333 |
| **EIF5** | Eukaryotic translation initiation factor 5 | 0.337302 | 0.00636 |
| **ITGA5** | Integrin alpha-5 | 0.517128 | 0.006417 |
| **CCNA2** | Cyclin-A2 | 0.611501 | 0.006439 |
| **POLR3G** | DNA-directed RNA polymerase III subunit RPC7 | 0.599953 | 0.006503 |

| | | | |
|---|---|---|---|
| **SAFB2** | Scaffold attachment factor B2 | 0.603695 | 0.006514 |
| **LYPLAL1** | Lysophospholipase-like protein 1 | 0.601273 | 0.00653 |
| **DHFR** | Dihydrofolate reductase | 0.514668 | 0.006534 |
| **PSMD7** | 26S proteasome non-ATPase regulatory subunit 7 | 0.654005 | 0.006557 |
| **SASH1** | SAM and SH3 domain-containing protein 1 | 0.374479 | 0.006579 |
| **CSNK2B** | Casein kinase II subunit beta | 0.465028 | 0.006594 |
| **RALY** | RNA-binding protein Raly | 0.805532 | 0.006598 |
| **NFATC2IP** | NFATC2-interacting protein | 0.488072 | 0.006637 |
| **LPXN** | Leupaxin | 0.66187 | 0.006724 |
| **ACBD7** | Acyl-CoA-binding domain-containing protein 7 | 0.59144 | 0.006805 |
| **FABP5** | Fatty acid-binding protein 5 | 0.767122 | 0.006821 |
| **FRMD4A** | FERM domain-containing protein 4A | 0.341462 | 0.006847 |
| **RABEP2** | Rab GTPase-binding effector protein 2 | 0.431172 | 0.006889 |
| **P4HB** | Protein disulfide-isomerase | 0.440326 | 0.0069 |
| **AAGAB** | Alpha- and gamma-adaptin-binding protein p34 | 1.118029 | 0.006908 |
| **DLGAP4** | Disks large-associated protein 4 | 0.376941 | 0.006929 |
| **EEF1D** | Elongation factor 1-delta | 0.432456 | 0.006971 |
| **RANBP1** | Ran-specific GTPase-activating protein | 1.28571 | 0.006981 |
| **GPATCH11** | G patch domain-containing protein 11 | 0.563851 | 0.007013 |
| **MAN2B1** | Lysosomal alpha-mannosidase | 0.342142 | 0.007021 |
| **CUSTOS** | Protein CUSTOS | 0.52359 | 0.007086 |
| **CMPK1** | UMP-CMP kinase | 0.548572 | 0.007108 |
| **XPC** | DNA repair protein complementing XP-C cells | 0.412314 | 0.007113 |
| **RBM38** | RNA-binding protein 38 | 0.914448 | 0.00712 |
| **HGH1** | Protein HGH1 homolog | 0.278059 | 0.007135 |

| | | | |
|---|---|---|---|
| **TRIR** | Telomerase RNA component interacting RNase | 0.313604 | 0.007155 |
| **KIF27** | Kinesin-like protein KIF27 | 0.61943 | 0.007213 |
| **HLA-A** | HLA class I histocompatibility antigen, A alpha chain | 0.808716 | 0.007254 |
| **HAUS8** | HAUS augmin-like complex subunit 8 | 0.643659 | 0.007257 |
| **MYL9** | Myosin regulatory light polypeptide 9 | 0.637184 | 0.007268 |
| **ZCCHC9** | Zinc finger CCHC domain-containing protein 9 | 0.532399 | 0.007307 |
| **PRCP** | Lysosomal Pro-X carboxypeptidase | 0.827345 | 0.00733 |
| **PAK1** | Serine/threonine-protein kinase PAK 1 | 0.32274 | 0.007342 |
| **TIPRL** | TIP41-like protein | 0.251699 | 0.007426 |
| **CD44** | CD44 antigen | 0.383079 | 0.007445 |
| **PDLIM4** | PDZ and LIM domain protein 4 | 0.993012 | 0.007485 |
| **BST1** | ADP-ribosyl cyclase/cyclic ADP-ribose hydrolase 2 | 0.634646 | 0.007493 |
| **TOM1L1** | TOM1-like protein 1 | 0.434268 | 0.007526 |
| **BRD7** | Bromodomain-containing protein 7 | 0.235195 | 0.007542 |
| **ABI1** | Abl interactor 1 | 0.468058 | 0.007552 |
| **LASP1** | LIM and SH3 domain protein 1 | 0.523333 | 0.007577 |
| **ZNF512B** | Zinc finger protein 512B | 0.305513 | 0.007601 |
| **NGFR** | Tumor necrosis factor receptor superfamily member 16 | 0.83421 | 0.007634 |
| **PLXNB2** | Plexin-B2 | 0.426447 | 0.007664 |
| **PPM1G** | Protein phosphatase 1G | 0.557119 | 0.007667 |
| **GUK1** | Guanylate kinase | 0.415347 | 0.007672 |
| **MAGEA10** | Melanoma-associated antigen 10 | 0.642673 | 0.007677 |
| **HROB** | Homologous recombination OB-fold protein | 0.256251 | 0.0077 |
| **ZC2HC1A** | Zinc finger C2HC domain-containing protein 1A | 0.481012 | 0.007709 |
| **CEP131** | Centrosomal protein of 131 kDa | 0.373361 | 0.007875 |

| | | | |
|---|---|---|---|
| **LZTS2** | Leucine zipper putative tumor suppressor 2 | 0.543715 | 0.007877 |
| **GYG1** | Glycogenin-1 | 0.357896 | 0.007913 |
| **KRT18** | Keratin, type I cytoskeletal 18 | 0.516055 | 0.007933 |
| **POLR1D** | Protein POLR1D, isoform 2 | 0.705259 | 0.007935 |
| **MESD** | LRP chaperone MESD | 0.504157 | 0.007961 |
| **FAM3C** | Protein FAM3C | 0.388541 | 0.007972 |
| **PDLIM2** | PDZ and LIM domain protein 2 | 0.773437 | 0.00802 |
| **CHMP2B** | Charged multivesicular body protein 2b | 0.507644 | 0.008027 |
| **PPP1R18** | Phostensin | 0.455158 | 0.008028 |
| **U2AF2** | Splicing factor U2AF 65 kDa subunit | 0.472983 | 0.008083 |
| **CCDC25** | Coiled-coil domain-containing protein 25 | 0.490155 | 0.008113 |
| **CLINT1** | Clathrin interactor 1 | 0.237311 | 0.008125 |
| **RTRAF** | RNA transcription, translation and transport factor protein | 0.60872 | 0.008127 |
| **POU2F3** | POU domain, class 2, transcription factor 3 | 1.005249 | 0.008303 |
| **RNASET2** | Ribonuclease T2 | 0.628825 | 0.00838 |
| **SUMF2** | Inactive C-alpha-formylglycine-generating enzyme 2 | 0.348738 | 0.008406 |
| **TPD52L1** | Tumor protein D53 | 0.603705 | 0.008434 |
| **UBAC1** | Ubiquitin-associated domain-containing protein 1 | 0.677589 | 0.008464 |
| **SNRPD2** | Small nuclear ribonucleoprotein Sm D2 | 0.380225 | 0.008478 |
| **RRBP1** | Ribosome-binding protein 1 | 0.455184 | 0.008486 |
| **TAX1BP1** | Tax1-binding protein 1 | 0.191694 | 0.00852 |
| **WDR55** | WD repeat-containing protein 55 | 0.506345 | 0.008548 |
| **BCAM** | Basal cell adhesion molecule | 0.299833 | 0.008641 |
| **MAPRE1** | Microtubule-associated protein RP/EB family member 1 | 0.625949 | 0.00871 |
| **CDK5RAP2** | CDK5 regulatory subunit-associated protein 2 | 0.139091 | 0.008821 |

| | | | |
|---|---|---|---|
| **ISCU** | Iron-sulfur cluster assembly enzyme ISCU | 0.543316 | 0.008825 |
| **CD2BP2** | CD2 antigen cytoplasmic tail-binding protein 2 | 0.561862 | 0.00897 |
| **IST1** | IST1 homolog | 0.679527 | 0.008972 |
| **NECTIN1** | Nectin-1 | 0.622975 | 0.009025 |
| **SPECC1** | Cytospin-B | 0.459141 | 0.009066 |
| **DENR** | Density-regulated protein | 0.942546 | 0.009112 |
| **MRPL58** | Peptidyl-tRNA hydrolase ICT1, mitochondrial | 1.20987 | 0.009126 |
| **ENY2** | Transcription and mRNA export factor ENY2 | 0.315718 | 0.009165 |
| **RAB18** | Ras-related protein Rab-18 | 0.122643 | 0.009171 |
| **PTK7** | Inactive tyrosine-protein kinase 7 | 0.640302 | 0.009183 |
| **CCDC102A** | Coiled-coil domain-containing protein 102A | 0.324326 | 0.009218 |
| **PPP1R12A** | Protein phosphatase 1 regulatory subunit 12A | 0.331296 | 0.009259 |
| **RILPL1** | RILP-like protein 1 | 0.525962 | 0.009275 |
| **FKBP4** | Peptidyl-prolyl cis-trans isomerase FKBP4 | 0.315803 | 0.00934 |
| **TOR1AIP1** | Torsin-1A-interacting protein 1 | 0.391302 | 0.009346 |
| **BAIAP2** | Brain-specific angiogenesis inhibitor 1-associated protein 2 | 0.528868 | 0.009371 |
| **SH3BGRL3** | SH3 domain-binding glutamic acid-rich-like protein 3 | 0.697878 | 0.009411 |
| **RBM34** | RNA-binding protein 34 | 0.562359 | 0.009413 |
| **TSSC4** | Protein TSSC4 | 0.568962 | 0.009559 |
| **HSPA5** | Endoplasmic reticulum chaperone BiP | 0.434012 | 0.009567 |
| **RPS12** | 40S ribosomal protein S12 | 0.563192 | 0.009608 |
| **CNOT3** | CCR4-NOT transcription complex subunit 3 | 0.488467 | 0.009664 |
| **CKB** | Creatine kinase B-type | 0.673446 | 0.0097 |
| **LLPH** | Protein LLP homolog | 1.358426 | 0.009815 |
| **YAP1** | Transcriptional coactivator YAP1 | 0.551585 | 0.009825 |
| **SLC30A1** | Proton-coupled zinc antiporter SLC30A1 | 0.507873 | 0.009836 |
| **CDCA8** | Borealin | 0.829272 | 0.009877 |
| **ING4** | Inhibitor of growth protein 4 | 0.600019 | 0.009884 |

| | | | |
|---|---|---|---|
| **CEP170B** | Centrosomal protein of 170 kDa protein B | 0.492368 | 0.009919 |
| **RRP1** | Ribosomal RNA processing protein 1 homolog A | 0.393114 | 0.009921 |
| **POMK** | Protein O-mannose kinase | 0.794931 | 0.009937 |
| **HIKESHI** | Protein Hikeshi | 0.284257 | 0.009938 |
| **TRAFD1** | TRAF-type zinc finger domain-containing protein 1 | 0.394277 | 0.00996 |
| **SREK1** | Splicing regulatory glutamine/lysine-rich protein 1 | 0.353795 | 0.009978 |
| **MANF** | Mesencephalic astrocyte-derived neurotrophic factor | 0.66346 | 0.009985 |
| **BCAS2** | Pre-mRNA-splicing factor SPF27 | 0.391941 | 0.009992 |
| **EIF4B** | Eukaryotic translation initiation factor 4B | 0.699888 | 0.009998 |

**Supplemental Table 6. Proteins which were downregulated by GEM treatment in A375 melanoma cells.**

| Downregulated proteins after 24 hours of treatment with 20 nM GEM | | | |
|---|---|---|---|
| **Gene name for protein** | **Description** | **Fold change (log2 form)** | **t-test P value** |
| **CHEK1** | Serine/threonine-protein kinase Chk1 | -0.9569 | 9.96E-06 |
| **FAM193A** | Protein FAM193A | -2.08635 | 3.94E-05 |
| **RABGAP1** | Rab GTPase-activating protein 1 | -0.41115 | 5.96E-05 |
| **CEBPZ** | CCAAT/enhancer-binding protein zeta | -0.14686 | 8.54E-05 |
| **ATF7** | Cyclic AMP-dependent transcription factor ATF-7 | -1.01636 | 1E-04 |
| **YARS2** | Tyrosine--tRNA ligase, mitochondrial | -0.37721 | 0.0001 |
| **CDC42BPA** | Serine/threonine-protein kinase MRCK alpha | -0.68028 | 0.00011 |
| **ARAP1** | Arf-GAP with Rho-GAP domain, ANK repeat and PH domain-containing protein 1 | -0.25689 | 0.000113 |
| **COG2** | Conserved oligomeric Golgi complex subunit 2 | -0.63424 | 0.000121 |
| **ATAD2** | ATPase family AAA domain-containing protein 2 | -0.94415 | 0.000123 |
| **SAPCD2** | Suppressor APC domain-containing protein 2 | -1.00513 | 0.000143 |
| **USP24** | Ubiquitin carboxyl-terminal hydrolase 24 | -0.53049 | 0.000186 |
| **SOGA1** | Protein SOGA1 | -0.69801 | 0.000192 |
| **PARP4** | Protein mono-ADP-ribosyltransferase PARP4 | -0.27733 | 0.000201 |
| **GTF3C1** | General transcription factor 3C polypeptide 1 | -0.6491 | 0.000234 |
| **UVRAG** | UV radiation resistance-associated gene protein | -0.56661 | 0.00024 |
| **TARS1** | Threonine--tRNA ligase 1, cytoplasmic | -0.37667 | 0.000257 |
| **RUFY1** | RUN and FYVE domain-containing protein 1 | -0.35716 | 0.00027 |
| **JAK1** | Tyrosine-protein kinase JAK1 | -0.51697 | 0.00028 |
| **KIF20A** | Kinesin-like protein KIF20A | -0.65178 | 0.000283 |
| **NDUFS8** | NADH dehydrogenase [ubiquinone] iron-sulfur protein 8, mitochondrial | -1.05445 | 0.000292 |
| **UCKL1** | Uridine-cytidine kinase-like 1 | -0.52169 | 0.000306 |
| **CEP192** | Centrosomal protein of 192 kDa | -0.74047 | 0.000311 |

| | | | |
|---|---|---|---|
| **KMT2D** | Histone-lysine N-methyltransferase 2D | -0.40264 | 0.000319 |
| **SPATA5L1** | Ribosome biogenesis protein SPATA5L1 | -0.81816 | 0.000345 |
| **PTPN9** | Tyrosine-protein phosphatase non-receptor type 9 | -0.52382 | 0.000366 |
| **CSDE1** | Cold shock domain-containing protein E1 | -0.50602 | 0.000385 |
| **OSTC** | Oligosaccharyltransferase complex subunit OSTC | -0.5642 | 0.000386 |
| **RASA1** | Ras GTPase-activating protein 1 | -0.64403 | 0.000401 |
| **CHD1** | Chromodomain-helicase-DNA-binding protein 1 | -0.35759 | 0.000443 |
| **DNAH11** | Dynein axonemal heavy chain 11 | -1.45757 | 0.000453 |
| **QRICH1** | Transcriptional regulator QRICH1 | -0.5238 | 0.000456 |
| **USP34** | Ubiquitin carboxyl-terminal hydrolase 34 | -0.31492 | 0.00049 |
| **H2BC1** | Histone H2B type 1-A | -1.74041 | 0.000515 |
| **PUS7L** | Pseudouridylate synthase PUS7L | -0.466 | 0.000521 |
| **GPHN** | Gephyrin | -0.41097 | 0.000544 |
| **ZKSCAN4** | Zinc finger protein with KRAB and SCAN domains 4 | -1.18341 | 0.000545 |
| **LRRC40** | Leucine-rich repeat-containing protein 40 | -0.57623 | 0.000545 |
| **OXSR1** | Serine/threonine-protein kinase OSR1 | -0.2193 | 0.000546 |
| **TRIP12** | E3 ubiquitin-protein ligase TRIP12 | -0.67423 | 0.000612 |
| **SSR3** | Translocon-associated protein subunit gamma | -0.4611 | 0.000628 |
| **DDX39A** | ATP-dependent RNA helicase DDX39A | -0.49684 | 0.000637 |
| **CDC123** | Cell division cycle protein 123 homolog | -0.47628 | 0.000663 |
| **SNAPC4** | snRNA-activating protein complex subunit 4 | -0.57696 | 0.000688 |
| **ADSS2** | Adenylosuccinate synthetase isozyme 2 | -0.23821 | 0.000692 |
| **METTL16** | RNA N6-adenosine-methyltransferase METTL16 | -0.54132 | 0.000694 |
| **KIF20B** | Kinesin-like protein KIF20B | -0.57787 | 0.000701 |
| **STT3B** | Dolichyl-diphosphooligosaccharide--protein glycosyltransferase subunit STT3B | -0.62551 | 0.000734 |
| **TXNL1** | Thioredoxin-like protein 1 | -0.31851 | 0.000761 |
| **TMED10** | Transmembrane emp24 domain-containing protein 10 | -0.4103 | 0.000768 |

| | | | |
|---|---|---|---|
| **TMX2** | Thioredoxin-related transmembrane protein 2 | -0.75396 | 0.000772 |
| **EIF3F** | Eukaryotic translation initiation factor 3 subunit F | -0.38827 | 0.000776 |
| **MALT1** | Mucosa-associated lymphoid tissue lymphoma translocation protein 1 | -0.67399 | 0.000804 |
| **GEMIN2** | Gem-associated protein 2 | -1.47252 | 0.00081 |
| **SIN3A** | Paired amphipathic helix protein Sin3a | -0.30825 | 0.00082 |
| **DCAF1** | DDB1- and CUL4-associated factor 1 | -0.38841 | 0.000824 |
| **SP2** | Transcription factor Sp2 | -1.09171 | 0.000863 |
| **DERL1** | Derlin-1 | -0.63972 | 0.000876 |
| **ACSL1** | Long-chain-fatty-acid--CoA ligase 1 | -0.52169 | 0.000878 |
| **SUPT6H** | Transcription elongation factor SPT6 | -0.30462 | 0.000878 |
| **CCT4** | T-complex protein 1 subunit delta | -0.46032 | 0.000948 |
| **RFC5** | Replication factor C subunit 5 | -0.40125 | 0.000966 |
| **TUBAL3** | Tubulin alpha chain-like 3 | -1.24279 | 0.000989 |
| **COASY** | Bifunctional coenzyme A synthase | -0.72324 | 0.000997 |
| **MACF1** | Microtubule-actin cross-linking factor 1, isoforms 1/2/3/4/5 | -0.46878 | 0.001012 |
| **CHD9** | Chromodomain-helicase-DNA-binding protein 9 | -0.49633 | 0.00102 |
| **WASHC5** | WASH complex subunit 5 | -0.41445 | 0.001024 |
| **GPAT3** | Glycerol-3-phosphate acyltransferase 3 | -0.43051 | 0.001038 |
| **LRRCC1** | Leucine-rich repeat and coiled-coil domain-containing protein 1 | -0.82491 | 0.001046 |
| **HECTD4** | Probable E3 ubiquitin-protein ligase HECTD4 | -0.55764 | 0.001057 |
| **ASNS** | Asparagine synthetase [glutamine-hydrolyzing] | -0.69765 | 0.001075 |
| **SRBD1** | S1 RNA-binding domain-containing protein 1 | -0.5732 | 0.00108 |
| **UBA5** | Ubiquitin-like modifier-activating enzyme 5 | -0.82359 | 0.001089 |
| **DIMT1** | Probable dimethyladenosine transferase | -0.43506 | 0.001168 |
| **PFAS** | Phosphoribosylformylglycinamidine synthase | -0.36986 | 0.001178 |
| **PPP2R1A** | Serine/threonine-protein phosphatase 2A 65 kDa regulatory subunit A alpha isoform | -0.44049 | 0.001208 |
| **XAB2** | Pre-mRNA-splicing factor SYF1 | -0.6813 | 0.001217 |

| | | | |
|---|---|---|---|
| **STK38L** | Serine/threonine-protein kinase 38-like | -0.63105 | 0.001219 |
| **POLR2A** | DNA-directed RNA polymerase II subunit RPB1 | -0.61483 | 0.00125 |
| **ARMC5** | Armadillo repeat-containing protein 5 | -1.32927 | 0.001254 |
| **ELMO2** | Engulfment and cell motility protein 2 | -0.87879 | 0.001274 |
| **MGRN1** | E3 ubiquitin-protein ligase MGRN1 | -0.37593 | 0.001279 |
| **TKFC** | Triokinase/FMN cyclase | -0.51365 | 0.001281 |
| **INPP4B** | Inositol polyphosphate 4-phosphatase type II | -0.83144 | 0.001318 |
| **CSNK2A1** | Casein kinase II subunit alpha | -0.26582 | 0.001341 |
| **JMJD1C** | Probable JmjC domain-containing histone demethylation protein 2C | -0.19686 | 0.001369 |
| **EXOSC1** | Exosome complex component CSL4 | -0.32289 | 0.001374 |
| **NOC3L** | Nucleolar complex protein 3 homolog | -0.27756 | 0.001383 |
| **RAB3GAP1** | Rab3 GTPase-activating protein catalytic subunit | -0.43761 | 0.001401 |
| **LTN1** | E3 ubiquitin-protein ligase listerin | -0.64415 | 0.001404 |
| **ZNF219** | Zinc finger protein 219 | -0.59931 | 0.001412 |
| **DDX47** | Probable ATP-dependent RNA helicase DDX47 | -0.44312 | 0.001417 |
| **PPP2R5E** | Serine/threonine-protein phosphatase 2A 56 kDa regulatory subunit epsilon isoform | -0.58499 | 0.00145 |
| **ECPAS** | Proteasome adapter and scaffold protein ECM29 | -0.42239 | 0.001456 |
| **ELP5** | Elongator complex protein 5 | -0.58483 | 0.001484 |
| **SBNO1** | Protein strawberry notch homolog 1 | -0.71683 | 0.001488 |
| **TMEM222** | Transmembrane protein 222 | -0.52428 | 0.001523 |
| **NKAPD1** | Uncharacterized protein NKAPD1 | -0.36959 | 0.00157 |
| **DHX30** | ATP-dependent RNA helicase DHX30 | -0.5025 | 0.001583 |
| **SRSF2** | Serine/arginine-rich splicing factor 2 | -1.25384 | 0.001603 |
| **TIMM10** | Mitochondrial import inner membrane translocase subunit Tim10 | -1.07738 | 0.00161 |
| **ZBTB21** | Zinc finger and BTB domain-containing protein 21 | -1.05629 | 0.001613 |
| **RANGAP1** | Ran GTPase-activating protein 1 | -0.43056 | 0.001672 |

| | | | |
|---|---|---|---|
| **ELAC2** | Zinc phosphodiesterase ELAC protein 2 | -0.37643 | 0.001677 |
| **TRIM41** | E3 ubiquitin-protein ligase TRIM41 | -0.4724 | 0.001681 |
| **RAB43** | Ras-related protein Rab-43 | -0.53921 | 0.001682 |
| **SPATA5** | Ribosome biogenesis protein SPATA5 | -0.45035 | 0.00172 |
| **IPO5** | Importin-5 | -0.44825 | 0.001722 |
| **PLK1** | Serine/threonine-protein kinase PLK1 | -0.4144 | 0.001729 |
| **DUS3L** | tRNA-dihydrouridine(47) synthase [NAD(P)(+)]-like | -0.34576 | 0.001749 |
| **TBCE** | Tubulin-specific chaperone E | -0.40778 | 0.001752 |
| **PAK4** | Serine/threonine-protein kinase PAK 4 | -0.49842 | 0.00176 |
| **DHPS** | Deoxyhypusine synthase | -0.5101 | 0.001792 |
| **ARAF** | Serine/threonine-protein kinase A-Raf | -0.47586 | 0.001798 |
| **DHX16** | Pre-mRNA-splicing factor ATP-dependent RNA helicase DHX16 | -0.45541 | 0.001801 |
| **CAMK2G** | Calcium/calmodulin-dependent protein kinase type II subunit gamma | -0.49349 | 0.001802 |
| **GALNS** | N-acetylgalactosamine-6-sulfatase | -0.65478 | 0.00184 |
| **XRCC6** | X-ray repair cross-complementing protein 6 | -0.50888 | 0.00185 |
| **KDM5B** | Lysine-specific demethylase 5B | -0.98875 | 0.001853 |
| **FAF1** | FAS-associated factor 1 | -0.36333 | 0.001868 |
| **TPM2** | Tropomyosin beta chain | -0.95094 | 0.001869 |
| **ATG9A** | Autophagy-related protein 9A | -0.36164 | 0.001893 |
| **SCML2** | Sex comb on midleg-like protein 2 | -0.55696 | 0.001923 |
| **COPG2** | Coatomer subunit gamma-2 | -0.57642 | 0.00194 |
| **MCM5** | DNA replication licensing factor MCM5 | -0.44755 | 0.001986 |
| **NOA1** | Nitric oxide-associated protein 1 | -0.48534 | 0.002004 |
| **CAP2** | Adenylyl cyclase-associated protein 2 | -0.36491 | 0.002094 |
| **CREBBP** | CREB-binding protein | -0.61914 | 0.002101 |
| **H2BC18** | Histone H2B type 2-F | -1.20127 | 0.002117 |
| **RRN3** | RNA polymerase I-specific transcription initiation factor RRN3 | -1.02042 | 0.002132 |
| **RGP1** | RAB6A-GEF complex partner protein 2 | -0.70301 | 0.002153 |
| **NUDT16L1** | Tudor-interacting repair regulator protein | -0.60371 | 0.002193 |
| **TSHZ2** | Teashirt homolog 2 | -1.42051 | 0.002221 |

| | | | |
|---|---|---|---|
| **LRPPRC** | Leucine-rich PPR motif-containing protein, mitochondrial | -0.44562 | 0.002226 |
| **TUBB3** | Tubulin beta-3 chain | -0.56657 | 0.002261 |
| **ATP5F1B** | ATP synthase subunit beta, mitochondrial | -0.37705 | 0.002267 |
| **DDX55** | ATP-dependent RNA helicase DDX55 | -0.48213 | 0.002276 |
| **ZNF146** | Zinc finger protein OZF | -0.58869 | 0.002292 |
| **SCAF8** | SR-related and CTD-associated factor 8 | -0.5127 | 0.00232 |
| **DARS1** | Aspartate--tRNA ligase, cytoplasmic | -0.21936 | 0.002331 |
| **POLR1B** | DNA-directed RNA polymerase I subunit RPA2 | -0.39915 | 0.002335 |
| **YME1L1** | ATP-dependent zinc metalloprotease YME1L1 | -0.37512 | 0.002337 |
| **MTHFD1L** | Monofunctional C1-tetrahydrofolate synthase, mitochondrial | -0.50202 | 0.002387 |
| **SLC25A13** | Electrogenic aspartate/glutamate antiporter SLC25A13, mitochondrial | -0.45292 | 0.002393 |
| **PPP2R5D** | Serine/threonine-protein phosphatase 2A 56 kDa regulatory subunit delta isoform | -0.87672 | 0.0024 |
| **BRCA2** | Breast cancer type 2 susceptibility protein | -0.55813 | 0.002419 |
| **NFIC** | Nuclear factor 1 C-type | -0.37583 | 0.002452 |
| **TIMELESS** | Protein timeless homolog | -0.45736 | 0.00247 |
| **EIF4A1** | Eukaryotic initiation factor 4A-I | -0.5566 | 0.002485 |
| **HRAS** | GTPase HRas | -0.37411 | 0.002486 |
| **EXT2** | Exostosin-2 | -1.01516 | 0.002495 |
| **C2CD5** | C2 domain-containing protein 5 | -0.88418 | 0.002499 |
| **ZNF385A** | Zinc finger protein 385A | -0.49552 | 0.002529 |
| **PAPOLG** | Poly(A) polymerase gamma | -0.84995 | 0.002547 |
| **EMC9** | ER membrane protein complex subunit 9 | -1.4173 | 0.002549 |
| **BICRA** | BRD4-interacting chromatin-remodeling complex-associated protein | -0.58687 | 0.002553 |
| **BCOR** | BCL-6 corepressor | -0.41448 | 0.002588 |
| **GFM2** | Ribosome-releasing factor 2, mitochondrial | -0.70975 | 0.002614 |
| **COX7A2L** | Cytochrome c oxidase subunit 7A-related protein, mitochondrial | -0.40406 | 0.002641 |
| **CUL3** | Cullin-3 | -0.44782 | 0.002699 |

| | | | |
|---|---|---|---|
| **VPS51** | Vacuolar protein sorting-associated protein 51 homolog | -0.717 | 0.002699 |
| **USP39** | U4/U6.U5 tri-snRNP-associated protein 2 | -0.55139 | 0.002706 |
| **HYLS1** | Centriolar and ciliogenesis-associated protein HYLS1 | -1.08308 | 0.002721 |
| **COA5** | Cytochrome c oxidase assembly factor 5 | -0.65452 | 0.002728 |
| **VAC14** | Protein VAC14 homolog | -0.48493 | 0.00274 |
| **PARP12** | Protein mono-ADP-ribosyltransferase PARP12 | -0.8424 | 0.002748 |
| **PRAF2** | PRA1 family protein 2 | -0.66993 | 0.00275 |
| **SLC25A12** | Electrogenic aspartate/glutamate antiporter SLC25A12, mitochondrial | -0.60082 | 0.002768 |
| **FASTKD5** | FAST kinase domain-containing protein 5, mitochondrial | -0.50499 | 0.002777 |
| **BCAR3** | Breast cancer anti-estrogen resistance protein 3 | -0.60324 | 0.002789 |
| **MAP4K4** | Mitogen-activated protein kinase kinase kinase kinase 4 | -0.43347 | 0.002806 |
| **OGT** | UDP-N-acetylglucosamine--peptide N-acetylglucosaminyltransferase 110 kDa subunit | -0.6665 | 0.002859 |
| **UBE4A** | Ubiquitin conjugation factor E4 A | -0.46892 | 0.002867 |
| **CDKN2AIPNL** | CDKN2AIP N-terminal-like protein | -0.46255 | 0.002929 |
| **ATP5PF** | ATP synthase-coupling factor 6, mitochondrial | -0.83529 | 0.002934 |
| **FBXO7** | F-box only protein 7 | -0.81567 | 0.002949 |
| **ATAD5** | ATPase family AAA domain-containing protein 5 | -0.63401 | 0.002954 |
| **ZMYM5** | Zinc finger MYM-type protein 5 | -0.61404 | 0.002957 |
| **GPS1** | COP9 signalosome complex subunit 1 | -0.40962 | 0.002964 |
| **VPS11** | Vacuolar protein sorting-associated protein 11 homolog | -0.61508 | 0.00299 |
| **AQR** | RNA helicase aquarius | -0.51851 | 0.002995 |
| **DLST** | Dihydrolipoyllysine-residue succinyltransferase component of 2-oxoglutarate dehydrogenase complex, mitochondrial | -0.41132 | 0.003004 |
| **FARSA** | Phenylalanine--tRNA ligase alpha subunit | -0.33482 | 0.003032 |
| **TMEM43** | Transmembrane protein 43 | -0.30548 | 0.003082 |
| **MBOAT7** | Lysophospholipid acyltransferase 7 | -0.97182 | 0.003096 |

| | | | |
|---|---|---|---|
| **HSPA8** | Heat shock cognate 71 kDa protein | -0.55301 | 0.003096 |
| **ZWILCH** | Protein zwilch homolog | -0.32509 | 0.003102 |
| **EXOC2** | Exocyst complex component 2 | -0.59648 | 0.003162 |
| **TEX10** | Testis-expressed protein 10 | -0.26227 | 0.003165 |
| **SUPV3L1** | ATP-dependent RNA helicase SUPV3L1, mitochondrial | -0.59014 | 0.00317 |
| **ACAA2** | 3-ketoacyl-CoA thiolase, mitochondrial | -0.26793 | 0.003174 |
| **GBF1** | Golgi-specific brefeldin A-resistance guanine nucleotide exchange factor 1 | -0.34648 | 0.003279 |
| **MLH1** | DNA mismatch repair protein Mlh1 | -0.55721 | 0.003281 |
| **NOL6** | Nucleolar protein 6 | -0.5038 | 0.003307 |
| **HSP90AB1** | Heat shock protein HSP 90-beta | -0.25855 | 0.00332 |
| **CNOT1** | CCR4-NOT transcription complex subunit 1 | -0.52072 | 0.003335 |
| **WDR46** | WD repeat-containing protein 46 | -0.38328 | 0.003343 |
| **SACS** | Sacsin | -0.54584 | 0.003352 |
| **APPL1** | DCC-interacting protein 13-alpha | -0.38473 | 0.003354 |
| **TEFM** | Transcription elongation factor, mitochondrial | -1.09676 | 0.003374 |
| **MCMBP** | Mini-chromosome maintenance complex-binding protein | -0.51433 | 0.003388 |
| **CYFIP1** | Cytoplasmic FMR1-interacting protein 1 | -0.45483 | 0.0034 |
| **WDR91** | WD repeat-containing protein 91 | -0.6158 | 0.003407 |
| **SMCHD1** | Structural maintenance of chromosomes flexible hinge domain-containing protein 1 | -0.49534 | 0.003439 |
| **PSMC2** | 26S proteasome regulatory subunit 7 | -0.40098 | 0.003445 |
| **DDX28** | Probable ATP-dependent RNA helicase DDX28 | -0.84904 | 0.003446 |
| **POLD1** | DNA polymerase delta catalytic subunit | -0.6532 | 0.003466 |
| **ZW10** | Centromere/kinetochore protein zw10 homolog | -0.5098 | 0.003467 |
| **ULK1** | Serine/threonine-protein kinase ULK1 | -1.0158 | 0.003481 |
| **EIF3L** | Eukaryotic translation initiation factor 3 subunit L | -0.50558 | 0.003517 |
| **APAF1** | Apoptotic protease-activating factor 1 | -0.38423 | 0.003537 |
| **FOXD3** | Forkhead box protein D3 | -0.82892 | 0.003538 |
| **MARS1** | Methionine--tRNA ligase, cytoplasmic | -0.72002 | 0.003544 |
| **NHLRC2** | NHL repeat-containing protein 2 | -0.89294 | 0.003544 |

| | | | |
|---|---|---|---|
| **TIMM17B** | Mitochondrial import inner membrane translocase subunit Tim17-B | -0.66598 | 0.003549 |
| **ABCF2** | ATP-binding cassette sub-family F member 2 | -0.47802 | 0.003574 |
| **LCMT2** | tRNA wybutosine-synthesizing protein 4 | -0.55754 | 0.00361 |
| **MVP** | Major vault protein | -0.47939 | 0.00363 |
| **KIF14** | Kinesin-like protein KIF14 | -0.38593 | 0.003663 |
| **SEC24C** | Protein transport protein Sec24C | -0.555 | 0.003678 |
| **EHD3** | EH domain-containing protein 3 | -0.41463 | 0.003688 |
| **MAP4K5** | Mitogen-activated protein kinase kinase kinase kinase 5 | -0.67523 | 0.003741 |
| **GNA13** | Guanine nucleotide-binding protein subunit alpha-13 | -0.42144 | 0.003752 |
| **SRCAP** | Helicase SRCAP | -0.52057 | 0.003769 |
| **RBL1** | Retinoblastoma-like protein 1 | -2.03506 | 0.003772 |
| **COMMD3** | COMM domain-containing protein 3 | -0.27069 | 0.003775 |
| **PTPRA** | Receptor-type tyrosine-protein phosphatase alpha | -0.78527 | 0.003861 |
| **PRUNE1** | Exopolyphosphatase PRUNE1 | -0.46868 | 0.003921 |
| **MTREX** | Exosome RNA helicase MTR4 | -0.51212 | 0.00394 |
| **HDAC3** | Histone deacetylase 3 | -0.47975 | 0.00394 |
| **ARF4** | ADP-ribosylation factor 4 | -0.50764 | 0.003943 |
| **AKIRIN2** | Akirin-2 | -0.63013 | 0.003944 |
| **VIRMA** | Protein virilizer homolog | -0.55072 | 0.003995 |
| **USP9X** | Probable ubiquitin carboxyl-terminal hydrolase FAF-X | -0.54246 | 0.003995 |
| **ARHGEF2** | Rho guanine nucleotide exchange factor 2 | -0.31827 | 0.003996 |
| **BRAT1** | BRCA1-associated ATM activator 1 | -0.66735 | 0.004019 |
| **ABCB10** | ATP-binding cassette sub-family B member 10, mitochondrial | -0.53239 | 0.004062 |
| **LRWD1** | Leucine-rich repeat and WD repeat-containing protein 1 | -0.83298 | 0.004092 |
| **CCAR2** | Cell cycle and apoptosis regulator protein 2 | -0.35545 | 0.00416 |
| **PPP6C** | Serine/threonine-protein phosphatase 6 catalytic subunit | -0.53218 | 0.004209 |
| **PGD** | 6-phosphogluconate dehydrogenase, decarboxylating | -0.3216 | 0.004224 |
| **ATP2C1** | Calcium-transporting ATPase type 2C member 1 | -0.573 | 0.004225 |
| **SH3GLB1** | Endophilin-B1 | -0.37262 | 0.004228 |

| | | | |
|---|---|---|---|
| **NCAPD2** | Condensin complex subunit 1 | -0.61216 | 0.004243 |
| **ACTL6A** | Actin-like protein 6A | -0.4378 | 0.004259 |
| **TENT4B** | Terminal nucleotidyltransferase 4B | -0.57588 | 0.004264 |
| **DHX37** | Probable ATP-dependent RNA helicase DHX37 | -0.47218 | 0.004268 |
| **HECTD1** | E3 ubiquitin-protein ligase HECTD1 | -0.56465 | 0.004301 |
| **UTP20** | Small subunit processome component 20 homolog | -0.6051 | 0.004319 |
| **EIF3E** | Eukaryotic translation initiation factor 3 subunit E | -0.4976 | 0.00435 |
| **POTEE** | POTE ankyrin domain family member E | -0.89521 | 0.00436 |
| **GTPBP1** | GTP-binding protein 1 | -0.34334 | 0.004369 |
| **ANAPC4** | Anaphase-promoting complex subunit 4 | -0.55659 | 0.004419 |
| **CDC27** | Cell division cycle protein 27 homolog | -0.54726 | 0.004443 |
| **FLOT1** | Flotillin-1 | -0.35181 | 0.004448 |
| **GTF2H4** | General transcription factor IIH subunit 4 | -0.36883 | 0.004451 |
| **XXYLT1** | Xyloside xylosyltransferase 1 | -0.67374 | 0.004471 |
| **UCK2** | Uridine-cytidine kinase 2 | -0.46321 | 0.004472 |
| **MET** | Hepatocyte growth factor receptor | -0.32872 | 0.004473 |
| **PYGB** | Glycogen phosphorylase, brain form | -0.31829 | 0.004476 |
| **BTAF1** | TATA-binding protein-associated factor 172 | -0.36206 | 0.004505 |
| **RPL10** | 60S ribosomal protein L10 | -0.30236 | 0.004516 |
| **TIMM50** | Mitochondrial import inner membrane translocase subunit TIM50 | -0.38793 | 0.004554 |
| **GNAI3** | Guanine nucleotide-binding protein G(i) subunit alpha-3 | -0.65407 | 0.004565 |
| **SEPTIN6** | Septin-6 | -0.61969 | 0.004594 |
| **H1-10** | Histone H1.10 | -1.06063 | 0.004598 |
| **STAT3** | Signal transducer and activator of transcription 3 | -0.35441 | 0.004606 |
| **VPS16** | Vacuolar protein sorting-associated protein 16 homolog | -0.54265 | 0.004613 |
| **MTA1** | Metastasis-associated protein MTA1 | -0.57323 | 0.004728 |
| **CRYBG1** | Beta/gamma crystallin domain-containing protein 1 | -0.34516 | 0.004742 |
| **ATP1A1** | Sodium/potassium-transporting ATPase subunit alpha-1 | -0.50365 | 0.004744 |
| **RAB5B** | Ras-related protein Rab-5B | -0.87776 | 0.004744 |

| | | | |
|---|---|---|---|
| **COPS6** | COP9 signalosome complex subunit 6 | -0.4532 | 0.00477 |
| **DDX51** | ATP-dependent RNA helicase DDX51 | -0.31865 | 0.004814 |
| **CHD8** | Chromodomain-helicase-DNA-binding protein 8 | -0.34664 | 0.004877 |
| **KYAT3** | Kynurenine--oxoglutarate transaminase 3 | -0.64078 | 0.00492 |
| **SLC6A6** | Sodium- and chloride-dependent taurine transporter | -0.81424 | 0.004925 |
| **GBA1** | Lysosomal acid glucosylceramidase | -0.1106 | 0.004963 |
| **TLE4** | Transducin-like enhancer protein 4 | -0.55496 | 0.005016 |
| **MCM2** | DNA replication licensing factor MCM2 | -0.36279 | 0.005041 |
| **EIF2B3** | Translation initiation factor eIF-2B subunit gamma | -0.22778 | 0.005072 |
| **SPNS1** | Protein spinster homolog 1 | -1.30355 | 0.005095 |
| **GTF3C4** | General transcription factor 3C polypeptide 4 | -0.34031 | 0.005123 |
| **ATM** | Serine-protein kinase ATM | -0.57026 | 0.005159 |
| **RARS1** | Arginine--tRNA ligase, cytoplasmic | -0.41479 | 0.005167 |
| **ATP6V0A1** | V-type proton ATPase 116 kDa subunit a 1 | -0.41514 | 0.005197 |
| **ORC3** | Origin recognition complex subunit 3 | -0.52881 | 0.005199 |
| **CPNE3** | Copine-3 | -0.36378 | 0.005225 |
| **SENP3** | Sentrin-specific protease 3 | -0.41565 | 0.005227 |
| **DDX3X** | ATP-dependent RNA helicase DDX3X | -0.27175 | 0.005235 |
| **FARSB** | Phenylalanine--tRNA ligase beta subunit | -0.61824 | 0.005249 |
| **TRIO** | Triple functional domain protein | -0.47834 | 0.00527 |
| **CDYL** | Chromodomain Y-like protein | -0.81879 | 0.005347 |
| **KANSL3** | KAT8 regulatory NSL complex subunit 3 | -0.87255 | 0.005349 |
| **SPAG9** | C-Jun-amino-terminal kinase-interacting protein 4 | -0.15887 | 0.005351 |
| **KDM1A** | Lysine-specific histone demethylase 1A | -0.52739 | 0.005378 |
| **ACADVL** | Very long-chain specific acyl-CoA dehydrogenase, mitochondrial | -0.45185 | 0.00538 |
| **NT5C2** | Cytosolic purine 5'-nucleotidase | -0.42717 | 0.005389 |
| **TSR1** | Pre-rRNA-processing protein TSR1 homolog | -0.30191 | 0.005401 |
| **THOC6** | THO complex subunit 6 homolog | -0.68393 | 0.005412 |

| | | | |
|---|---|---|---|
| **L2HGDH** | L-2-hydroxyglutarate dehydrogenase, mitochondrial | -0.45149 | 0.005417 |
| **GNL3L** | Guanine nucleotide-binding protein-like 3-like protein | -0.56984 | 0.005516 |
| **SLX4** | Structure-specific endonuclease subunit SLX4 | -0.25603 | 0.00553 |
| **MAP2K6** | Dual specificity mitogen-activated protein kinase kinase 6 | -0.60093 | 0.005533 |
| **ZC3HC1** | Zinc finger C3HC-type protein 1 | -0.51057 | 0.005541 |
| **PDCD6IP** | Programmed cell death 6-interacting protein | -0.42975 | 0.005543 |
| **DYNLL1** | Dynein light chain 1, cytoplasmic | -0.91485 | 0.005547 |
| **THOC2** | THO complex subunit 2 | -0.49652 | 0.005551 |
| **NIPBL** | Nipped-B-like protein | -0.31788 | 0.005567 |
| **NOL10** | Nucleolar protein 10 | -0.40895 | 0.005594 |
| **TSEN34** | tRNA-splicing endonuclease subunit Sen34 | -0.63513 | 0.0056 |
| **MSH6** | DNA mismatch repair protein Msh6 | -0.47538 | 0.0056 |
| **NIBAN2** | Protein Niban 2 | -0.51582 | 0.005625 |
| **HTRA2** | Serine protease HTRA2, mitochondrial | -0.36444 | 0.005625 |
| **PMS2CL** | Protein PMS2CL | -1.03672 | 0.005698 |
| **RBM12** | RNA-binding protein 12 | -0.35568 | 0.005734 |
| **AARS2** | Alanine--tRNA ligase, mitochondrial | -0.59212 | 0.005751 |
| **RANBP2** | E3 SUMO-protein ligase RanBP2 | -0.39538 | 0.005754 |
| **DPP3** | Dipeptidyl peptidase 3 | -0.42291 | 0.005795 |
| **WDR74** | WD repeat-containing protein 74 | -0.4848 | 0.005797 |
| **GPCPD1** | Glycerophosphocholine phosphodiesterase GPCPD1 | -0.601 | 0.005809 |
| **DDX52** | Probable ATP-dependent RNA helicase DDX52 | -0.46526 | 0.00581 |
| **AHCYL2** | Adenosylhomocysteinase 3 | -0.75422 | 0.005833 |
| **SCD** | Stearoyl-CoA desaturase | -0.76289 | 0.005844 |
| **XPO7** | Exportin-7 | -0.70178 | 0.005889 |
| **PSMD12** | 26S proteasome non-ATPase regulatory subunit 12 | -0.45935 | 0.005909 |
| **VWA8** | von Willebrand factor A domain-containing protein 8 | -0.87659 | 0.005922 |
| **DYNC1LI2** | Cytoplasmic dynein 1 light intermediate chain 2 | -0.12381 | 0.005926 |
| **ARL6IP5** | PRA1 family protein 3 | -0.13296 | 0.005948 |
| **TMA7** | Translation machinery-associated protein 7 | -1.05062 | 0.005954 |
| **PIAS2** | E3 SUMO-protein ligase PIAS2 | -0.90907 | 0.00597 |

| | | | |
|---|---|---|---|
| **ZCCHC4** | rRNA N6-adenosine-methyltransferase ZCCHC4 | -0.66183 | 0.005991 |
| **EFL1** | Elongation factor-like GTPase 1 | -0.60857 | 0.005991 |
| **ABCE1** | ATP-binding cassette sub-family E member 1 | -0.43902 | 0.006001 |
| **FLII** | Protein flightless-1 homolog | -0.56343 | 0.006005 |
| **RABGGTA** | Geranylgeranyl transferase type-2 subunit alpha | -0.57979 | 0.00602 |
| **CDK9** | Cyclin-dependent kinase 9 | -0.60707 | 0.006062 |
| **LONP1** | Lon protease homolog, mitochondrial | -0.31415 | 0.006074 |
| **CHD1L** | Chromodomain-helicase-DNA-binding protein 1-like | -0.50664 | 0.006091 |
| **ACACA** | Acetyl-CoA carboxylase 1 | -0.37494 | 0.006093 |
| **CCT2** | T-complex protein 1 subunit beta | -0.30877 | 0.0061 |
| **FH** | Fumarate hydratase, mitochondrial | -0.39741 | 0.006125 |
| **SYT11** | Synaptotagmin-11 | -0.52676 | 0.006129 |
| **COG1** | Conserved oligomeric Golgi complex subunit 1 | -0.48005 | 0.006132 |
| **R3HCC1** | R3H and coiled-coil domain-containing protein 1 | -0.57303 | 0.006134 |
| **ANKRD17** | Ankyrin repeat domain-containing protein 17 | -0.44549 | 0.006138 |
| **MPG** | DNA-3-methyladenine glycosylase | -0.56059 | 0.006187 |
| **BAZ2A** | Bromodomain adjacent to zinc finger domain protein 2A | -0.75546 | 0.006223 |
| **LDAH** | Lipid droplet-associated hydrolase | -1.15113 | 0.006283 |
| **MAN2A1** | Alpha-mannosidase 2 | -0.56566 | 0.006315 |
| **RPS27** | 40S ribosomal protein S27 | -0.29062 | 0.006329 |
| **TBC1D13** | TBC1 domain family member 13 | -0.43815 | 0.006346 |
| **PLEC** | Plectin | -0.33479 | 0.00636 |
| **SMAP** | Small acidic protein | -1.36555 | 0.006371 |
| **SCFD1** | Sec1 family domain-containing protein 1 | -0.47467 | 0.006392 |
| **DHX8** | ATP-dependent RNA helicase DHX8 | -0.4666 | 0.006484 |
| **ATG7** | Ubiquitin-like modifier-activating enzyme ATG7 | -0.59852 | 0.006536 |
| **KDSR** | 3-ketodihydrosphingosine reductase | -0.31928 | 0.00656 |
| **FASN** | Fatty acid synthase | -0.39102 | 0.006566 |
| **NCDN** | Neurochondrin | -0.47049 | 0.006679 |
| **NAT10** | RNA cytidine acetyltransferase | -0.35278 | 0.006701 |
| **RETREG1** | Reticulophagy regulator 1 | -0.54889 | 0.006777 |
| **PEPD** | Xaa-Pro dipeptidase | -0.53956 | 0.006813 |

| | | | |
|---|---|---|---|
| **EHMT1** | Histone-lysine N-methyltransferase EHMT1 | -0.32354 | 0.006825 |
| **HADHA** | Trifunctional enzyme subunit alpha, mitochondrial | -0.46522 | 0.006854 |
| **ALDH3A2** | Aldehyde dehydrogenase family 3 member A2 | -0.56305 | 0.006937 |
| **NNT** | NAD(P) transhydrogenase, mitochondrial | -0.67712 | 0.006959 |
| **MYO1F** | Unconventional myosin-If | -0.71368 | 0.006979 |
| **RNF111** | E3 ubiquitin-protein ligase Arkadia | -0.98348 | 0.007007 |
| **PAPOLA** | Poly(A) polymerase alpha | -0.33216 | 0.007034 |
| **URB2** | Unhealthy ribosome biogenesis protein 2 homolog | -0.8041 | 0.007045 |
| **SMG8** | Nonsense-mediated mRNA decay factor SMG8 | -0.47894 | 0.007081 |
| **FRYL** | Protein furry homolog-like | -0.61454 | 0.007083 |
| **PDS5A** | Sister chromatid cohesion protein PDS5 homolog A | -0.35414 | 0.007087 |
| **EIF2B2** | Translation initiation factor eIF-2B subunit beta | -0.45979 | 0.007098 |
| **HDGFL3** | Hepatoma-derived growth factor-related protein 3 | -0.75218 | 0.007154 |
| **TFRC** | Transferrin receptor protein 1 | -0.25779 | 0.007189 |
| **FAM120A** | Constitutive coactivator of PPAR-gamma-like protein 1 | -0.43595 | 0.007192 |
| **XPO4** | Exportin-4 | -0.43588 | 0.007214 |
| **USP47** | Ubiquitin carboxyl-terminal hydrolase 47 | -0.27995 | 0.007227 |
| **EXOC5** | Exocyst complex component 5 | -0.32817 | 0.007228 |
| **HIP1** | Huntingtin-interacting protein 1 | -0.26304 | 0.007231 |
| **EFTUD2** | 116 kDa U5 small nuclear ribonucleoprotein component | -0.21016 | 0.007236 |
| **POP1** | Ribonucleases P/MRP protein subunit POP1 | -0.27214 | 0.007276 |
| **CPSF2** | Cleavage and polyadenylation specificity factor subunit 2 | -0.35156 | 0.007276 |
| **THUMPD3** | tRNA (guanine(6)-N2)-methyltransferase THUMP3 | -0.79363 | 0.007283 |
| **TSC1** | Hamartin | -0.46424 | 0.007293 |
| **TUBGCP3** | Gamma-tubulin complex component 3 | -0.49594 | 0.007312 |
| **MCM6** | DNA replication licensing factor MCM6 | -0.47954 | 0.007315 |
| **EIF3C** | Eukaryotic translation initiation factor 3 subunit C | -0.2387 | 0.007325 |
| **MYH9** | Myosin-9 | -0.17106 | 0.007368 |

| | | | |
|---|---|---|---|
| **GOSR1** | Golgi SNAP receptor complex member 1 | -0.38968 | 0.007369 |
| **UBR4** | E3 ubiquitin-protein ligase UBR4 | -0.50044 | 0.007385 |
| **PSMB2** | Proteasome subunit beta type-2 | -0.48943 | 0.007404 |
| **NLN** | Neurolysin, mitochondrial | -0.47163 | 0.007494 |
| **EIF2B1** | Translation initiation factor eIF-2B subunit alpha | -0.38479 | 0.007535 |
| **AKAP1** | A-kinase anchor protein 1, mitochondrial | -0.31758 | 0.007542 |
| **L3HYPDH** | Trans-3-hydroxy-L-proline dehydratase | -0.5983 | 0.007574 |
| **PUF60** | Poly(U)-binding-splicing factor PUF60 | -0.43451 | 0.007585 |
| **KDM5A** | Lysine-specific demethylase 5A | -0.36936 | 0.007632 |
| **TYMS** | Thymidylate synthase | -0.48801 | 0.007635 |
| **INTS1** | Integrator complex subunit 1 | -0.75243 | 0.007655 |
| **TFB2M** | Dimethyladenosine transferase 2, mitochondrial | -0.49414 | 0.007676 |
| **TBL3** | Transducin beta-like protein 3 | -0.38503 | 0.007688 |
| **SEC24B** | Protein transport protein Sec24B | -0.71742 | 0.007709 |
| **PTPA** | Serine/threonine-protein phosphatase 2A activator | -0.45728 | 0.00775 |
| **ARMC8** | Armadillo repeat-containing protein 8 | -0.28111 | 0.007769 |
| **SPTLC2** | Serine palmitoyltransferase 2 | -0.51992 | 0.007821 |
| **PTK2** | Focal adhesion kinase 1 | -0.29274 | 0.007826 |
| **CKAP5** | Cytoskeleton-associated protein 5 | -0.32245 | 0.00785 |
| **USP48** | Ubiquitin carboxyl-terminal hydrolase 48 | -0.72829 | 0.007854 |
| **FAM172A** | Cotranscriptional regulator FAM172A | -0.57974 | 0.007884 |
| **EMC1** | ER membrane protein complex subunit 1 | -0.35178 | 0.007903 |
| **COG4** | Conserved oligomeric Golgi complex subunit 4 | -0.46161 | 0.007939 |
| **BPTF** | Nucleosome-remodeling factor subunit BPTF | -0.58401 | 0.007939 |
| **SRPK2** | SRSF protein kinase 2 | -0.37058 | 0.007954 |
| **RPS6KA3** | Ribosomal protein S6 kinase alpha-3 | -0.53031 | 0.007973 |
| **SMCR8** | Guanine nucleotide exchange protein SMCR8 | -0.49231 | 0.007982 |
| **RP2** | Protein XRP2 | -0.84907 | 0.007998 |
| **CLK3** | Dual specificity protein kinase CLK3 | -0.59431 | 0.008005 |
| **ACTR6** | Actin-related protein 6 | -0.54242 | 0.008007 |

| | | | |
|---|---|---|---|
| **MTPAP** | Poly(A) RNA polymerase, mitochondrial | -0.37004 | 0.008023 |
| **PPP4R1** | Serine/threonine-protein phosphatase 4 regulatory subunit 1 | -0.69727 | 0.008026 |
| **POTEF** | POTE ankyrin domain family member F | -0.73633 | 0.008056 |
| **PPP1CC** | Serine/threonine-protein phosphatase PP1-gamma catalytic subunit | -0.53472 | 0.00815 |
| **AP1G2** | AP-1 complex subunit gamma-like 2 | -0.31433 | 0.008184 |
| **AGPS** | Alkyldihydroxyacetonephosphate synthase, peroxisomal | -0.39474 | 0.008189 |
| **MTM1** | Myotubularin | -0.79615 | 0.008201 |
| **TUBG1** | Tubulin gamma-1 chain | -0.24794 | 0.008203 |
| **PPP3CA** | Protein phosphatase 3 catalytic subunit alpha | -0.11299 | 0.008301 |
| **CAND1** | Cullin-associated NEDD8-dissociated protein 1 | -0.44964 | 0.008335 |
| **RO60** | RNA-binding protein RO60 | -0.36522 | 0.008375 |
| **THNSL1** | Threonine synthase-like 1 | -0.52464 | 0.008441 |
| **SERPINB1** | Leukocyte elastase inhibitor | -0.37011 | 0.00854 |
| **PHIP** | PH-interacting protein | -0.32071 | 0.008553 |
| **COG7** | Conserved oligomeric Golgi complex subunit 7 | -0.46121 | 0.008555 |
| **HEATR5B** | HEAT repeat-containing protein 5B | -0.51627 | 0.008565 |
| **DNAJC13** | DnaJ homolog subfamily C member 13 | -0.24679 | 0.008571 |
| **CPNE1** | Copine-1 | -0.31289 | 0.008582 |
| **PKN2** | Serine/threonine-protein kinase N2 | -0.37914 | 0.008586 |
| **CASP8** | Caspase-8 | -0.4366 | 0.008588 |
| **ZNF592** | Zinc finger protein 592 | -0.31325 | 0.008591 |
| **ACD** | Adrenocortical dysplasia protein homolog | -0.34429 | 0.008648 |
| **PRR12** | Proline-rich protein 12 | -0.45017 | 0.008701 |
| **SSR1** | Translocon-associated protein subunit alpha | -0.89439 | 0.008721 |
| **ARFRP1** | ADP-ribosylation factor-related protein 1 | -0.50908 | 0.008747 |
| **MSH3** | DNA mismatch repair protein Msh3 | -0.27388 | 0.008796 |
| **RANBP6** | Ran-binding protein 6 | -0.65926 | 0.008829 |
| **SMARCA5** | SWI/SNF-related matrix-associated actin-dependent regulator of chromatin subfamily A member 5 | -0.61627 | 0.008863 |

| | | | |
|---|---|---|---|
| **MED13** | Mediator of RNA polymerase II transcription subunit 13 | -0.77139 | 0.008888 |
| **INTS4** | Integrator complex subunit 4 | -0.64956 | 0.008949 |
| **TFCP2** | Alpha-globin transcription factor CP2 | -0.60043 | 0.009041 |
| **PDCD11** | Protein RRP5 homolog | -0.38836 | 0.009063 |
| **UBA1** | Ubiquitin-like modifier-activating enzyme 1 | -0.39265 | 0.009153 |
| **TFDP1** | Transcription factor Dp-1 | -0.23077 | 0.009211 |
| **RDH11** | Retinol dehydrogenase 11 | -0.63851 | 0.009297 |
| **KYNU** | Kynureninase | -0.36251 | 0.009324 |
| **CDK1** | Cyclin-dependent kinase 1 | -0.20591 | 0.009357 |
| **MICAL3** | [F-actin]-monooxygenase MICAL3 | -0.22021 | 0.009388 |
| **APOB** | Apolipoprotein B-100 | -0.75227 | 0.009409 |
| **HACL1** | 2-hydroxyacyl-CoA lyase 1 | -0.37931 | 0.009427 |
| **VARS1** | Valine--tRNA ligase | -0.51974 | 0.009473 |
| **MGA** | MAX gene-associated protein | -0.50687 | 0.009478 |
| **EXOSC10** | Exosome component 10 | -0.42958 | 0.009479 |
| **KPNA6** | Importin subunit alpha-7 | -0.48993 | 0.009484 |
| **SRPRA** | Signal recognition particle receptor subunit alpha | -0.36278 | 0.00955 |
| **TMEM245** | Transmembrane protein 245 | -0.9294 | 0.009604 |
| **ZC3H3** | Zinc finger CCCH domain-containing protein 3 | -0.63509 | 0.009607 |
| **NARS1** | Asparagine--tRNA ligase, cytoplasmic | -0.32387 | 0.009615 |
| **CDC16** | Cell division cycle protein 16 homolog | -0.69502 | 0.009648 |
| **GCN1** | eIF-2-alpha kinase activator GCN1 | -0.44274 | 0.009649 |
| **AARSD1** | Alanyl-tRNA editing protein Aarsd1 | -0.21328 | 0.009678 |
| **SCYL2** | SCY1-like protein 2 | -0.45163 | 0.009707 |
| **DIAPH1** | Protein diaphanous homolog 1 | -0.29154 | 0.009742 |
| **UPF2** | Regulator of nonsense transcripts 2 | -0.62093 | 0.009772 |
| **ROCK2** | Rho-associated protein kinase 2 | -0.29278 | 0.009853 |
| **PSMB1** | Proteasome subunit beta type-1 | -0.37514 | 0.009874 |
| **EMC4** | ER membrane protein complex subunit 4 | -0.57283 | 0.00991 |
| **CNOT10** | CCR4-NOT transcription complex subunit 10 | -0.49587 | 0.009912 |
| **MTR** | Methionine synthase | -0.31589 | 0.009939 |
| **PPP1R3D** | Protein phosphatase 1 regulatory subunit 3D | -0.31936 | 0.00995 |

**Supplemental Table 7. Pathway Analysis for proteins upregulated by $JNK_i$ treatment in A375 melanoma cells.** See Table 3 for details.

**Biological Process (Gene Ontology)**

| GO-term | description | count in network | strength | false discovery rate |
|---|---|---|---|---|
| GO:0006695 | Cholesterol biosynthetic process | 6 of 38 | 1.73 | 4.98e-05 |
| GO:0008299 | Isoprenoid biosynthetic process | 4 of 29 | 1.67 | 0.0034 |
| GO:0006084 | acetyl-CoA metabolic process | 4 of 29 | 1.67 | 0.0034 |
| GO:0071379 | Cellular response to prostaglandin stimulus | 3 of 22 | 1.67 | 0.0389 |
| GO:0008203 | Cholesterol metabolic process | 8 of 119 | 1.36 | 4.98e-05 |
| GO:0006720 | Isoprenoid metabolic process | 5 of 119 | 1.15 | 0.0275 |
| GO:0008202 | Steroid metabolic process | 9 of 258 | 1.07 | 0.00015 |
| GO:0006066 | Alcohol metabolic process | 10 of 324 | 1.02 | 9.49e-05 |
| GO:0044283 | Small molecule biosynthetic process | 9 of 446 | 0.84 | 0.0074 |
| GO:0008610 | Lipid biosynthetic process | 10 of 577 | 0.77 | 0.0075 |
| GO:0006629 | Lipid metabolic process | 14 of 1210 | 0.59 | 0.0086 |
| GO:0044281 | Small molecule metabolic process | 15 of 1645 | 0.49 | 0.0436 |
| GO:0071310 | Cellular response to organic substance | 17 of 2019 | 0.46 | 0.0389 |
| GO:0070887 | Cellular response to chemical stimulus | 20 of 2609 | 0.42 | 0.0275 |
| GO:0010033 | Response to organic substance | 20 of 2692 | 0.4 | 0.0389 |

**Cellular Component (Gene Ontology)**

| GO-term | description | count in network | strength | false discovery rate |
|---|---|---|---|---|
| GO:0098802 | Plasma membrane signaling receptor complex | 5 of 194 | 0.94 | 0.0300 |
| GO:0005769 | Early endosome | 7 of 411 | 0.76 | 0.0280 |
| GO:0043235 | Receptor complex | 7 of 418 | 0.75 | 0.0280 |
| GO:0030141 | Secretory granule | 10 of 873 | 0.59 | 0.0280 |
| GO:0009986 | Cell surface | 10 of 894 | 0.58 | 0.0292 |
| GO:0005768 | Endosome | 11 of 1030 | 0.56 | 0.0280 |
| GO:0070062 | Extracellular exosome | 22 of 2096 | 0.55 | 0.00010 |
| GO:0005615 | Extracellular space | 25 of 3247 | 0.42 | 0.00058 |
| GO:0031982 | Vesicle | 30 of 3957 | 0.41 | 0.00010 |
| GO:0031410 | Cytoplasmic vesicle | 19 of 2482 | 0.41 | 0.0117 |
| GO:0012505 | Endomembrane system | 30 of 4721 | 0.33 | 0.0014 |
| GO:0005576 | Extracellular region | 26 of 4175 | 0.33 | 0.0112 |
| GO:0031090 | Organelle membrane | 22 of 3673 | 0.31 | 0.0457 |
| GO:0031224 | Intrinsic component of membrane | 31 of 5828 | 0.26 | 0.0216 |
| GO:0016021 | Integral component of membrane | 30 of 5670 | 0.25 | 0.0280 |
| GO:0016020 | Membrane | 42 of 9523 | 0.18 | 0.0274 |
| GO:0005737 | Cytoplasm | 50 of 12056 | 0.15 | 0.0074 |

**Local network cluster (STRING)**

| cluster | description | count in network | strength | false discovery rate |
|---|---|---|---|---|
| CL:9567 | Cholesterol biosynthesis pathway | 3 of 7 | 2.16 | 0.0031 |
| CL:9563 | Cholesterol biosynthesis | 5 of 22 | 1.89 | 6.43e-05 |
| CL:9386 | Steroid hormone biosynthesis, and Oxidation by cytochrome... | 6 of 129 | 1.2 | 0.0031 |

**KEGG Pathways**

| pathway | description | count in network | strength | false discovery rate |
|---|---|---|---|---|
| hsa00100 | Steroid biosynthesis | 3 of 20 | 1.71 | 0.0137 |
| hsa00900 | Terpenoid backbone biosynthesis | 3 of 22 | 1.67 | 0.0137 |
| hsa01100 | Metabolic pathways | 14 of 1435 | 0.52 | 0.0137 |

| | Reactome Pathways | | | |
|---|---|---|---|---|
| pathway | description | count in network | strength | false discovery rate |
| HSA-191273 | Cholesterol biosynthesis | 6 of 27 | 1.88 | 1.17e-06 |
| HSA-2426168 | Activation of gene expression by SREBF (SREBP) | 6 of 42 | 1.69 | 6.25e-06 |
| HSA-975634 | Retinoid metabolism and transport | 4 of 44 | 1.49 | 0.0045 |
| HSA-8957322 | Metabolism of steroids | 8 of 153 | 1.25 | 1.69e-05 |
| HSA-1430728 | Metabolism | 19 of 2092 | 0.49 | 0.0023 |

| | Subcellular localization (COMPARTMENTS) | | | |
|---|---|---|---|---|
| compartment | description | count in network | strength | false discovery rate |
| GOCC:0005769 | Early endosome | 7 of 253 | 0.97 | 0.0049 |
| GOCC:0098827 | Endoplasmic reticulum subcompartment | 8 of 502 | 0.73 | 0.0204 |
| GOCC:0005789 | Endoplasmic reticulum membrane | 8 of 538 | 0.7 | 0.0282 |
| GOCC:0005764 | Lysosome | 8 of 566 | 0.68 | 0.0321 |
| GOCC:0031984 | Organelle subcompartment | 10 of 739 | 0.66 | 0.0133 |
| GOCC:0098797 | Plasma membrane protein complex | 9 of 682 | 0.65 | 0.0281 |
| GOCC:0005794 | Golgi apparatus | 11 of 931 | 0.6 | 0.0152 |
| GOCC:0005615 | Extracellular space | 11 of 1027 | 0.56 | 0.0282 |
| GOCC:0031410 | Cytoplasmic vesicle | 16 of 1738 | 0.5 | 0.0100 |
| GOCC:0016021 | Integral component of membrane | 13 of 1507 | 0.47 | 0.0411 |
| GOCC:0031982 | Vesicle | 18 of 2125 | 0.46 | 0.0097 |
| GOCC:0012505 | Endomembrane system | 26 of 3156 | 0.45 | 0.00027 |
| GOCC:0005737 | Cytoplasm | 45 of 8195 | 0.27 | 6.30e-05 |
| GOCC:0016020 | Membrane | 30 of 5715 | 0.25 | 0.0321 |
| GOCC:0043227 | Membrane-bounded organelle | 45 of 9083 | 0.23 | 0.00079 |
| GOCC:0043226 | Organelle | 47 of 10113 | 0.2 | 0.0016 |
| GOCC:0005622 | Intracellular | 48 of 11512 | 0.15 | 0.0150 |
| GOCC:0110165 | Cellular anatomical entity | 54 of 14060 | 0.12 | 0.0111 |

| | Annotated Keywords (UniProt) | | | |
|---|---|---|---|---|
| keyword | description | count in network | strength | false discovery rate |
| KW-0427 | LDL | 2 of 8 | 1.93 | 0.0252 |
| KW-0152 | Cholesterol biosynthesis | 4 of 20 | 1.83 | 6.53e-05 |
| KW-0752 | Steroid biosynthesis | 6 of 43 | 1.68 | 4.18e-06 |
| KW-0153 | Cholesterol metabolism | 6 of 65 | 1.5 | 1.02e-05 |
| KW-0753 | Steroid metabolism | 7 of 98 | 1.38 | 7.91e-06 |
| KW-0444 | Lipid biosynthesis | 8 of 164 | 1.22 | 8.31e-06 |
| KW-0443 | Lipid metabolism | 11 of 485 | 0.89 | 1.96e-05 |
| KW-0732 | Signal | 20 of 3277 | 0.32 | 0.0464 |
| KW-0597 | Phosphoprotein | 38 of 8122 | 0.2 | 0.0125 |

**Supplemental Table 8. Pathway Analysis for proteins downregulated by $JNK_i$ treatment in A375 melanoma cells.** See Table 3 for details.

**Biological Process (Gene Ontology)**

| GO-term | description | count in network | strength | false discovery rate |
|---|---|---|---|---|
| GO:0051231 | Spindle elongation | 3 of 13 | 1.94 | 0.0098 |
| GO:0090307 | Mitotic spindle assembly | 5 of 51 | 1.57 | 0.00094 |
| GO:0007052 | Mitotic spindle organization | 6 of 97 | 1.37 | 0.00091 |
| GO:1902850 | Microtubule cytoskeleton organization involved in mitosis | 7 of 129 | 1.31 | 0.00034 |
| GO:0031398 | Positive regulation of protein ubiquitination | 5 of 121 | 1.19 | 0.0174 |
| GO:0007059 | Chromosome segregation | 8 of 286 | 1.03 | 0.0018 |
| GO:0051301 | Cell division | 11 of 527 | 0.9 | 0.00048 |
| GO:1903047 | Mitotic cell cycle process | 10 of 537 | 0.85 | 0.0024 |
| GO:0051276 | Chromosome organization | 17 of 968 | 0.82 | 4.20e-06 |
| GO:0000278 | Mitotic cell cycle | 11 of 631 | 0.82 | 0.0016 |
| GO:0006325 | Chromatin organization | 9 of 612 | 0.75 | 0.0257 |
| GO:0007049 | Cell cycle | 17 of 1246 | 0.71 | 9.09e-05 |
| GO:0022402 | Cell cycle process | 11 of 835 | 0.7 | 0.0099 |
| GO:0031401 | Positive regulation of protein modification process | 11 of 1018 | 0.61 | 0.0418 |
| GO:0051247 | Positive regulation of protein metabolic process | 15 of 1512 | 0.57 | 0.0081 |
| GO:0051173 | Positive regulation of nitrogen compound metabolic process | 22 of 3166 | 0.42 | 0.0087 |
| GO:0006996 | Organelle organization | 23 of 3470 | 0.4 | 0.0090 |
| GO:0009893 | Positive regulation of metabolic process | 24 of 3847 | 0.37 | 0.0119 |
| GO:0010604 | Positive regulation of macromolecule metabolic process | 22 of 3533 | 0.37 | 0.0292 |
| GO:0060255 | Regulation of macromolecule metabolic process | 31 of 6249 | 0.27 | 0.0257 |
| GO:0019222 | Regulation of metabolic process | 32 of 6784 | 0.25 | 0.0418 |

**Cellular Component (Gene Ontology)**

| GO-term | description | count in network | strength | false discovery rate |
|---|---|---|---|---|
| GO:0098687 | Chromosomal region | 7 of 365 | 0.86 | 0.0095 |
| GO:0005819 | Spindle | 7 of 425 | 0.8 | 0.0205 |
| GO:0005694 | Chromosome | 19 of 1850 | 0.59 | 0.00011 |
| GO:0005654 | Nucleoplasm | 29 of 4169 | 0.42 | 0.00011 |
| GO:0031981 | Nuclear lumen | 30 of 4526 | 0.4 | 0.00011 |
| GO:0043232 | Intracellular non-membrane-bounded organelle | 32 of 5191 | 0.37 | 0.00011 |
| GO:0005829 | Cytosol | 32 of 5438 | 0.35 | 0.00012 |
| GO:0070013 | Intracellular organelle lumen | 31 of 5660 | 0.32 | 0.00090 |
| GO:0005634 | Nucleus | 38 of 7672 | 0.27 | 0.00019 |
| GO:0043226 | Organelle | 48 of 14017 | 0.11 | 0.0277 |
| GO:0005622 | Intracellular anatomical structure | 50 of 14891 | 0.1 | 0.0130 |

**Local network cluster (STRING)**

| cluster | description | count in network | strength | false discovery rate |
|---|---|---|---|---|
| CL:6608 | Mixed, incl. Regulation of mitotic sister chromatid segregati… | 6 of 51 | 1.65 | 7.00e-06 |
| CL:6614 | Mixed, incl. Spindle elongation, and Polo-like kinase mediate… | 4 of 35 | 1.64 | 0.0019 |
| CL:6617 | Mixed, incl. Spindle elongation, and Polo-like kinase mediate… | 3 of 30 | 1.58 | 0.0406 |
| CL:6597 | Mixed, incl. Amplification of signal from the kinetochores, a… | 10 of 121 | 1.5 | 6.65e-09 |
| CL:6599 | Amplification of signal from the kinetochores, and Mitotic si… | 9 of 113 | 1.48 | 4.36e-08 |
| CL:6601 | Mixed, incl. Centromere, and Condensin complex | 8 of 104 | 1.46 | 5.80e-07 |
| CL:6604 | Mixed, incl. Amplification of signal from the kinetochores, a… | 7 of 93 | 1.46 | 7.00e-06 |

### Reactome Pathways

| pathway | description | count in network | strength | false discovery rate |
|---|---|---|---|---|
| HSA-3214841 | PKMTs methylate histone lysines | 4 of 47 | 1.51 | 0.0070 |
| HSA-141444 | Amplification of signal from unattached kinetochores via a ... | 5 of 94 | 1.3 | 0.0070 |
| HSA-9648025 | EML4 and NUDC in mitotic spindle formation | 5 of 116 | 1.21 | 0.0070 |
| HSA-2500257 | Resolution of Sister Chromatid Cohesion | 5 of 125 | 1.18 | 0.0070 |
| HSA-5663220 | RHO GTPases Activate Formins | 5 of 139 | 1.13 | 0.0084 |
| HSA-68877 | Mitotic Prometaphase | 6 of 201 | 1.05 | 0.0070 |
| HSA-2467813 | Separation of Sister Chromatids | 5 of 189 | 1.0 | 0.0197 |
| HSA-68882 | Mitotic Anaphase | 6 of 232 | 0.99 | 0.0084 |
| HSA-69620 | Cell Cycle Checkpoints | 6 of 272 | 0.92 | 0.0124 |
| HSA-195258 | RHO GTPase Effectors | 6 of 292 | 0.89 | 0.0171 |
| HSA-1640170 | Cell Cycle | 12 of 658 | 0.84 | 0.00029 |
| HSA-68886 | M Phase | 7 of 382 | 0.84 | 0.0112 |
| HSA-69278 | Cell Cycle, Mitotic | 9 of 526 | 0.81 | 0.0070 |
| HSA-194315 | Signaling by Rho GTPases | 9 of 672 | 0.71 | 0.0112 |

### Subcellular localization (COMPARTMENTS)

| compartment | description | count in network | strength | false discovery rate |
|---|---|---|---|---|
| GOCC:0099738 | Cell cortex region | 3 of 37 | 1.49 | 0.0332 |
| GOCC:0005694 | Chromosome | 11 of 951 | 0.64 | 0.0109 |
| GOCC:0031981 | Nuclear lumen | 15 of 1850 | 0.49 | 0.0157 |
| GOCC:0005634 | Nucleus | 32 of 4787 | 0.4 | 1.58e-05 |
| GOCC:0043232 | Intracellular non-membrane-bounded organelle | 22 of 3309 | 0.4 | 0.0066 |
| GOCC:0005829 | Cytosol | 19 of 3054 | 0.37 | 0.0340 |
| GOCC:0043231 | Intracellular membrane-bounded organelle | 36 of 8162 | 0.22 | 0.0139 |
| GOCC:0005622 | Intracellular | 49 of 11512 | 0.21 | 1.53e-05 |
| GOCC:0043229 | Intracellular organelle | 41 of 9609 | 0.21 | 0.0066 |
| GOCC:0043226 | Organelle | 42 of 10113 | 0.2 | 0.0066 |
| GOCC:0043227 | Membrane-bounded organelle | 38 of 9083 | 0.2 | 0.0172 |
| GOCC:0110165 | Cellular anatomical entity | 48 of 14060 | 0.11 | 0.0377 |

### Annotated Keywords (UniProt)

| keyword | description | count in network | strength | false discovery rate |
|---|---|---|---|---|
| KW-0995 | Kinetochore | 4 of 100 | 1.18 | 0.0072 |
| KW-0656 | Proto-oncogene | 8 of 225 | 1.13 | 2.15e-05 |
| KW-0498 | Mitosis | 8 of 275 | 1.04 | 7.85e-05 |
| KW-0137 | Centromere | 4 of 138 | 1.04 | 0.0210 |
| KW-0132 | Cell division | 10 of 384 | 0.99 | 1.12e-05 |
| KW-0160 | Chromosomal rearrangement | 8 of 312 | 0.99 | 0.00013 |
| KW-0158 | Chromosome | 9 of 433 | 0.9 | 0.00014 |
| KW-0156 | Chromatin regulator | 6 of 292 | 0.89 | 0.0066 |
| KW-0131 | Cell cycle | 11 of 651 | 0.81 | 9.16e-05 |
| KW-1017 | Isopeptide bond | 20 of 1717 | 0.64 | 1.24e-06 |
| KW-0832 | Ubl conjugation | 24 of 2399 | 0.58 | 5.46e-07 |
| KW-0175 | Coiled coil | 18 of 2166 | 0.5 | 0.00033 |
| KW-0863 | Zinc-finger | 14 of 1749 | 0.48 | 0.0066 |
| KW-0007 | Acetylation | 24 of 3362 | 0.43 | 9.56e-05 |
| KW-0862 | Zinc | 15 of 2347 | 0.38 | 0.0310 |
| KW-0597 | Phosphoprotein | 46 of 8122 | 0.33 | 1.32e-09 |
| KW-0539 | Nucleus | 30 of 5278 | 0.33 | 0.00017 |
| KW-0963 | Cytoplasm | 25 of 5095 | 0.27 | 0.0202 |

**Supplemental Table 9. Proteins which were upregulated by $JNK_i$ treatment in A375 melanoma cells.**

| Upregulated proteins after 24 hours of treatment with 5 µM $JNK_i$ | | | |
|---|---|---|---|
| **Gene name for protein** | **Description** | **Fold change (log2 form)** | **t-test P value** |
| **GLG1** | Golgi apparatus protein 1 | 0.606109 | 5.29E-05 |
| **HMGCS1** | Hydroxymethylglutaryl-CoA synthase, cytoplasmic | 2.182911 | 0.00014 |
| **TIMP3** | Metalloproteinase inhibitor 3 | 0.51054 | 0.000167 |
| **PDCD4** | Programmed cell death protein 4 | 0.653706 | 0.000302 |
| **IL13RA1** | Interleukin-13 receptor subunit alpha-1 | 1.002629 | 0.000412 |
| **PRSS23** | Serine protease 23 | 0.883382 | 0.000427 |
| **HLA-C** | HLA class I histocompatibility antigen, C alpha chain | 0.598935 | 0.000541 |
| **ZRANB3** | DNA annealing helicase and endonuclease ZRANB3 | 1.158247 | 0.00065 |
| **CPD** | Carboxypeptidase D | 0.320362 | 0.000717 |
| **FDFT1** | Squalene synthase | 1.670443 | 0.000852 |
| **RETREG1** | Reticulophagy regulator 1 | 0.830256 | 0.001284 |
| **TMEM59** | Transmembrane protein 59 | 0.833334 | 0.001351 |
| **IRS2** | Insulin receptor substrate 2 | 0.508446 | 0.001539 |
| **EPHA3** | Ephrin type-A receptor 3 | 0.211954 | 0.00157 |
| **LGALS3BP** | Galectin-3-binding protein | 0.440837 | 0.001626 |
| **PDHX** | Pyruvate dehydrogenase protein X component, mitochondrial | 0.105182 | 0.00171 |
| **HSPA2** | Heat shock-related 70 kDa protein 2 | 0.256839 | 0.001728 |
| **MSANTD2** | Myb/SANT-like DNA-binding domain-containing protein 2 | 0.87182 | 0.002199 |
| **LDLR** | Low-density lipoprotein receptor | 0.806592 | 0.002271 |
| **AKR1C2** | Aldo-keto reductase family 1 member C2 | 1.660153 | 0.002305 |

| | | | |
|---|---|---|---|
| **MRPL42** | 39S ribosomal protein L42, mitochondrial | 0.543379 | 0.002353 |
| **CKAP4** | Cytoskeleton-associated protein 4 | 0.314722 | 0.002377 |
| **APOB** | Apolipoprotein B-100 | 1.023775 | 0.002475 |
| **SSB** | Lupus La protein | 0.218659 | 0.002556 |
| **CLIC4** | Chloride intracellular channel protein 4 | 0.455844 | 0.002602 |
| **MGMT** | Methylated-DNA--protein-cysteine methyltransferase | 0.164415 | 0.002995 |
| **LSS** | Lanosterol synthase | 0.666856 | 0.003079 |
| **ACSS2** | Acetyl-coenzyme A synthetase, cytoplasmic | 1.316489 | 0.003104 |
| **IDI1** | Isopentenyl-diphosphate Delta-isomerase 1 | 1.016591 | 0.003173 |
| **SDC2** | Syndecan-2 | 1.480612 | 0.003649 |
| **B4GALT5** | Beta-1,4-galactosyltransferase 5 | 0.766622 | 0.003711 |
| **PRKAR1A** | cAMP-dependent protein kinase type I-alpha regulatory subunit | 0.580742 | 0.003998 |
| **ZBTB7A** | Zinc finger and BTB domain-containing protein 7A | 0.500812 | 0.00411 |
| **SCD** | Stearoyl-CoA desaturase | 0.633223 | 0.004135 |
| **ALDOC** | Fructose-bisphosphate aldolase C | 0.483884 | 0.004379 |
| **TOM1** | Target of Myb1 membrane trafficking protein | 0.470492 | 0.004628 |
| **ST8SIA4** | CMP-N-acetylneuraminate-poly-alpha-2,8-sialyltransferase | 1.234322 | 0.004958 |
| **SLAIN1** | SLAIN motif-containing protein 1 | 0.562937 | 0.005309 |
| **SLC31A1** | High affinity copper uptake protein 1 | 1.128583 | 0.005311 |
| **CD109** | CD109 antigen | 0.518803 | 0.005515 |
| **SFRP1** | Secreted frizzled-related protein 1 | 1.217803 | 0.005799 |
| **HSD17B7** | 3-keto-steroid reductase/17-beta-hydroxysteroid dehydrogenase 7 | 0.940309 | 0.006314 |

| | | | |
|---|---|---|---|
| **MVP** | Major vault protein | 0.580006 | 0.006446 |
| **IL6ST** | Interleukin-6 receptor subunit beta | 0.429649 | 0.006458 |
| **ITFG1** | T-cell immunomodulatory protein | 0.822078 | 0.006483 |
| **FAM210A** | Protein FAM210A | 0.320547 | 0.006578 |
| **EMILIN1** | EMILIN-1 | 0.848747 | 0.006728 |
| **TMX2** | Thioredoxin-related transmembrane protein 2 | 0.57766 | 0.006755 |
| **ITM2B** | Integral membrane protein 2B | 0.644165 | 0.006903 |
| **MVD** | Diphosphomevalonate decarboxylase | 1.010148 | 0.007304 |
| **ITGA1** | Integrin alpha-1 | 0.509668 | 0.007502 |
| **BLVRB** | Flavin reductase (NADPH) | 0.253048 | 0.008019 |
| **KYNU** | Kynureninase | 0.545873 | 0.008096 |
| **RAB11FIP5** | Rab11 family-interacting protein 5 | 0.313177 | 0.008208 |
| **STX8** | Syntaxin-8 | 0.34906 | 0.008552 |
| **AGRN** | Agrin | 0.479388 | 0.009351 |
| **EPS8** | Epidermal growth factor receptor kinase substrate 8 | 0.194475 | 0.00945 |
| **PDE3A** | cGMP-inhibited 3',5'-cyclic phosphodiesterase 3A | 1.033972 | 0.00985 |

**Supplemental Table 10. Proteins which were downregulated by $JNK_i$ treatment in A375 melanoma cells.**

| Downregulated proteins after 24 hours of treatment with 5 μM $JNK_i$ | | | |
|---|---|---|---|
| **Gene name for protein** | **Description** | **Fold change (log2 form)** | **t-test P value** |
| **CEP192** | Centrosomal protein of 192 kDa | -0.50994 | 4.15E-05 |
| **RNF40** | E3 ubiquitin-protein ligase BRE1B | -0.234 | 9.38E-05 |
| **KIF4A** | Chromosome-associated kinesin KIF4A | -0.36544 | 0.000387 |
| **DIDO1** | Death-inducer obliterator 1 | -0.22264 | 0.000897 |
| **CDR2** | Cerebellar degeneration-related protein 2 | -0.80042 | 0.000923 |
| **SUZ12** | Polycomb protein SUZ12 | -0.30105 | 0.001242 |
| **AFAP1L2** | Actin filament-associated protein 1-like 2 | -0.39762 | 0.001444 |
| **AHCTF1** | Protein ELYS | -0.31439 | 0.001888 |
| **LMNB1** | Lamin-B1 | -0.24942 | 0.001892 |
| **CTIF** | CBP80/20-dependent translation initiation factor | -0.26944 | 0.001951 |
| **CIT** | Citron Rho-interacting kinase | -0.28346 | 0.002341 |
| **ERCC6L** | DNA excision repair protein ERCC-6-like | -0.30645 | 0.00245 |
| **NSD2** | Histone-lysine N-methyltransferase NSD2 | -0.47433 | 0.002478 |
| **SP2** | Transcription factor Sp2 | -0.61668 | 0.002826 |
| **SOGA1** | Protein SOGA1 | -0.34956 | 0.002929 |
| **HELLS** | Lymphoid-specific helicase | -0.58403 | 0.002953 |
| **SLX4** | Structure-specific endonuclease subunit SLX4 | -0.61141 | 0.003466 |
| **DTL** | Denticleless protein homolog | -0.63157 | 0.003502 |
| **XIAP** | E3 ubiquitin-protein ligase XIAP | -0.19705 | 0.003818 |

| | | | |
|---|---|---|---|
| **XXYLT1** | Xyloside xylosyltransferase 1 | -0.46844 | 0.003935 |
| **TES** | Testin | -0.36648 | 0.00399 |
| **AXL** | Tyrosine-protein kinase receptor UFO | -0.5824 | 0.004006 |
| **MIDEAS** | Mitotic deacetylase-associated SANT domain protein | -0.29417 | 0.00417 |
| **AKAP13** | A-kinase anchor protein 13 | -0.14573 | 0.004304 |
| **CAMK2G** | Calcium/calmodulin-dependent protein kinase type II subunit gamma | -0.45292 | 0.004837 |
| **KIF11** | Kinesin-like protein KIF11 | -0.62196 | 0.005297 |
| **MACF1** | Microtubule-actin cross-linking factor 1, isoforms 6/7 | -0.42033 | 0.005297 |
| **CDCA2** | Cell division cycle-associated protein 2 | -0.69304 | 0.00548 |
| **PLIN2** | Perilipin-2 | -0.40746 | 0.005558 |
| **POLA1** | DNA polymerase alpha catalytic subunit | -0.38562 | 0.005786 |
| **SKA1** | Spindle and kinetochore-associated protein 1 | -0.85337 | 0.005951 |
| **ELL** | RNA polymerase II elongation factor ELL | -0.38677 | 0.006197 |
| **RACGAP1** | Rac GTPase-activating protein 1 | -0.48839 | 0.006345 |
| **TERF2** | Telomeric repeat-binding factor 2 | -0.17021 | 0.006512 |
| **GINS2** | DNA replication complex GINS protein PSF2 | -0.35967 | 0.006799 |
| **BCL3** | B-cell lymphoma 3 protein | -0.54918 | 0.006799 |
| **BDP1** | Transcription factor TFIIIB component B" homolog | -0.36833 | 0.007198 |
| **MALT1** | Mucosa-associated lymphoid tissue lymphoma translocation protein 1 | -0.37698 | 0.007252 |
| **HMGN3** | High mobility group nucleosome-binding domain-containing protein 3 | -0.68105 | 0.007393 |

| | | | |
|---|---|---|---|
| **DUS3L** | tRNA-dihydrouridine(47) synthase [NAD(P)(+)]-like | -0.27031 | 0.007458 |
| **ARG2** | Arginase-2, mitochondrial | -0.4996 | 0.007524 |
| **MPRIP** | Myosin phosphatase Rho-interacting protein | -0.14341 | 0.007802 |
| **SAPCD2** | Suppressor APC domain-containing protein 2 | -0.433 | 0.008035 |
| **INAVA** | Innate immunity activator protein | -0.57108 | 0.008266 |
| **NSD3** | Histone-lysine N-methyltransferase NSD3 | -0.45137 | 0.008267 |
| **NIBAN1** | Protein Niban 1 | -0.41561 | 0.008491 |
| **QRICH1** | Transcriptional regulator QRICH1 | -0.39836 | 0.008678 |
| **MMADHC** | Cobalamin trafficking protein CblD | -0.31197 | 0.008915 |
| **CDC20** | Cell division cycle protein 20 homolog | -0.74862 | 0.009388 |
| **CENPH** | Centromere protein H | -0.50451 | 0.009424 |
| **LENG8** | Leukocyte receptor cluster member 8 | -0.28083 | 0.009764 |
| **ADRM1** | Proteasomal ubiquitin receptor ADRM1 | -0.27613 | 0.009845 |
| **KMT2A** | Histone-lysine N-methyltransferase 2A | -0.27395 | 0.00988 |

**Supplemental Table 11. Pathway Analysis for proteins upregulated by JNK$_i$ co-treatment (with GEM) compared to single GEM treatment in A375 melanoma cells.** See Table 3 for details.

| Biological Process (Gene Ontology) | | | | |
|---|---|---|---|---|
| GO-term | description | count in network | strength | false discovery rate |
| GO:0034382 | Chylomicron remnant clearance | 3 of 5 | 2.05 | 0.0057 |
| GO:0032375 | Negative regulation of cholesterol transport | 4 of 14 | 1.73 | 0.0021 |
| GO:0033700 | Phospholipid efflux | 3 of 12 | 1.67 | 0.0274 |
| GO:0006695 | Cholesterol biosynthetic process | 7 of 38 | 1.54 | 2.69e-05 |
| GO:0032369 | Negative regulation of lipid transport | 5 of 30 | 1.5 | 0.0017 |
| GO:0034381 | Plasma lipoprotein particle clearance | 4 of 24 | 1.5 | 0.0078 |
| GO:0033344 | Cholesterol efflux | 4 of 27 | 1.44 | 0.0110 |
| GO:0034369 | Plasma lipoprotein particle remodeling | 4 of 29 | 1.41 | 0.0137 |
| GO:0090207 | Regulation of triglyceride metabolic process | 4 of 40 | 1.27 | 0.0358 |
| GO:0008203 | Cholesterol metabolic process | 10 of 119 | 1.2 | 2.59e-05 |
| GO:0042632 | Cholesterol homeostasis | 6 of 89 | 1.1 | 0.0070 |
| GO:0015918 | Sterol transport | 5 of 82 | 1.06 | 0.0401 |
| GO:0032368 | Regulation of lipid transport | 8 of 139 | 1.03 | 0.0017 |
| GO:1905952 | Regulation of lipid localization | 9 of 167 | 1.0 | 0.00071 |
| GO:1901617 | Organic hydroxy compound biosynthetic process | 8 of 172 | 0.94 | 0.0041 |
| GO:0055088 | Lipid homeostasis | 7 of 161 | 0.91 | 0.0150 |
| GO:0006066 | Alcohol metabolic process | 11 of 324 | 0.8 | 0.0020 |
| GO:0019216 | Regulation of lipid metabolic process | 9 of 346 | 0.69 | 0.0427 |
| GO:1901615 | Organic hydroxy compound metabolic process | 12 of 478 | 0.67 | 0.0070 |
| GO:0006629 | Lipid metabolic process | 20 of 1210 | 0.49 | 0.0046 |
| GO:0042127 | Regulation of cell population proliferation | 25 of 1669 | 0.45 | 0.0021 |
| GO:0051172 | Negative regulation of nitrogen compound metabolic process | 29 of 2403 | 0.35 | 0.0093 |
| GO:0031324 | Negative regulation of cellular metabolic process | 27 of 2265 | 0.35 | 0.0204 |
| GO:0009892 | Negative regulation of metabolic process | 35 of 2982 | 0.34 | 0.0026 |
| GO:0048523 | Negative regulation of cellular process | 52 of 4736 | 0.31 | 3.42e-05 |
| GO:0010605 | Negative regulation of macromolecule metabolic process | 30 of 2760 | 0.31 | 0.0358 |
| GO:0048519 | Negative regulation of biological process | 55 of 5313 | 0.29 | 6.69e-05 |
| GO:0010604 | Positive regulation of macromolecule metabolic process | 35 of 3533 | 0.27 | 0.0427 |
| GO:0009893 | Positive regulation of metabolic process | 37 of 3847 | 0.26 | 0.0447 |
| GO:0080090 | Regulation of primary metabolic process | 50 of 5899 | 0.2 | 0.0420 |

| Molecular Function (Gene Ontology) | | | | |
|---|---|---|---|---|
| GO-term | description | count in network | strength | false discovery rate |
| GO:0005515 | Protein binding | 62 of 7242 | 0.21 | 0.0153 |

| Cellular Component (Gene Ontology) | | | | |
|---|---|---|---|---|
| GO-term | description | count in network | strength | false discovery rate |
| GO:0034363 | Intermediate-density lipoprotein particle | 4 of 5 | 2.18 | 9.43e-05 |
| GO:0042627 | Chylomicron | 4 of 13 | 1.76 | 0.00043 |
| GO:0034362 | Low-density lipoprotein particle | 3 of 12 | 1.67 | 0.0076 |
| GO:0034361 | Very-low-density lipoprotein particle | 4 of 20 | 1.57 | 0.0014 |
| GO:0034364 | High-density lipoprotein particle | 4 of 29 | 1.41 | 0.0037 |
| GO:0030662 | Coated vesicle membrane | 6 of 193 | 0.77 | 0.0431 |
| GO:0005788 | Endoplasmic reticulum lumen | 9 of 312 | 0.73 | 0.0068 |
| GO:0005789 | Endoplasmic reticulum membrane | 17 of 1157 | 0.44 | 0.0116 |
| GO:0070062 | Extracellular exosome | 30 of 2096 | 0.43 | 0.00024 |
| GO:0005783 | Endoplasmic reticulum | 27 of 2021 | 0.4 | 0.0012 |
| GO:0031984 | Organelle subcompartment | 20 of 1479 | 0.4 | 0.0090 |
| GO:0005615 | Extracellular space | 35 of 3247 | 0.31 | 0.0029 |
| GO:0031982 | Vesicle | 40 of 3957 | 0.28 | 0.0027 |
| GO:0012505 | Endomembrane system | 42 of 4721 | 0.22 | 0.0150 |
| GO:0070013 | Intracellular organelle lumen | 47 of 5660 | 0.19 | 0.0254 |
| GO:0043227 | Membrane-bounded organelle | 95 of 13188 | 0.13 | 3.46e-05 |
| GO:0043231 | Intracellular membrane-bounded organelle | 87 of 12149 | 0.13 | 0.00049 |
| GO:0005737 | Cytoplasm | 83 of 12056 | 0.11 | 0.0081 |
| GO:0043229 | Intracellular organelle | 88 of 13231 | 0.1 | 0.0090 |
| GO:0005622 | Intracellular anatomical structure | 95 of 14891 | 0.08 | 0.0090 |

| | Local network cluster (STRING) | | | |
|---|---|---|---|---|
| *cluster* | *description* | *count in network* | *strength* | *false discovery rate* |
| CL:9567 | Cholesterol biosynthesis pathway | 4 of 7 | 2.03 | 0.00022 |
| CL:9565 | Steroid biosynthesis | 5 of 12 | 1.89 | 5.12e-05 |
| CL:9563 | Cholesterol biosynthesis | 6 of 22 | 1.71 | 3.01e-05 |

| | KEGG Pathways | | | |
|---|---|---|---|---|
| *pathway* | *description* | *count in network* | *strength* | *false discovery rate* |
| hsa00100 | Steroid biosynthesis | 5 of 20 | 1.67 | 6.24e-05 |
| hsa04979 | Cholesterol metabolism | 4 of 48 | 1.19 | 0.0279 |

| | Reactome Pathways | | | |
|---|---|---|---|---|
| *pathway* | *description* | *count in network* | *strength* | *false discovery rate* |
| HSA-8963901 | Chylomicron remodeling | 4 of 10 | 1.88 | 0.00033 |
| HSA-8963888 | Chylomicron assembly | 4 of 10 | 1.88 | 0.00033 |
| HSA-8964058 | HDL remodeling | 3 of 10 | 1.75 | 0.0076 |
| HSA-191273 | Cholesterol biosynthesis | 8 of 27 | 1.74 | 2.27e-08 |
| HSA-2426168 | Activation of gene expression by SREBF (SREBP) | 7 of 42 | 1.5 | 7.85e-06 |
| HSA-3000480 | Scavenging by Class A Receptors | 3 of 18 | 1.5 | 0.0256 |
| HSA-975634 | Retinoid metabolism and transport | 5 of 44 | 1.33 | 0.0017 |
| HSA-8957275 | Post-translational protein phosphorylation | 7 of 107 | 1.09 | 0.00081 |
| HSA-8957322 | Metabolism of steroids | 9 of 153 | 1.04 | 0.00012 |
| HSA-381426 | Regulation of Insulin-like Growth Factor (IGF) transport and ... | 7 of 124 | 1.02 | 0.0017 |
| HSA-1989781 | PPARA activates gene expression | 6 of 117 | 0.98 | 0.0084 |

| | Subcellular localization (COMPARTMENTS) | | | |
|---|---|---|---|---|
| *compartment* | *description* | *count in network* | *strength* | *false discovery rate* |
| GOCC:0034363 | Intermediate-density lipoprotein particle | 4 of 8 | 1.97 | 0.00038 |
| GOCC:0034360 | Chylomicron remnant | 2 of 4 | 1.97 | 0.0306 |
| GOCC:0062136 | Low-density lipoprotein receptor complex | 3 of 9 | 1.8 | 0.0054 |
| GOCC:0042627 | Chylomicron | 4 of 14 | 1.73 | 0.00083 |
| GOCC:0034361 | Very-low-density lipoprotein particle | 4 of 20 | 1.57 | 0.0021 |
| GOCC:0034362 | Low-density lipoprotein particle | 3 of 22 | 1.41 | 0.0251 |
| GOCC:0034364 | High-density lipoprotein particle | 4 of 30 | 1.4 | 0.0054 |
| GOCC:0005788 | Endoplasmic reticulum lumen | 6 of 172 | 0.82 | 0.0297 |
| GOCC:0070062 | Extracellular exosome | 10 of 428 | 0.64 | 0.0152 |
| GOCC:0098827 | Endoplasmic reticulum subcompartment | 11 of 502 | 0.61 | 0.0126 |
| GOCC:0065010 | Extracellular membrane-bounded organelle | 10 of 473 | 0.6 | 0.0219 |
| GOCC:0005789 | Endoplasmic reticulum membrane | 11 of 538 | 0.58 | 0.0183 |
| GOCC:0005783 | Endoplasmic reticulum | 21 of 1095 | 0.56 | 0.00038 |
| GOCC:0031984 | Organelle subcompartment | 13 of 739 | 0.52 | 0.0183 |
| GOCC:0005615 | Extracellular space | 16 of 1027 | 0.47 | 0.0152 |
| GOCC:0012505 | Endomembrane system | 34 of 3156 | 0.31 | 0.0053 |
| GOCC:0005634 | Nucleus | 42 of 4787 | 0.22 | 0.0240 |
| GOCC:0043231 | Intracellular membrane-bounded organelle | 69 of 8162 | 0.2 | 0.00038 |
| GOCC:0043227 | Membrane-bounded organelle | 75 of 9083 | 0.19 | 0.00033 |
| GOCC:0043226 | Organelle | 78 of 10113 | 0.16 | 0.00061 |
| GOCC:0005737 | Cytoplasm | 63 of 8195 | 0.16 | 0.0152 |
| GOCC:0043229 | Intracellular organelle | 73 of 9609 | 0.15 | 0.0033 |
| GOCC:0005622 | Intracellular | 84 of 11512 | 0.14 | 0.00083 |
| GOCC:0110165 | Cellular anatomical entity | 93 of 14060 | 0.09 | 0.0044 |

### Annotated Keywords (UniProt)

| keyword | description | count in network | strength | false discovery rate |
|---|---|---|---|---|
| KW-0380 | Hyperlipidemia | 2 of 4 | 1.97 | 0.0198 |
| KW-0162 | Chylomicron | 4 of 9 | 1.92 | 4.32e-05 |
| KW-0850 | VLDL | 3 of 10 | 1.75 | 0.0024 |
| KW-0152 | Cholesterol biosynthesis | 5 of 20 | 1.67 | 2.08e-05 |
| KW-0026 | Alzheimer disease | 3 of 18 | 1.5 | 0.0101 |
| KW-0752 | Steroid biosynthesis | 7 of 43 | 1.48 | 3.24e-06 |
| KW-0153 | Cholesterol metabolism | 8 of 65 | 1.36 | 3.24e-06 |
| KW-0444 | Lipid biosynthesis | 9 of 164 | 1.01 | 3.38e-05 |
| KW-0445 | Lipid transport | 5 of 110 | 0.93 | 0.0190 |
| KW-0443 | Lipid metabolism | 15 of 485 | 0.76 | 1.02e-05 |
| KW-0597 | Phosphoprotein | 68 of 8122 | 0.2 | 7.07e-05 |

**Supplemental Table 12. Pathway Analysis for proteins downregulated by JNK$_i$ co-treatment (with GEM) compared to single GEM treatment in A375 melanoma cells.** See Table 3 for details.

Biological Process (Gene Ontology)

| GO-term | description | count in network | strength | false discovery rate |
|---|---|---|---|---|
| GO:0051231 | Spindle elongation | 6 of 13 | 1.67 | 6.57e-06 |
| GO:0051256 | Mitotic spindle midzone assembly | 5 of 11 | 1.67 | 9.67e-05 |
| GO:0051987 | Positive regulation of attachment of spindle microtubules to... | 3 of 11 | 1.44 | 0.0383 |
| GO:0051988 | Regulation of attachment of spindle microtubules to kinetoc... | 4 of 19 | 1.33 | 0.0109 |
| GO:0090307 | Mitotic spindle assembly | 10 of 51 | 1.3 | 1.63e-07 |
| GO:0090266 | Regulation of mitotic cell cycle spindle assembly checkpoint | 4 of 21 | 1.29 | 0.0148 |
| GO:0051310 | Metaphase plate congression | 12 of 65 | 1.28 | 7.33e-09 |
| GO:0007052 | Mitotic spindle organization | 15 of 97 | 1.2 | 2.48e-10 |
| GO:0050000 | Chromosome localization | 13 of 84 | 1.2 | 7.33e-09 |
| GO:0051984 | Positive regulation of chromosome segregation | 4 of 26 | 1.2 | 0.0263 |
| GO:0007080 | Mitotic metaphase plate congression | 8 of 53 | 1.19 | 4.04e-05 |
| GO:1902850 | Microtubule cytoskeleton organization involved in mitosis | 18 of 129 | 1.15 | 5.72e-12 |
| GO:0000070 | Mitotic sister chromatid segregation | 16 of 122 | 1.13 | 3.24e-10 |
| GO:0032467 | Positive regulation of cytokinesis | 6 of 45 | 1.13 | 0.0020 |
| GO:0033047 | Regulation of mitotic sister chromatid segregation | 7 of 55 | 1.11 | 0.00054 |
| GO:0000819 | Sister chromatid segregation | 17 of 144 | 1.08 | 2.91e-10 |
| GO:0140014 | Mitotic nuclear division | 20 of 175 | 1.07 | 5.72e-12 |
| GO:1900087 | Positive regulation of G1/S transition of mitotic cell cycle | 5 of 44 | 1.06 | 0.0164 |
| GO:0051983 | Regulation of chromosome segregation | 14 of 128 | 1.05 | 5.77e-08 |
| GO:0010965 | Regulation of mitotic sister chromatid separation | 10 of 96 | 1.03 | 2.72e-05 |
| GO:1905818 | Regulation of chromosome separation | 11 of 109 | 1.01 | 8.50e-06 |
| GO:0030071 | Regulation of mitotic metaphase/anaphase transition | 9 of 91 | 1.0 | 0.00015 |
| GO:0033045 | Regulation of sister chromatid segregation | 10 of 105 | 0.99 | 5.55e-05 |
| GO:0032465 | Regulation of cytokinesis | 9 of 94 | 0.99 | 0.00019 |
| GO:0098813 | Nuclear chromosome segregation | 21 of 229 | 0.97 | 5.05e-11 |
| GO:0007059 | Chromosome segregation | 25 of 286 | 0.95 | 8.79e-13 |
| GO:1901992 | Positive regulation of mitotic cell cycle phase transition | 8 of 92 | 0.95 | 0.0013 |
| GO:0031638 | Zymogen activation | 5 of 59 | 0.94 | 0.0464 |
| GO:0000281 | Mitotic cytokinesis | 7 of 84 | 0.93 | 0.0054 |
| GO:0007098 | Centrosome cycle | 7 of 88 | 0.91 | 0.0069 |
| GO:0007088 | Regulation of mitotic nuclear division | 9 of 118 | 0.89 | 0.00095 |
| GO:1901879 | Regulation of protein depolymerization | 7 of 94 | 0.88 | 0.0098 |
| GO:0010389 | Regulation of G2/M transition of mitotic cell cycle | 7 of 99 | 0.86 | 0.0127 |
| GO:1903047 | Mitotic cell cycle process | 36 of 537 | 0.84 | 1.44e-15 |
| GO:0000280 | Nuclear division | 22 of 323 | 0.84 | 2.45e-09 |
| GO:0071103 | DNA conformation change | 7 of 104 | 0.84 | 0.0159 |
| GO:0031110 | Regulation of microtubule polymerization or depolymerizati... | 6 of 88 | 0.84 | 0.0388 |
| GO:0032508 | DNA duplex unwinding | 6 of 89 | 0.84 | 0.0405 |
| GO:0006260 | DNA replication | 13 of 203 | 0.82 | 6.03e-05 |
| GO:0071897 | DNA biosynthetic process | 6 of 92 | 0.82 | 0.0464 |
| GO:0051301 | Cell division | 33 of 527 | 0.81 | 1.79e-13 |
| GO:1901990 | Regulation of mitotic cell cycle phase transition | 21 of 332 | 0.81 | 2.22e-08 |
| GO:0033044 | Regulation of chromosome organization | 16 of 253 | 0.81 | 3.22e-06 |
| GO:0000278 | Mitotic cell cycle | 39 of 631 | 0.8 | 8.24e-16 |
| GO:0043244 | Regulation of protein-containing complex disassembly | 8 of 131 | 0.79 | 0.0108 |
| GO:0051302 | Regulation of cell division | 11 of 188 | 0.78 | 0.00095 |
| GO:0140694 | Non-membrane-bounded organelle assembly | 18 of 314 | 0.77 | 1.76e-06 |
| GO:0090068 | Positive regulation of cell cycle process | 14 of 251 | 0.76 | 9.67e-05 |
| GO:2000045 | Regulation of G1/S transition of mitotic cell cycle | 9 of 164 | 0.75 | 0.0085 |

| | | | | |
|---|---|---|---|---|
| GO:0000226 | Microtubule cytoskeleton organization | 29 of 542 | 0.74 | 2.55e-10 |
| GO:1901987 | Regulation of cell cycle phase transition | 23 of 431 | 0.74 | 5.57e-08 |
| GO:0010564 | Regulation of cell cycle process | 38 of 716 | 0.73 | 1.30e-13 |
| GO:0007346 | Regulation of mitotic cell cycle | 26 of 493 | 0.73 | 5.37e-09 |
| GO:0070507 | Regulation of microtubule cytoskeleton organization | 8 of 157 | 0.72 | 0.0285 |
| GO:0000075 | Cell cycle checkpoint signaling | 8 of 157 | 0.72 | 0.0285 |
| GO:0022402 | Cell cycle process | 41 of 835 | 0.7 | 1.04e-13 |
| GO:0007049 | Cell cycle | 56 of 1246 | 0.66 | 7.71e-18 |
| GO:0051656 | Establishment of organelle localization | 17 of 380 | 0.66 | 9.79e-05 |
| GO:0010948 | Negative regulation of cell cycle process | 12 of 272 | 0.65 | 0.0043 |
| GO:0006302 | Double-strand break repair | 9 of 204 | 0.65 | 0.0316 |
| GO:0051726 | Regulation of cell cycle | 46 of 1108 | 0.63 | 2.32e-13 |
| GO:0006281 | DNA repair | 21 of 497 | 0.63 | 1.18e-05 |
| GO:0032886 | Regulation of microtubule-based process | 11 of 261 | 0.63 | 0.0121 |
| GO:0007017 | Microtubule-based process | 31 of 803 | 0.6 | 6.10e-08 |
| GO:0045786 | Negative regulation of cell cycle | 14 of 359 | 0.6 | 0.0035 |
| GO:0097190 | Apoptotic signaling pathway | 12 of 318 | 0.59 | 0.0148 |
| GO:0006259 | DNA metabolic process | 29 of 785 | 0.58 | 6.11e-07 |
| GO:0006974 | Cellular response to DNA damage stimulus | 28 of 744 | 0.58 | 7.96e-07 |
| GO:0051640 | Organelle localization | 19 of 514 | 0.58 | 0.00028 |
| GO:0007010 | Cytoskeleton organization | 45 of 1229 | 0.57 | 2.24e-11 |
| GO:0051276 | Chromosome organization | 35 of 968 | 0.57 | 2.22e-08 |
| GO:0010639 | Negative regulation of organelle organization | 12 of 351 | 0.54 | 0.0307 |
| GO:0051493 | Regulation of cytoskeleton organization | 17 of 541 | 0.51 | 0.0059 |
| GO:0044089 | Positive regulation of cellular component biogenesis | 16 of 500 | 0.51 | 0.0080 |
| GO:0033043 | Regulation of organelle organization | 37 of 1190 | 0.5 | 2.67e-07 |
| GO:0070925 | Organelle assembly | 23 of 799 | 0.47 | 0.0012 |
| GO:0051129 | Negative regulation of cellular component organization | 20 of 691 | 0.47 | 0.0039 |
| GO:0030036 | Actin cytoskeleton organization | 15 of 547 | 0.45 | 0.0471 |
| GO:0051128 | Regulation of cellular component organization | 58 of 2365 | 0.4 | 1.69e-08 |
| GO:0018130 | Heterocycle biosynthetic process | 24 of 985 | 0.4 | 0.0080 |
| GO:0034654 | Nucleobase-containing compound biosynthetic process | 22 of 913 | 0.39 | 0.0166 |
| GO:0006996 | Organelle organization | 82 of 3470 | 0.38 | 1.78e-12 |
| GO:0090304 | Nucleic acid metabolic process | 51 of 2203 | 0.37 | 1.78e-06 |
| GO:0033554 | Cellular response to stress | 36 of 1572 | 0.37 | 0.00044 |
| GO:0019438 | Aromatic compound biosynthetic process | 23 of 992 | 0.37 | 0.0198 |
| GO:0044087 | Regulation of cellular component biogenesis | 22 of 971 | 0.36 | 0.0341 |
| GO:0044271 | Cellular nitrogen compound biosynthetic process | 33 of 1494 | 0.35 | 0.0023 |
| GO:0006139 | Nucleobase-containing compound metabolic process | 58 of 2722 | 0.34 | 2.16e-06 |
| GO:1901362 | Organic cyclic compound biosynthetic process | 24 of 1121 | 0.34 | 0.0389 |
| GO:0046483 | Heterocycle metabolic process | 61 of 2891 | 0.33 | 1.16e-06 |
| GO:0006725 | Cellular aromatic compound metabolic process | 60 of 2936 | 0.32 | 4.62e-06 |
| GO:1901360 | Organic cyclic compound metabolic process | 64 of 3181 | 0.31 | 2.27e-06 |
| GO:0016043 | Cellular component organization | 107 of 5436 | 0.3 | 1.07e-12 |
| GO:0034641 | Cellular nitrogen compound metabolic process | 68 of 3463 | 0.3 | 1.76e-06 |
| GO:0051649 | Establishment of localization in cell | 34 of 1756 | 0.3 | 0.0164 |
| GO:0071840 | Cellular component organization or biogenesis | 108 of 5639 | 0.29 | 4.09e-12 |
| GO:0044260 | Cellular macromolecule metabolic process | 48 of 2512 | 0.29 | 0.00095 |
| GO:0022607 | Cellular component assembly | 47 of 2467 | 0.29 | 0.0013 |
| GO:0044085 | Cellular component biogenesis | 49 of 2702 | 0.27 | 0.0027 |
| GO:0010467 | Gene expression | 38 of 2101 | 0.27 | 0.0228 |
| GO:0051641 | Cellular localization | 45 of 2677 | 0.23 | 0.0263 |
| GO:0051246 | Regulation of protein metabolic process | 44 of 2622 | 0.23 | 0.0316 |
| GO:0043170 | Macromolecule metabolic process | 90 of 5781 | 0.2 | 9.70e-05 |
| GO:0009893 | Positive regulation of metabolic process | 58 of 3847 | 0.19 | 0.0410 |
| GO:0010604 | Positive regulation of macromolecule metabolic process | 54 of 3533 | 0.19 | 0.0484 |
| GO:0006807 | Nitrogen compound metabolic process | 98 of 6643 | 0.18 | 0.00022 |
| GO:0048523 | Negative regulation of cellular process | 68 of 4736 | 0.17 | 0.0412 |
| GO:0048518 | Positive regulation of biological process | 88 of 6207 | 0.16 | 0.0058 |
| GO:0048522 | Positive regulation of cellular process | 79 of 5584 | 0.16 | 0.0185 |
| GO:0071704 | Organic substance metabolic process | 104 of 7522 | 0.15 | 0.0017 |
| GO:0044238 | Primary metabolic process | 99 of 7156 | 0.15 | 0.0034 |
| GO:0044237 | Cellular metabolic process | 90 of 6568 | 0.15 | 0.0148 |
| GO:0048519 | Negative regulation of biological process | 74 of 5313 | 0.15 | 0.0472 |
| GO:0008152 | Metabolic process | 108 of 7988 | 0.14 | 0.0025 |
| GO:0009987 | Cellular process | 173 of 14826 | 0.08 | 0.00011 |
| GO:0050789 | Regulation of biological process | 138 of 11655 | 0.08 | 0.0341 |

### Molecular Function (Gene Ontology)

| GO-term | description | count in network | strength | false discovery rate |
|---|---|---|---|---|
| GO:0003777 | Microtubule motor activity | 7 of 68 | 1.02 | 0.0027 |
| GO:0045296 | Cadherin binding | 21 of 334 | 0.81 | 6.76e-08 |
| GO:0008017 | Microtubule binding | 17 of 269 | 0.81 | 2.96e-06 |
| GO:0015631 | Tubulin binding | 19 of 380 | 0.71 | 1.07e-05 |
| GO:0050839 | Cell adhesion molecule binding | 22 of 560 | 0.6 | 3.98e-05 |
| GO:0016887 | ATP hydrolysis activity | 13 of 333 | 0.6 | 0.0071 |
| GO:0003779 | Actin binding | 16 of 448 | 0.56 | 0.0035 |
| GO:0008092 | Cytoskeletal protein binding | 35 of 1002 | 0.55 | 1.13e-07 |
| GO:0140657 | ATP-dependent activity | 17 of 543 | 0.5 | 0.0071 |
| GO:0019900 | Kinase binding | 21 of 785 | 0.44 | 0.0074 |
| GO:0019901 | Protein kinase binding | 19 of 702 | 0.44 | 0.0136 |
| GO:0003723 | RNA binding | 38 of 1672 | 0.37 | 0.00052 |
| GO:0032559 | Adenyl ribonucleotide binding | 35 of 1554 | 0.36 | 0.0015 |
| GO:0005524 | ATP binding | 33 of 1491 | 0.35 | 0.0033 |
| GO:0044877 | Protein-containing complex binding | 27 of 1261 | 0.34 | 0.0197 |
| GO:0019899 | Enzyme binding | 44 of 2084 | 0.33 | 0.00049 |
| GO:0032555 | Purine ribonucleotide binding | 38 of 1903 | 0.31 | 0.0048 |
| GO:0000166 | Nucleotide binding | 42 of 2168 | 0.3 | 0.0036 |
| GO:0035639 | Purine ribonucleoside triphosphate binding | 36 of 1834 | 0.3 | 0.0089 |
| GO:0043168 | Anion binding | 46 of 2404 | 0.29 | 0.0023 |
| GO:0042802 | Identical protein binding | 39 of 2144 | 0.27 | 0.0187 |
| GO:0036094 | Small molecule binding | 45 of 2507 | 0.26 | 0.0077 |
| GO:0097367 | Carbohydrate derivative binding | 41 of 2278 | 0.26 | 0.0157 |
| GO:0005515 | Protein binding | 123 of 7242 | 0.24 | 1.27e-10 |
| GO:0003676 | Nucleic acid binding | 65 of 4003 | 0.22 | 0.0033 |
| GO:0097159 | Organic cyclic compound binding | 92 of 6050 | 0.19 | 0.00043 |
| GO:1901363 | Heterocyclic compound binding | 90 of 5977 | 0.19 | 0.00074 |
| GO:0005488 | Binding | 172 of 12838 | 0.14 | 1.27e-10 |

### Cellular Component (Gene Ontology)

| GO-term | description | count in network | strength | false discovery rate |
|---|---|---|---|---|
| GO:0005971 | Ribonucleoside-diphosphate reductase complex | 2 of 2 | 2.01 | 0.0203 |
| GO:0097149 | Centralspindlin complex | 2 of 3 | 1.83 | 0.0324 |
| GO:0000940 | Outer kinetochore | 5 of 12 | 1.63 | 2.85e-05 |
| GO:0035371 | Microtubule plus-end | 3 of 18 | 1.23 | 0.0365 |
| GO:0005871 | Kinesin complex | 7 of 50 | 1.16 | 7.67e-05 |
| GO:0051233 | Spindle midzone | 5 of 36 | 1.15 | 0.0021 |
| GO:0001725 | Stress fiber | 7 of 65 | 1.04 | 0.00036 |
| GO:0000776 | Kinetochore | 14 of 165 | 0.94 | 2.21e-07 |
| GO:0000922 | Spindle pole | 14 of 169 | 0.93 | 2.82e-07 |
| GO:0005657 | Replication fork | 5 of 60 | 0.93 | 0.0159 |
| GO:0000775 | Chromosome, centromeric region | 19 of 252 | 0.89 | 3.15e-09 |
| GO:0072686 | Mitotic spindle | 13 of 180 | 0.87 | 3.79e-06 |
| GO:0030496 | Midbody | 14 of 201 | 0.85 | 1.89e-06 |
| GO:0005819 | Spindle | 29 of 425 | 0.84 | 3.15e-13 |
| GO:0045171 | Intercellular bridge | 6 of 91 | 0.83 | 0.0142 |
| GO:0000793 | Condensed chromosome | 18 of 275 | 0.82 | 7.50e-08 |
| GO:0000792 | Heterochromatin | 5 of 77 | 0.82 | 0.0408 |
| GO:0034451 | Centriolar satellite | 7 of 110 | 0.81 | 0.0058 |
| GO:0005875 | Microtubule associated complex | 10 of 168 | 0.78 | 0.00049 |
| GO:0090734 | Site of DNA damage | 6 of 102 | 0.78 | 0.0234 |
| GO:0098687 | Chromosomal region | 20 of 365 | 0.75 | 1.17e-07 |
| GO:0005912 | Adherens junction | 9 of 177 | 0.72 | 0.0036 |
| GO:0005813 | Centrosome | 26 of 609 | 0.64 | 7.50e-08 |
| GO:0005874 | Microtubule | 19 of 453 | 0.63 | 1.22e-05 |
| GO:0030027 | Lamellipodium | 8 of 203 | 0.6 | 0.0370 |
| GO:0005815 | Microtubule organizing center | 31 of 825 | 0.58 | 3.13e-08 |
| GO:0005911 | Cell-cell junction | 17 of 499 | 0.54 | 0.00058 |

| | | | | |
|---|---|---|---|---|
| GO:0005925 | Focal adhesion | 14 of 416 | 0.54 | 0.0035 |
| GO:0099513 | Polymeric cytoskeletal fiber | 25 of 757 | 0.53 | 1.15e-05 |
| GO:0015630 | Microtubule cytoskeleton | 44 of 1355 | 0.52 | 4.22e-10 |
| GO:0015629 | Actin cytoskeleton | 15 of 482 | 0.5 | 0.0043 |
| GO:0005856 | Cytoskeleton | 66 of 2369 | 0.45 | 4.98e-13 |
| GO:0099080 | Supramolecular complex | 38 of 1366 | 0.45 | 5.35e-07 |
| GO:0005730 | Nucleolus | 26 of 996 | 0.43 | 0.00036 |
| GO:0099512 | Supramolecular fiber | 26 of 1000 | 0.42 | 0.00037 |
| GO:0005694 | Chromosome | 44 of 1850 | 0.39 | 2.41e-06 |
| GO:0043232 | Intracellular non-membrane-bounded organelle | 116 of 5191 | 0.36 | 2.50e-19 |
| GO:0070161 | Anchoring junction | 29 of 1325 | 0.35 | 0.0021 |
| GO:0005829 | Cytosol | 114 of 5438 | 0.33 | 6.89e-17 |
| GO:0005654 | Nucleoplasm | 85 of 4169 | 0.32 | 3.17e-10 |
| GO:0031981 | Nuclear lumen | 89 of 4526 | 0.3 | 4.10e-10 |
| GO:0030054 | Cell junction | 40 of 2115 | 0.29 | 0.0018 |
| GO:0070013 | Intracellular organelle lumen | 96 of 5660 | 0.24 | 9.06e-08 |
| GO:0032991 | Protein-containing complex | 91 of 5506 | 0.23 | 1.08e-06 |
| GO:0005634 | Nucleus | 121 of 7672 | 0.21 | 4.21e-09 |
| GO:0005737 | Cytoplasm | 163 of 12056 | 0.14 | 4.10e-10 |
| GO:0043229 | Intracellular organelle | 173 of 13231 | 0.13 | 9.56e-11 |
| GO:0005622 | Intracellular anatomical structure | 187 of 14891 | 0.11 | 1.89e-13 |
| GO:0043226 | Organelle | 176 of 14017 | 0.11 | 1.68e-09 |
| GO:0043231 | Intracellular membrane-bounded organelle | 152 of 12149 | 0.11 | 2.03e-05 |
| GO:0043227 | Membrane-bounded organelle | 159 of 13188 | 0.09 | 7.76e-05 |
| GO:0110165 | Cellular anatomical entity | 190 of 18293 | 0.03 | 0.0160 |

**Local network cluster (STRING)**

| *cluster* | *description* | *count in network* | *strength* | *false discovery rate* |
|---|---|---|---|---|
| CL:6796 | Outer kinetochore | 3 of 5 | 1.79 | 0.0154 |
| CL:6657 | Chromosome passenger complex | 3 of 5 | 1.79 | 0.0154 |
| CL:6617 | Mixed, incl. Spindle elongation, and Polo-like kinase mediate... | 15 of 30 | 1.71 | 4.48e-17 |
| CL:6619 | Mixed, incl. Spindle elongation, and Outer kinetochore | 12 of 24 | 1.71 | 2.09e-13 |
| CL:6643 | Chromosome passenger complex, and Outer kinetochore | 6 of 12 | 1.71 | 4.96e-06 |
| CL:6620 | Mixed, incl. Spindle elongation, and Axon hillock | 6 of 12 | 1.71 | 4.96e-06 |
| CL:6614 | Mixed, incl. Spindle elongation, and Polo-like kinase mediate... | 16 of 35 | 1.67 | 1.00e-17 |
| CL:6644 | Mixed, incl. Activation of NIMA Kinases NEK9, NEK6, NEK7, ... | 3 of 7 | 1.64 | 0.0276 |
| CL:6622 | Mixed, incl. Mitotic centrosome separation, and Nuclear mic... | 3 of 7 | 1.64 | 0.0276 |
| CL:6610 | Mixed, incl. Spindle elongation, and Polo-like kinase mediate... | 17 of 41 | 1.63 | 2.67e-18 |
| CL:6608 | Mixed, incl. Regulation of mitotic sister chromatid segregati... | 19 of 51 | 1.58 | 2.57e-19 |
| CL:6606 | Mitotic sister chromatid segregation, and Mitotic spindle ch... | 20 of 61 | 1.52 | 2.57e-19 |
| CL:6601 | Mixed, incl. Centromere, and Condensin complex | 23 of 104 | 1.35 | 2.57e-19 |
| CL:6597 | Mixed, incl. Amplification of signal from the kinetochores, a... | 24 of 121 | 1.31 | 2.57e-19 |
| CL:6596 | Mixed, incl. Mitotic Spindle Checkpoint, and Mitotic sister ch... | 25 of 151 | 1.23 | 3.08e-19 |
| CL:25994 | Mixed, incl. Centriole replication, and Centriolar subdistal ap... | 5 of 49 | 1.02 | 0.0402 |
| CL:25993 | Anchoring of the basal body to the plasma membrane, and ... | 6 of 61 | 1.0 | 0.0154 |

**KEGG Pathways**

| *pathway* | *description* | *count in network* | *strength* | *false discovery rate* |
|---|---|---|---|---|
| hsa00480 | Glutathione metabolism | 5 of 52 | 0.99 | 0.0381 |
| hsa04115 | p53 signaling pathway | 6 of 72 | 0.93 | 0.0373 |

**Reactome Pathways**

| *pathway* | *description* | *count in network* | *strength* | *false discovery rate* |
|---|---|---|---|---|
| HSA-9755779 | SARS-CoV-2 targets host intracellular signalling and regulat... | 3 of 12 | 1.41 | 0.0182 |
| HSA-75035 | Chk1/Chk2(Cds1) mediated inactivation of Cyclin B:Cdk1 co... | 3 of 13 | 1.37 | 0.0214 |
| HSA-5358606 | Mismatch repair (MMR) directed by MSH2:MSH3 (MutSbeta) | 3 of 14 | 1.34 | 0.0253 |
| HSA-5358565 | Mismatch repair (MMR) directed by MSH2:MSH6 (MutSalph... | 3 of 14 | 1.34 | 0.0253 |
| HSA-174414 | Processive synthesis on the C-strand of the telomere | 4 of 19 | 1.33 | 0.0048 |
| HSA-9735871 | SARS-CoV-1 targets host intracellular signalling and regulat... | 3 of 15 | 1.31 | 0.0284 |

| | | | | |
|---|---|---|---|---|
| HSA-69183 | Processive synthesis on the lagging strand | 3 of 15 | 1.31 | 0.0284 |
| HSA-111447 | Activation of BAD and translocation to mitochondria | 3 of 15 | 1.31 | 0.0284 |
| HSA-156711 | Polo-like kinase mediated events | 3 of 16 | 1.28 | 0.0301 |
| HSA-6804114 | TP53 Regulates Transcription of Genes Involved in G2 Cell C... | 3 of 18 | 1.23 | 0.0364 |
| HSA-174417 | Telomere C-strand (Lagging Strand) Synthesis | 5 of 34 | 1.18 | 0.0032 |
| HSA-5651801 | PCNA-Dependent Long Patch Base Excision Repair | 3 of 21 | 1.16 | 0.0464 |
| HSA-983189 | Kinesins | 7 of 60 | 1.08 | 0.00055 |
| HSA-180786 | Extension of Telomeres | 6 of 51 | 1.08 | 0.0018 |
| HSA-9696264 | RND3 GTPase cycle | 5 of 42 | 1.08 | 0.0059 |
| HSA-9696273 | RND1 GTPase cycle | 4 of 42 | 0.99 | 0.0358 |
| HSA-9696270 | RND2 GTPase cycle | 4 of 43 | 0.98 | 0.0371 |
| HSA-69473 | G2/M DNA damage checkpoint | 7 of 77 | 0.97 | 0.0018 |
| HSA-4615885 | SUMOylation of DNA replication proteins | 4 of 45 | 0.96 | 0.0425 |
| HSA-141444 | Amplification of signal from unattached kinetochores via a ... | 8 of 94 | 0.94 | 0.00078 |
| HSA-8854518 | AURKA Activation by TPX2 | 6 of 72 | 0.93 | 0.0068 |
| HSA-73893 | DNA Damage Bypass | 4 of 48 | 0.93 | 0.0467 |
| HSA-69618 | Mitotic Spindle Checkpoint | 9 of 111 | 0.92 | 0.00046 |
| HSA-9013106 | RHOC GTPase cycle | 6 of 73 | 0.92 | 0.0072 |
| HSA-6791312 | TP53 Regulates Transcription of Cell Cycle Genes | 4 of 49 | 0.92 | 0.0495 |
| HSA-68877 | Mitotic Prometaphase | 16 of 201 | 0.91 | 2.25e-07 |
| HSA-2500257 | Resolution of Sister Chromatid Cohesion | 10 of 125 | 0.91 | 0.00018 |
| HSA-157579 | Telomere Maintenance | 7 of 93 | 0.89 | 0.0044 |
| HSA-5693607 | Processing of DNA double-strand break ends | 6 of 80 | 0.88 | 0.0109 |
| HSA-380259 | Loss of Nlp from mitotic centrosomes | 5 of 69 | 0.87 | 0.0301 |
| HSA-6811434 | COPI-dependent Golgi-to-ER retrograde traffic | 7 of 99 | 0.86 | 0.0054 |
| HSA-9013026 | RHOB GTPase cycle | 5 of 70 | 0.86 | 0.0302 |
| HSA-69620 | Cell Cycle Checkpoints | 19 of 272 | 0.85 | 5.20e-08 |
| HSA-9648025 | EML4 and NUDC in mitotic spindle formation | 8 of 116 | 0.85 | 0.0026 |
| HSA-2565942 | Regulation of PLK1 Activity at G2/M Transition | 6 of 87 | 0.85 | 0.0145 |
| HSA-69481 | G2/M Checkpoints | 10 of 149 | 0.84 | 0.00055 |
| HSA-453279 | Mitotic G1 phase and G1/S transition | 10 of 148 | 0.84 | 0.00055 |
| HSA-6781827 | Transcription-Coupled Nucleotide Excision Repair (TC-NER) | 5 of 78 | 0.82 | 0.0425 |
| HSA-1640170 | Cell Cycle | 42 of 658 | 0.81 | 2.35e-18 |
| HSA-69278 | Cell Cycle, Mitotic | 33 of 526 | 0.81 | 7.40e-14 |
| HSA-69275 | G2/M Transition | 12 of 195 | 0.8 | 0.00018 |
| HSA-69206 | G1/S Transition | 8 of 130 | 0.8 | 0.0047 |
| HSA-5693567 | HDR through Homologous Recombination (HRR) or Single S... | 7 of 113 | 0.8 | 0.0100 |
| HSA-380270 | Recruitment of mitotic centrosome proteins and complexes | 5 of 80 | 0.8 | 0.0438 |
| HSA-5693532 | DNA Double-Strand Break Repair | 9 of 149 | 0.79 | 0.0023 |
| HSA-2467813 | Separation of Sister Chromatids | 11 of 189 | 0.77 | 0.00057 |
| HSA-5663220 | RHO GTPases Activate Formins | 8 of 139 | 0.77 | 0.0061 |
| HSA-69239 | Synthesis of DNA | 7 of 120 | 0.77 | 0.0129 |
| HSA-68886 | M Phase | 21 of 382 | 0.75 | 2.25e-07 |
| HSA-5696398 | Nucleotide Excision Repair | 6 of 109 | 0.75 | 0.0323 |
| HSA-3108232 | SUMO E3 ligases SUMOylate target proteins | 9 of 166 | 0.74 | 0.0044 |
| HSA-68882 | Mitotic Anaphase | 12 of 232 | 0.72 | 0.00062 |
| HSA-983231 | Factors involved in megakaryocyte development and platele... | 8 of 155 | 0.72 | 0.0109 |
| HSA-9018519 | Estrogen-dependent gene expression | 6 of 119 | 0.71 | 0.0438 |
| HSA-69242 | S Phase | 8 of 162 | 0.7 | 0.0132 |
| HSA-2132295 | MHC class II antigen presentation | 6 of 122 | 0.7 | 0.0464 |
| HSA-73894 | DNA Repair | 15 of 310 | 0.69 | 0.00018 |
| HSA-8980692 | RHOA GTPase cycle | 7 of 148 | 0.68 | 0.0302 |
| HSA-8939211 | ESR-mediated signaling | 8 of 190 | 0.63 | 0.0288 |
| HSA-195258 | RHO GTPase Effectors | 12 of 292 | 0.62 | 0.0037 |
| HSA-3700989 | Transcriptional Regulation by TP53 | 13 of 361 | 0.57 | 0.0054 |
| HSA-194315 | Signaling by Rho GTPases | 24 of 672 | 0.56 | 2.88e-05 |
| HSA-9012999 | RHO GTPase cycle | 13 of 449 | 0.47 | 0.0264 |
| HSA-5663205 | Infectious disease | 20 of 917 | 0.35 | 0.0302 |
| HSA-162582 | Signal Transduction | 52 of 2540 | 0.32 | 4.80e-05 |
| HSA-74160 | Gene expression (Transcription) | 27 of 1476 | 0.27 | 0.0438 |
| HSA-1643685 | Disease | 30 of 1702 | 0.26 | 0.0426 |

**Subcellular localization (COMPARTMENTS)**

| compartment | description | count in network | strength | false discovery rate |
|---|---|---|---|---|
| GOCC:0097149 | Centralspindlin complex | 3 of 4 | 1.88 | 0.0018 |
| GOCC:0000940 | Condensed chromosome outer kinetochore | 5 of 13 | 1.59 | 4.84e-05 |
| GOCC:0035371 | Microtubule plus-end | 3 of 17 | 1.26 | 0.0342 |
| GOCC:0051233 | Spindle midzone | 5 of 31 | 1.22 | 0.0015 |
| GOCC:0005871 | Kinesin complex | 7 of 46 | 1.19 | 5.69e-05 |
| GOCC:0001725 | Stress fiber | 5 of 39 | 1.12 | 0.0036 |
| GOCC:0000922 | Spindle pole | 13 of 118 | 1.05 | 7.38e-08 |
| GOCC:0099738 | Cell cortex region | 4 of 37 | 1.04 | 0.0256 |
| GOCC:0072686 | Mitotic spindle | 13 of 133 | 1.0 | 2.50e-07 |
| GOCC:0000779 | Condensed chromosome, centromeric region | 11 of 112 | 1.0 | 3.54e-06 |
| GOCC:0000776 | Kinetochore | 11 of 115 | 0.99 | 3.98e-06 |
| GOCC:0005819 | Spindle | 27 of 289 | 0.98 | 2.22e-15 |
| GOCC:0000775 | Chromosome, centromeric region | 14 of 160 | 0.95 | 2.36e-07 |
| GOCC:0030496 | Midbody | 12 of 142 | 0.94 | 3.56e-06 |
| GOCC:0045171 | Intercellular bridge | 7 of 92 | 0.89 | 0.0029 |
| GOCC:0005657 | Replication fork | 5 of 67 | 0.88 | 0.0259 |
| GOCC:0034451 | Centriolar satellite | 7 of 105 | 0.83 | 0.0055 |
| GOCC:0070160 | Tight junction | 6 of 92 | 0.82 | 0.0165 |
| GOCC:0005875 | Microtubule associated complex | 9 of 150 | 0.79 | 0.0016 |
| GOCC:0000793 | Condensed chromosome | 12 of 205 | 0.78 | 9.88e-05 |
| GOCC:0043296 | Apical junction complex | 6 of 101 | 0.78 | 0.0246 |
| GOCC:0005813 | Centrosome | 22 of 470 | 0.68 | 3.35e-07 |
| GOCC:1902911 | Protein kinase complex | 7 of 160 | 0.65 | 0.0455 |
| GOCC:0005874 | Microtubule | 10 of 233 | 0.64 | 0.0063 |
| GOCC:0005815 | Microtubule organizing center | 25 of 607 | 0.62 | 3.33e-07 |
| GOCC:0005925 | Focal adhesion | 11 of 269 | 0.62 | 0.0048 |
| GOCC:0015630 | Microtubule cytoskeleton | 39 of 973 | 0.61 | 2.33e-11 |
| GOCC:0099513 | Polymeric cytoskeletal fiber | 15 of 382 | 0.6 | 0.00052 |
| GOCC:0005694 | Chromosome | 35 of 951 | 0.57 | 4.79e-09 |
| GOCC:0070161 | Anchoring junction | 20 of 553 | 0.57 | 5.93e-05 |
| GOCC:0005911 | Cell-cell junction | 11 of 299 | 0.57 | 0.0101 |
| GOCC:0005856 | Cytoskeleton | 56 of 1575 | 0.56 | 5.65e-15 |
| GOCC:0061695 | Transferase complex, transferring phosphorus-containing gr... | 11 of 322 | 0.54 | 0.0178 |
| GOCC:0005730 | Nucleolus | 20 of 606 | 0.53 | 0.00021 |
| GOCC:0000785 | Chromatin | 15 of 476 | 0.51 | 0.0048 |
| GOCC:0099080 | Supramolecular complex | 26 of 844 | 0.5 | 2.94e-05 |
| GOCC:0015629 | Actin cytoskeleton | 11 of 363 | 0.49 | 0.0411 |
| GOCC:0043232 | Intracellular non-membrane-bounded organelle | 99 of 3309 | 0.48 | 1.33e-24 |
| GOCC:0070062 | Extracellular exosome | 12 of 428 | 0.46 | 0.0457 |
| GOCC:0005829 | Cytosol | 76 of 3054 | 0.4 | 4.51e-13 |
| GOCC:0031981 | Nuclear lumen | 44 of 1850 | 0.39 | 3.54e-06 |
| GOCC:0030054 | Cell junction | 25 of 1053 | 0.38 | 0.0027 |
| GOCC:0005654 | Nucleoplasm | 26 of 1146 | 0.36 | 0.0036 |
| GOCC:0070013 | Intracellular organelle lumen | 56 of 2902 | 0.29 | 2.94e-05 |
| GOCC:1902494 | Catalytic complex | 33 of 1710 | 0.29 | 0.0066 |
| GOCC:0005634 | Nucleus | 88 of 4787 | 0.27 | 2.30e-08 |
| GOCC:0032991 | Protein-containing complex | 93 of 5325 | 0.25 | 6.39e-08 |
| GOCC:0005622 | Intracellular | 181 of 11512 | 0.21 | 2.56e-25 |
| GOCC:0043226 | Organelle | 159 of 10113 | 0.21 | 1.77e-16 |
| GOCC:0043229 | Intracellular organelle | 154 of 9609 | 0.21 | 4.29e-16 |
| GOCC:0043227 | Membrane-bounded organelle | 133 of 9083 | 0.17 | 2.88e-08 |
| GOCC:0043231 | Intracellular membrane-bounded organelle | 119 of 8162 | 0.17 | 1.65e-06 |
| GOCC:0005737 | Cytoplasm | 114 of 8195 | 0.15 | 6.07e-05 |
| GOCC:0110165 | Cellular anatomical entity | 181 of 14060 | 0.12 | 1.42e-12 |

### Annotated Keywords (UniProt)

| keyword | description | count in network | strength | false discovery rate |
|---|---|---|---|---|
| KW-0995 | Kinetochore | 10 of 100 | 1.01 | 6.63e-06 |
| KW-0159 | Chromosome partition | 4 of 45 | 0.96 | 0.0348 |
| KW-0235 | DNA replication | 8 of 98 | 0.92 | 0.00036 |
| KW-0498 | Mitosis | 22 of 275 | 0.91 | 1.30e-11 |
| KW-0137 | Centromere | 11 of 138 | 0.91 | 1.14e-05 |
| KW-0132 | Cell division | 27 of 384 | 0.86 | 4.03e-13 |
| KW-0440 | LIM domain | 5 of 70 | 0.86 | 0.0238 |
| KW-0493 | Microtubule | 18 of 280 | 0.82 | 5.24e-08 |
| KW-0131 | Cell cycle | 39 of 651 | 0.79 | 6.55e-17 |
| KW-0347 | Helicase | 7 of 141 | 0.7 | 0.0187 |
| KW-0206 | Cytoskeleton | 50 of 1235 | 0.62 | 1.56e-15 |
| KW-0234 | DNA repair | 13 of 325 | 0.61 | 0.00098 |
| KW-0158 | Chromosome | 16 of 433 | 0.58 | 0.00035 |
| KW-0227 | DNA damage | 14 of 386 | 0.57 | 0.0013 |
| KW-0009 | Actin-binding | 9 of 271 | 0.53 | 0.0440 |
| KW-0007 | Acetylation | 99 of 3362 | 0.48 | 1.36e-24 |
| KW-1017 | Isopeptide bond | 49 of 1717 | 0.46 | 6.89e-10 |
| KW-0832 | Ubl conjugation | 66 of 2399 | 0.45 | 2.99e-13 |
| KW-0945 | Host-virus interaction | 14 of 540 | 0.42 | 0.0297 |
| KW-0175 | Coiled coil | 51 of 2166 | 0.38 | 1.58e-07 |
| KW-0067 | ATP-binding | 32 of 1379 | 0.37 | 0.00026 |
| KW-0597 | Phosphoprotein | 164 of 8122 | 0.31 | 6.05e-33 |
| KW-0963 | Cytoplasm | 103 of 5095 | 0.31 | 8.42e-14 |
| KW-0547 | Nucleotide-binding | 34 of 1776 | 0.29 | 0.0044 |
| KW-0539 | Nucleus | 85 of 5278 | 0.22 | 1.14e-05 |
| KW-0025 | Alternative splicing | 130 of 10313 | 0.11 | 0.00068 |

**Supplemental Table 13. Proteins which were upregulated by $JNK_i$ co-treatment (with GEM) compared to single GEM treatment in A375 melanoma cells.**

| Upregulated proteins after 24 hours of co-treatment with 20 nM GEM and 5 µM $JNK_i$ (compared to single GEM treatment) | | | |
|---|---|---|---|
| **Gene name for protein** | **Description** | **Fold change (log2 form)** | **t-test P value** |
| **VGF** | Neurosecretory protein VGF | 1.076204 | 3.8E-05 |
| **HMGCS1** | Hydroxymethylglutaryl-CoA synthase, cytoplasmic | 1.904062 | 4.43E-05 |
| **SREBF2** | Sterol regulatory element-binding protein 2 | 0.78664 | 0.000155 |
| **FDFT1** | Squalene synthase | 1.635764 | 0.000233 |
| **APOB** | Apolipoprotein B-100 | 1.561881 | 0.000546 |
| **HSD17B7** | 3-keto-steroid reductase/17-beta-hydroxysteroid dehydrogenase 7 | 0.926148 | 0.000569 |
| **MRTFA** | Myocardin-related transcription factor A | 0.546328 | 0.000652 |
| **H1-10** | Histone H1.10 | 1.349269 | 0.000834 |
| **CSNK1D** | Casein kinase I isoform delta | 0.338482 | 0.000887 |
| **IRS2** | Insulin receptor substrate 2 | 0.701803 | 0.000896 |
| **ACSL1** | Long-chain-fatty-acid--CoA ligase 1 | 0.423161 | 0.000966 |
| **CBLB** | E3 ubiquitin-protein ligase CBL-B | 0.223528 | 0.001503 |
| **GPBP1** | Vasculin | 0.618754 | 0.001769 |
| **ATF7** | Cyclic AMP-dependent transcription factor ATF-7 | 1.123079 | 0.0018 |
| **LGALS8** | Galectin-8 | 0.538766 | 0.001887 |
| **SUPT6H** | Transcription elongation factor SPT6 | 0.317942 | 0.002013 |
| **HLA-E** | HLA class I histocompatibility antigen, alpha chain E | 0.765484 | 0.002125 |
| **SEZ6L2** | Seizure 6-like protein 2 | 0.25854 | 0.002247 |
| **KYNU** | Kynureninase | 0.539452 | 0.002274 |
| **FTH1** | Ferritin heavy chain | 0.593421 | 0.002307 |
| **ACSS2** | Acetyl-coenzyme A synthetase, cytoplasmic | 0.84308 | 0.002329 |

| | | | |
|---|---|---|---|
| **CDH13** | Cadherin-13 | 1.026841 | 0.002434 |
| **APOE** | Apolipoprotein E | 1.113001 | 0.002453 |
| **PBXIP1** | Pre-B-cell leukemia transcription factor-interacting protein 1 | 0.742713 | 0.002462 |
| **TGOLN2** | Trans-Golgi network integral membrane protein 2 | 0.959824 | 0.002587 |
| **ZKSCAN4** | Zinc finger protein with KRAB and SCAN domains 4 | 0.70483 | 0.002608 |
| **RPL35A** | 60S ribosomal protein L35a | 0.410125 | 0.002648 |
| **FAM234A** | Protein FAM234A | 1.034991 | 0.002667 |
| **MAF** | Transcription factor Maf | 1.22661 | 0.002781 |
| **SLC6A6** | Sodium- and chloride-dependent taurine transporter | 1.210694 | 0.002821 |
| **CYP51A1** | Lanosterol 14-alpha demethylase | 0.630215 | 0.003048 |
| **ZRANB3** | DNA annealing helicase and endonuclease ZRANB3 | 1.051233 | 0.003106 |
| **IFT74** | Intraflagellar transport protein 74 homolog | 0.426132 | 0.003237 |
| **VAT1** | Synaptic vesicle membrane protein VAT-1 homolog | 0.248862 | 0.003324 |
| **LSS** | Lanosterol synthase | 0.720207 | 0.003422 |
| **MRPL27** | 39S ribosomal protein L27, mitochondrial | 0.394434 | 0.003433 |
| **SERF2** | Small EDRK-rich factor 2 | 0.722597 | 0.003444 |
| **SFRP1** | Secreted frizzled-related protein 1 | 0.952052 | 0.003483 |
| **B4GALT5** | Beta-1,4-galactosyltransferase 5 | 0.579079 | 0.003515 |
| **ITFG1** | T-cell immunomodulatory protein | 0.552719 | 0.003552 |
| **DNAJB9** | DnaJ homolog subfamily B member 9 | 0.624891 | 0.003722 |
| **SLC39A10** | Zinc transporter ZIP10 | 0.510029 | 0.003757 |
| **NCOA3** | Nuclear receptor coactivator 3 | 0.342713 | 0.003784 |
| **SPRY2** | Protein sprouty homolog 2 | 0.464587 | 0.00386 |

| | | | |
|---|---|---|---|
| **MID1IP1** | Mid1-interacting protein 1 | 0.460689 | 0.003892 |
| **CD109** | CD109 antigen | 0.498302 | 0.003941 |
| **GRN** | Progranulin | 0.815831 | 0.004074 |
| **RETREG1** | Reticulophagy regulator 1 | 0.951965 | 0.004276 |
| **N4BP2L2** | NEDD4-binding protein 2-like 2 | 0.59233 | 0.004313 |
| **ADGRG1** | Adhesion G-protein coupled receptor G1 | 0.668457 | 0.00454 |
| **SAPCD2** | Suppressor APC domain-containing protein 2 | 0.589782 | 0.004603 |
| **MPG** | DNA-3-methyladenine glycosylase | 0.503491 | 0.004671 |
| **SNCA** | Alpha-synuclein | 0.704363 | 0.004843 |
| **MPLKIP** | M-phase-specific PLK1-interacting protein | 0.442562 | 0.004906 |
| **HABP4** | Intracellular hyaluronan-binding protein 4 | 0.393632 | 0.004923 |
| **GEMIN2** | Gem-associated protein 2 | 0.493845 | 0.004973 |
| **AAGAB** | Alpha- and gamma-adaptin-binding protein p34 | 1.361742 | 0.00503 |
| **EID2** | EP300-interacting inhibitor of differentiation 2 | 0.435428 | 0.005031 |
| **NDUFAF4** | NADH dehydrogenase [ubiquinone] 1 alpha subcomplex assembly factor 4 | 0.27505 | 0.005347 |
| **CDR2L** | Cerebellar degeneration-related protein 2-like | 0.457765 | 0.005373 |
| **KDM5B** | Lysine-specific demethylase 5B | 0.733259 | 0.005379 |
| **APOC2** | Apolipoprotein C-II | 1.614503 | 0.005522 |
| **FOXD3** | Forkhead box protein D3 | 0.773875 | 0.005569 |
| **ST8SIA4** | CMP-N-acetylneuraminate-poly-alpha-2,8-sialyltransferase | 1.583714 | 0.005584 |

| | | | |
|---|---|---|---|
| **EID1** | EP300-interacting inhibitor of differentiation 1 | 0.853687 | 0.005594 |
| **CKS2** | Cyclin-dependent kinases regulatory subunit 2 | 0.70465 | 0.005829 |
| **GRAMD1B** | Protein Aster-B | 0.585055 | 0.005843 |
| **DHCR24** | Delta(24)-sterol reductase | 0.556278 | 0.005874 |
| **ERMP1** | Endoplasmic reticulum metallopeptidase 1 | 0.632951 | 0.006174 |
| **MVP** | Major vault protein | 0.511428 | 0.006199 |
| **ZNF608** | Zinc finger protein 608 | 0.442066 | 0.006256 |
| **DNAJC1** | DnaJ homolog subfamily C member 1 | 0.303546 | 0.00636 |
| **EML3** | Echinoderm microtubule-associated protein-like 3 | 0.226787 | 0.006421 |
| **TUT1** | Speckle targeted PIP5K1A-regulated poly(A) polymerase | 0.446787 | 0.006656 |
| **NIPBL** | Nipped-B-like protein | 0.194847 | 0.006676 |
| **IL13RA1** | Interleukin-13 receptor subunit alpha-1 | 1.056727 | 0.006689 |
| **VGLL4** | Transcription cofactor vestigial-like protein 4 | 0.467464 | 0.006714 |
| **DAG1** | Dystroglycan 1 | 0.696923 | 0.006745 |
| **TRMT1L** | TRMT1-like protein | 0.393929 | 0.00684 |
| **WASF2** | Actin-binding protein WASF2 | 0.396543 | 0.0069 |
| **DUSP4** | Dual specificity protein phosphatase 4 | 0.685071 | 0.006931 |
| **APOC3** | Apolipoprotein C-III | 1.403981 | 0.006945 |
| **AMOTL1** | Angiomotin-like protein 1 | 0.240062 | 0.007094 |
| **RRP36** | Ribosomal RNA processing protein 36 homolog | 0.368129 | 0.007095 |
| **PDE3A** | cGMP-inhibited 3',5'-cyclic phosphodiesterase 3A | 0.893008 | 0.007103 |
| **ITIH2** | Inter-alpha-trypsin inhibitor heavy chain H2 | 1.005178 | 0.007336 |
| **H2BC18** | Histone H2B type 2-F | 0.988771 | 0.00734 |
| **ANKRD16** | Ankyrin repeat domain-containing protein 16 | 0.409963 | 0.007468 |

| | | | |
|---|---|---|---|
| **TMX2** | Thioredoxin-related transmembrane protein 2 | 0.773985 | 0.007923 |
| **MCFD2** | Multiple coagulation factor deficiency protein 2 | 0.503236 | 0.007987 |
| **NDUFS5** | NADH dehydrogenase [ubiquinone] iron-sulfur protein 5 | 0.559474 | 0.008104 |
| **MAP2K6** | Dual specificity mitogen-activated protein kinase kinase 6 | 0.438644 | 0.008368 |
| **ELF2** | ETS-related transcription factor Elf-2 | 0.880941 | 0.008421 |
| **NHSL1** | NHS-like protein 1 | 0.458558 | 0.008513 |
| **LSM1** | U6 snRNA-associated Sm-like protein LSm1 | 0.569069 | 0.008581 |
| **ADAM10** | Disintegrin and metalloproteinase domain-containing protein 10 | 0.247531 | 0.008915 |
| **MVD** | Diphosphomevalonate decarboxylase | 0.894227 | 0.008969 |
| **SEMA3C** | Semaphorin-3C | 0.531036 | 0.008986 |
| **NUCB1** | Nucleobindin-1 | 0.575475 | 0.009105 |
| **EPS8** | Epidermal growth factor receptor kinase substrate 8 | 0.154915 | 0.009277 |
| **HSPA13** | Heat shock 70 kDa protein 13 | 0.458041 | 0.009605 |
| **MGRN1** | E3 ubiquitin-protein ligase MGRN1 | 0.17567 | 0.009636 |
| **SDC4** | Syndecan-4 | 0.891376 | 0.009671 |
| **MON2** | Protein MON2 homolog | 0.240008 | 0.009723 |
| **TBL1XR1** | F-box-like/WD repeat-containing protein TBL1XR1 | 0.190625 | 0.009925 |

**Supplemental Table 14. Proteins which were downregulated by JNK$_i$ co-treatment (with GEM) compared to single GEM treatment in A375 melanoma cells.**

| Downregulated proteins after 24 hours of co-treatment with 20 nM GEM and 5 µM JNK$_i$ (compared to single GEM treatment) | | | |
|---|---|---|---|
| **Gene name for protein** | **Description** | **Fold change (log2 form)** | **t-test P value** |
| **TK1** | Thymidine kinase, cytosolic | -1.82128 | 1.84E-06 |
| **CCP110** | Centriolar coiled-coil protein of 110 kDa | -0.88667 | 2.86E-05 |
| **SMTN** | Smoothelin | -2.19613 | 3.26E-05 |
| **CLSPN** | Claspin | -1.46465 | 5.4E-05 |
| **ANLN** | Anillin | -1.80827 | 5.58E-05 |
| **TOP2A** | DNA topoisomerase 2-alpha | -0.94888 | 9.25E-05 |
| **RRM2** | Ribonucleoside-diphosphate reductase subunit M2 | -1.9824 | 9.63E-05 |
| **CENPF** | Centromere protein F | -0.69282 | 0.000101 |
| **KRT8** | Keratin, type II cytoskeletal 8 | -0.86501 | 0.000109 |
| **PSMD14** | 26S proteasome non-ATPase regulatory subunit 14 | -0.15159 | 0.000147 |
| **LAP3** | Cytosol aminopeptidase | -0.18471 | 0.000175 |
| **RAD18** | E3 ubiquitin-protein ligase RAD18 | -0.52672 | 0.000187 |
| **SPOUT1** | Putative methyltransferase C9orf114 | -0.30733 | 0.000199 |
| **ERCC6L** | DNA excision repair protein ERCC-6-like | -0.44313 | 0.000208 |
| **BLM** | RecQ-like DNA helicase BLM | -1.38949 | 0.000216 |
| **KIF11** | Kinesin-like protein KIF11 | -1.36297 | 0.000225 |
| **KRT18** | Keratin, type I cytoskeletal 18 | -0.76407 | 0.000226 |
| **DDB2** | DNA damage-binding protein 2 | -0.60393 | 0.000306 |
| **SYNPO** | Synaptopodin | -0.82166 | 0.00033 |
| **JUN** | Transcription factor Jun | -1.74694 | 0.000343 |
| **LIG1** | DNA ligase 1 | -0.46564 | 0.000386 |
| **CAV1** | Caveolin-1 | -1.0398 | 0.000393 |
| **KIF20A** | Kinesin-like protein KIF20A | -0.29367 | 0.000411 |

| | | | |
|---|---|---|---|
| **SGO1** | Shugoshin 1 | -1.36462 | 0.000427 |
| **PHLDB1** | Pleckstrin homology-like domain family B member 1 | -0.53259 | 0.000439 |
| **MTPAP** | Poly(A) RNA polymerase, mitochondrial | -0.25858 | 0.000442 |
| **CKAP2** | Cytoskeleton-associated protein 2 | -1.48414 | 0.000461 |
| **KIFC1** | Kinesin-like protein KIFC1 | -0.88221 | 0.000462 |
| **INCENP** | Inner centromere protein | -0.94315 | 0.00053 |
| **RACGAP1** | Rac GTPase-activating protein 1 | -0.59728 | 0.000542 |
| **GPRIN1** | G protein-regulated inducer of neurite outgrowth 1 | -0.81161 | 0.000546 |
| **MICALL1** | MICAL-like protein 1 | -0.59601 | 0.000587 |
| **UAP1** | UDP-N-acetylhexosamine pyrophosphorylase | -0.4606 | 0.000588 |
| **RAI14** | Ankycorbin | -0.35196 | 0.000609 |
| **SPART** | Spartin | -0.2785 | 0.000626 |
| **TULP3** | Tubby-related protein 3 | -0.55411 | 0.000727 |
| **RECQL** | ATP-dependent DNA helicase Q1 | -0.25219 | 0.000752 |
| **HADH** | Hydroxyacyl-coenzyme A dehydrogenase, mitochondrial | -0.28188 | 0.000801 |
| **PRC1** | Protein regulator of cytokinesis 1 | -0.90329 | 0.000809 |
| **AFAP1L2** | Actin filament-associated protein 1-like 2 | -0.73943 | 0.00081 |
| **EPB41L1** | Band 4.1-like protein 1 | -0.60983 | 0.000815 |
| **DPYSL3** | Dihydropyrimidinase-related protein 3 | -0.40922 | 0.000827 |
| **POLR1C** | DNA-directed RNA polymerases I and III subunit RPAC1 | -0.16365 | 0.000836 |
| **CGN** | Cingulin | -0.43401 | 0.000846 |
| **STK10** | Serine/threonine-protein kinase 10 | -0.45042 | 0.000857 |
| **TACC3** | Transforming acidic coiled-coil-containing protein 3 | -1.72011 | 0.000888 |
| **GPSM3** | G-protein-signaling modulator 3 | -0.2675 | 0.001007 |

| | | | |
|---|---|---|---|
| **PPME1** | Protein phosphatase methylesterase 1 | -0.28318 | 0.001065 |
| **DDX3X** | ATP-dependent RNA helicase DDX3X | -0.22434 | 0.00108 |
| **KIF22** | Kinesin-like protein KIF22 | -0.79877 | 0.001114 |
| **SNX1** | Sorting nexin-1 | -0.25249 | 0.001192 |
| **ECM1** | Extracellular matrix protein 1 | -0.79251 | 0.001199 |
| **LIMCH1** | LIM and calponin homology domains-containing protein 1 | -0.61897 | 0.001228 |
| **FOSB** | Protein FosB | -1.30022 | 0.001232 |
| **EXOSC1** | Exosome complex component CSL4 | -0.28963 | 0.001295 |
| **PARD3** | Partitioning defective 3 homolog | -0.30373 | 0.001332 |
| **RBMS2** | RNA-binding motif, single-stranded-interacting protein 2 | -0.71618 | 0.001458 |
| **SCRIB** | Protein scribble homolog | -0.34913 | 0.001533 |
| **RPL10** | 60S ribosomal protein L10 | -0.32942 | 0.001536 |
| **RBBP5** | Retinoblastoma-binding protein 5 | -0.42443 | 0.00158 |
| **CDK5RAP2** | CDK5 regulatory subunit-associated protein 2 | -0.55681 | 0.001581 |
| **TOE1** | Target of EGR1 protein 1 | -0.72607 | 0.001713 |
| **SH3BGRL2** | SH3 domain-binding glutamic acid-rich-like protein 2 | -0.38407 | 0.001748 |
| **YWHAH** | 14-3-3 protein eta | -0.50751 | 0.001767 |
| **PNP** | Purine nucleoside phosphorylase | -0.22033 | 0.001781 |
| **SPICE1** | Spindle and centriole-associated protein 1 | -0.49299 | 0.001785 |
| **MPRIP** | Myosin phosphatase Rho-interacting protein | -0.34064 | 0.001859 |
| **TAGLN3** | Transgelin-3 | -1.12891 | 0.001869 |
| **CBX2** | Chromobox protein homolog 2 | -0.38799 | 0.001889 |
| **KIF23** | Kinesin-like protein KIF23 | -0.48452 | 0.002042 |
| **DIDO1** | Death-inducer obliterator 1 | -0.45747 | 0.002058 |

| | | | |
|---|---|---|---|
| **SFN** | 14-3-3 protein sigma | -0.88485 | 0.002071 |
| **TRIP6** | Thyroid receptor-interacting protein 6 | -0.74867 | 0.002106 |
| **MMP14** | Matrix metalloproteinase-14 | -0.50706 | 0.002133 |
| **CC2D1A** | Coiled-coil and C2 domain-containing protein 1A | -0.14756 | 0.002252 |
| **KIAA1217** | Sickle tail protein homolog | -0.4297 | 0.002256 |
| **PPFIBP1** | Liprin-beta-1 | -0.24214 | 0.002294 |
| **PCLAF** | PCNA-associated factor | -1.45784 | 0.002312 |
| **DSCC1** | Sister chromatid cohesion protein DCC1 | -0.67415 | 0.002353 |
| **ACBD5** | Acyl-CoA-binding domain-containing protein 5 | -0.46416 | 0.002419 |
| **BAZ1B** | Tyrosine-protein kinase BAZ1B | -0.42266 | 0.002472 |
| **TGM2** | Protein-glutamine gamma-glutamyltransferase 2 | -0.42558 | 0.002473 |
| **CDCA5** | Sororin | -1.3623 | 0.002601 |
| **ANXA2** | Annexin A2 | -0.4298 | 0.00264 |
| **HSPH1** | Heat shock protein 105 kDa | -0.27913 | 0.002661 |
| **KIF2C** | Kinesin-like protein KIF2C | -1.11756 | 0.002677 |
| **MYBL2** | Myb-related protein B | -1.03676 | 0.002678 |
| **WNK1** | Serine/threonine-protein kinase WNK1 | -0.17436 | 0.0028 |
| **KLHDC4** | Kelch domain-containing protein 4 | -0.29965 | 0.002832 |
| **CASP8** | Caspase-8 | -0.29281 | 0.002849 |
| **PLAT** | Tissue-type plasminogen activator | -0.86225 | 0.00285 |
| **CCNB1** | G2/mitotic-specific cyclin-B1 | -1.58392 | 0.002865 |
| **NCKAP5L** | Nck-associated protein 5-like | -0.90701 | 0.003001 |
| **DLGAP5** | Disks large-associated protein 5 | -0.5274 | 0.003042 |
| **RAB11FIP1** | Rab11 family-interacting protein 1 | -0.36414 | 0.003086 |
| **VPS28** | Vacuolar protein sorting-associated protein 28 homolog | -0.40365 | 0.00315 |

| | | | |
|---|---|---|---|
| **DHFR** | Dihydrofolate reductase | -0.50568 | 0.003181 |
| **NSD2** | Histone-lysine N-methyltransferase NSD2 | -0.33049 | 0.00335 |
| **DNAJB4** | DnaJ homolog subfamily B member 4 | -0.54973 | 0.003418 |
| **NES** | Nestin | -0.60709 | 0.003507 |
| **TPX2** | Targeting protein for Xklp2 | -1.06717 | 0.003549 |
| **RPA3** | Replication protein A 14 kDa subunit | -0.33312 | 0.003556 |
| **SKA2** | Spindle and kinetochore-associated protein 2 | -0.67486 | 0.003644 |
| **ADAR** | Double-stranded RNA-specific adenosine deaminase | -0.24791 | 0.003648 |
| **PRMT1** | Protein arginine N-methyltransferase 1 | -0.23326 | 0.003677 |
| **DNLZ** | DNL-type zinc finger protein | -0.35441 | 0.003747 |
| **TP53BP1** | TP53-binding protein 1 | -0.34282 | 0.003842 |
| **ELL** | RNA polymerase II elongation factor ELL | -0.31559 | 0.00387 |
| **CKB** | Creatine kinase B-type | -0.76866 | 0.00399 |
| **MPHOSPH8** | M-phase phosphoprotein 8 | -0.38426 | 0.004025 |
| **CSNK2A1** | Casein kinase II subunit alpha | -0.23513 | 0.004107 |
| **RASAL2** | Ras GTPase-activating protein nGAP | -0.38389 | 0.004172 |
| **DDX5** | Probable ATP-dependent RNA helicase DDX5 | -0.28642 | 0.004295 |
| **LIMA1** | LIM domain and actin-binding protein 1 | -0.55185 | 0.004307 |
| **ALMS1** | Centrosome-associated protein ALMS1 | -0.22884 | 0.004549 |
| **SKA3** | Spindle and kinetochore-associated protein 3 | -0.95862 | 0.004727 |
| **EIF4G1** | Eukaryotic translation initiation factor 4 gamma 1 | -0.28975 | 0.004865 |
| **TJP2** | Tight junction protein ZO-2 | -0.45612 | 0.004891 |
| **TOX4** | TOX high mobility group box family member 4 | -0.31327 | 0.005139 |

| | | | |
|---|---|---|---|
| **CTNND1** | Catenin delta-1 | -0.22068 | 0.005172 |
| **DNAJC6** | Putative tyrosine-protein phosphatase auxilin | -0.2371 | 0.005183 |
| **BAD** | Bcl2-associated agonist of cell death | -0.47067 | 0.005262 |
| **GMNN** | Geminin | -1.1551 | 0.005429 |
| **GPSM1** | G-protein-signaling modulator 1 | -0.4301 | 0.005435 |
| **PYCR2** | Pyrroline-5-carboxylate reductase 2 | -0.29879 | 0.005499 |
| **CEMIP2** | Cell surface hyaluronidase | -0.58174 | 0.005622 |
| **KIF14** | Kinesin-like protein KIF14 | -0.25253 | 0.00565 |
| **UBE2S** | Ubiquitin-conjugating enzyme E2 S | -0.80305 | 0.005778 |
| **ZFYVE16** | Zinc finger FYVE domain-containing protein 16 | -0.12913 | 0.005832 |
| **CLPP** | ATP-dependent Clp protease proteolytic subunit, mitochondrial | -0.23682 | 0.005926 |
| **HMOX2** | Heme oxygenase 2 | -0.45149 | 0.005942 |
| **GGCT** | Gamma-glutamylcyclotransferase | -0.44977 | 0.005965 |
| **RRM1** | Ribonucleoside-diphosphate reductase large subunit | -0.50731 | 0.005981 |
| **ADD2** | Beta-adducin | -0.25186 | 0.006035 |
| **POLD3** | DNA polymerase delta subunit 3 | -0.59734 | 0.006068 |
| **ACOT7** | Cytosolic acyl coenzyme A thioester hydrolase | -0.18099 | 0.00613 |
| **FAM98B** | Protein FAM98B | -0.41672 | 0.006134 |
| **GSPT1** | Eukaryotic peptide chain release factor GTP-binding subunit ERF3A | -0.40748 | 0.006287 |
| **CNN2** | Calponin-2 | -0.63909 | 0.006362 |
| **CDK2** | Cyclin-dependent kinase 2 | -0.45948 | 0.006615 |
| **CALB2** | Calretinin | -0.50327 | 0.00694 |
| **MAP1S** | Microtubule-associated protein 1S | -0.24913 | 0.007028 |
| **NUP214** | Nuclear pore complex protein Nup214 | -0.26916 | 0.007147 |
| **KPNA2** | Importin subunit alpha-1 | -0.58475 | 0.007158 |

| | | | |
|---|---|---|---|
| **MKI67** | Proliferation marker protein Ki-67 | -0.71878 | 0.007177 |
| **TRIM28** | Transcription intermediary factor 1-beta | -0.38276 | 0.007226 |
| **PPM1F** | Protein phosphatase 1F | -0.29415 | 0.007258 |
| **UCK2** | Uridine-cytidine kinase 2 | -0.40873 | 0.007288 |
| **PCM1** | Pericentriolar material 1 protein | -0.24618 | 0.007335 |
| **UBN1** | Ubinuclein-1 | -0.29158 | 0.007374 |
| **PDLIM2** | PDZ and LIM domain protein 2 | -0.58838 | 0.007385 |
| **CSPP1** | Centrosome and spindle pole-associated protein 1 | -0.6448 | 0.00742 |
| **FSCN1** | Fascin | -0.43478 | 0.007428 |
| **ERLIN2** | Erlin-2 | -0.27513 | 0.007548 |
| **ANPEP** | Aminopeptidase N | -0.44891 | 0.007555 |
| **TARS1** | Threonine--tRNA ligase 1, cytoplasmic | -0.14973 | 0.007604 |
| **TTK** | Dual specificity protein kinase TTK | -0.45232 | 0.007719 |
| **PITPNB** | Phosphatidylinositol transfer protein beta isoform | -0.22495 | 0.007739 |
| **STRAP** | Serine-threonine kinase receptor-associated protein | -0.30358 | 0.007794 |
| **DUS3L** | tRNA-dihydrouridine(47) synthase [NAD(P)(+)]-like | -0.20995 | 0.007812 |
| **SMARCC2** | SWI/SNF complex subunit SMARCC2 | -0.12862 | 0.007871 |
| **RABL6** | Rab-like protein 6 | -0.30159 | 0.008017 |
| **SPATS2L** | SPATS2-like protein | -0.43053 | 0.008061 |
| **SEC63** | Translocation protein SEC63 homolog | -0.32367 | 0.008097 |
| **CDCA8** | Borealin | -0.75198 | 0.008107 |
| **TRIOBP** | TRIO and F-actin-binding protein | -0.37629 | 0.00816 |
| **TSC22D2** | TSC22 domain family protein 2 | -0.23431 | 0.008214 |
| **CLCC1** | Chloride channel CLIC-like protein 1 | -0.33194 | 0.008217 |
| **ANXA1** | Annexin A1 | -0.39299 | 0.008291 |

| | | | |
|---|---|---|---|
| **ZNF598** | E3 ubiquitin-protein ligase ZNF598 | -0.29186 | 0.0083 |
| **RPL28** | 60S ribosomal protein L28 | -0.65084 | 0.00834 |
| **HPF1** | Histone PARylation factor 1 | -0.24543 | 0.008541 |
| **KDM2A** | Lysine-specific demethylase 2A | -0.39738 | 0.008591 |
| **CHAF1B** | Chromatin assembly factor 1 subunit B | -0.70203 | 0.008614 |
| **SPDL1** | Protein Spindly | -0.37771 | 0.008623 |
| **POLR2H** | DNA-directed RNA polymerases I, II, and III subunit RPABC3 | -0.27039 | 0.00866 |
| **EPHA2** | Ephrin type-A receptor 2 | -0.52844 | 0.008718 |
| **DDX46** | Probable ATP-dependent RNA helicase DDX46 | -0.30318 | 0.008743 |
| **VSIR** | V-type immunoglobulin domain-containing suppressor of T-cell activation | -0.87863 | 0.008822 |
| **BCAR1** | Breast cancer anti-estrogen resistance protein 1 | -0.56308 | 0.009048 |
| **GNB4** | Guanine nucleotide-binding protein subunit beta-4 | -0.51743 | 0.009124 |
| **VTA1** | Vacuolar protein sorting-associated protein VTA1 homolog | -0.51884 | 0.009243 |
| **DDX21** | Nucleolar RNA helicase 2 | -0.41741 | 0.00927 |
| **CDC5L** | Cell division cycle 5-like protein | -0.29532 | 0.009411 |
| **RPL21** | 60S ribosomal protein L21 | -0.20701 | 0.009478 |
| **WDR70** | WD repeat-containing protein 70 | -0.2001 | 0.009577 |
| **MNX1** | Motor neuron and pancreas homeobox protein 1 | -0.47617 | 0.0096 |
| **CDKN2AIP** | CDKN2A-interacting protein | -0.42883 | 0.00965 |
| **NKRF** | NF-kappa-B-repressing factor | -0.32741 | 0.00969 |
| **MRPL47** | 39S ribosomal protein L47, mitochondrial | -0.19809 | 0.009801 |

| | | | |
|---|---|---|---|
| **DCUN1D3** | DCN1-like protein 3 | -1.05152 | 0.00993 |
| **CEP131** | Centrosomal protein of 131 kDa | -0.32141 | 0.009942 |
| **CDCA3** | Cell division cycle-associated protein 3 | -0.80778 | 0.00997 |
